# A druggable redox switch on SHP1 controls macrophage inflammation

**DOI:** 10.64898/2026.02.19.706790

**Authors:** Mei Ying Ng, Meredith N. Nix, Guangyan Du, Ivan Davidek, Nils Burger, Sanghee Shin, Sean Toenjes, Haruna Takeda, Megan Cheah Xin Yan, Bingsen Zhang, Haopeng Xiao, Shelley Wei, Hyuk-Soo Seo, Sirano Dhe-Paganon, Thomas E. Wales, John R. Engen, Evanna L. Mills, Jianwei Che, Tinghu Zhang, Nathanael S. Gray, Edward T. Chouchani

**Author notes:** These authors contributed equally: Mei Ying Ng, Meredith N. Nix.

## Abstract

Immunological proteins are major disease targets, yet most remain undrugged. Post-translational redox modification of cysteine residues has emerged as an important mode of immune cell regulation, particularly in macrophage cytokine responses. Here, we develop a strategy for systematic discovery and small-molecule functionalization of redox-regulated cysteines on immunological proteins. Using deep redox proteomics, we annotate 788 in vivo redox-regulated cysteines across diverse immune-relevant protein domains. We demonstrate how these sites enable cysteine-directed pharmacology through discovery of a novel cysteine activation site on the immune regulator SHP1. Targeting Cys102, we develop a highly selective covalent agonist, SCA, which binds the N-SH2 domain to relieve autoinhibition and activate SHP1. In mouse and human macrophages, SCA selectively engages SHP1 Cys102, antagonizing IRAK signaling and LPS-induced pro-inflammatory cytokine production. Together, this work identifies a druggable cysteine redox switch controlling macrophage cytokine responses and provides a compendium of redox-regulated sites for therapeutic development.

## Introduction

A wide range of immunological processes are regulated through the generation of reactive oxygen species (ROS) and related redox-active molecules^1-5^. Redox-active metabolites regulate immune cells through covalent modification of protein cysteine residues, consequently modifying protein function^2-4,6^. These redox modifications occur due to the distinct reactivity of protein cysteines compared to other amino acids, rendering them selective targets for this form of post-translational modification. A prominent example of this type of regulation is found in the context of macrophage cytokine production. Oxidation of protein cysteines by ROS regulates production of pro-inflammatory cytokines by macrophages^5-8^. Conversely, covalent modification of distinct protein cysteines by other redox active metabolites inhibits pro-inflammatory cytokine production and drives an anti-inflammatory phenotype^6,9-11^. Despite the widespread importance of protein cysteine redox regulation over immune cell function, there has been a persistent lack of rigorous information regarding the specific protein modifications that explain the molecular basis for these processes. Elucidating the cysteine targets on immunological proteins that regulate their function is fundamental for understanding the mechanistic basis for redox regulation of the immune system.

In addition, a more complete understanding of redox regulated cysteine sites on immune targets could provide a foundation for systematic therapeutic targeting of the immunological proteome. Indeed, the intrinsic reactivity of redox regulated cysteines can be harnessed for the development of covalent tool compounds and drugs^12,13^. In clinical oncology, numerous individual redox-active cysteines have been targeted therapeutically using electrophilic small molecules, aiming to inhibit protein activity or interfere with protein-protein interactions^12^. The effectiveness of these strategies has demonstrated the potential for design of electrophilic molecules to selectively modify specific protein cysteines.

In this work, we develop a strategy for discovery and small molecule functionalization of *in vivo* redox regulated cysteines on immunological proteins. Using deep redox proteomics, we annotate cysteines across diverse functional domains of 1,125 immune-relevant proteins to identify those that are redox regulated *in vivo*. We demonstrate how this redox regulated cysteine landscape can be queried with high-throughput chemical biology to manipulate the function of immunological proteins that previously were pharmacologically challenging, particularly in the development of positive allosteric modulators of protein phosphatases for which there are few successful examples. We develop a selective covalent agonist of the central immune regulator SHP1, which functions by targeting a redox regulated cysteine to relieve intrinsic auto-inhibition. This previously unknown mechanism of SHP1 activation inhibits pro-inflammatory cytokine production and inflammation in macrophages. We provide a compendium of redox regulated cysteines across the immunological proteome (provided as an online resource at https://oximmune-chouchani-lab.dfci.harvard.edu/) as a roadmap to engineer pharmacological regulators of currently undrugged immune-relevant proteins.

## Results

### Discovery of redox regulated cysteines on immunological proteins

Protein cysteine redox regulation plays a central role in immunology. To identify functional nodes through which ROS and related species exert their physiological roles, it is important to quantify the stoichiometry of reversible modification on a particular cysteine residue. Sites subject to dynamic modification, with the capacity to exhibit large absolute changes in modification state, represent a core property of redox regulated cysteines^14-17^. Indeed, proteomic studies of macrophages indicate a wide dynamic range of protein cysteine oxidation stoichiometry depending on cellular context^18,19^. Quantifying these regulated sites *in vivo* is a particular challenge for immunological proteins, which exhibit low abundance in tissues.

We recently developed Oximouse, a quantitative and comprehensive characterization of protein cysteine oxidation *in vivo*. This technology makes use of cysteine derivatization and enrichment chemistry to determine % redox modification of ∼171,000 protein cysteines sites across ten tissues in mice **(Extended Data Figure 1a)**^14^. Here, we evolved this compendium to explore the biological role of individual redox regulated protein cysteines across the immunological proteome *in vivo*, and in doing so we develop a resource called OxImmune. We defined the immunological proteome to include all proteins contained in InnateDB. InnateDB is comprised of 1,125 proteins with experimentally verified roles in the innate immune response^20^. To profile redox regulation across this immunological proteome *in vivo*, we examined the redox modification state of ∼34,000 unique cysteines in living mouse tissues **(Figure 1a)**. From this population, we focused our analysis to 13,018 cysteines on InnateDB proteins conserved between mouse and human **(Figure 1a & Supplementary Table 1)**. For each cysteine residue, OxImmune tracks the modification state across 10 tissues and two ages. It is appreciated that the local tissue environment exerts widely divergent effects on immunological proteins and processes. On this basis, we hypothesized that by tracking the natural heterogeneity of cysteine modification state inherent to different tissue environments, we could reveal cysteine sites subject to dynamic modification that depends on local milieu.

**Fig. 1.**
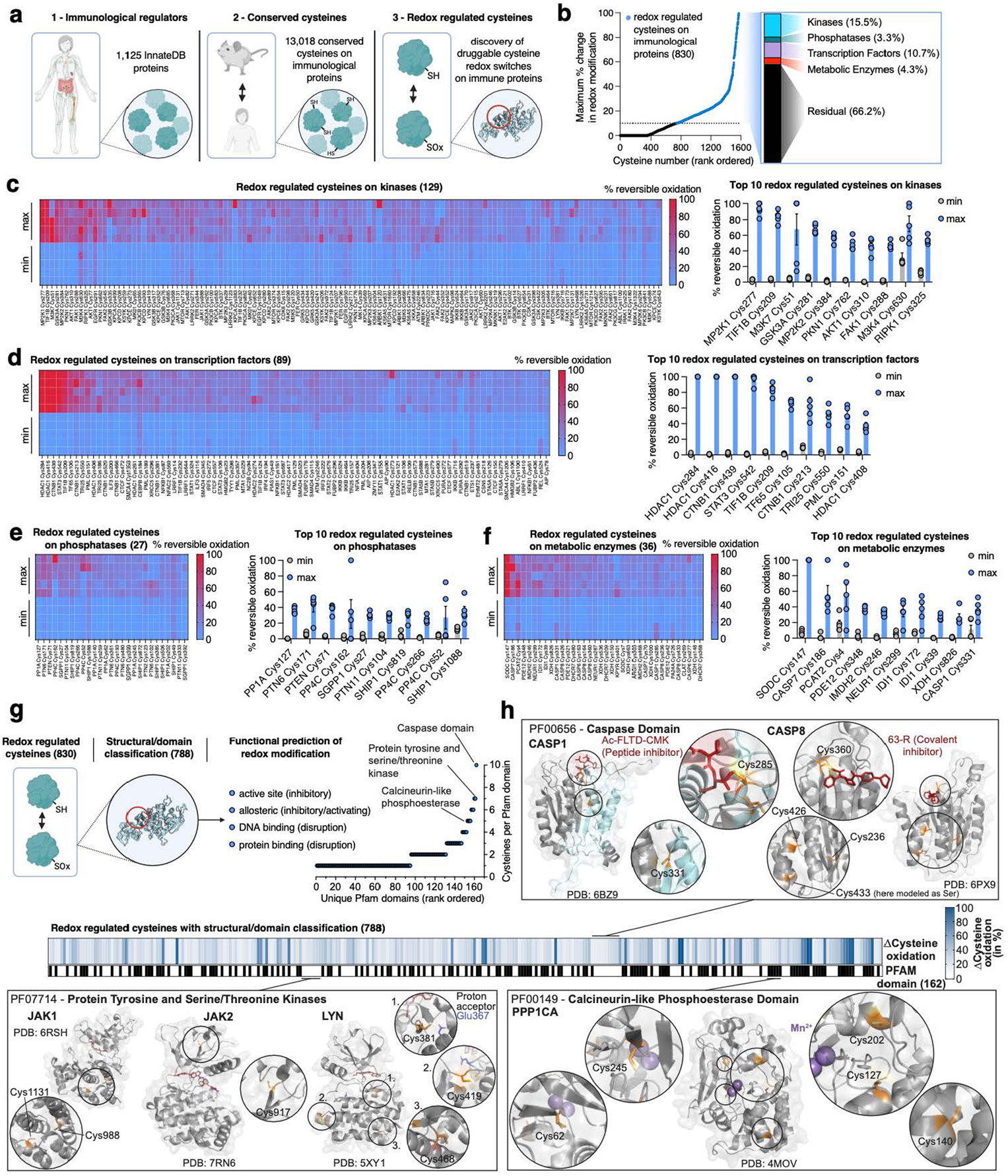
Systematic discovery of redox regulated cysteines on immunological proteins. **a**, Strategy defining redox regulated cysteines on immunological proteins. Immune proteins defined by InnateDB were aligned to identify cysteines conserved between mouse and human. Conserved cysteines were classified for their propensity to be oxidized *in vivo* and functionally and structurally annotated. Created in BioRender. Ng, M. (2025) https://BioRender.com/e74b682. **b**, Categorization of immunological cysteines subject to dynamic oxidation (>10% difference in oxidation between any two tissues/conditions in Oximouse (data from ref. 14)). **c-f**, Top 10 redox regulated cysteines on kinases **(c)**, transcription factors **(d)**, phosphatases **(e)**, and metabolic enzymes **(f)** with percent reversible oxidation across tissues/conditions with lowest and highest average oxidation in the Oximouse dataset (data from ref. 14). Min is defined as the minimum oxidation state for a site observed in Oximouse, max is defined as the maximum degree of oxidation observed in the Oximouse dataset. Data are mean ± s.e.m., *n* = 5 for each tissue. **g**, Mapping of redox regulated cysteines to experimentally determined structural and functional domains across the immunological proteome. Created in BioRender. Ng, M. (2025) https://BioRender.com/e74b682. **h**, Prominent examples of high value immune targets with redox regulated cysteines on structural or functional domains, such as active and allosteric sites.

We tracked % modification of each protein cysteine under multiple tissue conditions to define each site based on capacity for dynamic modification **(Figure 1b)**. We categorized each mapped cysteine across the immunological proteome as follows:

1. Stable redox state – a cysteine never exhibiting more than 10% difference in redox modification status between any two tissue conditions.
2. Dynamic redox state – a cysteine exhibiting more than 10% difference in redox modification status between any two tissue conditions.

From this analysis, ∼6% (830) of all mapped cysteines on immunological proteins were defined as dynamically redox regulated across tissues *in vivo* **(Figure 1b)**. Dynamic endogenous redox modification often marks regulatory sites on target proteins^14,21^, so we next classified these residues across the immunological proteome. Redox regulated cysteines were found on a wide range of protein families, including kinases, transcription factors, phosphatases, and metabolic enzymes **(Figure 1c-f)**. Across all families, ∼95% (788) of regulated cysteines mapped to established functional domains within the protein, including enzyme active sites, allosteric regulatory sites, nucleotide binding regions, and protein-protein interaction sites **(Figure 1g & Supplementary Table 1)**. We mapped redox regulated cysteines onto these domains, as well as experimentally determined structures across the immunological proteome, which enabled mechanistic prediction of the consequence of cysteine modification for each site **(Figure 1h & Supplementary Table 1)**. Among the most prominently redox regulated enzyme active sites were caspases, which are known to frequently feature redox modified active site cysteines^22-23^. Notably, several of these cysteine targets have been targeted previously by covalent caspase inhibitors^24-28^, underpinning the potential of targeting this class of regulatory cysteines. Likewise, protein tyrosine and serine/threonine kinases are known to be redox regulated and exhibit important regulatory action on immune processes^29-32^. Many members of this enzyme class contain cysteines subject to dynamic oxidation *in vivo*, both in active and allosteric sites **(Figure 1h & Supplementary Table 1)**. This was also found to apply to phosphatases, exemplified by the catalytic domain of protein phosphatase 1 **(Figure 1h & Supplementary Table 1)**^33-35^. Surveying the major protein families described above revealed that phosphatases were well represented, underscoring their critical role in major immune processes including dephosphorylation of proteins in the toll-like receptor (TLR), nuclear factor kappa-light-chain-enhancer of activated B cells (NF-κB), and stimulator of interferon genes (STING) inflammatory pathways^36,37^. We found that the top 10 highlighted redox-regulated cysteines in this family reported on established mechanisms of phosphatase redox control **(Figure 1e)**. Specifically, PP1A (PPP1CA) Cys127 can undergo inhibitory dimerization via disulfide formation between Cys127 and Cys39^33^, while oxidation of PTEN Cys71 can inhibit phosphatase activity by disulfide formation with the catalytic Cys124^38,39^. In addition, PP4C Cys266 corresponds to Cys269 in a highly conserved region of the closely related phosphatase PP2A, which is covalently modified by the inhibitor phoslactomycin A (PLMA), underpinning the relevance of this residue^40^. Beyond these examples, many other protein classes such as proteases, and central components of immune relevant signaling cascades, contain dynamically oxidized cysteines **(Supplementary Table 1)**. Together, structural and domain annotation of the redox regulated immunological proteome identified 788 instances of regulatory cysteines on established functional domains of major immune regulatory proteins **(Figure 1h)**. We provide this dataset, including target class and domain and structural annotations, via an online interface, which we use as a basis to better understand the biological role of redox regulation of immunological proteins (https://oximmune-chouchani-lab.dfci.harvard.edu/).

### Chemical manipulation of cysteines on immunological proteins

By uncovering hundreds of protein cysteines that are subject to oxidative modification *in vivo* **(Figure 1),** we reasoned that this provides an opportunity to leverage these same sites for therapeutic manipulation. The high nucleophilicity and redox activity that render cysteines amenable to dynamic oxidative modification allows for selective targeting by small molecules containing an appropriately oriented electrophilic warhead. It is now well appreciated that solvent-accessible, nucleophilic cysteine thiolates can be selectively targeted by small molecule electrophiles^13,41,42^. In principle, if such cysteine residues are localized to functional domains within a disease-relevant protein, such chemical targeting may be therapeutically useful. Based on our structural and domain annotation of the redox regulated immunological proteome, we explored whether redox regulated cysteines identified in OxImmune could be exploited for pharmacological targeting of central immunological processes.

We focused our attention to protein classes that have been challenging to target pharmacologically. A prominent example of this is found in protein phosphatases, which are central to immune cell function, and for which positive allosteric modulators have been extremely challenging to develop. This is exemplified by the SH2 domain containing protein tyrosine phosphatase-1 (SHP1) **(Figure 2a)**. SHP1 is a core inhibitor of autoimmunity^43^. SHP1 activation is central to inhibition of macrophage, monocyte, and T cell-driven inflammation^43,44^. Mice deficient for SHP1 suffer from microbe-driven inflammation and spontaneous autoimmunity^45^. T cells from patients with rheumatoid arthritis show tonic inhibition of SHP1^46^, while active SHP1 appears to be required for the anti-inflammatory action of IFNβ in multiple sclerosis^18,19^. Moreover, transgenic SHP1 overexpression protects mice from inflammatory arthritis without noticeable adverse effects^47^. In the context of a pro-inflammatory stimulus, activated SHP1 directly binds to interleukin-1 receptor-associated kinase 1 (IRAK1) and antagonizes IRAK1-mediated NF-κB signaling^48,49^. SHP1 activity is known to be suppressed by auto-inhibition, whereby the N-SH2 domain blocks access of substrate to the active site^45^.

**Fig. 2.**
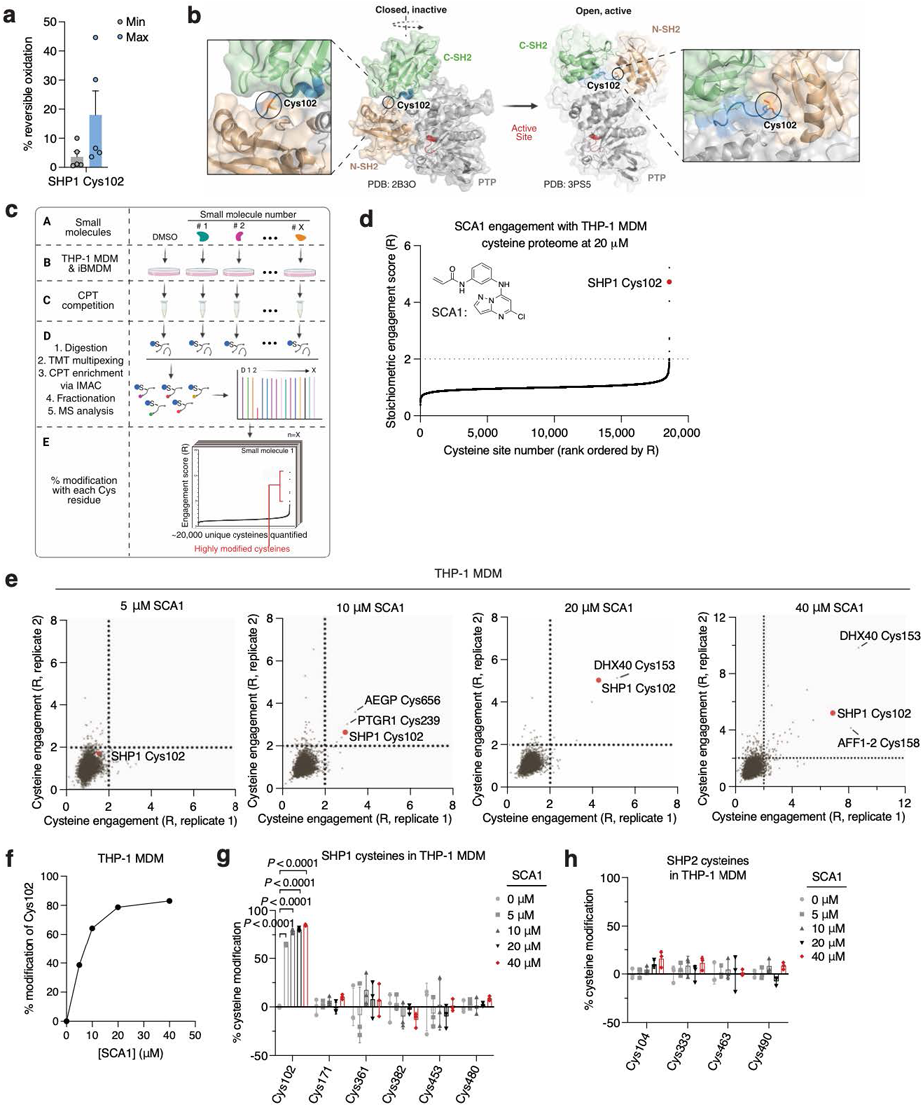
Deep chemoproteomics functionalizes the redox-sensitive Cys102 on SHP1. **a**, Oximouse redox proteomics minimum (Min) and maximum (Max) percent reversible oxidation of SHP1 Cys102 across all tissues studied (*n* = 5 for each tissue). See Oximouse data from ref. 14. **b**, Crystal structure of human SHP1 highlighting Cys102 (orange) in the closed, inactive form (left, PDB ID: 2B3O), residing within an N-SH2 proximal pocket involved in rearrangement to the open, active form (right, PDB ID: 3PS5). N-SH2: N-terminal SH2 domain (green); C-SH2: C-terminal SH2 domain (brown); PTP: protein tyrosine phosphatase domain (grey). Disordered regions are shown in blue; active site pocket is represented in red. Created in BioRender. Ng, M. (2025) https://BioRender.com/sa5mdvt. **c**, Schematic of CPT-MS workflow for quantification of cysteine sites engagement with 53 cysteine-reactive small molecules, illustrating CPT labeling, TMT multiplexing, IMAC enrichment, fractionation, and MS analysis. Created in BioRender. Ng, M. (2025) https://BioRender.com/e74b682. **d**, Stoichiometric engagement score (R) of each cysteine quantified in human THP-1 MDM cysteine proteome treated with SCA1 (20 μM) (*n* = 3), showing SHP1 Cys102 as a top hit exhibiting >50% engagement by SCA1. R = (S/N of DMSO) / (S/N of SCA1), where S/N is signal-to-noise ratio. **e**, Pairwise comparisons of R for each cysteine quantified in human THP-1 MDM replicates treated with 5, 10, 20, or 40 μM SCA1 for 3 h (37°C), followed by CPT enrichment. **f**, SHP1 Cys102 site-specific percent modification by SCA1 in THP-1 MDMs at the indicated concentrations (*n* = 2). **g**,**h**, SHP1 **(g)** or SHP2 **(h)** cysteines site-specific percent modification by SCA1 in THP-1 MDMs at the indicated concentrations (*n* = 3). Data are mean ± s.e.m. (in **a**) or s.d. in (**d**-**h**). *P* values calculated using two-way analysis of variance (ANOVA) for multiple comparisons or two-tailed Student’s *t*-test for unpaired comparison.

Analysis of OxImmune identified redox regulated Cys102 that exists in an N-SH2 proximal pocket involved in the structural rearrangement of SHP1 from an auto-inhibited state to an active conformation^50^ **(Figure 2a,b)**. On this basis, we hypothesized that modification of SHP1 Cys102 could drive a conformational change to the N-SH2 domain leading to elevation of phosphatase activity. First, to better understand whether SHP1 Cys102 is redox regulated in macrophages, we monitored SHP1 Cys102 oxidation state in human THP-1 monocyte-derived macrophages (MDM) **(Extended Data Figure 1b & Supplementary Table 2)**. We used a series of well-established perturbations known to modify cellular ROS levels and thiol redox state. We applied lipopolysaccharide (LPS) which is a well-known trigger for endogenous macrophage ROS production^7^. Separately, we applied redox modifying agents including N-acetylcysteine (NAC) and cell permeable TCEP (tmTCEP)^51^. As expected, we found SHP1 Cys102 oxidation increased in the presence of LPS, and reduced in the presence of chemical reducing agents NAC and tmTCEP, indicating that SHP1 Cys102 is sensitive to redox regulation in macrophages **(Extended Data Figure 1c)**. To further characterize temporal redox regulation of SHP1 Cys102 in an inflammatory context, we also monitored Cys102 accessibility in THP-1 MDMs over a time course of LPS stimulation. These data demonstrate that SHP1 Cys102 is redox regulated in a time-dependent manner with oxidation peaking 60 min post-LPS in THP-1 MDMs, supporting its regulation in the context of the acute macrophage cytokine response **(Extended Data Figure 1d)**.

Having demonstrated that SHP1 Cys102 is redox regulated in macrophages and *in vivo*, we set out to discover selective cysteine-targeted ligands of SHP1 Cys102 with the hypothesis that chemical modification of SHP1 Cys102 could drive a conformational change to the N-SH2 domain leading to elevation of phosphatase activity. We applied deep cysteine chemoproteomic mapping (CPT-MS)^14^ of both human THP-1 MDMs and mouse iBMDMs to query the cysteine proteome under native conditions for engagement with cysteine-reactive small molecules. CPT-MS makes use of cysteine-reactive phosphate tags (CPTs) that facilitate enrichment of cysteine-containing peptides using metal affinity chromatography, allowing for deep mapping of the cysteine proteome^14,52,53^. CPT-MS quantified small-molecule engagement with ∼20,000 unique cysteines in a single experiment **(Figure 2c & Supplementary Table 3,4)**. Using CPT-MS, we screened for small molecules that could selectively engage SHP1 Cys102 in human THP-1 MDMs and mouse iBMDMs. We identified a molecule, henceforth referred to as SCA1, that covalently modified SHP1 Cys102 **(Figure 2d & Extended Data Figure 1e)**. SCA1 exhibited a very high degree of proteome-wide selectivity for SHP1 Cys102 at low micromolar concentrations. Between 5-40 μM, SCA1 reproducibly and selectively labelled SHP1 Cys102 in THP-1 MDMs **(Figure 2e & Supplementary Table 3)** and iBMDMs **(Extended Data Figure 1e & Supplementary Table 4)**. At these concentrations, SCA1 did not reproducibly label other cysteine sites with higher stoichiometry compared to SHP1 Cys102, with the exception of DHX40 Cys153 that was non-specifically engaged at higher concentrations. More generally, these data indicated a near complete lack of reproducible off-target engagement of the observable cysteine proteome **(Figure 2e, Extended Data Figure 1e & Supplementary Table 3,4)**. SCA1 engagement of SHP1 Cys102 in THP-1 MDMs and iBMDMs was concentration dependent and readily achieved above 50% stoichiometry between 10-20 μM **(Figure 2f & Extended Data Figure 1f)**, and exhibited high selectivity compared to the closely related PTP SHP2 **(Figure 2g,h & Extended Data Figure 1g,h)**.

We confirmed direct labeling of Cys102 by SCA1 by examining the intact mass of recombinant human SHP1. Incubation of SHP1 with 5 or 10 molar equivalents of SCA1 resulted in a mass shift of the protein corresponding to the mass of a SCA1 adduct **(Figure 3a)**. Peptide analysis of recombinant SHP1 determined that this labeling was attributable to covalent modification of Cys102 **(Figure 3b)**. Moreover, covalent modification of SHP1 by SCA1 was completely lost upon mutagenesis of SHP1 Cys102 to serine **(Figure 3c)**. We additionally confirmed labeling of SHP1 by SCA1 in iBMDMs and THP-1 MDMs by generating a desthiobiotin-conjugated SCA1 analog (desthiobiotinylated SCA1), which we established could also label SHP1 **(Figure 3d)**. Cells were lysed and incubated with desthiobiotinylated SCA1 followed by desthiobiotin enrichment, which demonstrated robust enrichment for SHP1 at as low as 1 μM desthiobiotinylated SCA1 **(Figure 3d)**. Conversely, pre-treatment of cells with titrated SCA1 followed by 10 μM desbiotinylated SCA1 demonstrated that desthiobiotin enrichment of SHP1 could be fully competed by SCA1 **(Figure 3e)**. Taken together, these data uncover SCA1 as a selective electrophile targeting Cys102 of SHP1 *in vitro* and in mouse and human macrophages.

**Fig. 3.**
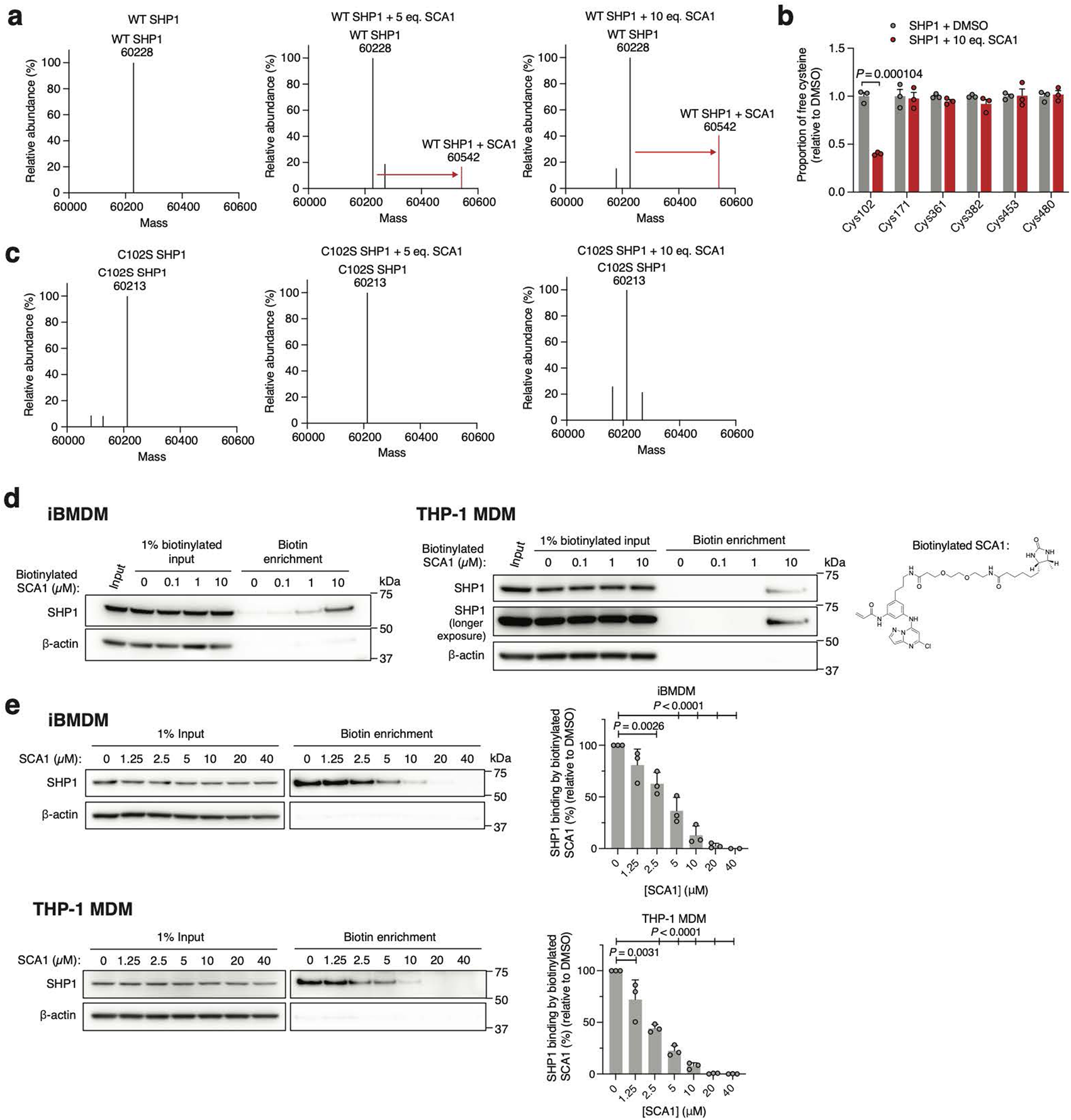
SCA1 selectively engages Cys102 on SHP1. **a,c**, Intact protein MS of human recombinant wild type (WT) or Cys102 to Ser102 mutant (C102S) SHP1 (2 μM) incubated with DMSO, 5 or 10 molar equivalents of SCA1 for 24 h (4°C). **b**, Quantitative MS determination of the proportion of free cysteine in human recombinant SHP1 incubated with DMSO or 10 molar equivalents of SCA1 for 2 h (room temperature) (*n* = 3). **d**, SHP1 binding by desthiobiotinylated SCA1 in iBMDM or THP-1 MDM lysates treated overnight (4°C) at the indicated concentrations and enriched by streptavidin pulldown. Immunoblots shown are representative of three independent experiments. **e**, SHP1 binding by SCA1 in iBMDM or THP-1 MDM cells treated at the indicated concentrations for 3 h (37°C) followed by competition for SHP1 binding with desthiobiotinylated SCA1 (10 μM). Densitometry analyses of SHP1 immunoblots from *n* = 3 independent experiments each are shown. Data are mean ± s.e.m. (in **e**) or s.d. in (**b**). *P* values calculated using one-way ANOVA for multiple comparisons involving one independent variable or two-tailed Student’s *t*-tests for unpaired comparisons.

### Structural models for SCA1-mediated modification of SHP1

To understand how SCA1 binds to SHP1 and consequently could modulate its function, we used a combination of covalent docking and molecular dynamics simulations to study its interactions with Cys102 and the surrounding residues. There are two distinct conformations of full length SHP1 reported, corresponding to the auto-inhibited (PDB ID: 2B3O) and active (PDB ID: 3PS5) form, respectively^50^. SHP1 activation requires N-SH2 dissociation from the active site^50^. Cys102 is located at the hinge connecting N-SH2 and C-SH2 domains. Through covalent docking and subsequent 1 μs molecular dynamics simulations, the proposed binding mode of SCA1 in the active form of SHP1 revealed strong cation-π interactions between the SCA1 phenyl ring and SHP1 Arg7 during most of the simulation time **(Figure 4a)**. In addition, a weaker but still significant edge-to-π interaction between the pyrazolo[1,5-a]pyrimidine ring and Phe39 persisted in 37% of the simulation length. One of the most important interactions for SCA1 is the hydrogen bond between the amine of the acrylamide and Phe5 backbone. This hydrogen bond persisted almost throughout the entire simulation, indicating its critical role. This model predicts that this interaction could effectively couple the N-terminal tail with the C-SH2 domain, which would be predicted to stabilize the active conformation, or facilitate the transition from inactive to active state by promoting the concerted motion of the two SH2 domains. The binding mode also showed limited space between the Cl atom and His114, Leu139 and Met1, and solvent exposure of the meta position of the phenyl ring, suggesting opportunities for further modification and chemical optimization of SCA1. The binding pocket is formed primarily by residues in β strands in the two SH2 domains and the loops between them, such as Arg3 to Arg7, Pro31, Ser32, Phe39, and His114 **(Figure 4a)**. In comparison with apo active state SHP1 simulation, residues from 209 to 219 became much more flexible with SCA1 bound, even though they are not in direct contact with the compound, presumably due to structural rearrangements of the internal loop in C-SH2 domain to accommodate compound binding.

**Fig. 4.**
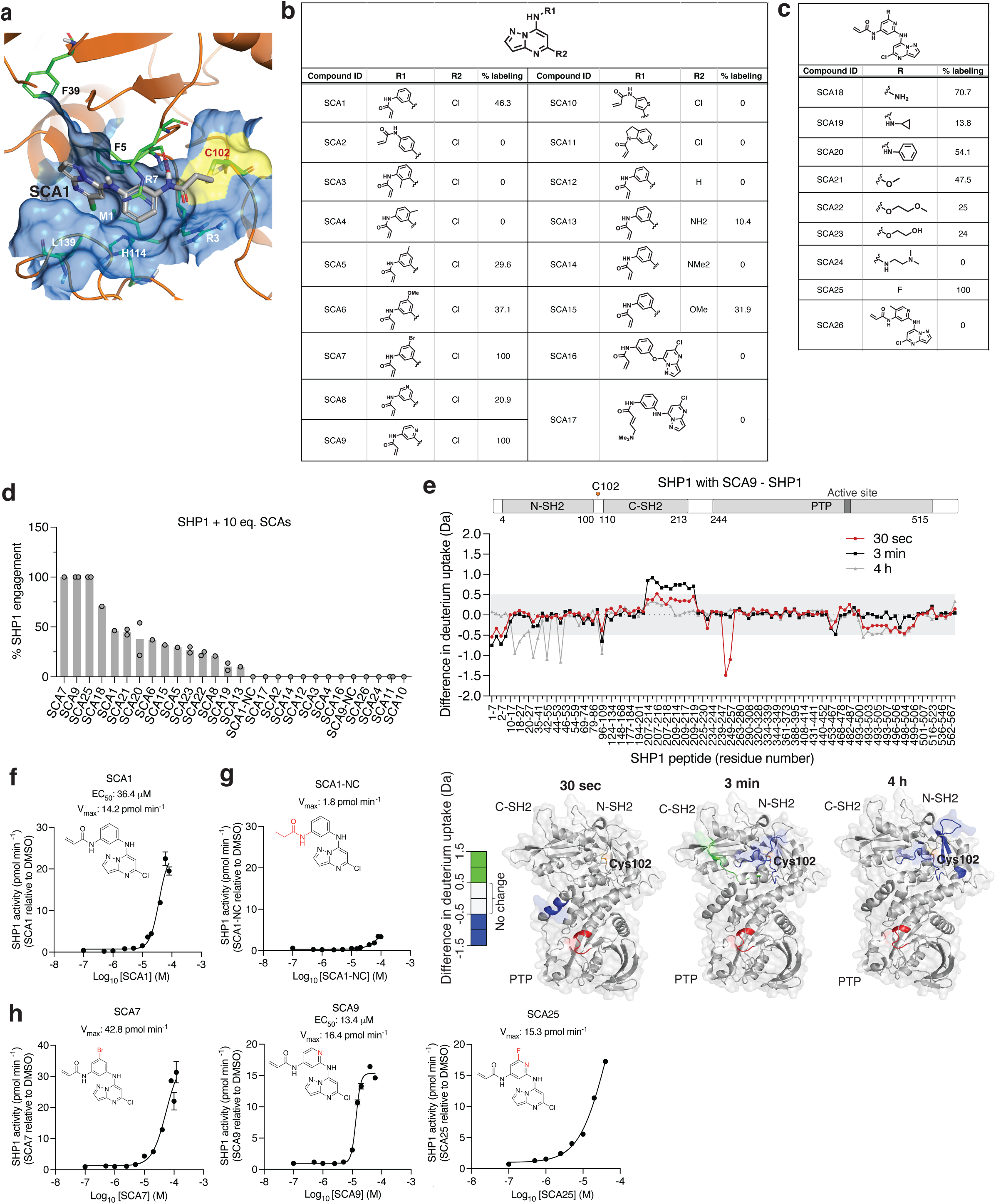
Structural consequence for SHP1 Cys102 modification by SCA1 and analogs. **a**, Covalent docking and molecular dynamics simulations of SCA1 to active conformation SHP1 (PDB ID: 3PS5). Surface representation of SCA1 (gray, sticks) and Cys102 (yellow) binding pocket consisting of residues on both SH2 domains including Arg3 to Arg7, Pro31 to Phe39 and the loop between them (brown). Stabilizing cation-π interactions between SCA1 phenyl ring and Arg7 (green, sticks), edge-to-π interaction between pyrazolopyrimidine and Phe39 (green, sticks), and hydrogen bonds between acrylamide amine and Phe5 (red, dashes) are represented. Space rigidity between Cl atom and His114, Leu139, and Met1 (green, sticks) and solvent exposure of phenyl ring meta position are indicated. **b,c**, Chemical optimization of SCA1 and corresponding SHP1 target engagement measured by intact protein MS of human recombinant SHP1 (2 μM) incubated with the indicated SCA1 analogs at 10 molar equivalents (20 μM) for 24 h (4°C). Percent SHP1 engagement represents relative abundance of small molecule engaged SHP1 to relative abundance of unlabeled SHP1. **d**, Percent SHP1 engagement of human recombinant SHP1 (2 μM) incubated with the full breadth of SCA1 analogs at 10 molar equivalents (20 μM) for 24 h (4°C) and measured by intact protein MS. Percent SHP1 engagement represents relative abundance of small molecule engaged SHP1 to relative abundance of unlabeled SHP1. Data are mean ± s.d. of two independent experiments. **e**, Deuterium uptake difference plot between SHP1 (10 μM) incubated with SCA9 (100 μM) and SHP1 alone for a subset of SHP1 peptides (top panel) and represented on SHP1 open, active crystal structure (PDB ID: 3PS5) (bottom panel). Red ribbon indicates the SHP1 active site pocket. Full list of identified SHP1 peptides is provided in **Supplementary Table 5**. **f-h**, Recombinant human SHP1 phosphatase enzymatic activity following 2 h pre-incubation (room temperature) with DMSO, SCA1 (**f**), non-covalent reversible analog of SCA1 (SCA1-NC) (**g**), or SCA7, SCA9 and SCA25 (**h**) at the indicated concentrations. Phosphatase activity as a function of SCA1 or SCA1-NC concentration relative to DMSO control measured at 37°C is shown. All data are mean ± s.e.m., *n* = 4 technical replicates.

To further understand whether SCA1 binding is compatible with the inactive conformation of SHP1, we covalently docked SCA1 to the auto-inhibited form of SHP1 (PDB ID: 2B3O) **(Extended Data Figure 2a)**. Unlike in the active form where the compound situates in a typical ligand binding pocket, the inactive form only provides a shallow binding surface formed by 3 domains of SHP1. There were no strong hydrophobic interactions to anchor the compound for sufficient residence time for the covalent bond formation. Therefore, we believe that SCA1 was not able to bind to the inactive conformation of SHP1. Using the close analog SHP2 as the example, SHP2 allosteric inhibitors bind to the inactive form that is very similar to the apo auto-inhibited form and stabilize the inactive conformation **(Extended Data Figure 2b, left)**. It is unlikely that SCA1 could induce and stabilize an unobserved inactive configuration of SHP1. Comparing with the SHP2 inactive conformation, the C-SH2 orientation of SHP1 is significantly different from SHP2 C-SH2 domain **(Extended Data Figure 2b, right).** The allosteric site of auto-inhibited SHP2 does not exist in the SHP1 inactive configuration. In fact, the covalent SCA1 occupies a different location when SHP1 and SHP2 are aligned **(Extended Data Figure 2c)**. Moreover, the hypothetical binding mode of SCA1 for inactive SHP1 would clash with C-SH2 of SHP2 if it were to covalently bind to the hinge cysteine in SHP2 **(Extended Data Figure 2d)**. Taken together, SCA1 selectively reacts with Cys102 in SHP1 and induces a binding site that is distinct from allosteric pockets observed in SHP2. The binding is believed to stabilize the active conformation rather than the auto-inhibited conformation. It is possible that a different scaffold or molecule can also activate SHP2 via a similar mechanism, but that remains to be seen.

Additionally, given the sequence homology between SHP1 and SHP2, we also examined whether SCA1 demonstrates selectivity for either phosphatase using structural modeling, discussed in Supplementary Information Note (**Extended Data Figure 2c,e)**. Taken together, structural modeling and the binding model of SCA1 can sufficiently rationalize its selectivity.

### Optimization of SCA1 to enhance engagement with SHP1

Using the SHP1-SCA1 docking model as a guide, we next explored the tolerability of the SCA1 scaffold to modification to enhance SHP1 binding, and developed analogs SCA9, SCA7 and SCA25 as more potent SHP1 Cys102 covalent binders for subsequent studies, discussed in further details in Supplementary Information Note **(Figure 4a-d & Extended Data Figure 3a,b)**.

### Engagement of Cys102 by SCAs relieves SHP1 auto-inhibition

In our structural model, the conformation of the SCA1-SHP1 complex suggested that Cys102 modification could facilitate rearrangements of the N-SH2 domain. To examine the SCA1 binding mode and characterize the structural effects of SCA1 binding to SHP1, we used hydrogen/deuterium exchange mass spectrometry (HDX-MS) to probe the effects of Cys102 modification on structural dynamics of SHP1. HDX-MS analysis of human SHP1 alone indicated distinct HDX profiles for the N- and C-SH2 domains with highly dynamic inter-domain linker regions and C-terminal tail **(Supplementary Table 5)**. In addition, low deuterium incorporation into the backbone was observed for the majority of the PTP domain including peptides that span the active site, suggesting a stable structure, which is expected for the auto-inhibited form of the enzyme **(Supplementary Table 5)**. In order to produce uniformly labeled SHP1, we used the SCA1 analog SCA9, which labeled Cys102 to 100% stoichiometry **(Figure 4d)**. Not surprisingly, protection from deuterium incorporation was evident for SCA9 modified SHP1 in those peptide regions proximal to Cys102, due to reduced flexibility resulting from SCA9 anchorage in the binding pocket **(Figure 4e)**. This protection was either at the earliest time points for Met1-Asp8 and Phe248-Gln254 in the N-SH2 and C-SH2-PTP linker region, or at the later time points for Leu18-Gly20, Leu28-Leu41, and Ile54-Gln55 in the N-SH2 domain. Incubation with SCA9 resulted in an early deprotection of the C-SH2 domain, and increased protection of the N-SH2 domain over time **(Figure 4e & Supplementary Table 5)**. Specifically, increases in deuterium incorporation for the extreme C-terminus of the C-SH2 domain for Leu209-Asn219 was observed. These increased exchange rates were also supported by molecular dynamics simulations **(Extended Data Figure 3c & Supplementary Table 6)**.

The binding mode of SCA1, and consequent structural effects on SHP1 by SCA9, suggests that these molecules could destabilize the auto-inhibited state of SHP1 and enhance SHP1 phosphatase activity. We examined recombinant human SHP1 activity and found that SCA1 activated SHP1, exhibiting an apparent EC_50_ of 36.4 μM and maximal activity of 14.2 pmol min^-1^ **(Figure 4f)**. We examined the kinetics of activation and engagement of SHP1 by SCA1, observing time and concentration dependence **(Extended Data Figure 3d,e)**. SCA1 activation occurred in the presence of thiol reducing agents, and a SCA1 analog lacking the acrylamide warhead (SCA1-NC) largely abrogated activation **(Figure 4g)**. SCA9, SCA7 and SCA25, all of which afforded improved SHP1 engagement also further enhanced SHP1 phosphatase activity compared to SCA1 **(Figure 4d,h)**. It is worth noting the lower solubility of SCA25 and SCA7 precluded increasing concentrations beyond 40 µM and 120 µM respectively, limiting the assessment of EC_50_ across a wider concentration range. As such, the observed SAR is consistent with the predicted binding mode. Together, HDX-MS and biochemical activity data supported a model whereby SCA1 and analogs destabilized the auto-inhibited form of SHP1 at the C-SH2-PTP domain linker to lower the activation barrier to rearrangement that is necessary to access phosphorylated substrate.

### SCAs engage inhibit TLR4-driven cytokine production

In macrophages and dendritic cells, activated SHP1 binds cytokine receptors and IRAKs to antagonize IRAK-mediated downstream NF-κB signaling^48,49^. Of note, SHP1 mediated antagonism of IRAK1 signaling is thought to be attributable in part to its phosphatase activity as well as to its binding with IRAK immunoreceptor tyrosine-based inhibitory motifs (ITIM), both of which are driven by the active conformation SHP1 upon relief of auto-inhibition^48,49^. To examine whether SCA1 binding to Cys102 on SHP1 alters the ability of the tandem SH2 domains to access phosphorylated ITIM and activate SHP1 phosphatase activity, we first aligned the crystal structure of the phosphatase domain of SHP1 bound to phosphorylated Tyr469 signal regulatory protein alpha (SIRPα) ITIM peptide on PDB domain (PDB ID: 1FPR) with our SCA1-SHP1 binding model **(Extended Data Figure 4a)**. In this structural alignment, the binding site of the phospho-peptide (orange ribbon) on SHP1 phosphatase domain (gray cartoon) shows considerable distance from the SCA1 small molecule (stick-ball mode) allosteric binding site in the SCA1-bound SHP1 model (marine cartoon). While the peptide would clash with the inactive state of SHP1 in the crystal structure, it is entirely structurally compatible with the native active state and SCA1 activated state. To further investigate the phospho-tyrosine (pY) peptide binding to SH2 domains, we next aligned C-SH2 of SHP1 with pY NKG2A peptide (PDB ID: 2YU7, NMR structure, orange ribbon, gray cartoon) with our full-length model (marine cartoon) **(Extended Data Figure 4b)**. Since there are no available co-crystal or NMR structures of N-SH2 with pY peptides, an N-SH2 SHP2 domain co-crystal of pY peptide (PDB ID: 3TL0, green cartoon) was used to illustrate the binding site of N-SH2 domain in SHP1. In both C-SH2 and N-SH2, it is clearly seen that our covalent agonist SCA1 binds to a distinct site from all peptide binding clefts. To further evaluate this model in which SCA and ITIM binding to SHP1 are compatible, we also experimentally measured the binding affinity of a pY-IRAK1 ITIM peptide to titrated concentration of a catalytically dead SHP1 (SHP1 C453S) to prevent spontaneous hydrolysis of the phospho-peptide, that was pre-incubated with optimized SCA analogs or a known SHP1/SHP2 inhibitor NSC-87877, and tracked fluorescence polarization of the pY-ITIM peptide. We show that the binding affinity of the pY-ITIM peptide to SHP1 C453S showed no change when pre-incubated with or without SCAs **(Extended Data Figure 4c)**. We also further measured the binding affinity of the pY-IRAK1 ITIM peptide to SHP1 C453S in the presence of increasing concentrations of the optimized SCA analogs or NSC-87877 inhibitor. The data showed that the binding affinity of pY-ITIM peptide to SHP1 C453S also showed no change with SCA analogs, in contrast to the SHP1/SHP2 inhibitor NSC-87877 that competes with pY-ITIM peptide for SHP1 binding, together indicating that Cys102 agonism by SCA and ITIM binding are not dependent on one another, further supporting our structural binding model **(Extended Data Figure 4d)**.

To understand the functional consequences of SHP1 Cys102 engagement, we next examined the cellular effects of SCAs on SHP1 regulated immunological processes by evaluating the signaling responsible for production of pro-inflammatory cytokines in mouse iBMDMs treated with SCA1 and the SHP1 inactive SCA1-NC derivative. Following stimulation of TLR4 signaling by LPS, macrophage cytokine production is initiated by IRAK1 potentiation of IKK-mediated downstream degradation of phosphorylated IκBα, which leads to NF-κB phosphorylation^48^. We found near complete depletion of IκBα by 15 min following LPS stimulation, which recovered by 120 min, matching the established kinetics previously reported^54^ **(Figure 5a)**. Low micromolar SCA1 prevented acute degradation of IκBα, and prevented phosphorylation of the NF-κB p65 subunit upon stimulation by LPS (**Figure 5a)**. The SCA1-NC derivative lacking the warhead required to modify SHP1 Cys102 had no effect on this pathway **(Figure 5b)**.

**Fig. 5.**
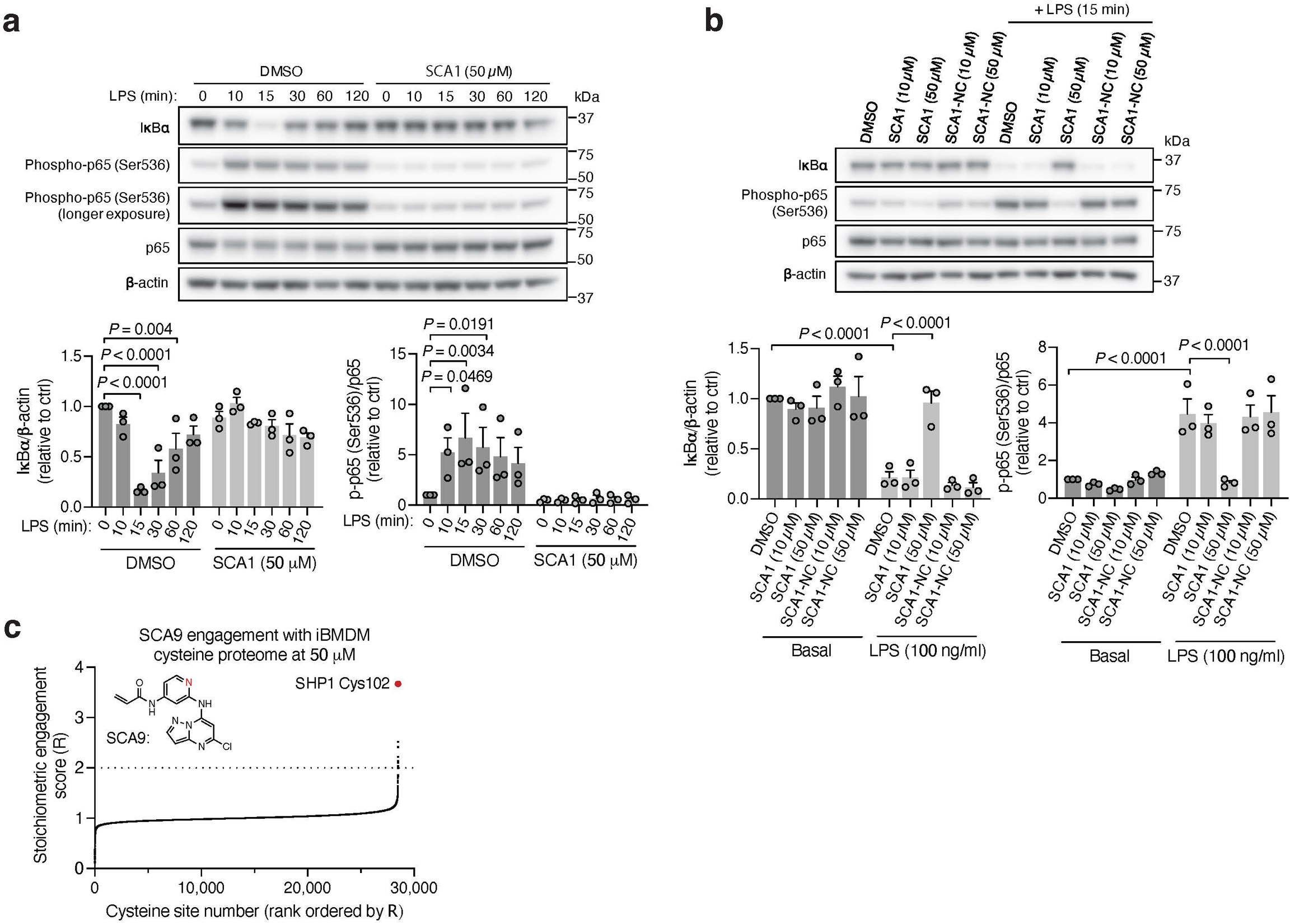
Covalent SHP1 agonists antagonize TLR4 signaling in macrophages. **a**, LPS-induced (100 ng/ml) IκBα degradation and NF-κB p65 phosphorylation over a time course of 2 h in iBMDMs pre-treated with DMSO or SCA1 (50 μM) for 3 h. Immunoblots shown are representative of three independent experiments. **b**, LPS-induced (100 ng/ml) IκBα degradation and NF-κB p65 phosphorylation in iBMDMs following 3 h pre-incubation with DMSO, SCA1 or its non-covalent reversible analog of SCA1 (SCA1-NC) at the indicated concentrations. Immunoblots shown are representative of three independent experiments. **c**, Stoichiometric engagement score (R) of each cysteine quantified in iBMDM cysteine proteome treated with SCA9 (50 μM), showing SHP1 Cys102 as top hit exhibiting >50% engagement by SCA9. R = (S/N of DMSO) / (S/N of SCA9), where S/N is signal-to-noise ratio. Data are mean ± s.e.m. *P* values calculated using two-way ANOVA for multiple comparisons.

We next examined whether optimized SCA1 derivatives effectively engaged SHP1 in cells by assessing competitive SHP1 binding with desthiobiotinylated SCA1, as described in **Figure 3**. Notably, optimized SHP1 agonists SCA9, SCA7 and SCA25 also effectively engaged SHP1 in iBMDMs, while SCA1-NC did not **(Extended Data Figure 3b & 5a,b).** We further examined if engagement of SHP1 by the SCA1 derivatives generalized in other cellular models using primary BMDMs, THP-1 MDMs and THP-1 monocytes. In all cases we observed robust reproducible engagement of SHP1 by SCA9, SCA7 and SCA25 **(Extended Data Figure 5c-e)**. Additionally, we selected two of the improved SCAs (SCA9 and SCA5) and the SCA1-NC derivative for CPT-MS to query proteome wide engagement in iBMDMs under native conditions. Like SCA1, we found SCA9 and SCA5 also exhibited proteome wide selectivity for SHP1 Cys102 while inactive SCA1-NC showed no engagement with SHP1 Cys102 **(Figure 5c, Extended Data Figure 5f & Supplementary Table 7)**.

Interestingly, SCA9 and SCA5 did not engage the only reproducible off-target observed for SCA1, Cys153 on DHX40 **(Supplementary Table 7)**. Next, we profiled a panel of SHP1 agonists and structurally similar SHP1 inactive SCA1 derivatives for modulation of TLR4-dependent IRAK1 signaling. SCA derivatives SCA9, SCA7 and SCA25 exhibiting both improved SHP1 engagement **(Extended Data Figure 3b & 5b & Figure 5c)** and enhanced SHP1 phosphatase activation **(Figure 4h),** also demonstrated more robust inhibitory effects on TLR4-mediated IRAK1 signaling and downstream NF-κB activation at low micromolar concentrations **(Figure 6a)**. Conversely, structurally similar SCA1 derivatives SCA1-NC that lack the covalent warhead, or SCA6 that is a weak SHP1 labeler **(Figure 4d & Extended Data Figure 3a,b)** exhibited no effect on macrophage TLR4 signaling **(Figure 6a)**. The observed cellular structure-activity relationship is consistent with the predicted binding mode and provided a strong correlation between capacity for SHP1 Cys102 engagement and effects on TLR4-mediated IRAK1 signaling. Together, these data support that the effects of SCAs on TLR4-mediated IRAK1 signaling depended on engagement of SHP1 Cys102.

**Fig. 6.**
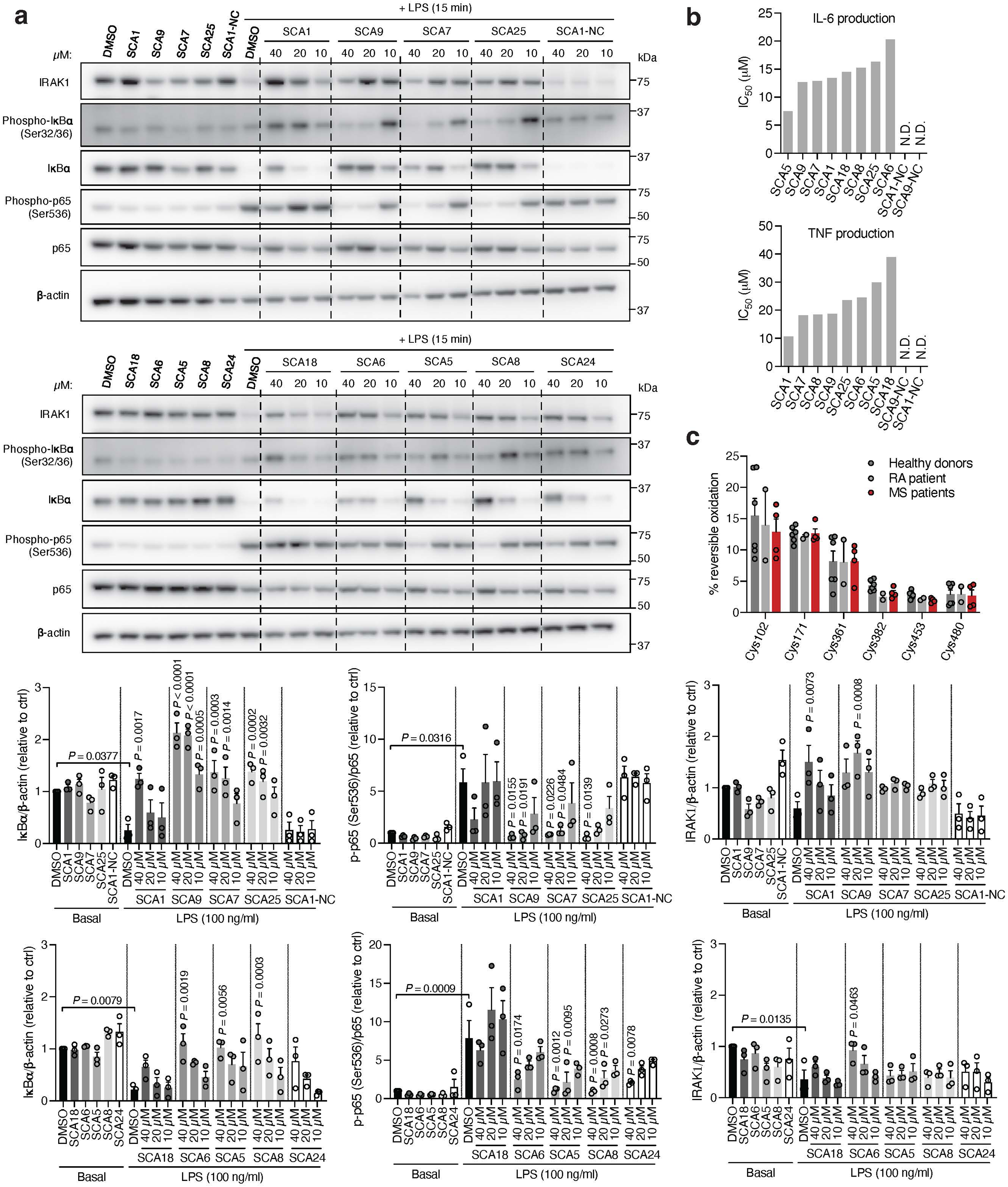
Covalent SHP1 agonists inhibit TLR4-mediated pro-inflammatory cytokine response in macrophages and *in vivo*. **a**, LPS-induced (100 ng/ml) IκBα degradation and NF-κB p65 phosphorylation in iBMDMs following 3 h pre-incubation with DMSO or SCA derivatives at the indicated concentrations. Immunoblots shown are representative of three independent experiments. **b**, Pro-inflammatory cytokine IL-6 and TNF levels in cell supernatants of iBMDMs pre-incubated 3 h with SCA1, SCA1-NC or the respective derivatives over a dose response (0.313-80 μM) followed by 6 h LPS stimulation (100 ng/ml). Normalized percent cytokine production as a function of concentration are used to calculate IC_50_ values. **c**, Percent reversible oxidation of SHP1 cysteines in CD14+CD16-monocytes of healthy donors (*n* = 6) and rheumatoid (RA) (*n* = 2) and multiple sclerosis (MS) (*n* = 4) patients derived PBMCs. Data are mean ± s.e.m. *P* values calculated using two-way ANOVA for multiple comparisons.

Our findings thus far indicate that SCA-mediated SHP1 phosphatase activation is required to drive downstream inhibitory effects on TLR4-mediated signaling. We further examined if this effect coincides with modulation of SHP1 phosphorylation. As previously reported, phosphorylation on Tyr536 and Tyr564 contributes to relief of basal SHP1 auto-inhibition, while phosphorylation on Ser591 negatively regulates phosphatase activity^55,56^. We observed Tyr536 phosphorylation within 15 min following LPS, and inhibition of this modification by SCA1 **(Extended Data Figure 6a)**. We observed no significant difference in Tyr564 phosphorylation levels inducible by LPS, and an inhibition of the baseline phosphorylation at this site by SCA1. In contrast, we did not observe differences in Ser591 phosphorylation by LPS or SCA1. These findings suggest that one consequence of SCA1 engagement of SHP1 may be to remodel the N-SH2 and antagonize phosphorylation at Tyr536 and Tyr564. This would be consistent with our structural binding mode in which Cys102 engagement by SCA1 drives N-SH2 structural rearrangement to activate SHP1.

Given that SHP1 regulates signaling from multiple immune cell surface receptors, we also examined whether SCA1 modulation of SHP1 affects other SHP1-dependent signaling pathways. Specifically, a number of reports have highlighted the role of STAT3 phosphorylation as an intermediary response to TLR4 agonism in macrophages, as well as an underlying signaling driving inflammation in myeloid cells of an autoimmune animal model of multiple sclerosis^57-59^. We observed phosphorylation of STAT3 Tyr705 and Ser727 after LPS exposure, consistent with previous reports **(Extended Data Figure 6b)**^58^. We also observed that SCA1 inhibits LPS-induced phosphorylation at both STAT3 sites, indicating that SHP1 activation by SCA1 can modulate STAT3 signaling in the context of TLR4 agonism.

We next examined effects of SCAs on pro-inflammatory cytokine production downstream of TLR4-mediated activation of NF-κB. SCA1 inhibited production of NF-κB dependent pro-inflammatory cytokines including IL6 and TNF following LPS stimulation in a concentration-dependent manner **(Extended Data Figure 6c)**. SCA1 treatment alone showed no significant effect on baseline IL-6 and TNF levels **(Extended Data Figure 6d)**. For SCA1 derivatives showing improved SHP1 engagement and activation, NF-κB inhibitory effects correlated with improved IC_50_ on cytokine production **(Figure 6b)**. Moreover, in the context of TLR-dependent IL-6 and IL-1β production, SCA1 derivatives SCA5 and SCA9 inhibited IL-6 and IL-1β levels with single digit micromolar potency **(Extended Data Figure 6e,f)**. While the profile of SHP1 engagement by the SCA series correlated well with cellular efficacy on NF-κB signaling and cytokine production, it is possible that SCAs may also engage other targets relevant to the macrophage cytokine response. Nonetheless, CPT-MS of SCA1 and SCA9 indicated highly selective target engagement with SHP1 in cells and failed to identify reproducible off-targets **(Figure 2e and Extended Data Figure 1e & 5f)**. We also explored whether SCA mediated activation of SHP1 affects LPS-induced phagocytosis in iBMDMs^60^. We observed that LPS-induced phagocytic activity of macrophages was significantly inhibited by optimized SCA derivatives SCA7 and SCA25 **(Extended Data Figure 6g)**. We also assessed whether SCAs were cytotoxic to macrophages. Monitoring major markers of apoptosis by protein-MS demonstrated they were not significantly altered by SCA1 **(Extended Data Figure 6h)**.

Given the importance of SHP1 in autoimmune conditions, we next considered whether SCA1 and its derivatives could be useful in this setting. The utility of SCAs in the context of autoimmune disorders would require that sufficient SHP1 Cys102 exists in the reduced state to be available for modification. We therefore first examined the redox state of SHP1 cysteines in CD14+CD16-monocytes isolated from peripheral blood mononuclear cell (PBMCs) of healthy controls and patients with both rheumatoid arthritis (RA) and multiple sclerosis (MS) **(Extended Data Figure 7a,b)**. We found that the baseline cysteine oxidation state of all SHP1 cysteines including Cys102 in MS or RA monocytes were not significantly different when compared to that of healthy controls **(Figure 6c)**. Importantly these data support the notion that a substantial proportion of Cys102 is present in the unmodified state in the context of autoimmune disease, and is therefore amenable to pharmacological targeting with covalent ligands.

Given that myeloid cells from patients with autoimmune disease exhibit decreased SHP1 expression or activity that correlate with a hyper-inflamed state and elevation in pro-inflammatory response, we considered whether covalent SHP1 agonists could be therapeutically useful in lowering pro-inflammatory cytokine production in these contexts^18,19,43,44,46^. To examine this, we investigated the effect of optimized SCAs on monocytes derived from individuals with RA or MS, and healthy donors. We found that optimized SCA agonists markedly inhibited TNF, IL-6 and IL-1β cytokine secretion in monocytes under proinflammatory stimulation by LPS, supporting the notion that these ligands may be efficacious in the context of autoimmune diseases where TLR-driven cytokine production plays a pathogenic role **(Extended Data Figure 7c)**. Notably, the baseline proinflammatory cytokine levels in the patient monocytes were representative of the non-activated state of these CD14+CD16-monocyte populations, and SCA agonists show no significant baseline effect. Together these findings show that SCA-mediated targeting of the Cys102 redox switch on SHP1 regulates the monocyte and macrophage cytokine response to an acute pro-inflammatory stimulus.

## Discussion

In this work, we developed OxImmune to systematically identify and classify hundreds of *in vivo* redox regulated cysteines on functional pockets of immune-relevant proteins. We provide a roadmap to use OxImmune to functionalize the immunological proteome by targeting redox regulated cysteine residues, as a basis for tool compound and therapeutics development. To exemplify this approach, we focused our attention on SHP1, a central inhibitor of macrophage cytokine production. Redox regulated Cys102 was found to mark a druggable pocket that modulates the auto-inhibited state of SHP1. Developing SCA1 and its derivatives to selectively target Cys102 demonstrated that engagement of this site is sufficient to relieve auto-inhibition of the protein and enhance phosphatase activity. SHP1 controls signaling from multiple immune cell surface receptors, and in macrophages we demonstrated robust regulation of TLR4-mediated activation of NF-κB. Given the broad regulatory role of SHP1, it will be interesting to examine how modulation of Cys102 affects other SHP1-dependent signaling pathways in other immune cell types, for example the JAK/STAT cytokine pathways, and Src-kinase mediated cell adhesion and integrin pathways. Specifically, it will be important to examine established TCR targets such as TCRζ, LCK, FYN, and zeta chain-associated protein of 70 kDa (ZAP70)^61,62^. More broadly, a more complete understanding of cell- and context-dependent targets of activated SHP1 will be an interesting application of the SCA series. Significant evidence supports a role for activation of SHP1 in antagonizing autoimmune disorders including rheumatoid arthritis and multiple sclerosis^18,19,45^.

Further development of SCAs in this context could also provide new therapeutic approaches in these diseases. However, in this context it will be important to monitor the endogenous redox state of SHP1 Cys102 and how this affects the efficacy of covalent SHP1 agonists. OxImmune analysis of SHP1 Cys102 determined that this site is subject to endogenous redox regulation *in vivo*. It is therefore likely that this site is subject to endogenous regulation by redox active metabolites, which could modulate SHP1 activity in the contexts described above. We observed low levels of oxidation in monocytes originating from individuals with RA/MS, however it may be that in other contexts competitive modifications exist which may affect the efficacy of covalent agonists. Indeed, we show in this work that modification of macrophages redox state using chemical probes or inflammatory stimulus renders this site sensitive to modulation of its oxidation state. Further examining the relevance of endogenous regulation of Cys102, or suppression of this modification, could illuminate the biological contexts where SHP1 activity is relevant in physiology or pathophysiology, and where covalent agonists may be most therapeutically useful. More broadly, using the path taken with SHP1 as an exemplar, we expect that OxImmune will provide a foundation for therapeutic exploitation of functional pockets on proteins that regulate the immune system.

## Acknowledgements

This work was supported by the Claudia Adams Barr Program (E.T.C.), the Lavine Family Fund (E.T.C.), the Pew Charitable Trust (E.T.C.), NIH DK123095 (E.T.C.), NIH AG071966 (E.T.C.), NIH AI175317 (E.T.C.), The Smith Family Foundation (E.T.C.), the American Federation for Aging Research (E.T.C.), The Dana-Farber Cancer Institute Innovation Research Fund (E.T.C.). E.T.C. is an HHMI Investigator. Cartoon illustrations in Figures 1a,g, 2b,c, and Extended Data Figure 1a were created with BioRender.com.

## Author Contributions

M.Y.N., J.C., T.Z., N.S.G. and E.T.C. conceived of and designed the study. M.Y.N. and M.C.X.Y. performed cellular experiments and analyzed data. M.N.N., G.D., I.D., S.T., J.C. and T.Z. designed and conducted chemical syntheses. M.Y.N., N.B. and S.W. developed and designed OxImmune resource and web application. H.X. provided Oximouse dataset. M.Y.N., S.S., B.Z. and H.X. carried out and analyzed data from CPT mass spectrometry experiments. H.X. developed CPT-MS technology. M.Y.N. and S.S. carried out and analyzed data from SHP1 intact protein and SHP1 cysteine site engagement mass spectrometry experiments. H.-S.S. and S.D.-P. carried out and oversaw SHP1 proteins expression and purification. J.C. performed molecular modeling. M.Y.N., T.W. and J.E. designed and carried out HDX mass spectrometry experiments and analyzed data. M.Y.N., H.T. and E.L.M. performed and analyzed data from *in vivo* experiments. M.Y.N., J.C., T.Z., N.S.G. and E.T.C. directed the research, oversaw the experiments, and wrote the manuscript with assistance from the other authors.

## Methods

### Generation of the innate immune cysteine dataset

The innate immune cysteine dataset was generated using R (Version 4.2.1) and for all dataset assembly operations the “tidyverse” package (v.2.0.0) was used. The innate immune gene dataset was obtained from the InnateDB website^20^ (downloaded February 2023: https://www.innatedb.com/). The human proteome (Uniprot: downloaded February 2023: https://www.uniprot.org) was filtered for gene names found in InnateDB to establish a dataset of human innate immune proteins. To match conserved mouse cysteine residues to the corresponding human cysteines, mouse and human protein sequences were obtained from Uniprot (downloaded February 2023: https://www.uniprot.org). Sequence datasets were matched by Uniprot names and sequences of matched proteins were aligned using the msa package (v.1.32.0) in R and the ClustalOmega sequence alignment function^63^. Cysteine positions of conserved cysteines in mouse were extracted and matched to corresponding cysteines in human. To determine the redox regulatory potential of cysteines, reversible cysteine oxidation data was retrieved from the Oximouse dataset^14^ (https://oximouse.hms.harvard.edu). The average cysteine oxidation for each tissue and age group within the oximouse was calculated and the delta percent oxidation between the maximum and minimum average oxidation was determined for each cysteine. Cysteine oxidation values were then matched to mouse cysteines that are conserved in human and the dataset was filtered for cysteines within innate immune proteins. Pfam domain annotations (downloaded from https://www.ebi.ac.uk/interpro/: Human February 2024, Mouse April 2023) were matched to all human cysteines and all conserved mouse cysteines. Identifiers of available protein structure (AlphaFold and PDB, obtained from Uniprot: downloaded February 2023: https://www.uniprot.org) for each species were mapped to the cysteines. Cysteines were further annotated by mapping the following database annotations to the generated innate immune cysteine dataset: Phosphatases (downloaded February 2023 from the DEPOD website^64^: https://depod.bioss.uni-freiburg.de), kinases (downloaded February 2023 from the KinaseMD website^65^: https://bioinfo.uth.edu/kmd/index.html), Human Protein Atlas (Protein Class information downloaded February 2023 from the Human Protein Atlas website^66^: https://www.proteinatlas.org), transcription factors (downloaded February 2024 from the TFLink website^67^: https://tflink.net), and metabolic enzymes (downloaded February 2024 from the Mammalian Metabolic Enzymes Database website^68^: https://esbl.nhlbi.nih.gov/Databases/KSBP2/Targets/Lists/MetabolicEnzymes/MetabolicEnzymeDatabase.html).

### OxImmune web application

The OxImmune web application (https://oximmune-chouchani-lab.dfci.harvard.edu/) was developed using the Shiny R package (<u>v1.8.0;</u> https://shiny.posit.co/). Data visualizations were constructed with the following packages in R: ‘DT’: (https://rstudio.github.io/DT/), ‘shinyjs’: (https://github.com/daattali/shinyjs), ‘shinydashboard’: (https://rstudio.github.io/shinydashboard/), ‘glue’: (https://glue.tidyverse.org/), ‘png’: (https://www.rforge.net/png/), ‘readxl’: (https://readxl.tidyverse.org/), ‘readr’: (https://readr.tidyverse.org/), and ‘plotly’: (https://plotly-r.com/). 3D protein structural models were accessed from the AlphaFold Database (https://alphafold.ebi.ac.uk/) using the ‘shiny.molstar’ package. The OxImmune web application is based on the innate immune cysteine dataset generated above.

### Cell culture

Immortalized bone marrow-derived macrophages (iBMDMs) (prepared from C57BL/6J wild type mice, a gift from Prof. J. Kagan (Boston Children’s Hospital, Boston, MA, USA)^69^) were cultured in Dulbecco’s modified eagle medium (DMEM), high glucose, GlutaMAX supplement (Gibco, #10566016) supplemented with 10% (vol/vol) heat-inactivated fetal bovine serum (FBS) (Hyclone), 100 U/ml penicillin and 100 μg/ml streptomycin (1% penicillin/streptomycin) (Gibco, #15140122), 1 mM sodium pyruvate, in a humidified 37°C, 5% CO_2_ tissue culture incubator. THP-1 monocytes (ATCC #TIB-202) were cultured in RPMI 1640, GlutaMAX supplement, HEPES (Gibco, #72400047) supplemented with 10% (vol/vol) heat-inactivated FBS (Hyclone) and 1% penicillin/streptomycin (Gibco, #15140122), in a humidified 37°C, 5% CO_2_ tissue culture incubator. THP-1 monocytes were differentiated into macrophages with 10 ng/ml phorbol 12-myristate 13-acetate (PMA) for 24 h (Sigma Aldrich, #P1585).

### Gene expression analysis of THP-1

Total RNA from differentiated and undifferentiated THP-1 were extracted using TRIzol (Invitrogen, #15596026), purified with PureLink™ RNA Mini Kit (Invitrogen, #12183025) according to manufacturer’s protocol, and quantified using a Nanodrop™ One/One^C^ Microvolume UV-Vis Spectrophotometer. cDNA was prepared from 120 ng total RNA via reverse transcription-polymerase chain reaction (RT-PCR) using a High-Capacity cDNA Reverse Transcription Kit (Applied Biosystems, #4368814) according to manufacturer’s protocol. Real-time quantitative PCR (qPCR) was performed on cDNA using SYBR Green probes in 384-well format with a QuantStudio™ 6 Flex Real-Time PCR System (Applied Biosystems) using GoTaq qPCR Master Mix (Promega, #A6001). Fold changes in expression were calculated by the Delta Delta Ct method using human glyceraldehyde-3-phosphate dehydrogenase (GAPDH) as an endogenous control for mRNA expression. All fold changes are expressed normalized to the respective time point DMSO control. SYBR green primer pair sequences are provided in **Supplementary Table 2**.

### Quantification of cysteine oxidation state using Cysteine-reactive phosphate tag mass spectrometry (CPT-MS)

For details on quantification of cysteine oxidation state in THP-1 MDM using CPT-MS, see Supplementary Methods.

### Screening of cysteine-reactive small molecules using CPT-MS

For details on cysteine-reactive small molecules screen in iBMDMs and THP-1 MDMs using CPT-MS, see Supplementary Methods.

### SHP1 protein expression and purification

C-terminal 6x His-tag constructs of human wild type (WT) or C102S full-length (residues 1-595) and residues 1-524 SHP1 were expressed using pET 21b(+) vector in Rosetta 2 and purified via affinity and size-exclusion chromatography. Briefly, transformed cells were cultured at 37°C in Terrific Broth (TB) medium for 6h, and auto-induced at 20°C for 18 h. Cell pellets were lysed by high pressure microfluidizer in lysis buffer containing 50 mM HEPES, 500 mM NaCl, 1 mM TCEP, 20 mM imidazole, pH 7.5, 0.1% IGEPAL, 10% glycerol, 1 mM phenylmethylsulfonyl fluoride (PMSF). Following lysis, the supernatant was applied to Ni-NTA for affinity binding at 4°C for 1 h with rocking, and the resin was washed by fast protein liquid chromatography (FPLC) with wash buffer containing 25 mM HEPES, 1.5 M NaCl, 1 mM TCEP, 20 mM imidazole, pH 7.5, and eluted with elution buffer containing 25 mM HEPES, 500 mM NaCl, 1 mM TCEP, 400 mM imidazole, 10% glycerol, pH 7.5. The final protein was purified by size exclusion using a HiLoad 16/600 Superdex 200 pg column (Cytiva) in buffer containing 20 mM HEPES, 200 mM NaCl, 1 mM TCEP, pH 7.5. The resultant SHP1 containing fractions were concentrated to 10 mg/ml and stored at -80°C.

### Intact protein mass spectrometry

WT or C102S SHP1 recombinant protein (final concentration 2 μM) was incubated with 5 equivalents (10 μM) or 10 equivalents (20 μM) of the indicated SHP1 agonists at 4°C for 24 h in reaction buffer (25 mM HEPES, pH 7.4, 50 mM NaCl, 2.5 mM EDTA, 100 μM TCEP). The samples were then diluted 1 in 2 with Buffer A (0.1% formic acid, 5% acetonitrile, 95% H_2_O), and 3.5 μg per sample was subjected to LC-MS analysis using a PLRP-S 100A, 2.1 x 50 mM, 5 μM column (Agilent) on a Q-Exactive HF-X. Separation buffers A (0.1% formic acid, 5% acetonitrile, 95% H_2_O) and B (0.1% formic acid in 95% acetonitrile, 5% H_2_O) were used for the runs at 60°C, 0.3 ml min^-1^. The gradient used was 15% Buffer B, 0.5 min, ramped up to 95% over 4.5 min. Positive ion modes were collected with full scan analysis over 900 – 2600 *m/z* at 7500 resolution, 1 x 10^5^ AGC, 25 ms maximum ion accumulation time and 60 eV in-source collision-induced dissociation. Data processing was performed using BioPharma Finder software version 4.1 (Thermo Fisher Scientific).

### Recombinant SHP1 cysteine site engagement

Recombinant SHP1 at 8.6 μM was incubated with 86 μM SCA1 or DMSO (in 10 mM HEPES, pH 7.4, 50 mM NaCl, 2.5 mM EDTA, 100 μM TCEP) for 2 h at room temperature. Following incubation, the protein was subjected to methanol-chloroform precipitation, and the resultant protein pellets were washed twice with methanol. The pellets were then resuspended in a buffer containing 2% SDS, 50 mM HEPES pH 8.5, 40 mM iodoacetamide and 5 mM TCEP, and incubated for 1 h at 37°C in the dark. The samples were then subjected to methanol-chloroform precipitation and the pellets were washed twice with methanol. The resultant protein pellets were reconstituted with 50 μl 200 mM EPPS buffer, pH 8, and digested with Lys-C and trypsin at 100:1 substrate:enzyme ratio overnight (19 h) at 37°C. Overnight digested samples were clarified by centrifugation at 16,000 g for 10 min, and subjected to protein concentration quantification by microBCA assay (Thermo Scientific Micro BCA protein assay kit, #23235). 13.8 μg peptides for each sample were labeled with TMTpro-18plex reagents (Thermo Scientific, #A44520) in 30% ACN/EPPS solution for 1 h at room temperature. TMT labelling was then quenched with 5% hydroxylamine for 15 min and samples were acidified with 20% FA (pH<3). 2 μl of each sample was pooled, diluted to 5% ACN using 1% FA, desalted by StageTip and lyophilized by speed vac. The sample was reconstituted in 5% FA/5% ACN and analyzed by LC-MS.

### Desthiobiotin enrichment assay

For SCA1 competitive desthiobiotin enrichment, iBMDMs or THP-1 MDMs seeded in 10-cm dishes were pre-treated with 1.25, 2.5, 5, 10, 20 or 40 μM SCA1 or DMSO for 3 h (37°C). Following pre-treatment, cells were washed twice with ice-cold phosphate-buffered saline (PBS), lysed with 1 ml/dish lysis buffer (50 mM Tris pH 8.0, 150 mM NaCl, 1% NP-40, 5 mM EDTA, Roche EDTA-free protease inhibitor cocktail (Roche, #11836170001)), and incubated 30 min on ice. Clarified lysates (centrifugation at 16,000 g, 10 min, 4°C) were incubated with 10 μM desthiobiotinylated SCA1 overnight (16-18 h), 4°C, with rotation. For non-competitive desthoibiotin enrichment, iBMDMs or THP-1 MDMs seeded in 10-cm dishes were lysed and clarified as above, and incubated with 0.1, 1 or 10 μM desthiobiotinylated SCA1 overnight (16-18 h), 4°C, with rotation. 30 μl slurry streptavidin beads (Pierce high capacity streptavidin agarose, #20357) were added the next day, incubated for 3 h, 4°C, with rotation. Samples were washed 7 times, each with 1 ml lysis buffer indicated above. Beads were pelleted, reduced with 2X LDS (4X NuPAGE LDS sample buffer (Invitrogen, #NP0007) diluted 1 in 2) containing 5% β-mercaptoethanol, boiled at 95°C for 10 min, and subjected to western blotting analysis with anti-SHP1 (Cell Signaling, #3759, rabbit mAb; 1:1000 dilution) and anti-β-actin (Cell Signaling, #3700, mouse mAb; 1:5000 dilution) antibodies. 1% volume of pre-purified overnight desthiobiotinylated samples were included as input. Input samples were reduced and denatured as indicated above for desthiobiotin enriched samples.

### Molecular modeling

Covalent docking of SCA1 was performed using CovDock in Schrödinger suite (release 2023-4) using default parameters. Cys102 was defined as the reactive residue in all covalent docking calculations. Molecular dynamics (MD) simulations were performed with Desmond Molecular Dynamics System in Schrödinger suite with OPLS-4 forcefield and NPT ensemble^71^. A selected binding mode after visual inspection was used as initial structure for MD simulations. Orthorhombic simulation box with buffer of 12 Å and TIP3P water model were applied. Default equilibration protocol was used, and simulations were carried out at 293 K at 1 atm pressure. Production run was 1 μs, and the last 500 ns trajectory was used to analyze the interactions and identify the most populated configurations through trajectory clustering. Solvent accessible surface areas of selected atoms (i.e. backbone amine proton) were calculated using customized python scripts with Schrödinger suite API.

### Hydrogen/deuterium exchange mass spectrometry

Complete HDX-MS methods are included in **Supplementary Table 5**. In brief, prior to HDX-MS analyses, SHP1 recombinant protein (10 μM) was incubated with 10 equivalents (100 μM) of SCA9 at 4°C for 24 h in reaction buffer (25 mM HEPES, pH 7.4, 50 mM NaCl, 2.5 mM EDTA, 100 μM TCEP). Deuterium labeling was initiated with an 18-fold dilution into D_2_O labeling buffer (36 μl, 10 mM HEPES-KOH, 50 mM NaCl, 100 µM TCEP, 2.5 mM EDTA, pD 7.4, 99.9% D_2_O). The labeling reaction was quenched at time points 10 seconds, 30 seconds, 1 minute, 3 minutes, 10 minutes, 1 hour, and 4 hours with the addition of an equal volume of ice-cold quenching buffer (150 mM potassium phosphate pH 2.4, H_2_O) and immediately injected into a Waters HDX system. Deuterated and control samples were digested online at 15°C using a AffiPro Nep-2 column (AffiPro, AP-PC-004) and peptides were trapped and desalted on a VanGuard Pre-Column trap [2.1 mm × 5 mm, ACQUITY UPLC BEH C18, 1.7 μm (Waters, #186002346)] for 3 minutes at 100 μl/min. Peptides were eluted from the trap using a 5%-35% gradient of acetonitrile with 0.1% formic acid over 6 minutes at a flow rate of 100 μl/min, and separated using an ACQUITY UPLC HSS T3, 1.8 μm, 1.0 mm × 50 mm column. Mass spectra were acquired using a Waters Synapt G2-Si HDMS^E^ mass spectrometer in ion mobility mode. Peptides were identified from replicate analyses of undeuterated control samples using PLGS 3.0.1 and relative amounts of deuterium incorporated into each peptide were determined using DynamX 3.0. The deuterium uptake values used to generate all difference maps presented in **Figure 4e** are provided in **Supplementary Table 5**. Deuterium levels were not corrected for back exchange and thus reported as relative^72^. All HDX-MS data have been deposited to the ProteomeXchange Consortium via the PRIDE^73^ partner repository with dataset identifier PXD055006.

### SHP1 phosphatase activity assay

SHP1 phosphatase activity was assessed by monitoring dephosphorylation of the synthetic substrate DiFMUP (6,8-Difluoro-4-Methylumbelliferyl Phosphate) (Thermo Fisher Scientific, #D6567) to the fluorogenic product DiFMU (6,8-Difluoro-7-Hydroxy-4-Methylcoumarin) (Thermo Fisher Scientific, #D6566). Briefly, 100 nM recombinant SHP1 was incubated with SCA1, SCA1-NC, SCA7, SCA9, or SCA25 at the indicated concentrations for 2 h at room temperature in activity buffer (25 mM HEPES, pH 7.4, 50 mM NaCl, 2.5 mM EDTA, 100 μM TCEP). Following incubation, the reaction was initiated by addition of 80 μl pre-incubated enzyme to 20 μl DiFMUP substrate (final concentration 50 μM) in a 96-well plate (Corning, #3603) at 37°C. Fluorescence at excitation/emission wavelengths of 355/460 nm was continuously measured in a fluorescence-based SpectraMax M5 microplate reader (Molecular Devices), and converted to phosphate equivalents released using a DiFMU reference standard curve. The turnover to one picomole of DiFMU is equivalent to release of one picomole of phosphate. Rate of SHP1 activity in the presence of each agonist was calculated as picomole phosphate release per min in fold change relative to DMSO control, and the rates fitted on a non-linear regression model on GraphPad Prism 9, with calculated EC_50_ and V_max_ of the reactions as indicated.

### Fluorescence polarization assay

For non-competitive assay assessing the binding affinity of phospho-tyrosine ITIM peptide to SHP1, assay was performed in black 384-well plates with 33 μl assay volume. Catalytically dead SHP1 (SHP1 C453S) titrated from 10 nM to 25 μM were pre-incubated with SCA9, SCA7, SCA25 or the SHP1/SHP2 inhibitor NSC-87877 (50 μM) for 3 h at room temperature in assay buffer (20 mM HEPES, pH 7.6, 20 mM NaCl, 100 μM TCEP, 0.01% pluronic F-68). 20 nM FITC-labeled phospho-tyrosine IRAK1 ITIM peptide (^LVpYGFL$, where ^=FITC, $=NH_2_) was then added, and fluorescence at excitation/emission wavelengths of 482/530 nm was measured using a CLARIOstar Plus plate reader (BMG Labtech). Normalized fluorescence polarization of pY-ITIM peptide (to 10 nM protein) expressed in millipolarization (mP) as a function of SHP1 concentration is fitted using non-linear regression model on GraphPad Prism 9 to calculate binding affinity IC_50_. For competitive ligand-binding assay, assay was similarly performed as described above, except that SHP1 C453S at 1 μM was pre-incubated with the ligands above, titrated from 20 nM to 50 μM, for 1 h at room temperature, and 200 nM of FITC-labeled pY-IRAK1 ITIM peptide was added and fluorescence measured. Normalized fluorescence polarization of pY-ITIM peptide (to 20 nM ligand) expressed in mP as a function of ligand concentration is fitted using non-linear regression model on GraphPad Prism 9 to calculate binding affinity IC_50_.

### Cellular NF-κB activation assay

iBMDMs seeded in 24-well plates at 2×10^6^/ml were allowed to adhere overnight. The following day, cells were pre-treated with the indicated concentrations of SCA1 or derivatives for 3 h, followed by LPS induction at 100 ng/ml for the indicated time course or15 min. Cells were washed with ice-cold phosphate-buffered saline (PBS) and lysed in 80 μl/well of Triton-lysis buffer (25 mM Tris-HCl, pH 7.5, 100 mM NaCl, 2.5 mM EDTA, 2.5 mM EGTA, 20 mM NaF, 1 mM Na3VO4, 20 mM sodium β-glycerophosphate, 10 mM sodium pyrophosphate, 0.5% Triton X-100, Roche EDTA-free protease inhibitor cocktail (Roche, #11836170001) and 0.1% β-mercaptoethanol). Lysates were pre-cleared by centrifugation at 21,000 g, 10 min, 4°C, reduced with 4x NuPAGE LDS sample buffer (Invitrogen, #NP0007) containing 5% β-mercaptoethanol, and boiled at 95°C for 10 min. Equal amounts of proteins were resolved using NuPAGE 4-12% Bis-Tris 15-well or 26-well gels (Invitrogen, #NP0336BOX or #WG1403BOX), transferred onto PVDF membranes (Thermo Fisher Scientific, iBlot 2 Gel Transfer Device, #IB21001), and immunoblotted with anti-IRAK1 (Cell Signaling, #4504, rabbit mAb; 1:1000 dilution), anti-phospho-IκBα (Ser32/36) (Cell Signaling, #9246; mouse mAb; 1:1000 dilution), anti-IκBα (Cell Signaling, #9242, rabbit pAb; 1:1000 dilution), anti-phospho-NFκB p65 (Ser536) (Cell Signaling, #3033, rabbit mAb; 1:1000 dilution), anti-NFκB p65 (Cell Signaling, #8242, rabbit mAb; 1:1000 dilution), anti-SHP1 (Cell Signaling, #3759, rabbit mAb; 1:1000 dilution), anti-phospho-SHP1 (Tyr536) (Invitrogen, #PA5-36682, rabbit pAb; 1:1000 dilution), anti-phospho-SHP1 (Tyr564) (Cell Signaling, #8849, rabbit mAb; 1:1000 dilution), anti-phospho-SHP1 (Ser591) (Invitrogen, #PA5-143764, rabbit pAb; 1:1000 dilution), anti-STAT3 (Cell Signaling, #4904, rabbit mAb; 1:1000 dilution), anti-phospho-STAT3 (Tyr705) (Cell Signaling, #9145, rabbit mAb; 1:1000 dilution), anti-phospho-STAT3 (Ser727) (Cell Signaling, #9136, mouse mAb; 1:1000 dilution) and anti-β-actin (Cell Signaling, #3700, mouse mAb; 1:3000 dilution) antibodies, diluted in TBS-T (Boston BioProducts, Inc., #IBB-180X) containing 5% BSA, followed by incubation with horseradish peroxidase-conjugated anti-rabbit or anti-mouse secondary antibodies (Promega, #W4011 or #W4021), diluted 1:10000 in TBS-T containing 5% milk, and visualized using ECL western blotting substrates (Pierce ECL Western Blotting Substrate, #32106).

### Isolation of human peripheral blood mononuclear cells (PBMCs)

For details on isolation of human PBMCs, see Supplementary Methods.

### Isolation of human monocytes from PBMCs and flow cytometry

For details on isolation of monocytes from human PBMCs and flow cytometry assessment, see Supplementary Methods.

### Isolation of mouse primary bone marrow-derived macrophages (BMDMs)

For details on isolation of mouse BMDMs, see Supplementary Methods.

### Cytokine production quantification via ELISA assays

iBMDMs seeded in 96-well plates at 2×10^6^/ml were allowed to adhere overnight. The following day, cells were pre-treated with the indicated concentrations of SCA1 or derivatives for 3 h, followed by LPS induction at 100 ng/ml for 6 h. IL-6 and TNF cytokine levels in cell culture supernatants were quantified using mouse IL-6 or TNF DuoSet ELISA kits (R&D Systems, #DY406 or #DY410) according to manufacturer’s protocols. For measurement of pro-inflammatory cytokine IL-1β levels in cell supernatants, iBMDMs seeded and allowed to adhere as above are pre-treated with LPS (100 ng/ml) for 3 h followed by treatment with DMSO or SCA9 at the indicated concentrations for 45 min and primed with adenosine triphosphate (ATP, 5 mM) for 45 min. IL-1β levels in cell supernatants were quantified using mouse IL-1β DuoSet ELISA kit (R&D Systems, #DY401) according to manufacturer’s protocols. For RA and MS patients ELISA assays, monocytes were plated in U-shaped-bottom 96-well plates (Fisherbrand, # FB012932) at 0.8×10^6^/ml and treated with 20 μM SCA1, SCA9, SCA7 or SCA25 for 3 h, followed by LPS stimulation (100 ng/ml) for 6 h. Cells were then pelleted at 500 g for 5 min, and TNF and IL-6 cytokine levels in the supernatants were quantified using human TNF or IL-6 DuoSet ELISA kits (R&D Systems, #DY210 or #DY206) according to manufacturer’s protocols. For measurement of IL-1β levels, monocytes were additionally treated with ATP (5 mM) for 45 min following LPS stimulation, pelleted at 500 g for 5 min, and supernatant IL-1β levels assayed using human IL-1β DuoSet ELISA kit (R&D Systems, #DY201). Absorbance values at 450 and 540 nm were determined with a microplate reader (BMG LABTECH FLUOstar Omega microplate reader), and wavelength corrected absorbances were used to plot standard curves from which cytokine levels from supernatant samples were calculated by intrapolation using a 4-parameter logistic (4PL) regression on GraphPad Prism 9.

### Phagocytosis assay

iBMDMs seeded in 96-well black/clear bottom plates at 1×10^5^/ml were allowed to adhere overnight. The following day, cells were pre-treated with SCA1, SCA9, SCA7 or SCA25 (10 μM) for 3 h, followed by LPS stimulation (100 ng/ml) for 6 h. Phagocytosis was assessed using the Vybrant Phagocytosis Assay Kit (Molecular Probes, #V-6694) according to manufacturer’s protocol. Briefly, cell supernatant was removed, and reconstituted fluorescent BioParticle suspension was added to all wells and incubated for 2 h at 37°C. BioParticle suspension was then aspirated, and trypan blue suspension added to all wells and incubated for 1 min at room temperature to quench extracellular fluorescence. Fluorescence at excitation/emission wavelengths of 480/520 nm was immediately measured using a CLARIOstar Plus plate reader (BMG Labtech). Phagocytic activity was calculated using net background corrected fluorescence of each treatment condition relative to DMSO control, and expressed as percent phagocytic activity.

### Quantification of protein abundance using TMT proteomics

For assessing protein abundance in iBMDMs, cells plated in 24-well plate at 2×10^6^/ml were allowed to adhere overnight. The following day, cells were pre-treated with SCA1 at 50 μM for 3 h, followed by LPS stimulation at 100 ng/ml for 15 min. TMT proteomics was then performed on these samples as described previously^74^. For further details, see Supplementary Methods.

### Chemical syntheses

Details for chemical syntheses are provided in Supplementary Methods.

### Figures and illustrations

Figures were prepared in Adobe Illustrator 2024, GraphPad Prism 9, PyMOL (Version 2.3.1), ChemDraw (Version 22.2.0), R (Version 4.2.1), and illustrations in BioRender (https://www.biorender.com/) with publication licenses. Protein structures (indicated PDB IDs) were obtained from the PDB website (https://www.rcsb.org).

### Statistical analyses

Data were expressed as mean ± s.e.m. or s.d. and *P* values were calculated using two-tailed Student’s *t*-test for unpaired comparisons of variables, one-way ANOVA for multiple comparison of one independent variable, and two-way ANOVA involving two independent variables. ANOVA analyses were subjected to Bonferroni post-hoc test for multiple comparisons. For all experiments, all stated replicates are biological replicates unless otherwise specified.

## Data Availability

The mass spectrometry proteomics data have been deposited to the ProteomeXchange Consortium via the PRIDE partner repository with the dataset identifier PXD072062 and PXD055006. The OxImmune compendium of redox regulated cysteines on immune proteins is provided as an online resource (https://oximmune-chouchani-lab.dfci.harvard.edu/). Source data are provided with this paper.

---

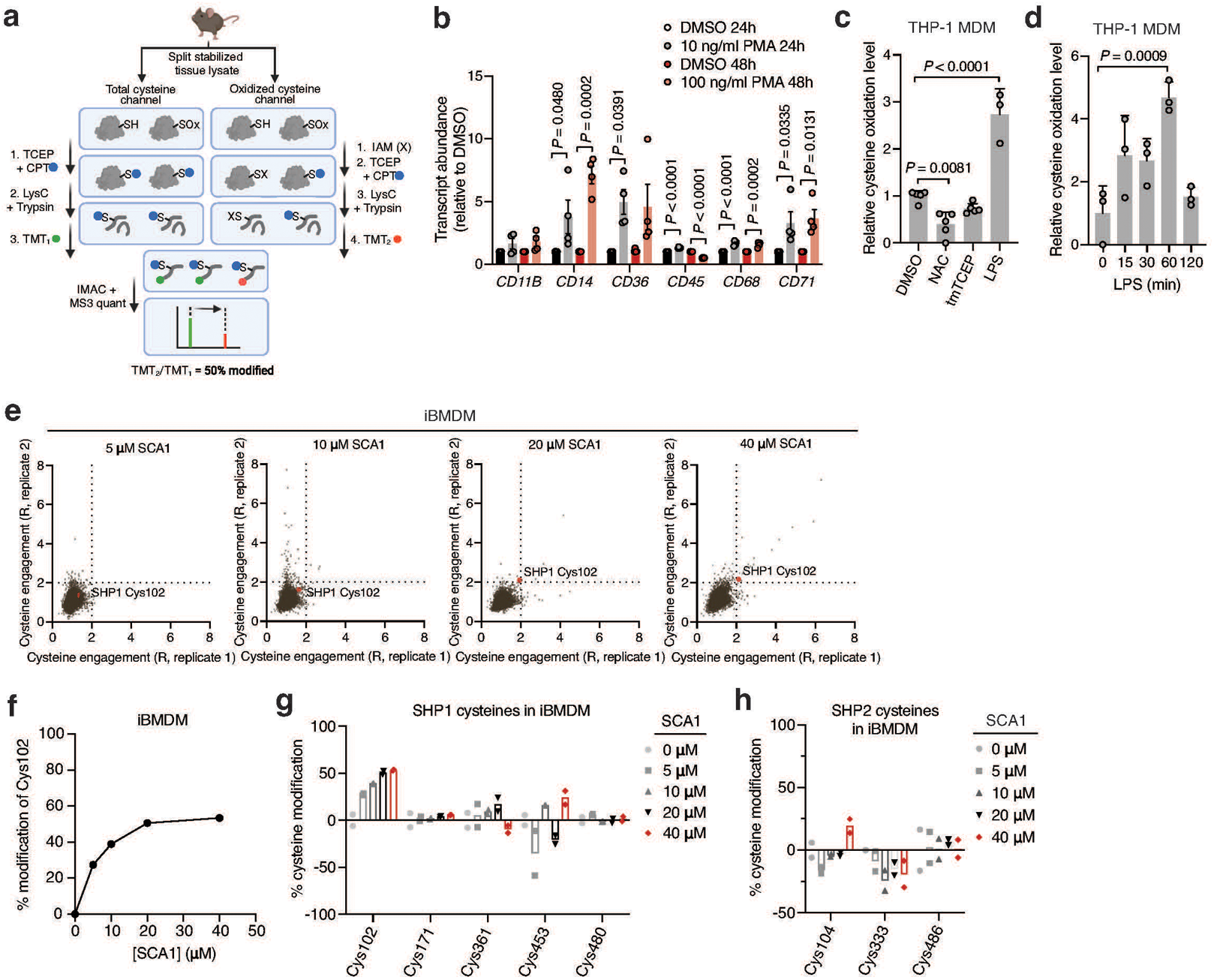

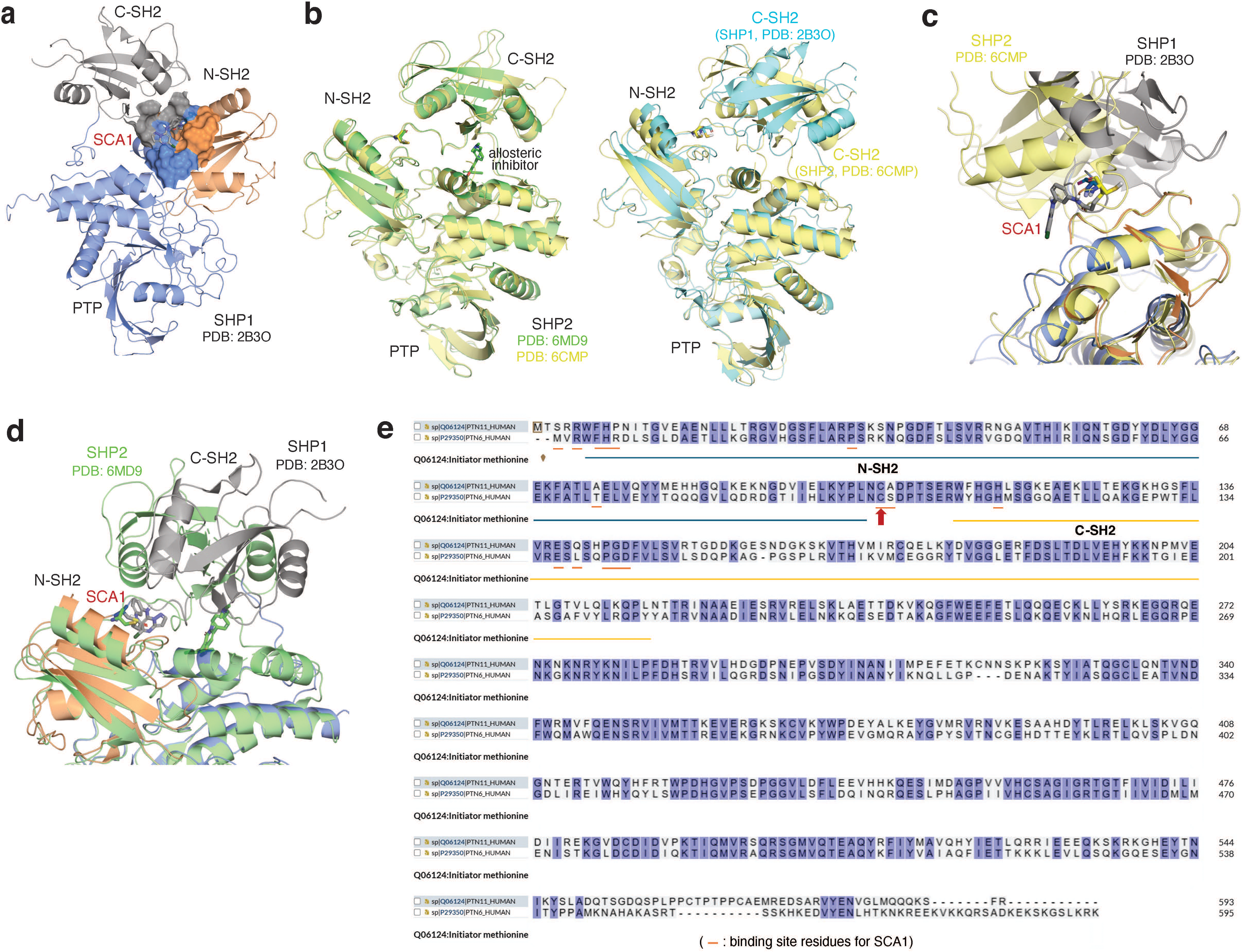

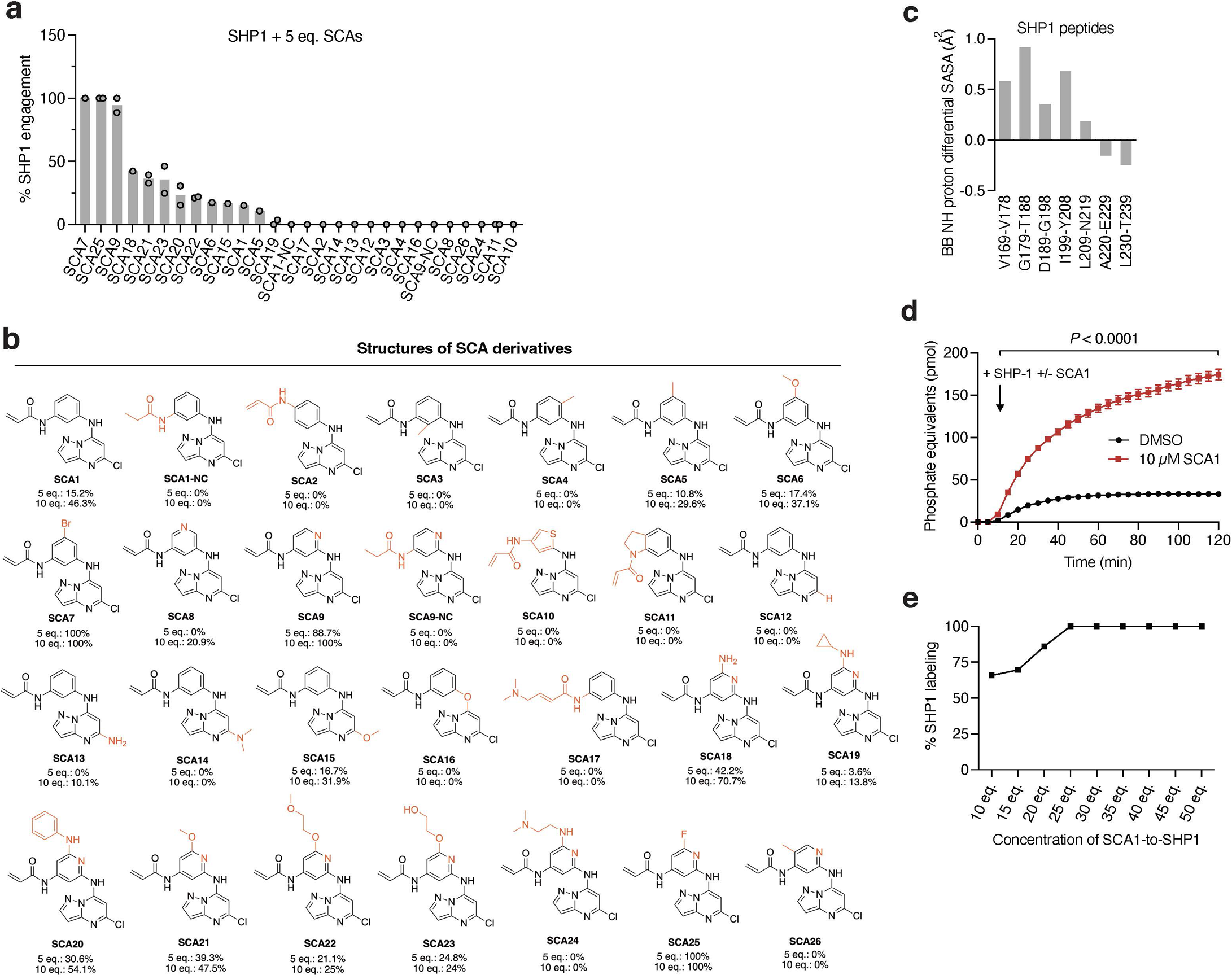

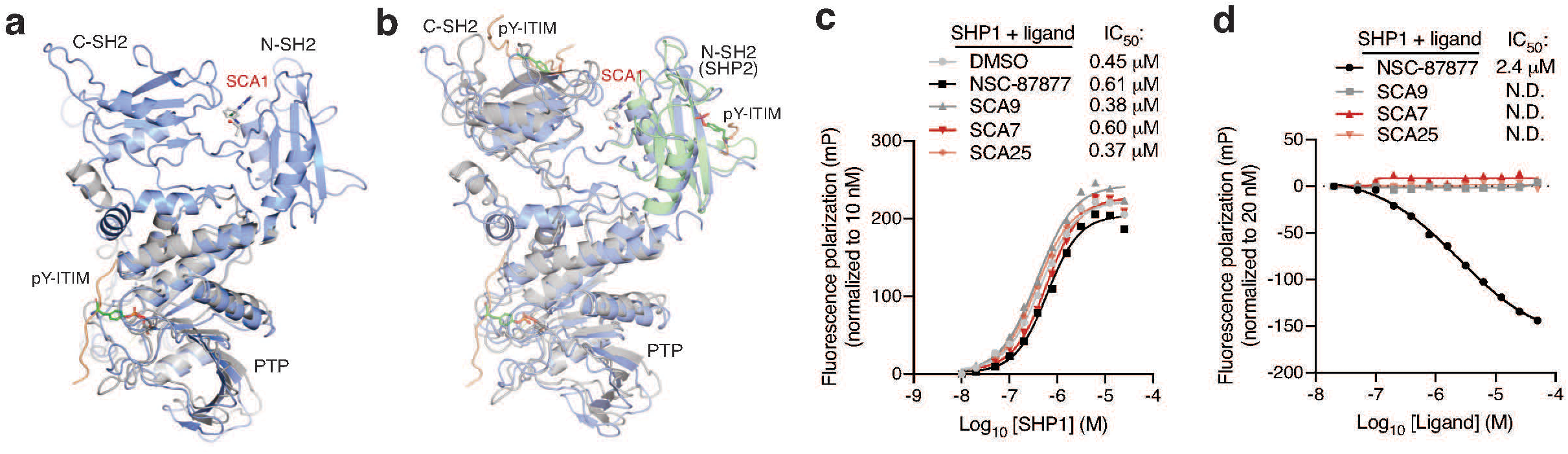

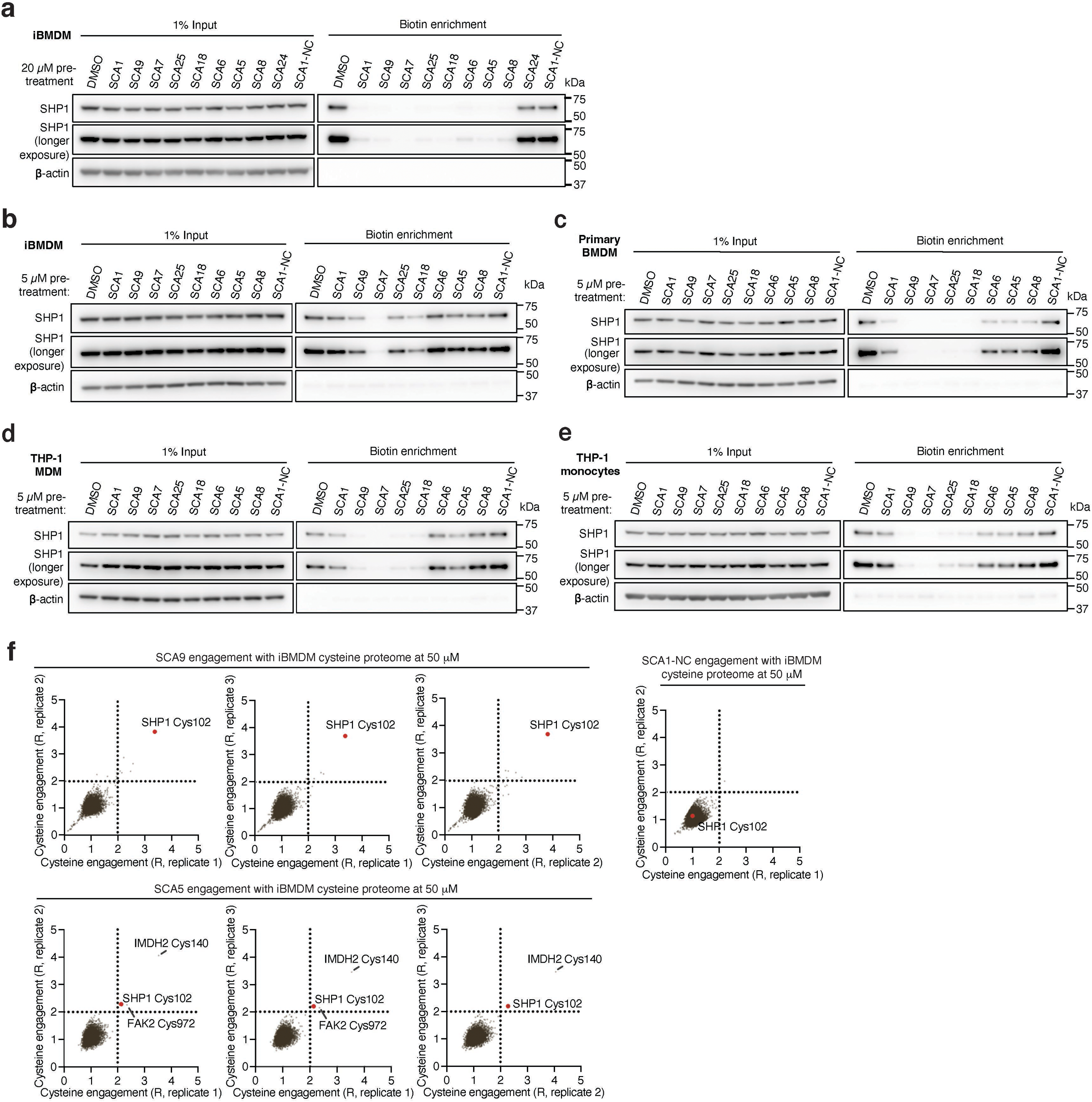

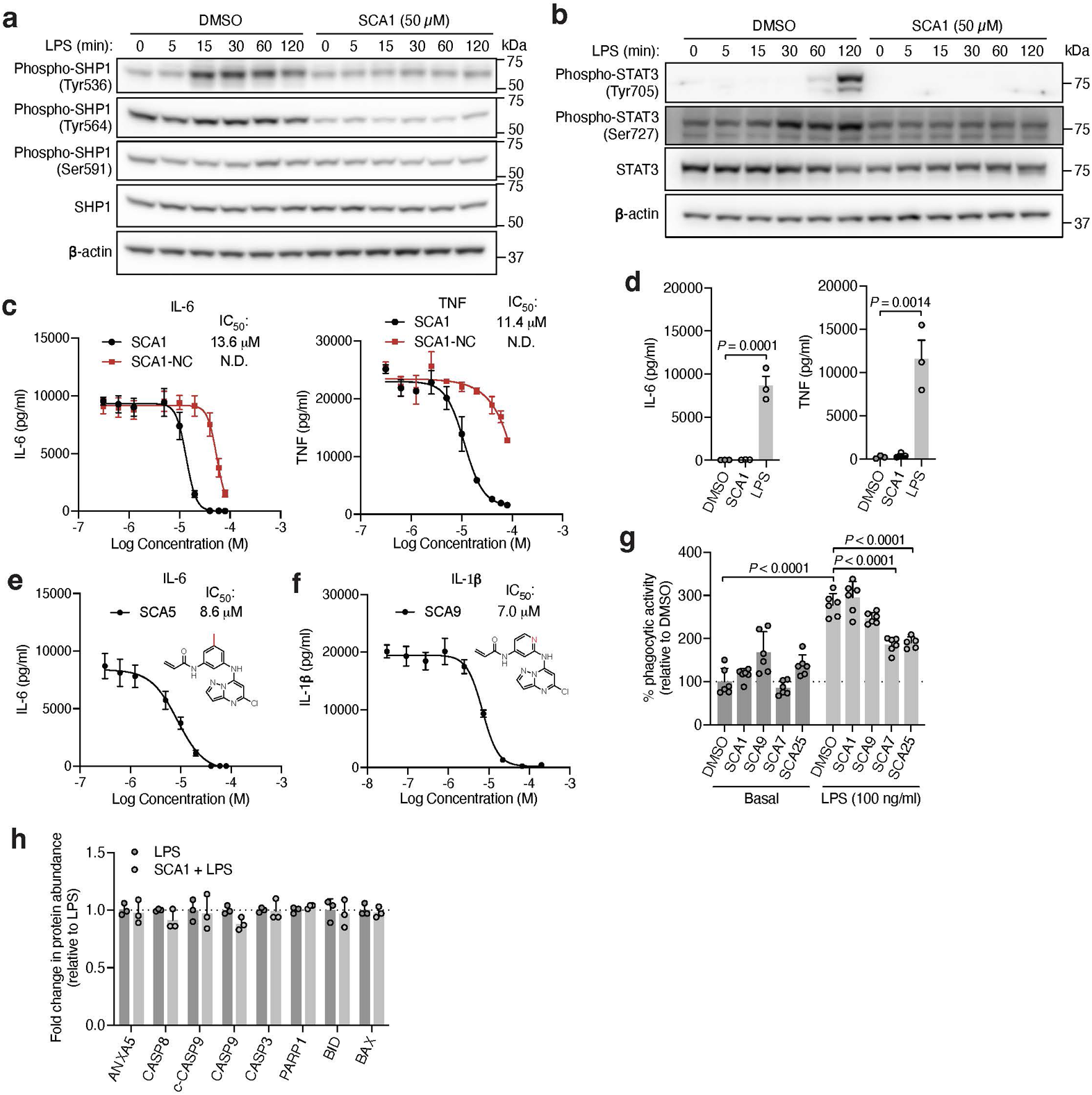

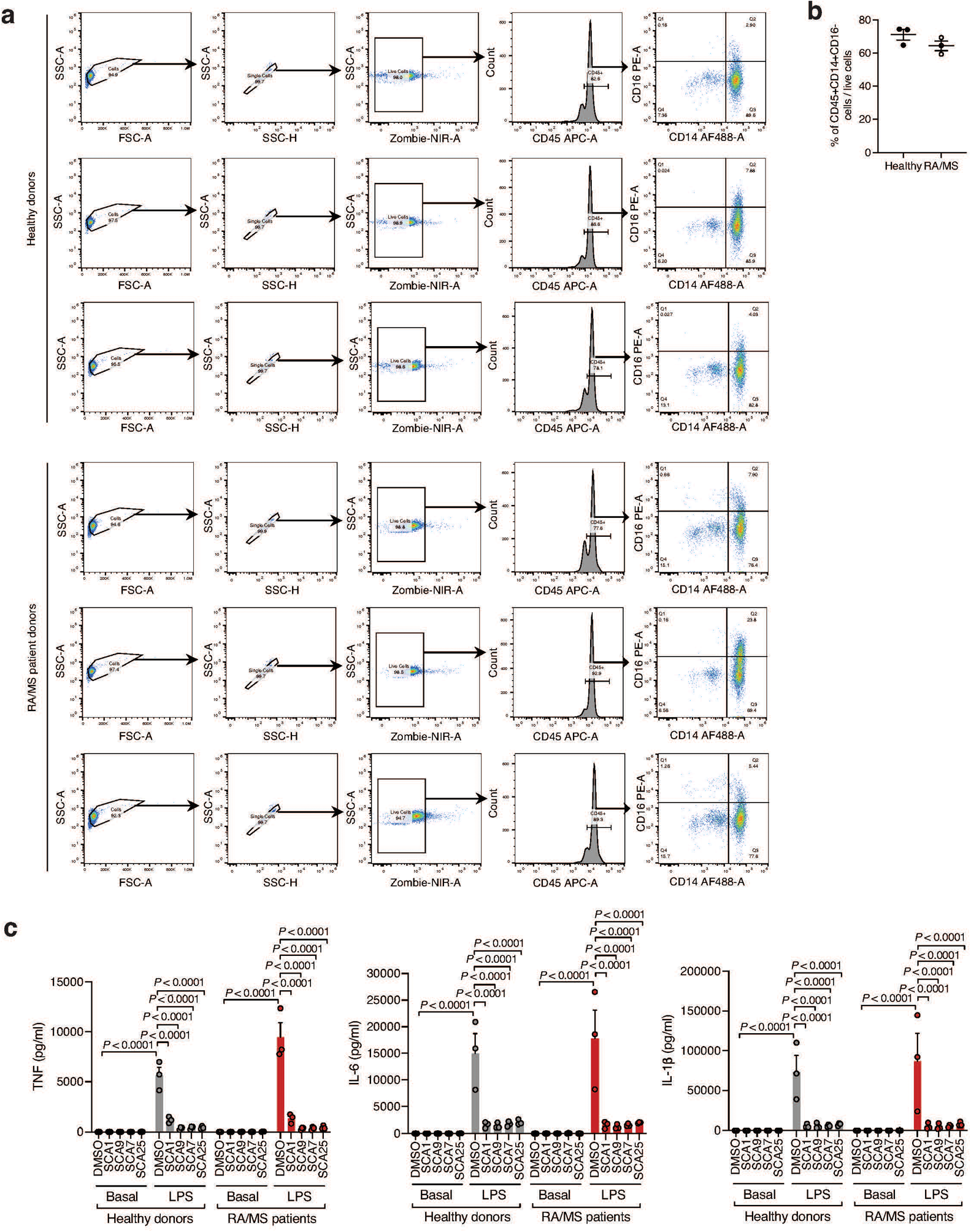

## References

1 O’Neill, L. A. & Pearce, E. J. Immunometabolism governs dendritic cell and macrophage function. J Exp Med 213, 15–23 (2016). 10.1084/jem.20151570

2 Tran, N. & Mills, E. L. Redox regulation of macrophages. Redox Biol 72, 103123 (2024). 10.1016/j.redox.2024.103123

3 Lennicke, C. & Cochemé, H. M. Redox metabolism: ROS as specific molecular regulators of cell signaling and function. Molecular Cell 81, 3691–3707 (2021). 10.1016/j.molcel.2021.08.018

4 Schieber, M. & Chandel, N. S. ROS function in redox signaling and oxidative stress. Curr Biol 24, R453–R462 (2014). 10.1016/j.cub.2014.03.034

5 West, A. P., Shadel, G. S. & Ghosh, S. Mitochondria in innate immune responses. Nat Rev Immunol 11, 389–402 (2011). 10.1038/nri2975

6 Murphy, M. P. & O’Neill, L. A. J. Krebs Cycle Reimagined: The Emerging Roles of Succinate and Itaconate as Signal Transducers. Cell 174, 780–784 (2018). 10.1016/j.cell.2018.07.030

7 Mills, E. L. et al. Succinate Dehydrogenase Supports Metabolic Repurposing of Mitochondria to Drive Inflammatory Macrophages. Cell 167, 457–470 e413 (2016). 10.1016/j.cell.2016.08.064

8 Garaude, J. et al. Mitochondrial respiratory-chain adaptations in macrophages contribute to antibacterial host defense. Nat Immunol 17, 1037–1045 (2016). 10.1038/ni.3509

9 Mills, E. L. et al. Itaconate is an anti-inflammatory metabolite that activates Nrf2 via alkylation of KEAP1. Nature 556, 113–117 (2018). 10.1038/nature25986

10 Bambouskova, M. et al. Electrophilic properties of itaconate and derivatives regulate the IκBζ-ATF3 inflammatory axis. Nature 556, 501–504 (2018). 10.1038/s41586-018-0052-z

11 Vinogradova, E. V. et al. An Activity-Guided Map of Electrophile-Cysteine Interactions in Primary Human T Cells. Cell 182, 1009–1026.e29 (2020). 10.1016/j.cell.2020.07.001

12 Boike, L., Henning, N. J. & Nomura, D. K. Advances in covalent drug discovery. Nature Reviews Drug Discovery 21, 881–898 (2022). 10.1038/s41573-022-00542-z

13 Weerapana, E. et al. Quantitative reactivity profiling predicts functional cysteines in proteomes. Nature 468, 790–795 (2010). 10.1038/nature09472

14 Xiao, H. et al. A Quantitative Tissue-Specific Landscape of Protein Redox Regulation during Aging. Cell 180, 968–983 e924 (2020). 10.1016/j.cell.2020.02.012

15 Kisty, E. A., Saart, E. C. & Weerapana, E. Identifying Redox-Sensitive Cysteine Residues in Mitochondria. Antioxidants (Basel) 12, 992 (2023). 10.3390/antiox12050992

16 Carroll, K. Defining and refining the cysteine redoxome with sulfur chemical biology. ChemRxiv [Preprint] (2022). 10.26434/chemrxiv-2022-pmk47

17 Duan, J. et al. Stochiometric quantification of the thiol redox proteome of macrophages reveals subcellular compartmentalization and susceptibility to oxidative perturbations. Redox Biology 36, 101649 (2020). 10.1016/j.redox.2020.101649

18 Christophi, G. P. et al. Interferon-beta treatment in multiple sclerosis attenuates inflammatory gene expression through inducible activity of the phosphatase SHP-1. Clin Immunol 133, 27–44 (2009). 10.1016/j.clim.2009.05.019

19 Christophi, G. P. et al. Macrophages of multiple sclerosis patients display deficient SHP-1 expression and enhanced inflammatory phenotype. Lab Invest 89, 742–759 (2009). 10.1038/labinvest.2009.32

20 Breuer, K. et al. InnateDB: systems biology of innate immunity and beyond--recent updates and continuing curation. Nucleic Acids Res 41, D1228–1233 (2013). 10.1093/nar/gks1147

21 Fu, L. et al. Nucleophilic covalent ligand discovery for the cysteine redoxome. Nat Chem Biol 19, 1309–1319 (2023). 10.1038/s41589-023-01330-5

22 Scheer, J. M., Romanowski, M. J. & Wells, J. A. A common allosteric site and mechanism in caspases. Proc Natl Acad Sci U S A 103, 7595–600 (2006). 10.1073/pnas.0602571103

23 Bozonet, S. M. et al. Oxidation of caspase-8 by hypothiocyanous acid enables TNF-mediated necroptosis. J Biol Chem 299, 104792 (2023). 10.1016/j.jbc.2023.104792

24 Yang, J. et al. Mechanism of gasdermin D recognition by inflammatory caspases and their inhibition by a gasdermin D-derived peptide inhibitor. Proc Natl Acad Sci U S A 115, 6792–6797 (2018). 10.1073/pnas.1800562115

25 Brady, K. D. et al. A catalytic mechanism for caspase-1 and for bimodal inhibition of caspase-1 by activated aspartic ketones. Bioorg Med Chem 7, 621–631 (1999). 10.1016/s0968-0896(99)00009-7

26 Xu, J. H. et al. Integrative X-ray Structure and Molecular Modeling for the Rationalization of Procaspase-8 Inhibitor Potency and Selectivity. ACS Chem Biol 15, 575–586 (2020). 10.1021/acschembio.0c00019

27 Xu, G. et al. Covalent inhibition revealed by the crystal structure of the caspase-8/p35 complex. Nature 410, 494–497 (2001). 10.1038/35068604

28 Wang, Z. et al. Kinetic and structural characterization of caspase-3 and caspase-8 inhibition by a novel class of irreversible inhibitors. Biochim Biophys Acta 1804, 1817–1831 (2010). 10.1016/j.bbapap.2010.05.007

29 Smith, J. K. et al. Identification of a redox-sensitive switch within the JAK2 catalytic domain. Free Radic Biol Med 52, 1101–1110 (2012). 10.1016/j.freeradbiomed.2011.12.025

30 Truong, T. H. & Carroll, K. S. Redox regulation of protein kinases. Crit Rev Biochem Mol Biol 48, 332–356 (2013). 10.3109/10409238.2013.790873

31 Castelo-Soccio, L., et al. Protein kinases: drug targets for immunological disorders. Nat Rev Immunol 23, 787–806 (2023). 10.1038/s41577-023-00877-7 Erratum in: Nat Rev Immunol 24, 79 (2024). 10.1038/s41577-023-00976-5

32 Ghoreschi, K., Laurence, A. & O’Shea, J. J. Janus kinases in immune cell signaling. Immunol Rev 228, 273–287 (2009). 10.1111/j.1600-065X.2008.00754.x

33 Singh, S. et al. The reduced activity of PP-1α under redox stress condition is a consequence of GSH-mediated transient disulfide formation. Sci Rep 8, 17711 (2018). 10.1038/s41598-018-36267-6

34 Chu, Y. et al. Effect of activation of protein phosphatase 1 on sulfhydryl reactivity. Arch Biochem Biophys 334, 83–88 (1996). 10.1006/abbi.1996.0432

35 Kim, H. S. et al. Constitutive induction of p-Erk1/2 accompanied by reduced activities of protein phosphatases 1 and 2A and MKP3 due to reactive oxygen species during cellular senescence. J Biol Chem 278, 37497–37510 (2003). 10.1074/jbc.M211739200

36 Gu, M. et al. Protein phosphatase PP1 negatively regulates the Toll-like receptor- and RIG-I-like receptor-triggered production of type I interferon by inhibiting IRF3 phosphorylation at serines 396 and 385 in macrophage. Cell Signal 26, 2930–2939 (2014). 10.1016/j.cellsig.2014.09.007

37 Yang, T. et al. Macrophage PTEN controls STING-induced inflammation and necroptosis through NICD/NRF2 signaling in APAP-induced liver injury. Cell Commun Signal. 21, 160 (2023). 10.1186/s12964-023-01175-4

38 Kwon, J. et al. Reversible oxidation and inactivation of the tumor suppressor PTEN in cells stimulated with peptide growth factors. Proc Natl Acad Sci U S A 101, 16419–16424 (2004). 10.1073/pnas.0407396101

39 Lee, S. R. et al. Reversible inactivation of the tumor suppressor PTEN by H2O2. J Biol Chem 277, 20336–20342 (2002). 10.1074/jbc.M111899200

40 Teruya, T. et al. Phoslactomycin targets cysteine-269 of the protein phosphatase 2A catalytic subunit in cells. FEBS Lett 579, 2463–2468 (2005). 10.1016/j.febslet.2005.03.049

41 Visscher, M., Arkin, M. R. & Dansen, T. B. Covalent targeting of acquired cysteines in cancer. Current opinion in chemical biology 30, 61–67 (2016) 10.1016/j.cbpa.2015.11.004

42 Shannon, D. A. & Weerapana, E. Covalent protein modification: the current landscape of residue-specific electrophiles. Current opinion in chemical biology 24, 18–26 (2015). 10.1016/j.cbpa.2014.10.021

43 Stanford, S. M. & Bottini, N. Targeting protein phosphatases in cancer immunotherapy and autoimmune disorders. Nature Reviews Drug Discovery 22, 273–294 (2023). 10.1038/s41573-022-00618-w

44 Nandan, D. & Reiner, N. E. Leishmania donovani engages in regulatory interference by targeting macrophage protein tyrosine phosphatase SHP-1. Clin Immunol 114, 266–277 (2005). 10.1016/j.clim.2004.07.017

45 Croker, B. A. et al. Inflammation and autoimmunity caused by a SHP1 mutation depend on IL-1, MyD88, and a microbial trigger. Proc Natl Acad Sci U S A 105, 15028-15033 (2008). 10.1073/pnas.0806619105

46 Singh, K. et al. ERK-dependent T cell receptor threshold calibration in rheumatoid arthritis. J Immunol 183, 8258–8267 (2009). 10.4049/jimmunol.0901784

47 Markovics, A., Toth, D. M., Glant, T. T. & Mikecz, K. Regulation of autoimmune arthritis by the SHP-1 tyrosine phosphatase. Arthritis Research & Therapy 22, 160 (2020). 10.1186/s13075-020-02250-8

48 An, H. et al. Phosphatase SHP-1 promotes TLR- and RIG-I-activated production of type I interferon by inhibiting the kinase IRAK1. Nature Immunology 9, 542–550 (2008). 10.1038/ni.1604

49 Abram, C. L. & Lowell, C. A. Shp1 function in myeloid cells. J Leukoc Biol 102, 657–675 (2017). 10.1189/jlb.2MR0317-105R

50 Wang, W. et al. Crystal Structure of Human Protein Tyrosine Phosphatase SHP-1 in the Open Conformation. J Cell Biochem 112, 2062–2071 (2011). 10.1002/jcb.23125

51 Takahashi, M. et al. DrugMap: A quantitative pan-cancer analysis of cysteine ligandability. Cell 187, 2536–2556.e30. (2024). 10.1016/j.cell.2024.03.027

52 Darabedian, N. et al. Depletion of creatine phosphagen energetics with a covalent creatine kinase inhibitor. Nat Chem Biol 19, 815–824 (2023). 10.1038/s41589-023-01273-x

53 Burger, N. et al. A comprehensive landscape of the zinc-regulated human proteome. bioRxiv [Preprint] 2024.01.04.574225 (2024). 10.1101/2024.01.04.574225

54 Yang, F. et al. IKK beta plays an essential role in the phosphorylation of RelA/p65 on serine 536 induced by lipopolysaccharide. J Immunol 170, 5630–5635 (2003). 10.4049/jimmunol.170.11.5630

55 Zhang, Z. et al. The role of C-terminal tyrosine phosphorylation in the regulation of SHP-1 explored via expressed protein ligation. J Biol Chem 278, 4668–4674 (2003). 10.1074/jbc.M210028200

56 Jones, M. L. et al. Regulation of SHP-1 Tyrosine Phosphatase in Human Platelets by Serine Phosphorylation at Its C Terminus. J Biol Chem 279, 40475–40483 (2004). 10.1074/jbc.M402970200

57 Xu, S. et al. Phospho-Tyr705 of STAT3 is a therapeutic target for sepsis through regulating inflammation and coagulation. Cell Commun Signal 18, 104 (2020). 10.1186/s12964-020-00603-z

58 Balic, J. J. et al. STAT3 serine phosphorylation is required for TLR4 metabolic reprogramming and IL-1β expression. Nat Commun 11, 3816 (2020). 10.1038/s41467-020-17669-5

59 Lu, H. C. et al. STAT3 signaling in myeloid cells promotes pathogenic myelin-specific T cell differentiation and autoimmune demyelination. Proc Natl Acad Sci U S A 117, 5430–5441 (2020). 10.1073/pnas.1913997117

60 Myers, D. R. et al. Shp1 Loss Enhances Macrophage Effector Function and Promotes Anti-Tumor Immunity. Front Immunol 11, 576310 (2020). 10.3389/fimmu.2020.576310

61 Lorenz, U. SHP-1 and SHP-2 in T cells: two phosphatases functioning at many levels. Immunol Rev 228, 342–359 (2009). 10.1111/j.1600-065X.2008.00760.x

62 Pao, L. I. et al. Nonreceptor protein-tyrosine phosphatases in immune cell signaling. Annu Rev Immunol 25, 473–523 (2007). 10.1146/annurev.immunol.23.021704.115647

## Methods References

63 Bodenhofer, U. et al. msa: an R package for multiple sequence alignment. Bioinformatics 31, 3997–3999 (2015). 10.1093/bioinformatics/btv494

64 Damle, N. P. & Köhn, M. The human DEPhOsphorylation Database DEPOD: 2019 update. Database (Oxford) 2019, baz133 (2019). 10.1093/database/baz133

65 Hu, R. et al. KinaseMD: kinase mutations and drug response database. Nucleic Acids Res 49, D552–561 (2021). 10.1093/nar/gkaa945

66 Uhlén, M. et al. Proteomics. Tissue-based map of the human proteome. Science 347, 1260419 (2015). 10.1126/science.1260419

67 Liska, O. et al. TFLink: an integrated gateway to access transcription factor-target gene interactions for multiple species. Database (Oxford) 2022, baac083 (2022). 10.1093/database/baac083

68 Corcoran, C. C. et al. From 20th century metabolic wall charts to 21st century systems biology: database of mammalian metabolic enzymes. Am J Physiol Renal Physiol 312, F533–542 (2017). 10.1152/ajprenal.00601.2016

69 Evavold, C. L. et al. Control of gasdermin D oligomerization and pyroptosis by the Ragulator-Rag-mTORC1 pathway. Cell 184, 4495–4511.e19 (2021). 10.1016/j.cell.2021.06.028

70 Eng, J. K., Jahan, T. A. & Hoopmann, M. R. Comet: an open-source MS/MS sequence database search tool. Proteomics 13, 22–24 (2013). 10.1002/pmic.201200439

71 Bowers, K. J. et al. Scalable algorithms for molecular dynamics simulations on commodity clusters. Proceedings of the ACM/IEEE Conference on Supercomputing (SC06), Tampa, Florida (2006). 10.1145/1188455.1188544

72 Wales, T. E. & Engen, J. R. Hydrogen exchange mass spectrometry for the analysis of protein dynamics. Mass Spectrom Rev 25, 158-170 (2006). 10.1002/mas.20064

73 Perez-Riverol, Y., et al. The PRIDE database at 20 years: 2025 update. Nucleic Acids Res 53, D543–D553 (2025). 10.1093/nar/gkae1011

74 Mills, E. L. et al. Cysteine 253 of UCP1 regulates energy expenditure and sex-dependent adipose tissue inflammation. Cell Metab 34, 140–157.e8 (2022). 10.1016/j.cmet.2021.11.003

