## Supplementary material for "A druggable redox switch on SHP1 controls macrophage inflammation": SI Note

**Supplementary Note**

**Structural models for SCA1-mediated modification of SHP1.**
Given the sequence homology between SHP1 and SHP2, we also examined whether SCA1 demonstrates selectivity for either phosphatase using structural modeling. The overall sequence similarity between SHP1 and SHP2 is 55.8% **(Extended Data Figure 2e)**. Although the catalytic sites are identical between them, the allosteric site for SCA1 has different contact residues despite the conserved Cys102 (**Extended Data Figure 2c,e**). Specifically as previously illustrated, one of the most important cation-π interaction of SCA1 central phenyl ring is established with Arg7 of SHP1, which however, is displaced with a proline in SHP2, resulting in loss of the stabilizing interaction between SCA1 and active form of SHP1. Furthermore, the chloro group on SCA1 pyrazolopyrimidine core that makes hydrophobic contacts with Met1 and Leu139 is incapable of establishing similar hydrophobic interactions with the hydrophilic residues Ser3 and Gln141 on SHP2. These differences in the SCA1 allosteric binding pocket between SHP1 and SHP2 render distinct selectivity of SCA1 for SHP1. Allosteric compound selectivity is surprising for SHP1 and SHP2, as it has been observed for SHP2 allosteric inhibitors that are selective against SHP1. While it is challenging to obtain selective catalytic binders, selective allosteric binders are not uncommon given their distinct regulatory roles. In addition, many activating mutations in SHP2 (PDB ID: 6CRF) and tandem SH2 domains with bound phospho-peptide (PDB ID: 5DF6) result in a different open conformation to the SHP1 apo active conformation. Taken together, structural modeling and the binding model of SCA1 can sufficiently rationalize its selectivity.

**Optimization of SCA1 to enhance engagement with SHP1.** Using the SHP1-SCA1 docking model as a guide, we next explored the tolerability of the SCA1 scaffold to modification to enhance SHP1 binding **(Figure 4b-d)**. The binding model of SCA1 with SHP1 revealed solvent exposure at the 5$^{th}$-position of the phenyl ring that facilitates tolerability to modifications, while considerable space rigidity and constraint at the 5$^{th}$-position of the pyrazolo[1,5-a]pyrimidine core limits amenability to modification **(Figure 4a)**. On this basis, we set out on a medicinal chemistry campaign to improve SHP1 conjugation by derivatizing SCA1. First, we fixed the R2 group with chloride at the 5$^{th}$-position and then screened synthesized aniline-containing analogs that bear an acrylamide warhead. As shown in **Figure 4b**, the para-regioisomer (SCA2) failed to label SHP1, likely owing to an incompatible trajectory of the warhead to Cys102 as suggested by the binding model **(Figure 4a)**. Similarly, the model indicated that modifications to the ortho (SCA3) and para (SCA4) positions could significantly alter the phenyl orientation due to steric repulsion with the core, resulting in loss of SHP1 conjugation **(Figure 4b)**. In contrast, modifications to the meta position including the installation of a methyl (SCA5), methoxy (SCA6), or bromo (SCA7) group retained SHP1 labeling as predicted by the model, with the bromo group further enhancing labeling. Replacing aniline with a 3-pyridyl group resulted in a slight reduction in conjugation efficiency (SCA8). In contrast, replacing aniline with a 4-pyridyl group significantly improved labeling up to 100% under the same labeling conditions, indicating a potential polar interaction or improved cation-π interaction with SHP1 Arg7 (SCA9). Although thiophene is a phenyl bio-isostere (SCA10), it failed to form an adduct with SHP1, likely due to altered warhead orientation preventing a Michael addition reaction. Similarly, rigidifying the warhead by cyclization resulted in complete loss of SHP1 conjugation (SCA11), further supporting the requirement of the hydrogen bond between the amine of the acrylamide and Phe5 backbone in the model. In addition, we fixed the warhead and replaced Cl with H, $NH_2$, $NMe_2$ or OMe. As shown in **Figure 4b**, while H and $NMe_2$ (SCA12 and SCA14) completely abolished the conjugation with SHP1, $NH_2$ and OMe modifications (SCA13 and SCA15) retained 10% and 32% labeling respectively, consistent with a small hydrophobic cavity in this region. We also explored whether conjugation could be realized through an oxygen linker. However, compound SCA16 proved to be incompetent for conjugation, presumably due to electrostatic repulsion between the oxygen and carbonyl of Pro142. Adding a methylene dimethyl amine to the terminal end of the warhead also blocked the warhead's proximity to Cys102, likely as a result of steric hindrance or incompatible electrostatic environment due to multiple positively charged residues residing in the pocket (SCA17).

We next advanced SCA9 as a new lead compound and conducted a focused structure-activity relationship (SAR) study around the pyridine ring. Installing polar groups ortho to the nitrogen pyridine reduced SHP1 conjugation for SCA18, SCA19 and SCA20, which bear amino, cyclopropane amine, and aniline substituents respectively

(**Figure 4c**). Installation of a methoxy group in SCA21, and of extended polar groups in SCA22, SCA23 and SCA24 similarly reduced conjugation. Consistent with tolerability for a halogen (bromide) at this position in SCA7, introduction of a fluoride in SCA25 retained productive conjugation with SHP1. Finally, a methyl group introduced to the ortho position of the warhead dampened conjugation (SCA26). Thus, we collectively identified SCA9, SCA7 and SCA25 as the most potent SHP1 covalent binders with Cys102. We then advanced these compounds to biochemical and functional SHP1 studies. Analogs representing the full breadth of observed SHP1 engagement are provided in the SI **(Figure 4d & Extended Data Figure 3a,b)**.

**Supplementary Methods**

**Quantification of cysteine oxidation state using Cysteine-reactive phosphate tag mass spectrometry (CPT-MS)**

For quantification of cysteine oxidation state in THP-1 MDM, cells were plated at $2x10^6$/ml in 10-cm dishes with 10 ng/ml PMA for 24 h. The following day, cells were washed twice with culture media to remove PMA, and treated with either lipopolysaccharide (LPS, 100 ng/ml) (InvivoGen, LPS-EK Ultrapure, #tlrl-peklps), N-acetylcysteine (NAC, 10 mM) (Sigma Aldrich, #A9165) and Tris(2-methoxycarbonylethyl)phosphine hydrochloride (cell permeable TCEP, tmTCEP, 1 mM) (Sigma Aldrich, #ATE448430611) for 1 h, or LPS (100 ng/ml) for a time course (15 min, 30 min, 60 min, 120 min). Following treatment, cells were processed for CPT-MS quantification as described previously[14], with the exception that CPT-labeled peptides were not subjected to tandem mass tags (TMT)-labeling, and individual samples were purified via immobilized metal affinity chromatography (IMAC) using PTMScan® Phospho-Enrichment IMAC Fe-NTA Magnetic Beads (Cell Signaling, #20432), followed by label-free MS quantification described as follows. Peptides from individual samples were analyzed using an Evosep One (EVOSEP) liquid chromatography system coupled to a timsTOF HT (Bruker) mass spectrometer. Samples were reconstituted in water containing 0.1% formic acid and loaded onto Evotip. Peptides were separated using a PepSep C18 Column (15 cm × 150 µm, particle size 1.5 µm) and a 30 SPD method (44-min gradient) on Evosep One. CaptiveSpray was operated at 1600 V with dry gas flow 3 L/min at 180 °C. The mass spectrometer was operated under parallel accumulation serial fragmentation (PASEF) mode for data-dependent acquisition using the following parameters: polarity positive, scan *m/z* range 100–1700, mobility ($1/K_0$) range 0.60–1.60 $V·s/cm^2$, ramp time 100 ms, accumulation time 2 ms, number of PASEF MS/MS scans 10, target intensity 20000, intensity threshold 2500, exclusion release after 0.4 min. Linear mobility-dependent collision energy was applied with 20 eV at 0.60 $V·s/cm^2$ and 59 eV at 1.6 $V·s/cm^2$. Data analyses were performed using FragPipe package (version 23.0) including MSFragger (version 4.3) and IonQuant (version 1.11.11). TimsTOF data files (.d) from one experiment were searched and quantified using default settings of a label-free post-translational modification method except the following changes: (1) cysteine carbamidomethylation +57.02146 (maximum 2 occurrence in a peptide) and CPT +221.08169 (maximum 2 occurrence in a peptide) were selected as variable modifications in addition to methionine oxidation and protein N-terminal acetylation; (2) PSM validation with Percolator, PTMProphet, ProteinProphet, and FDR filter at 0.01 were used in the Validation workflow; (3) data normalization across runs was not selected during quantification. Post quantification, cysteine oxidation level for each site was normalized to total CPT signal for each sample, and expressed as relative oxidation level for treated cells over DMSO control. For quantification of baseline cysteine oxidation state in monocytes of rheumatoid arthritis (RA) and multiple sclerosis (MS) patient and healthy donors, $4x10^6$ cells per donor were used for CPT-MS quantification as described previously[14].

**Screening of cysteine-reactive small molecules using CPT-MS**

For cysteine-reactive small molecules screen in iBMDMs and THP-1 MDMs, cells were plated in 10-cm dishes and allowed to adhere overnight. Cells were then lysed in 500 µl/dish PBS, tip-sonicated (QSonica Q500 Sonicator, 10 pulses, 30% amplitude for 10sec per pulse, 5 sec pause between pulses), and protein concentration quantified by bicinchoninic acid (BCA) assay (Pierce BCA protein assay kit, #23225). Lysates were split into 8 300 µg protein aliquots, and treated with 5, 10, 20 or 40 µM SCA1 or DMSO in duplicates, or 50 µM SCA9, SCA5 or SCA1-NC in replicates, for 3 h at 37°C on a shaking incubator. Samples were then subjected to methanol-chloroform precipitation and the pellets were washed twice with methanol. Following precipitation, protein pellets were re-dissolved in cysteine-reactive phosphate tag (CPT) (in-house) labelling buffer (50 mM

HEPES, pH 8.5, 2% SDS, 10 mM CPT, 5 mM TCEP) and incubated for 2 h at 37°C on a shaking incubator in the dark. Proteins were then subjected to repeated methanol-chloroform precipitation to remove CPT and the pellets were washed twice with methanol. The resultant protein pellets were reconstituted with 100 µl 200 mM EPPS buffer, pH 8 (Sigma Aldrich, #E9502), and digested with Lys-C (Wako Chemicals, #125-05061) and trypsin (Promega, #V5113) at 100:1 substrate:enzyme ratio overnight (19 h) at 37°C.

Overnight digested samples were clarified by centrifugation at 16,000 g for 10 min, and subjected to protein concentration quantification by microBCA assay (Thermo Scientific Micro BCA protein assay kit, #23235). 250 µg peptides for each sample were labelled with tandem mass tag (TMT)pro-18plex reagents (Thermo Scientific, #A44520) in 30% acetonitrile (ACN)/EPPS solution for 1 h at room temperature. TMT labelling was then quenched with 5% hydroxylamine for 15 min and samples were acidified with 1% formic acid (FA) (pH<3). A ratio check was performed by mixing 1% (vol/vol) sample from each TMT channel, and standardized TMT-labelled peptides were evenly pooled according to the ratio check. Pooled sample was then desalted by Sep-Pak C18 cartridge (Waters, #WAT054945, 200 mg). Briefly, cartridge was equilibrated with 6 ml 100% ACN, 2 ml 70% ACN/1% FA, 6 ml 1% FA, loaded with sample, washed with 6 ml 1% FA, and eluted with 0.6 ml 40% ACN/1% FA followed by 1.2 ml 70% ACN/1% FA, all steps allowed for gravity flow. Elution sample was frozen and lyophilized by speed vac. Following which, phosphate groups on endogenously phosphorylated peptides were removed through Lambda phosphatase treatment by redissolving the peptides in 500 µl phosphatase treatment buffer (50 mM HEPES, pH 7.5, 100 mM NaCl, 1 mM $MnCl_2$, 12 µl Lambda phosphatase (Santa Cruz Biotechnology, #sc-200312)), and incubated for 2 h at 30°C on a shaking incubator. Phosphatase-treated samples were acidified with 10% trifluoroacetic acid (TFA) (pH<3) desalted once more by Sep-Pak C18 cartridge (Waters, #WAT054955, 50 mg). Briefly, cartridge was equilibrated with 0.9 ml 100% ACN, 0.3 ml 70% ACN/1% FA, 0.9 ml 0.1% TFA, loaded with sample, washed with 0.9 ml 0.1% TFA, and eluted with 0.3 ml 40% ACN/1% FA followed by 0.35 ml 70% ACN/1% FA, all steps allowed for gravity flow. Elution sample was frozen and lyophilized by speed vac.

CPT-labeled peptides were then purified by IMAC using High-Select Fe-NTA Phosphopeptide Enrichment Kit (Thermo Scientific, #A32992) following manufacturer's protocol. Briefly, IMAC column storage buffer was removed by centrifugation at 1000 g for 30 sec, equilibrated with 200 µl Binding/Wash Buffer (provided in kit), centrifuged at 1000 g for 30 sec, this equilibration step repeated once, and column incubated with CPT-labelled peptides resuspended in 300 µl Binding/Wash Buffer for 30 min at room temperature on a shaking incubator. Column was centrifuged at 1000 g for 30 sec, sample washed with 200 µl Binding/Wash Buffer thrice followed by 100 µl LC-MS (liquid chromatography-mass spectrometry) grade water, and eluted with 150 µl 50 mM $HK_2PO_4$ (dibasic potassium phosphate, pH 10) twice by incubation for 1 min prior to centrifugation at 1000 g for 30 sec. Purified peptides were acidified with 10% TFA (pH<3) and desalted by Sep-Pak C18 50 mg cartridge as described above, and lyophilized by speed vac.

Peptides were fractionated with Agilent 1100 quaternary HPLC system. Peptides were separated with 57 min linear gradients from 3-32% ACN in 10 mM ammonium bicarbonate, pH 8, at 0.25 ml/min flow rate, into 96 fractions, which were then combined into 24 fractions. Fractionated and combined samples were lyophilized by speed vac and desalted using StageTip, and reconstituted in 5% FA/5% ACN for liquid chromatography tandem mass spectrometry (LC-MS/MS) analysis. Protein abundances in 2 µg peptides from each fraction were measured using Orbitrap Eclipse instrument coupled to Easy-nLC 1200 (Thermo Scientific, #LC140) ultrahigh-pressure liquid chromatography (HPLC) pump and analyzed using 180 min gradients comprising 2-23% ACN, 0.125% FA, at a flow rate of 500 nl/min. A High-Frequency Asymmetric-waveform ion mobility mass spectrometry (FAIMS)Pro (Thermo Scientific, #FMS02-10001) device was used for precursors separation and operated under default settings and compensation voltages 40V/-60V/-80V. At each voltage, peptide ions were collected in data dependent mode with a mass range of m/z 400-1600 and 2 sec cycles. MS1 resolution was set to 120,000 with standard automatic gain control target, ions were selected and subjected to fragmentation at a normalized collisional energy of 35% for MS2 with a dynamic exclusion of 120 sec. MultiNotch synchronous precursor selection (SPS)-MS3 was used for isolating MS3 precursors and MS3-based TMT quantification. Ten notches were selected and the MS3 spectra were obtained in the Orbitrap using higher-energy-collisional dissociation and normalized collisional energy of 65% as previously described[14].

MS/MS spectra were searched using the Comet algorithm[70] against a database of Mus musculus protein sequences from UniProt (http://www.uniprot.org, 2020) or homo sapiens protein sequences from UniProt (http://www.uniprot.org, 2020). Reversed peptide sequences were used as decoys for false discovery rate (FDR) filtering, and common protein contaminants including human keratins and trypsin were included. Peptides search was performed following these parameters: 25 ppm precursor mass tolerance, 1.0 Da product ion mass tolerance, full tryptic digestion, ≤3 missed cleavages, variable modifications of oxidation on methionines (+15.9949), and CPT on cysteines (+221.08169), static modifications of TMT (+304.2071) on peptide N-termini and lysines[14]. To control for FDR, the target-decoy method was used. Linear discriminant analysis was employed for peptide-level FDR control at <1% to discriminate incorrect from correct peptide identifications. Peptides with <6 amino acids were excluded. Protein-level FDR was controlled at 1%, which further decreases peptide reverse hits. Peptides were matched to the least number of sites. TMT reporter ion signal-to-noise ratios for all peptides that were matched to the same site were used for site quantification. For SCA9, SCA5 and SCA1-NC CPT-MS experiment, sites were filtered for R with coefficient of variation <0.3 across replicates.

**Isolation of human peripheral blood mononuclear cells (PBMCs)**
Peripheral blood mononuclear cells (PBMCs) were isolated from either fresh human leukopaks (for RA and MS patient donors; Stemcell Technologies) or leukocyte-enriched red blood cells (for healthy donors; Brigham and Women's Hospital). For RA/MS patient samples, leukopak contents were diluted 1:2 with wash buffer (PBS, 2% FBS, 1 mM EDTA), layered over Lymphoprep™ (Stemcell Technologies, #18060; sample:Lymphoprep = 2:1) in 50 ml conical tubes, and centrifuged at 800 g for 12 min at room temperature, brake off. The upper two layers containing the buffy coat were collected, washed once in wash buffer, and red blood cell lysis was performed using ammonium-chloride-potassium (ACK) lysing buffer (Gibco, #A1049201) for 3 min at room temperature. After quenching the lysis with 10× volume of PBS, cells were filtered through a 100 µm strainer, centrifuged, resuspended in wash buffer, counted by trypan blue exclusion, cryopreserved in Bambanker™ Freezing Medium (Nippon Genetics, #BB05), and stored at −80°C. For healthy donor samples, leukocyte-enriched red blood cells were processed using SepMate™ tubes (Stemcell Technologies, #85450) and Lymphoprep™ as per manufacturer's instructions. Briefly, cell suspensions were diluted 2:1 with wash buffer, layered over Lymphoprep in SepMate™ tubes (sample:Lymphoprep = 2:1), and centrifuged at 1200 g for 20 min at room temperature, brake on. The mononuclear cell fraction was collected, washed, subjected to ACK lysis and quenching as described above, filtered, centrifuged, resuspended in wash buffer, counted, cryopreserved in Bambanker, and stored at −80°C until further isolation and analysis.

**Isolation of human monocytes from PBMCs and flow cytometry**
Monocytes were isolated from PBMCs using the EasySep™ Human Monocyte Isolation Kit (Stemcell Technologies, #19359), according to manufacturer's protocol. Briefly, PBMCs were incubated with the antibody cocktails and magnetic particles supplied in the kit, and monocytes were separated by negative selection using the EasySep™ magnet. Monocyte purity was assessed by flow cytometry. $0.2 \times 10^6$ cells were initially incubated with human Fc receptor blocking solution for 10 min at room temperature (BioLegend, #422302; 1:20 dilution), followed by staining with Zombie-NIR fixable viability dye for 10 min at room temperature (BioLegend, # 423105; 1:1000 dilution), and then with fluorophore-conjugated antibodies against CD45 (Stemcell Technologies, #60018AZ.1), CD14 (Stemcell Technologies, #60004AD), and CD16 (Stemcell Technologies, #60041PE.1) for 15 min at room temperature. Following staining, cells were washed with PBS containing 2% FBS and 1 mM EDTA, and resuspended in the same buffer for flow cytometry analysis. Data were acquired with a Sony ID7000 analyzer and analyzed using FlowJo software (BD Biosciences). Monocytes were gated as live singlets (Zombie-$NIR^-$) and $CD45^+$ $CD14^+$ $CD16^-$ cells. The gating strategy is provided in **Extended Data Figure 7a,b**. Spectral unmixing was performed using the instrument's algorithm with single-stained UltraComp eBeads (Invitrogen, #01-2222-42) for each antibody-fluorophore, alongside unstained UltraComp eBeads as the negative control. For Zombie-NIR, PBMCs stained only with Zombie-NIR and unstained PBMCs were used as positive and negative controls, respectively.

**Isolation of mouse primary bone marrow-derived macrophages (BMDMs)**
Female C57BL/6J 7-9 weeks mice (The Jackson Laboratory) were euthanized by $CO_2$ and death confirmed by cervical dislocation. Bone marrow cells were extracted from femurs and tibiae by flushing with PBS containing 2% heat-inactivated FBS and 1% penicillin/streptomycin, passed through a 25G needle and filtered through a 100 µm cell strainer. Cells were pelleted at 500 g for 5min, resuspended with ACK lysing buffer (Gibco, #A1049201) for 3 min, quenched, filtered through a 100 µm cell strainer and pelleted at 500 g for 5 min. White blood cells pellet was washed twice and differentiated in DMEM (Corning, #10-017-CV) supplemented with 10% heat-inactivated FBS, 1% penicillin/streptomycin and 40 ng/ml macrophage colony-stimulating factor (M-CSF) (PeproTech, #315-02) for 6 days, at which time they were counted and replated in 10-cm dishes at $2x10^6$/ml with 40 ng/ml M-CSF for desthiobiotin enrichment assays. All animal-related experiments were approved by Institutional Animal Care and Use Committee of the Beth Israel Deaconess Medical Center.

**Quantification of protein abundance using TMT proteomics**
TMT proteomics was performed on SCA1 and/or LPS treated samples as described previously[74]. Briefly, treated cells were lysed in lysis buffer (50 mM HEPES pH 8.5, 2% SDS, 8 M urea, Roche EDTA-free protease inhibitor cocktail (Roche, #11836170001)), protein purified by methanol-chloroform precipitation, digested with Lys-C (Wako Chemicals, #125-05061) and trypsin (Promega, #V5113) in 200 mM EPPS buffer, pH 8 (Sigma Aldrich, #E9502) at 100:1 substrate:enzyme ratio overnight (19 h) at 37°C, and then labelled with TMTpro-16plex reagents (Thermo Scientific, #A44520), and quenched by adding 5% hydroxylamine. Samples were desalted by Sep-Pak C18 cartridge (Waters, #WAT054955, 50 mg) and fractionated with Agilent 1100 quaternary HPLC system as described above, then lyophilized and desalted using StageTip, and reconstituted in 5% FA/5% ACN for LC-MS/MS analysis and quantification as described above, except that peptides search was performed without CPT modification on cysteines.

**Supplementary Figure 20.** $^{1}H$ and $^{13}C$ NMR of SCA12……………………………………………..……52
**Supplementary Figure 21.** $^{1}H$ and $^{13}C$ NMR of SCA13……………………………………………..……53
**Supplementary Figure 22.** $^{1}H$ and $^{13}C$ NMR of SCA14……………………………………………..……54
**Supplementary Figure 23.** $^{1}H$ and $^{13}C$ NMR of SCA15……………………………………………..……55
**Supplementary Figure 24.** $^{1}H$ and $^{13}C$ NMR of SCA16……………………………………………..……56
**Supplementary Figure 25.** $^{1}H$ and $^{13}C$ NMR of SCA17……………………………………………..……57
**Supplementary Figure 26.** $^{1}H$ and $^{13}C$ NMR of SCA18……………………………………………..……58
**Supplementary Figure 27.** $^{1}H$ and $^{13}C$ NMR of SCA19..……………………………………………...59
**Supplementary Figure 28.** $^{1}H$ and $^{13}C$ NMR of SCA20..……………………………………………..60
**Supplementary Figure 29.** $^{1}H$ and $^{13}C$ NMR of SCA21……………………………………………….61
**Supplementary Figure 30.** $^{1}H$ and $^{13}C$ NMR of SCA22……………………………………………….62
**Supplementary Figure 31.** $^{1}H$ and $^{13}C$ NMR of SCA23……………………………………………….63
**Supplementary Figure 32.** $^{1}H$ and $^{13}C$ NMR of SCA24…………………………………………….....64
**Supplementary Figure 33.** $^{1}H$, $^{13}C$, and $^{19}F$ NMR of SCA25..……………………………………......65
**Supplementary Figure 34.** $^{1}H$ and $^{13}C$ NMR of SCA26..……………………………………….......67
**Supplementary Figure 35.** $^{1}H$ and $^{13}C$ NMR of Desthiobiotinylated SCA1…….………………………….68

**Chemical Synthesis:**

All reactions were conducted under standard atmospheric conditions unless otherwise specified. Reagents and solvents were purchased from commercial suppliers without further modification. Reactions were monitored by LC-MS using a Waters Acuity UPLC. Reverse phase chromatography was performed on a Teledyne ISCO CombiFlash system using C18 columns. Normal phase chromatography was performed on a Teledyne ISCO CombiFlash System using silica columns. Final compounds were purified using a Waters 2489 HPLC System. $^{1}H$, $^{19}F$, and $^{13}C$ NMR spectra were obtained on a 500 MHz Bruker A500. NMR solvents were purchased by Cambridge Isotopes Laboratories Inc. $^{1}H$ NMR signals are reported as chemical shift (ppm), multiplicity (s = singlet, d = doublet, t = triplet, q = quartet, m = multiplet, dd = doublet of doublets), integration value, and coupling constant (J) in Hz. All compounds were obtained in a purity of over 95% as assessed by LC-MS and $^{1}H$ NMR. X-ray Crystal structures were obtained at the Synchrotron Radiation Light-Source at SLAC National Accelerator Laboratory. Chemical abbreviations are as follows: DMF = *N*,*N*-Dimethylformamide; DCM = Dichloromethane; TFA = Trifluoroacetic Acid; DIPEA = N,N-Diisopropylethylamine; TEA = Triethylamine; HATU = Hexafluorophosphate Azabenzotriazole Tetramethyl Uronium; THF = Tetrahydrofuran; $NH_4Cl$ = Ammonium chloride; EtOH = Ethanol; DMSO = Dimethyl sulfoxide; NaH = Sodium hydride; NMP = N-Methyl-2-pyrrolidone; DPPA = Diphenylphosphoryl azide; $NaHCO_3$ = Sodium bicarbonate; AcOH = Acetic acid; Fe = Iron; HCl = Hydrochloric acid; t-BuOH = Tert-butanol; IPA = Isopropyl alcohol; EtOAc = Ethyl acetate; MeOH = Methanol; $H_2O$ = Water; DMAP = 4-Dimethylaminopyridine; $Na_2SO_4$ = Sodium sulfate; $PMBNH_2$ = 4-methoxybenzylamine; Pd/C = Palladium on carbon; t-BuONa = Tert-butoxide; BOP = Benzotriazol-1-yloxytris(dimethylamino)phosphonium hexafluorophosphate; NaOMe = Sodium methoxide; and $N_2$ = anhydrous Nitrogen gas.

**Synthesis of SCA1 (N-(3-((5-chloropyrazolo[1,5-a]pyrimidin-7-yl)amino)phenyl)acrylamide):**

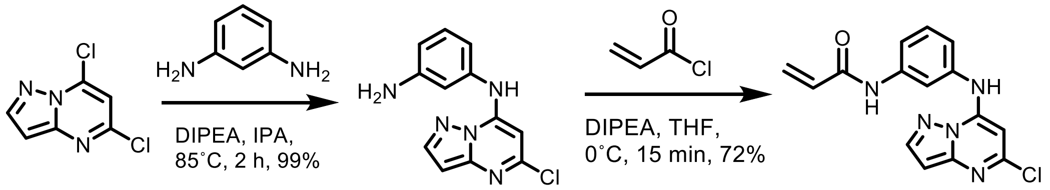

**Step 1:** To a solution of 5,7-dichloropyrazolo[1,5-a]pyrimidine (410.0 mg, 2.67 mmol, 1.0 equiv) and 5-benzene-1,3-diamine (289 mg, 2.67 mmol, 1.0 equiv) in isopropanol (8.09 mL, 0.33 M) was added DIPEA (1.86 mL, 10.7 mmol, 4.0 equiv). The reaction was allowed to stir at 85˚ C for 2 hours. The reaction was concentrated *in vacuo* and purified using normal phase purification (0-100% ethyl acetate: hexanes). The product was additionally purified by prep-HPLC (85% to 10% water (0.05% TFA) to MeOH (0.05% TFA)) to afford N1-(5-chloropyrazolo[1,5-a]pyrimidin-7-yl)benzene-1,3-diamine (595 mg, 99%, 2.64 mmol) as a white solid. LC-MS (ESI) m/z: 259.8 $[M+H]^+$. $^{1}H$ NMR (500 MHz, DMSO) δ 10.38 (s, 1H), 9.26 (s, 2H), 8.23 (d, *J* = 2.3 Hz, 1H), 7.43 – 7.35 (m, 1H), 7.17 – 7.11 (m, 2H), 7.01 – 6.95 (m, 1H), 6.52 (d, *J* = 2.2 Hz, 1H), 6.18 (s, 1H). $^{13}C$ NMR (126 MHz, DMSO) δ 150.41, 147.75, 146.11, 144.54, 140.00, 137.63, 130.55, 119.08, 117.26, 115.21, 95.51, 85.79.

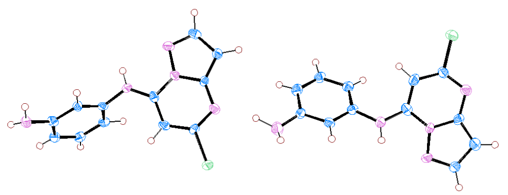

**X-ray Crystal Structure of N1-(5-chloropyrazolo[1,5-a]pyrimidin-7-yl)benzene-1,3-diamine**

**Step 2:** To a solution of N1-(5-chloropyrazolo[1,5-a]pyrimidin-7-yl)benzene-1,3-diamine (260.0 mg, 1.00 mmol, 1.0 equiv) in THF (9.94 mL, 0.02 M) was added DIPEA (0.104 mL, 0.596 mmol, 3.0 equiv). The mixture was cooled to 0˚ C and acryloyl chloride (0.0800 mL, 1.00 mmol, 1.0 equiv) was added dropwise and the reaction was allowed to stir for 15 minutes. The reaction was concentrated *in vacuo* and purified by prep-HPLC (85% to 10% water (0.05% TFA) to MeOH (0.05% TFA)) to afford N-(3-((5-chloropyrazolo[1,5-a]pyrimidin-7-yl)amino)phenyl)propionamide (226 mg, 72%, 0.0720 mmol) as a white solid. LC-MS (ESI) m/z: 314.00 $[M+H]^+$. $^1H$ NMR (500 MHz, DMSO) δ 6.72 (s, 1H), 6.65 (s, 1H), 4.57 (d, $J$ = 2.3 Hz, 1H), 4.16 (t, $J$ = 2.1 Hz, 1H), 3.92 (ddd, $J$ = 8.3, 2.1, 1.0 Hz, 1H), 3.76 (t, $J$ = 8.1 Hz, 1H), 3.51 (ddd, $J$ = 8.0, 2.3, 1.0 Hz, 1H), 2.86 (d, $J$ = 2.2 Hz, 1H), 2.78 (dd, $J$ = 17.0, 10.2 Hz, 1H), 2.61 (dd, $J$ = 17.0, 2.0 Hz, 1H), 2.50 (s, 1H), 2.11 (dd, $J$ = 10.1, 2.0 Hz, 1H). $^{13}C$ NMR (126 MHz, DMSO) δ 163.36, 150.33, 147.68, 146.18, 144.50, 139.98, 136.88, 131.70, 129.90, 127.29, 119.88, 117.24, 115.38, 95.43, 85.52.

**Synthesis of SCA1-NC (N-(3-((5-chloropyrazolo[1,5-a]pyrimidin-7-yl)amino)phenyl)propionamide):**

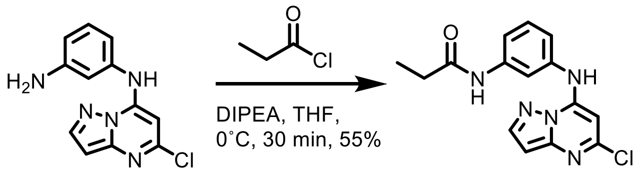

To a solution of N1-(5-chloropyrazolo[1,5-a]pyrimidin-7-yl)benzene-1,3-diamine (37.0 mg, 0.142 mmol, 1.0 equiv) in THF (7.12 mL, 0.02 M) was added DIPEA (0.0745 mL, 0.427 mmol, 3.0 equiv). The mixture was cooled to 0˚ C and a solution of propionyl chloride (0.0123 mL, 0.142 mmol, 1.0 equiv) diluted in THF (1.0 mL) was added dropwise. The reaction was concentrated *in vacuo* and purified by prep-HPLC (85% to 10% water (0.05% TFA) to MeOH (0.05% TFA)) to afford N-(3-((5-chloropyrazolo[1,5-a]pyrimidin-7-yl)amino)phenyl)propionamide (24.7 mg, 55%, 0.0782 mmol) as a white solid. LC-MS (ESI) m/z: 316.10 $[M+H]^+$. $^1H$ NMR (500 MHz, DMSO) δ 10.36 (s, 1H), 10.04 (s, 1H), 8.23 (d, $J$ = 2.2 Hz, 1H), 7.76 (t, $J$ = 2.1 Hz, 1H), 7.52 (dt, $J$ = 8.6, 1.2 Hz, 1H), 7.39 (t, $J$ = 8.0 Hz, 1H), 7.13 (ddd, $J$ = 8.0, 2.1, 1.0 Hz, 1H), 6.53 (d, $J$ = 2.2 Hz, 1H), 6.14 (s, 1H), 2.35 (q, $J$ = 7.5 Hz, 2H), 1.09 (t, $J$ = 7.5 Hz, 3H). $^{13}C$ NMR (126 MHz, DMSO) δ 172.25, 150.32, 147.68, 146.22, 144.48, 140.36, 136.76, 129.77, 119.33, 116.91, 115.08, 95.41, 85.47, 29.55, 9.58.

**Synthesis of SCA2 (N-(4-(5-chloropyrazolo[1,5-a]pyrimidin-7-ylamino)phenyl)acrylamide):**

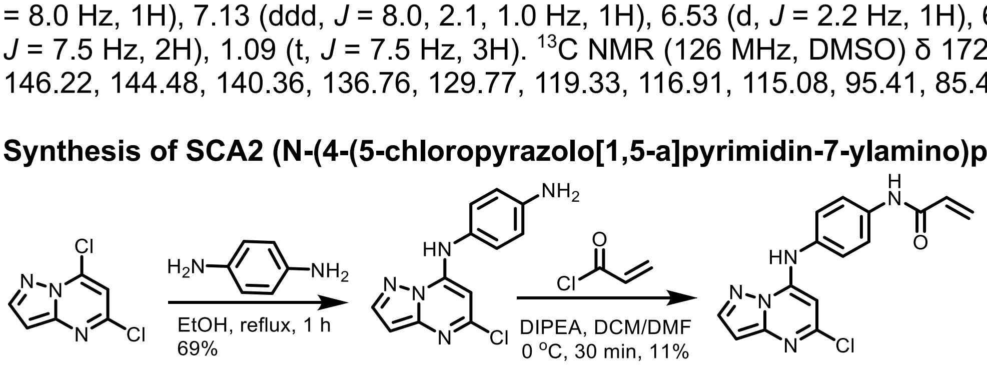

**Step 1:** A mixture of 5,7-dichloropyrazolo[1,5-a]pyrimidine (940.0 mg, 5.00 mmol, 1.0 equiv) and benzene-1,4-diamine (540.0 mg, 5.00 mmol, 1.0 equiv) in EtOH (20.0 mL, 0.25 M) was stirred at reflux for 2 h. After cooling to room temperature, the resulting suspension was filtered and the

cake was washed with cold EtOH (2.0 mL x 2) and dried to afford N1-(5-chloropyrazolo[1,5-a]pyrimidin-7-yl)benzene-1,4-diamine as a yellow solid (894 mg, 69%, 3.44 mmol). LC-MS (ESI) m/z: 260 $[M+H]^+$.

**Step 2:** To a solution of N1-(5-chloropyrazolo[1,5-a]pyrimidin-7-yl)benzene-1,4-diamine (259 mg, 1.00 mmol, 1.0 equiv) in DMF (4.00 ml, 0.25 M) was added DIPEA (258 mg, 2.00 mmol, 2.0 equiv). A solution of acryloyl chloride (270.0 mg, 3.00 mmol, 3.0 equiv) in DCM (2.00 ml, 0.5 M) was added dropwise at 0˚ C. The mixture was stirred at 0˚ C for 30 min. After removal of the solvent, the residue was purified with reverse phase chromatography (acetonitrile/0.5% $NH_3.H_2O$, 0:100%) to yield the crude product, which was triturated with DCM (1.5 mL) and dried under vacuum to afford compound (N-(4-(5-chloropyrazolo[1,5-a]pyrimidin-7-ylamino)phenyl)acrylamide) as a yellow solid (35.0 mg, 11%, 0.112 mmol). LC-MS (ESI) m/z: 314.05 $[M+H]^+$. $^1$H NMR (500 MHz, DMSO) δ 10.34 (s, 1H), 10.29 (s, 1H), 8.23 (d, *J* = 2.3 Hz, 1H), 7.82 – 7.75 (m, 2H), 7.45 – 7.39 (m, 2H), 6.52 (d, *J* = 2.3 Hz, 1H), 6.47 (dd, *J* = 16.9, 10.1 Hz, 1H), 6.29 (dd, *J* = 17.0, 2.0 Hz, 1H), 6.03 (s, 1H), 5.78 (dd, *J* = 10.1, 2.0 Hz, 1H). $^{13}$C NMR (126 MHz, DMSO) δ 163.20, 150.34, 147.67, 146.53, 144.48, 137.44, 131.77, 131.59, 127.07, 125.64 (2C), 120.24 (2C), 95.30, 85.27.

**Synthesis of SCA3 (N-(3-((5-chloropyrazolo[1,5-a]pyrimidin-7-yl)amino)-2-methylphenyl)acrylamide):**

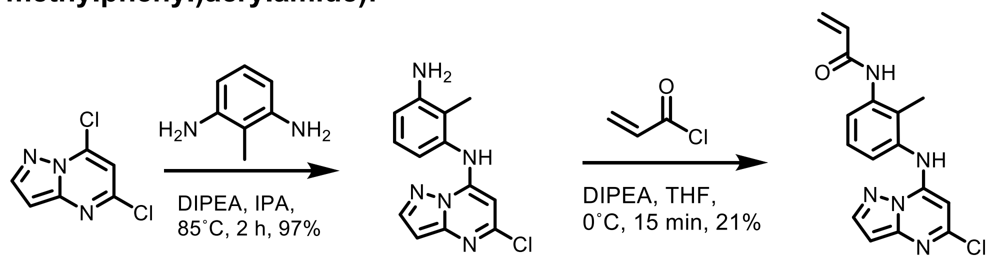

**Step 1:** To a solution of 5,7-dichloropyrazolo[1,5-a]pyrimidine (500.0 mg, 2.66 mmol, 1.0 equiv) and 2-methylbenzene-1,3-diamine (325 mg, 2.66 mmol, 1.0 equiv) in isopropanol (8.06 mL, 0.33 M) was added DIPEA (1.85 mL, 10.6 mmol, 4.0 equiv). The reaction was allowed to stir at 85˚ C for 2 hours. The reaction was concentrated *in vacuo* and purified through normal phase chromatography (0-100% ethyl acetate: hexane) to afford N1-(5-chloropyrazolo[1,5-a]pyrimidin-7-yl)-2-methylbenzene-1,3-diamine as a beige solid (701 mg, 96%, 2.66 mmol). LC-MS (ESI) m/z: 274.13 $[M+H]^+$.

**Step 2:** To a solution of N1-(5-chloropyrazolo[1,5-a]pyrimidin-7-yl)-2-methylbenzene-1,3-diamine (45.0 mg, 0.160 mmol, 1.0 equiv) in THF (8.20 mL, 0.05 M) was added DIPEA (0.0860 mL, 0.490 mmol, 3.0 equiv). The mixture was cooled to 0˚ C and acryloyl chloride (0.0130 mL, 0.160 mmol, 1.0 equiv) was added dropwise and the reaction was allowed to stir for 15 minutes. The reaction was concentrated *in vacuo* and purified by prep-HPLC (85% to 10% water (0.05% TFA) to MeOH (0.05% TFA)) to afford N-(3-((5-chloropyrazolo[1,5-a]pyrimidin-7-yl)amino)-2-methylphenyl)acrylamide (11.5 mg, 21%, 0.160 mmol) as a white solid. LC-MS (ESI) m/z: 328.00 $[M+H]^+$. $^1$H NMR (500 MHz, DMSO) δ 10.24 (s, 1H), 9.70 (s, 1H), 8.24 (d, *J* = 2.3 Hz, 1H), 7.56 (d, *J* = 8.0 Hz, 1H), 7.34 (t, *J* = 7.9 Hz, 1H), 7.25 (dd, *J* = 7.9, 1.3 Hz, 1H), 6.60 – 6.54 (m, 1H), 6.52 (d, *J* = 2.2 Hz, 1H), 6.28 (dd, *J* = 17.1, 2.0 Hz, 1H), 5.77 (dd, *J* = 10.2, 2.0 Hz, 1H), 5.47 (s, 1H), 2.09 (s, 3H). $^{13}$C NMR (126 MHz, DMSO) δ 163.39, 150.23, 147.67, 147.10, 144.62, 137.53, 135.25, 131.57, 130.16, 126.93, 126.57, 124.97, 95.25, 84.98, 12.77.

**Synthesis of SCA4 (N-(3-(5-chloropyrazolo[1,5-a]pyrimidin-7-ylamino)-4-methylphenyl)acrylamide):**

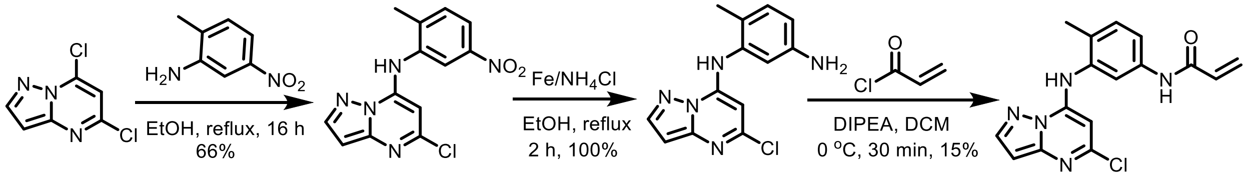

**Step 1:** A mixture of 5,7-dichloropyrazolo[1,5-a]pyrimidine (470.0 mg, 2.50 mmol, 1.0 equiv) and 2-methyl-5-nitroaniline (380.0 mg, 2.50 mmol, 1.0 equiv) in EtOH (10.0 mL, 0.25 M) was stirred at reflux for 2 h. After cooling to room temperature, the precipitate was collected by filtration, washed with EtOH (2.0 mL x 2) and dried under vacuum to afford 5-chloro-N-(2-methyl-5-nitrophenyl)pyrazolo[1,5-a]pyrimidin-7-amine as a yellow solid (500.0 mg, 66%, 1.64 mmol). LC-MS (ESI) m/z: 304 $[M+H]^+$.

**Step 2:** To a solution of 5-chloro-N-(2-methyl-5-nitrophenyl)pyrazolo[1,5-a]pyrimidin-7-amine (500.0 mg, 1.65 mmol, 1.0 equiv) in EtOH (15.0 mL, 0.11 M) and water (1.00 mL, 1.65 M) was added Fe powder (924 mg, 16.5 mmol, 10.0 equiv) and $NH_4Cl$ (441 mg, 8.25 mmol, 5.0 equiv). The mixture was stirred at reflux overnight. After cooling to room temperature, the mixture was filtered through Celite and washed with EtOH (10 mL x 2). The filtrate was evaporated to afford N1-(5-chloropyrazolo[1,5-a]pyrimidin-7-yl)-6-methylbenzene-1,3-diamine as a yellow solid (450.0 mg, 100%, 1.65 mmol). LC-MS (ESI) m/z: 274 $[M+H]^+$.

**Step 3:** To a solution of N1-(5-chloropyrazolo[1,5-a]pyrimidin-7-yl)-6-methylbenzene-1,3-diamine (450.0 mg, 1.65 mmol, 1.0 equiv) in DCM (10.0 ml, 0.165 M) was added DIPEA (1.06 mL, 6.60 mmol, 4.0 equiv). A solution of acryloyl chloride (0.265 mL, 3.30 mmol, 2.0 equiv) in DCM (1.00 ml, 1.65 M) was added dropwise at 0° C. The mixture was stirred at 0° C for 30 min. After removal of the solvent, the residue was purified with prep-HPLC (85% to 10% water to MeOH) to afford N-(3-(5-chloropyrazolo[1,5-a]pyrimidin-7-ylamino)-4-methylphenyl)acrylamide as a white solid (80.0 mg, 15%, 0.244 mmol). LC-MS (ESI) m/z: 328.10 $[M+H]^+$. $^1$H NMR (500 MHz, DMSO) δ 10.25 (s, 1H), 8.21 (d, *J* = 2.2 Hz, 1H), 7.70 (d, *J* = 2.2 Hz, 1H), 7.57 (dd, *J* = 8.2, 2.3 Hz, 1H), 7.34 (d, *J* = 8.4 Hz, 1H), 6.49 (d, *J* = 2.3 Hz, 1H), 6.43 (dd, *J* = 17.0, 10.1 Hz, 1H), 6.25 (dd, *J* = 17.0, 2.0 Hz, 1H), 5.76 (dd, *J* = 10.1, 2.0 Hz, 1H), 5.54 (s, 1H), 2.16 (s, 3H). $^{13}$C NMR (126 MHz, DMSO) δ 163.18, 150.26, 147.76, 146.96, 144.47, 137.94, 135.38, 131.74, 131.38, 129.96, 127.03, 118.61, 118.06, 95.12, 84.76, 16.80.

**Synthesis of SCA5 (N-(3-(5-chloropyrazolo[1,5-a]pyrimidin-7-ylamino)-5-methylphenyl)acrylamide):**

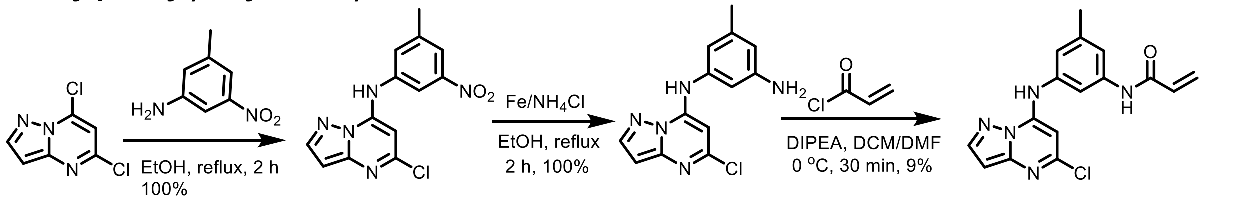

**Step 1:** A mixture of 5,7-dichloropyrazolo[1,5-a]pyrimidine (470.0 mg, 2.50 mmol, 1.0 equiv) and 3-methyl-5-nitroaniline (380.0 mg, 2.50 mmol, 1.0 equiv) in EtOH (10.0 mL, 0.25 M) was stirred at reflux for 2 h. After cooling to room temperature, the precipitate was collected by filtration, washed with EtOH (2.0 mL x 2) and dried under vacuum to afford 5-chloro-N-(3-methyl-5-nitrophenyl)pyrazolo[1,5-a]pyrimidin-7-amine as a yellow solid (758 mg, 100%, 2.49 mmol). LC-MS (ESI) m/z: 304 $[M+H]^+$.

**Step 2:** To a solution of 5-chloro-N-(3-methyl-5-nitrophenyl)pyrazolo[1,5-a]pyrimidin-7-amine (758 mg, 2.50 mmol, 1.0 equiv) in EtOH (20.0 mL, 0.25 M) and water (1.00 mL, 2.5 M) was added Fe powder (1.40 g, 25.0 mmol, 10.0 equiv) and $NH_4Cl$ (669 mg, 12.5 mmol, 5.0 equiv). The mixture was stirred at reflux for 2 h. After cooling to room temperature, the mixture was filtered through Celite and washed with EtOH (10 mL x 2). The combined filtrate and wash was evaporated to afford N1-(5-chloropyrazolo[1,5-a]pyrimidin-7-yl)-5-methylbenzene-1,3-diamine as a yellow solid (683 mg, 100%, 2.50 mmol). LC-MS (ESI) m/z: 274 $[M+H]^+$.

**Step 3:** To a solution of N1-(5-chloropyrazolo[1,5-a]pyrimidin-7-yl)-5-methylbenzene-1,3-diamine (683 mg, 2.50 mmol, 1.0 equiv) in DCM (10.0 ml, 0.25 M) was added DIPEA (1.61 mL, 10.0 mmol, 4.0 equiv). A solution of acryloyl chloride (0.402 mL, 5.00 mmol, 2.0 equiv) in DCM (2.0 ml, 1.25 M) was added dropwise at 0˚ C. The mixture was stirred at 0˚ C for 30 min. After removal of the solvent, the residue was purified by prep-HPLC (85% to 10% water to MeOH) to afford N-(3-(5-chloropyrazolo[1,5-a]pyrimidin-7-ylamino)-5-methylphenyl)acrylamide as a white solid (71.0 mg, 9%, 0.216 mmol). LC-MS (ESI) m/z: 328.05 $[M+H]^+$. $^1H$ NMR (500 MHz, DMSO) δ 10.29 (s, 1H), 10.25 (s, 1H), 8.23 (d, *J* = 2.2 Hz, 1H), 7.61 (d, *J* = 2.1 Hz, 1H), 7.44 (d, *J* = 1.9 Hz, 1H), 7.02 – 6.97 (m, 1H), 6.53 (d, *J* = 2.2 Hz, 1H), 6.44 (dd, *J* = 16.9, 10.1 Hz, 1H), 6.27 (dd, *J* = 17.0, 2.0 Hz, 1H), 6.14 (s, 1H), 5.77 (dd, *J* = 10.1, 2.0 Hz, 1H), 2.33 (s, 3H). $^{13}C$ NMR (126 MHz, DMSO) δ 163.31, 150.32, 147.67, 146.27, 144.49, 139.81, 139.49, 136.70, 131.75, 127.19, 120.58, 117.89, 112.79, 95.39, 85.52, 21.22.

**Synthesis of SCA6 (N-(3-((5-chloropyrazolo[1,5-a]pyrimidin-7-yl)amino)-5-methoxyphenyl)acrylamide):**

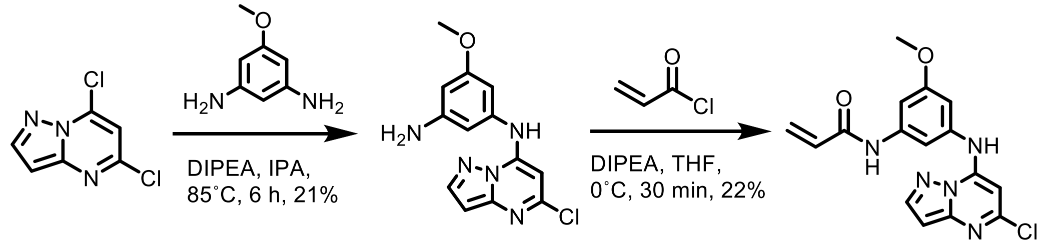

**Step 1:** To a solution of 5,7-dichloropyrazolo[1,5-a]pyrimidine (100.0 mg, 0.53 mmol, 1.0 equiv) and 5-methoxybenzene-1,3-diamine (73.0 mg, 0.53 mmol, 1.0 equiv) in isopropanol (2.00 mL, 0.265 M) was added DIPEA (0.276 mL, 1.59 mmol, 3.0 equiv). The reaction was allowed to stir at 85˚ C for 6 hours. The reaction was concentrated *in vacuo* and purified using normal phase purification (0-100% ethyl acetate: hexanes) to afford N1-(5-chloropyrazolo[1,5-a]pyrimidin-7-yl)-5-methoxybenzene-1,3-diamine as a brown solid (33.0 mg, 21%, 0.114 mmol). LC-MS (ESI) m/z: 290.10$[M+H]^+$.

**Step 2:** To a solution of N1-(5-chloropyrazolo[1,5-a]pyrimidin-7-yl)-5-methoxybenzene-1,3-diamine (18.0 mg, 0.0620 mmol, 1.0 equiv) in THF (3.1 mL, 0.02 M) was added DIPEA (0.0320 mL, 0.190 mmol, 3.0 equiv). The mixture was cooled to 0˚ C and a solution of acryloyl chloride (0.005 mL, 0.062 mmol, 2.0 equiv) in THF (0.5 mL) was added dropwise. The reaction was concentrated *in vacuo* and purified by prep-HPLC (85% to 10% water (0.05% TFA) to MeOH (0.05% TFA)) to afford N-(3-((5-chloropyrazolo[1,5-a]pyrimidin-7-yl)amino)-5-methoxyphenyl)acrylamide (5.01 mg, 23%, 0.025 mmol) as a white solid. LC-MS (ESI) m/z: 344.06 $[M+H]^+$. $^1H$ NMR (500 MHz, DMSO) δ 10.33 (s, 1H), 10.31 (s, 1H), 8.24 (d, *J* = 2.2 Hz, 1H), 7.37 (t, *J* = 1.8 Hz, 1H), 7.32 (t, *J* = 2.0 Hz, 1H), 6.80 (t, *J* = 2.1 Hz, 1H), 6.54 (d, *J* = 2.2 Hz, 1H), 6.44 (dd, *J* = 16.9, 10.1 Hz, 1H), 6.28 (dd, *J* = 16.9, 2.0 Hz, 1H), 6.24 (s, 1H), 5.78 (dd, *J* =

10.0, 2.0 Hz, 1H). $^{13}C$ NMR (126 MHz, DMSO) δ 163.41, 160.22, 150.33, 147.66, 146.04, 144.50, 140.74, 137.77, 131.69, 127.38, 107.45, 105.80, 102.88, 95.46, 85.89, 55.31.

**Synthesis of SCA7 (N-(3-bromo-5-((5-chloropyrazolo[1,5-a]pyrimidin-7-yl)amino)phenyl)acrylamide):**

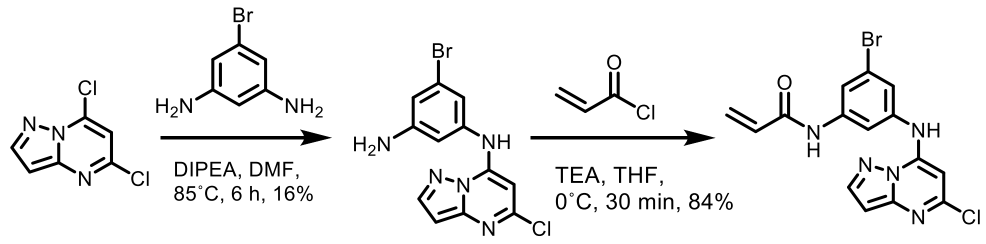

**Step 1:** To a solution of 5,7-dichloropyrazolo[1,5-a]pyrimidine (930.0 mg, 4.95 mmol, 1.0 equiv) and 5-bromobenzene-1,3-diamine (925.0 mg, 4.95 mmol, 1.0 equiv) in isopropanol (49.5 mL, 0.1 M) was added DIPEA (2.58 mL, 14.8 mmol, 3.0 equiv). The reaction was allowed to stir at 85˚ C for 6 hours. The reaction was concentrated *in vacuo* and purified using normal phase purification (0-100% ethyl acetate: hexanes) to afford 5-bromo-N1-(5-chloropyrazolo[1,5-a]pyrimidin-7-yl)benzene-1,3-diamine as a brown solid (267 mg, 16%, 0.789 mmol). LC-MS (ESI) m/z: 339.91 $[M+H]^+$. $^1H$ NMR (500 MHz, DMSO) δ 10.20 (s, 1H), 8.21 (d, *J* = 2.2 Hz, 1H), 6.73 (t, *J* = 1.8 Hz, 1H), 6.69 (t, *J* = 1.9 Hz, 1H), 6.64 (t, *J* = 1.9 Hz, 1H), 6.51 (d, *J* = 2.3 Hz, 1H), 6.15 (s, 1H), 5.67 (s, 2H). $^{13}C$ NMR (126 MHz, DMSO) δ 151.24, 150.24, 147.66, 146.05, 144.45, 138.58, 122.29, 113.95 (2C), 108.33, 95.41, 85.95.

**Step 2:** To a solution of 5-bromo-N1-(5-chloropyrazolo[1,5-a]pyrimidin-7-yl)benzene-1,3-diamine (100.0 mg, 0.295 mmol, 1.0 equiv) in THF (14.8 mL, 0.02 M) was added DIPEA (0.154 mL, 0.886 mmol, 3.0 equiv). The mixture was cooled to 0˚ C and a 1% solution of acryloyl chloride in THF (1.19 mL, 0.148 mmol, 0.5 equiv) was added dropwise. 15 minutes later an additional 0.6 equiv of the 1% acryloyl chloride DCM solution (1.43 mL, 0.177 mmol, 0.6 equiv) was added dropwise and the reaction was allowed to stir for an additional 15 minutes. The reaction was concentrated *in vacuo* and purified by prep-HPLC (85% to 10% water (0.05% TFA): MeOH (0.05% TFA)) to afford N-(3-bromo-5-((5-chloropyrazolo[1,5-a]pyrimidin-7-yl)amino)phenyl)acrylamide as a white solid (97.4 mg, 84%, 0.241 mmol). LC-MS (ESI) m/z: 393.98 $[M+H]^+$. $^1H$ NMR (500 MHz, DMSO) δ 10.47 (s, 1H), 10.45 (s, 1H), 8.25 (d, *J* = 2.3 Hz, 1H), 7.93 (t, *J* = 1.8 Hz, 1H), 7.71 (t, *J* = 1.9 Hz, 1H), 7.40 (t, *J* = 1.8 Hz, 1H), 6.56 (d, *J* = 2.2 Hz, 1H), 6.42 (dd, *J* = 16.9, 10.1 Hz, 1H), 6.30 (dd, *J* = 17.0, 2.0 Hz, 1H), 6.30 (s, 1H), 5.82 (dd, *J* = 10.0, 2.0 Hz, 1H). $^{13}C$ NMR (126 MHz, DMSO) δ 163.60, 150.35, 147.65, 145.76, 144.55, 141.13, 138.64, 131.37, 127.96, 121.99, 121.96, 119.19, 113.65, 95.58, 86.18.

**Synthesis of SCA8 (N-(5-((5-chloropyrazolo[1,5-a]pyrimidin-7-yl)amino)pyridin-3-yl)acrylamide):**

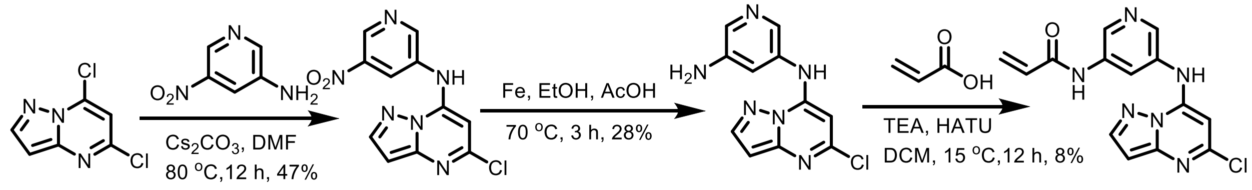

T1

**Step 1:** A mixture of 5,7-dichloropyrazolo[1,5-a]pyrimidine (1.00 g, 5.32 mmol, 1.2 equiv), 5-nitropyridin-3-amine (617 mg, 4.43 mmol, 1.0 equiv), and $Cs_2CO_3$ (2.89 g, 8.86 mmol, 2.0 equiv) in DMF (20.0 mL, 0.22 M) was degassed and purged with $N_2$ for 3 times, and then the mixture was stirred at 80 °C for 12 hours under a $N_2$ atmosphere. The reaction mixture was diluted with $H_2O$ (20 mL) and extracted with EtOAc (30 mL * 2). The combined organic layers were dried over $Na_2SO_4$, filtered, and the filtrate was concentrated under reduced pressure. The resulting residue was purified by column chromatography ($SiO_2$, Petroleum ether/Ethyl acetate=100/1 to 0/1). 5-chloro-N-(5-nitro-3-pyridyl)pyrazolo[1,5-a]pyrimidin-7-amine (600.0 mg, 47%, 2.06 mmol) was obtained as a yellow solid. LC-MS (ESI) m/z = 291.09 $[M+H]^+$. $^1H$ NMR (500 MHz, DMSO) δ 10.76 (s, 1H), 9.23 (d, *J* = 2.3 Hz, 1H), 9.08 (d, *J* = 2.3 Hz, 1H), 8.69 (t, *J* = 2.3 Hz, 1H), 8.28 (d, *J* = 2.3 Hz, 1H), 6.58 (d, *J* = 2.3 Hz, 1H), 6.55 (s, 1H). $^{13}C$ NMR (126 MHz, DMSO) δ 150.83, 150.54, 147.60, 145.65, 144.67, 144.54, 140.99, 134.63, 126.39, 95.71, 86.95.

**Step 2:** A mixture of 5-chloro-N-(5-nitro-3-pyridyl)pyrazolo[1,5-a]pyrimidin-7-amine (600.0 mg, 2.06 mmol, 1.0 equiv), Fe (576 mg, 10.3 mmol, 5.0 equiv) in EtOH (5.00 mL, 0.41 M) and AcOH (5.00 mL, 0.412 M) was degassed and purged with $N_2$ for 3 times, and then the mixture was stirred at 70 °C for 3 hours under $N_2$ atmosphere. The reaction mixture was filtered and the filtrate was concentrated under reduced pressure to yield a residue. The residue was diluted with $H_2O$ (20 mL) and extracted with DCM 60 mL (30 mL * 2). The combined organic layers were dried over $Na_2SO_4$, filtered, and the filtrate was concentrated under reduced pressure. The resulting residue was purified by column chromatography ($SiO_2$, Petroleum ether/Ethyl acetate=100/1 to 0/1). N5-(5-chloropyrazolo[1,5-a]pyrimidin-7-yl)pyridine-3,5-diamine (150.0 mg, 28%, 575 µmol) was obtained as a yellow solid. LC-MS (ESI) m/z = 261.13 $[M+H]^+$. $^1H$ NMR (500 MHz, DMSO) δ 10.27 (s, 1H), 8.23 (d, *J* = 2.2 Hz, 1H), 7.89 (d, *J* = 2.4 Hz, 1H), 7.84 (d, *J* = 2.3 Hz, 1H), 7.01 (t, *J* = 2.3 Hz, 1H), 6.52 (d, *J* = 2.2 Hz, 1H), 6.10 (s, 1H), 5.59 (s, 2H). $^{13}C$ NMR (126 MHz, DMSO) δ 150.32, 147.67, 146.36, 145.50, 144.53, 134.35, 133.52, 133.13, 115.38, 95.42, 85.68.

**Step 3:** A mixture of N5-(5-chloropyrazolo[1,5-a]pyrimidin-7-yl)pyridine-3,5-diamine (220.0 mg, 844 µmol, 1.0 equiv), acrylic acid (0.0598 mL, 928 µmol, 1.1 equiv), TEA (0.235, 1.69 mmol, 2.0 equiv), and HATU (481 mg, 1.27 mmol, 1.5 equiv) in DCM (10.0 mL, 0.0844 M) was degassed and purged with $N_2$ for 3 times, and then the mixture was stirred at 15 °C for 12 hours under a $N_2$ atmosphere. The reaction mixture was concentrated under reduced pressure. The resulting residue was purified by prep-HPLC (85% to 10% water (0.05% TFA) to MeOH (0.05% TFA)) to afford SCA8 (20.0 mg, 8%, 63.6 µmol) as a white solid. LC-MS (ESI) m/z = 315.05 $[M+H]^+$. $^1H$ NMR (500 MHz, DMSO) δ 10.55 (s, 1H), 10.53 (s, 1H), 8.74 (d, *J* = 2.2 Hz, 1H), 8.43 (d, *J* = 2.3 Hz, 1H), 8.27 (d, *J* = 2.3 Hz, 1H), 8.25 (t, *J* = 2.3 Hz, 1H), 6.57 (d, *J* = 2.2 Hz, 1H), 6.46 (dd, *J* = 17.0, 10.1 Hz, 1H), 6.32 (dd, *J* = 17.0, 1.9 Hz, 1H), 6.27 (s, 1H), 5.84 (dd, *J* = 10.1, 1.9 Hz, 1H). $^{13}C$ NMR (126 MHz, DMSO) δ 163.88, 150.41, 147.64, 145.96, 144.65, 139.96, 137.47, 136.25, 133.75, 131.13, 128.16, 122.31, 95.63, 86.08.

**Synthesis of SCA9 (N-(2-((5-chloropyrazolo[1,5-a]pyrimidin-7-yl)amino)pyridin-4-yl)acrylamide):**

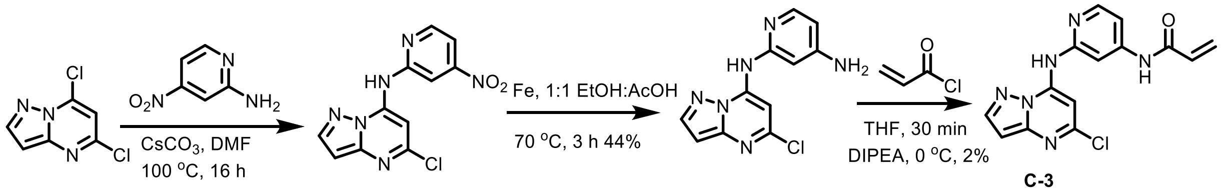

**Step 1:** To a solution of 5,7-dichloropyrazolo[1,5-a]pyrimidine (2.10 g, 11.3 mmol, 1.0 equiv) and 4-nitropyridin-2-amine (1.87 g, 13.4 mmol, 1.2 equiv) in DMF (55.9 mL, 0.2 M) was added cesium carbonate (10.9 g, 33.5 mmol, 3.0 equiv) and the reaction was allowed to stir at 100 ˚C for 16 hours. Upon cooling, the reaction was diluted with 30 mL of water and the product was extracted into the organic layer using 3 x 30 mL ethyl acetate. The organic layer was washed with brine, dried over $Na_2SO4$, filtered, concentrated, and purified through normal phase liquid-liquid chromatography (0-100% ethyl acetate: hexane) to afford 5-chloro-N-(4-nitropyridin-2-yl)pyrazolo[1,5-a]pyrimidin-7-amine as a yellow solid (2.53 g, 77%, 8.69 mmol). LC-MS (ESI) m/z: 291.04 $[M+H]^+$. $^1H$ NMR (500 MHz, DMSO) δ 11.31 (s, 1H), 8.78 (dd, $J$ = 5.5, 0.6 Hz, 1H), 8.55 (dd, $J$ = 2.1, 0.6 Hz, 1H), 8.32 (d, $J$ = 2.2 Hz, 1H), 8.03 (s, 1H), 7.82 (dd, $J$ = 5.5, 2.0 Hz, 1H), 6.66 (d, $J$ = 2.3 Hz, 1H). $^{13}C$ NMR (126 MHz, DMSO) δ 154.96, 154.32, 150.49, 150.19, 147.25, 144.33, 142.40, 110.99, 108.13, 96.23, 92.14.

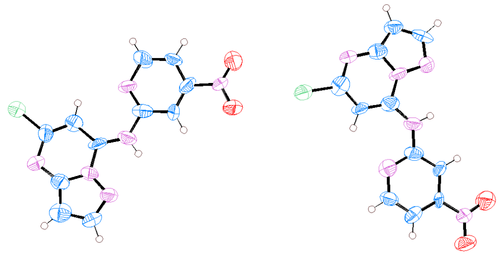

**X-Ray Crystal Structure of 5-chloro-N-(4-nitropyridin-2-yl)pyrazolo[1,5-a]pyrimidin-7-amine**

**Step 2:** To a solution of 5-chloro-N-(4-nitropyridin-2-yl)pyrazolo[1,5-a]pyrimidin-7-amine (400.0 mg, 1.38 mmol, 1.0 equiv) in EtOH (4.17 mL, 0.33 M) and acetic acid (4.17 mL, 0.33 M) was added Fe powder (769 mg, 13.8 mmol, 10.0 equiv). The mixture was stirred at 70 ˚C for three hours. After cooling to room temperature, the mixture was filtered through Celite and the filtrate was concentrated to dryness *in vacuo.* The residue was purified via reverse phase chromatography (0:100% water: acetonitrile) to afford N2-(5-chloropyrazolo[1,5-a]pyrimidin-7-yl)pyridine-2,4-diamine as a brown solid (158 mg, 44%, 0.606 mmol). LC-MS (ESI) m/z: 261.08 $[M+H]^+$. $^1H$ NMR (500 MHz, DMSO) δ 8.30 (d, $J$ = 2.1 Hz, 1H), 7.99 (d, $J$ = 6.8 Hz, 1H), 7.81 (s, 2H), 6.95 (s, 1H), 6.78 (d, $J$ = 2.1 Hz, 1H), 6.64 (d, $J$ = 2.1 Hz, 1H), 6.62 (dd, $J$ = 6.8, 2.1 Hz, 1H). $^{13}C$ NMR (126 MHz, DMSO) δ 160.14, 150.28, 147.48, 146.13, 144.83, 143.87, 140.94, 106.67, 100.10, 96.10, 90.39.

**Step 3:** To a solution of N2-(5-chloropyrazolo[1,5-a]pyrimidin-7-yl)pyridine-2,4-diamine (245 mg, 0.937 mmol, 1.0 equiv) in THF (4.0 ml, 0.23 M) was added DIPEA (0.303 mL, 1.88 mmol, 2.0 equiv). A solution of acryloyl chloride (0.0903 mL, 1.10 mmol, 1.2 equiv) in THF (1.00 ml) was added dropwise at 0˚ C. The mixture was stirred at 0˚ C for 30 minutes, concentrated, and purified by prep-HPLC ((85% to 10% water (0.05% TFA) to MeOH (0.05% TFA)) to afford (N-(2-((5-chloropyrazolo[1,5-a]pyrimidin-7-yl)amino)pyridin-4-yl)acrylamide) as a white solid (5.00 mg, 2%, 0.0159 mmol). LC-MS (ESI) m/z: 315.05 $[M+H]^+$. $^1H$ NMR (500 MHz, DMSO) δ 10.75 (s, 1H),

10.61 (s, 1H), 8.33 (d, *J* = 5.7 Hz, 1H), 8.27 (d, *J* = 2.3 Hz, 1H), 7.99 (d, *J* = 1.8 Hz, 1H), 7.94 (s, 1H), 7.37 (dd, *J* = 5.7, 1.8 Hz, 1H), 6.60 (d, *J* = 2.3 Hz, 1H), 6.49 (dd, *J* = 17.0, 10.1 Hz, 1H), 6.34 (dd, *J* = 17.0, 1.9 Hz, 1H), 5.86 (dd, *J* = 10.1, 1.8 Hz, 1H). $^{13}$C NMR (126 MHz, DMSO) δ 164.08, 153.25, 150.40, 148.08, 147.37, 147.34, 144.11, 143.18, 131.23, 128.52, 109.88, 104.09, 95.80, 91.11.

**Synthesis of SCA9-NC (N-(2-((5-chloropyrazolo[1,5-a]pyrimidin-7-yl)amino)pyridin-4-yl) propionamide):**

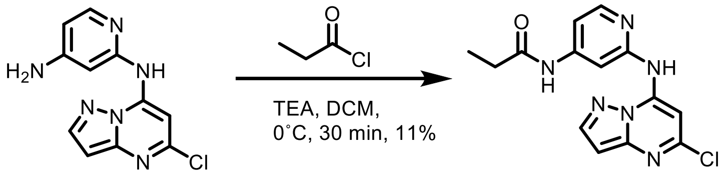

**Step 1:** To a solution of N2-(5-chloropyrazolo[1,5-a]pyrimidin-7-yl)pyridine-2,4-diamine (74.0 mg, 0.280 mmol, 1.0 equiv) dissolved in DCM (14.0 mL, 0.02 M) was added DIPEA (0.150 mL, 0.850 mmol, 3.0 equiv) and the reaction was brought to 0˚ C on ice. Then a 1% solution of propionyl chloride dissolved in DCM was added dropwise (0.015 mL, 0.28 mmol, 1.0 equiv) and the reaction was allowed to stir for 15 minutes. The reaction was concentrated and purified by prep-HPLC (85% to 10% water (0.05% TFA) to MeOH (0.05% TFA)) to afford N-(2-((5-chloropyrazolo[1,5-a]pyrimidin-7-yl)amino)pyridin-4-yl)propionamide (9.59 mg, 11%, 0.0303 mmol) as a white solid. LC-MS (ESI) m/z: 317.05 [M+H]$^+$. $^1$H NMR (500 MHz, DMSO) δ 10.68 (s, 1H), 10.35 (s, 1H), 8.31 – 8.24 (m, 2H), 7.92 (d, *J* = 2.7 Hz, 2H), 7.29 (d, *J* = 5.2 Hz, 1H), 6.59 (s, 1H), 2.39 (t, *J* = 7.4 Hz, 2H), 1.09 (t, *J* = 7.3 Hz, 3H). $^{13}$C NMR (126 MHz, DMSO) δ 173.19, 153.13, 150.37, 147.89, 147.62, 147.30, 144.06, 143.17, 109.56, 103.64, 95.74, 91.01, 29.61, 9.24.

**SCA10 (N-(5-((5-chloropyrazolo[1,5-a]pyrimidin-7-yl)amino)thiophen-3-yl)acrylamide):**

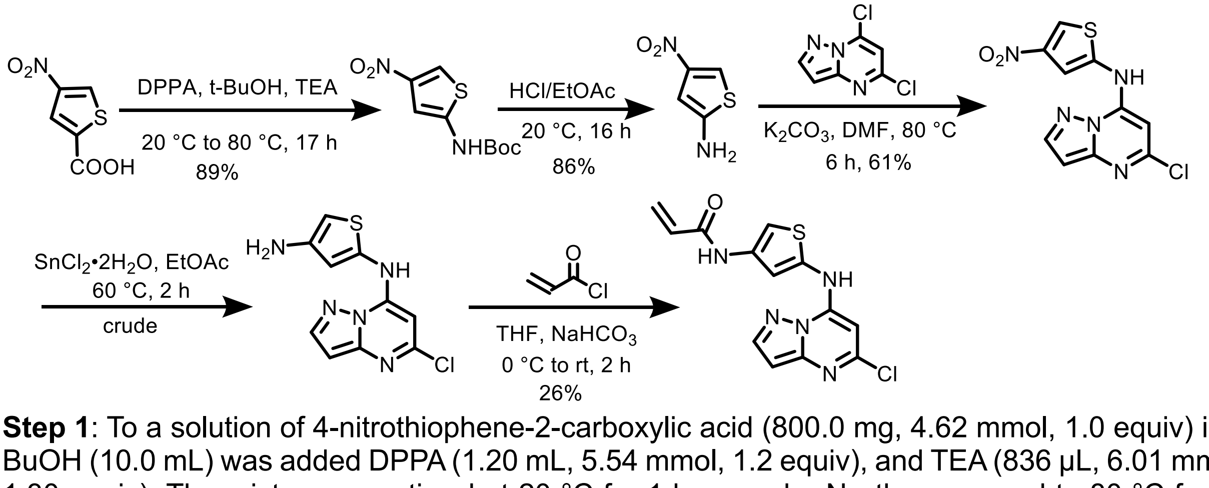

**Step 1**: To a solution of 4-nitrothiophene-2-carboxylic acid (800.0 mg, 4.62 mmol, 1.0 equiv) in t-BuOH (10.0 mL) was added DPPA (1.20 mL, 5.54 mmol, 1.2 equiv), and TEA (836 µL, 6.01 mmol, 1.30 equiv). The mixture was stirred at 20 °C for 1 hour under $N_2$, then warmed to 80 °C for 16 hours. The solvent was removed under vacuum. The residue was purified by column chromatography ($SiO_2$, Petroleum ether/Ethyl acetate = 100/0 to 3/1). Tert-butyl (4-nitrothiophen-2-yl)carbamate (1.00 g, 89%, 4.09 mmol) was obtained as yellow oil. $^1$H NMR: 400 MHz, DMSO-$d_6$ δ 10.92 (br s, 1 H), 8.16 (d, J = 2.0 Hz, 1 H), 6.93 (d, J = 2.0 Hz, 1 H), 1.48 (s, 9 H).

**Step 2:** A mixture of tert-butyl (4-nitrothiophen-2-yl)carbamate (1.00 g, 4.09 mmol, 1.0 equiv) in HCl/EtOAc (10.0 mL, 0.4 M) was stirred at 20 °C for 16 hours. The mixture was concentrated under vacuum, then diluted with $NaHCO_3$ (aq, 15.0 mL) and extracted with EtOAc (10.0 mL * 3). The combined organic layers were washed with brine (10.0 mL), dried over $Na_2SO_4$, filtered, and concentrated under reduced pressure. 4-nitrothiophen-2-amine (505 mg, 86%, 3.50 mmol) was obtained as brown solid. $^1$H NMR: 400 MHz, DMSO-$d_6$ δ 7.64 (d, J = 2.0 Hz, 1 H), 6.30 (d, J = 2.0 Hz, 1 H), 6.18 (s, 2 H).

**Step 3**: To a mixture of 4-nitrothiophen-2-amine (500.0 mg, 3.47 mmol, 1.0 equiv) in DMF (6.00 mL) was added $K_2CO_3$ (958 mg, 6.94 mmol, 2.0 equiv) and 5,7-dichloropyrazolo[1,5-a]pyrimidine (913 mg, 4.86 mmol, 1.4 equiv). The mixture was stirred at 80 °C for 6 hours. The reaction mixture was quenched by the addition of ice-water (30.0 mL), then extracted with EtOAc (10.0 mL * 3). The combined organic layers were washed with brine (15.0 mL), dried over $Na_2SO_4$, filtered, and concentrated under reduced pressure. The residue was purified by column chromatography ($SiO_2$, Petroleum ether/Ethyl acetate = 100/1 to 0/1). 5-chloro-N-(4-nitrothiophen-2-yl)pyrazolo[1,5-a]pyrimidin-7-amine (630 mg, 61%, 2.13 mmol) was obtained as yellow solid. $^1$H NMR: 400 MHz, DMSO-$d_6$ δ 10.89 (br s, 1 H), 8.65 (d, J = 1.6 Hz, 1 H), 8.28 (d, J = 2.4 Hz, 1 H), 7.74 (d, J = 1.6 Hz, 1 H), 6.59 (d, J = 2.4 Hz, 1 H), 6.38 (s, 1 H).

**Step 4:** To a solution of 5-chloro-N-(4-nitrothiophen-2-yl)pyrazolo[1,5-a]pyrimidin-7-amine (200.0 mg, 676 µmol, 1.0 equiv) in EtOAc (5.00 mL) was added $SnCl_2$•$2H_2O$ (610.0 mg, 2.71 mmol, 4.0 equiv). The mixture was stirred at 60 °C for 2 hours. The reaction mixture was quenched by the addition of $NaHCO_3$ (aq, 5.00 mL) at 0 °C, and then extracted with EtOAc (5.00 mL * 3). The combined organic layers were washed with brine (5.00 mL), dried over $Na_2SO_4$, filtered, and concentrated under reduced pressure. N2-(5-chloropyrazolo[1,5-a]pyrimidin-7-yl)thiophene-2,4-diamine (120.0 mg, crude) was obtained as brown solid and used to the next step directly. MS (ESI) m/z = 266.0 $[M+H]^+$.

**Step 5**: To a solution of N2-(5-chloropyrazolo[1,5-a]pyrimidin-7-yl)thiophene-2,4-diamine (120 mg, 353 µmol, 1.0 equiv) in THF (4.00 mL) was added $NaHCO_3$ (54.9 µL, 1.41 mmol, 4.0 equiv) and acryloyl chloride (57.3 µL, 706 µmol, 2.0 equiv) in THF (1.00 mL) at 0 °C. The mixture was stirred at 20 °C for 2 hours. The reaction mixture was quenched by the addition of iced water (10.0 mL) at 0 °C, and then extracted with EtOAc (4.00 mL * 3). The combined organic layers were washed with brine (5.00 mL), dried over $Na_2SO_4$, filtered, and concentrated under reduced pressure. The residue was purified by prep-HPLC (85% to 10% water (0.05% TFA) to MeOH (0.05% TFA). N-(5-((5-chloropyrazolo[1,5-a]pyrimidin-7-yl)amino)thiophen-3-yl)acrylamide (30.0 mg, 26%, 92.0 µmol) was obtained as yellow solid. MS (ESI) m/z = 320.0 $[M+H]^+$. $^1$H NMR (500 MHz, DMSO) δ 10.69 (s, 1H), 10.58 (s, 1H), 8.25 (d, *J* = 2.2 Hz, 1H), 7.49 (d, *J* = 1.7 Hz, 1H), 7.13 (d, *J* = 1.8 Hz, 1H), 6.56 (d, *J* = 2.3 Hz, 1H), 6.37 (dd, *J* = 17.0, 10.0 Hz, 1H), 6.27 (dd, *J* = 17.0, 2.1 Hz, 1H), 6.20 (s, 1H), 5.76 (dd, *J* = 9.9, 2.1 Hz, 1H). $^{13}$C NMR (126 MHz, DMSO) δ 162.56, 150.32, 147.57, 146.69, 144.77, 136.83, 134.73, 131.17, 127.02, 117.28, 106.01, 95.58, 86.15.

**Synthesis of SCA11 (1-(6-((5-chloropyrazolo[1,5-a]pyrimidin-7-yl)amino)indolin-1-yl)prop-2-en-1-one):**

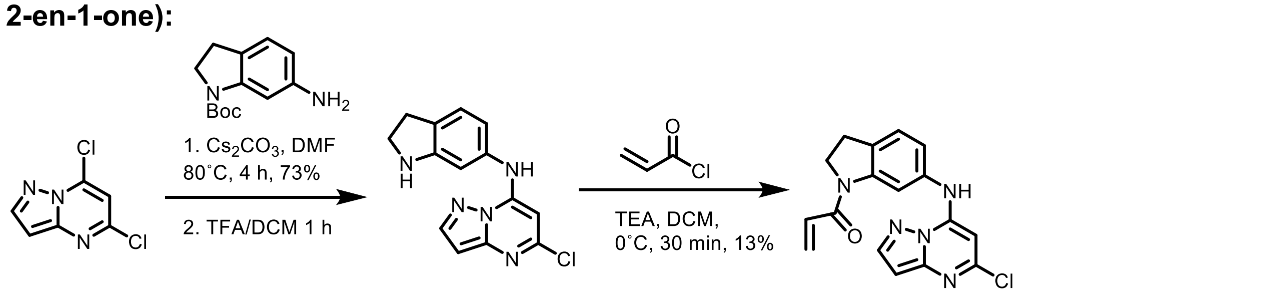

**Step 1:** To a solution of tert-butyl 6-aminoindoline-1-carboxylate (100.0 mg, 0.427 mmol, 1.0 equiv) in DMF (4.27 mL, 0.1 M) was added cesium carbonate (417 mg, 1.28 mmol, 3.0 equiv) and 5,7-dichloropyrazolo[1,5-a]pyrimidine (88.3 mg, 0.469 mmol, 1.1 equiv). The solution was heated to 80˚ C for four hours at which point starting material was consumed. The reaction was diluted in 10 mL of water and product was extracted from the aqueous later with 3 x 10 mL ethyl acetate. The organic layer was washed with brine, dried over $Na_2SO4$, filtered, and concentrated *in vacuo*. The residue was purified via normal phase chromatography (0-60% ethyl acetate: hexane) to afford 5-chloro-N-(indolin-6-yl)pyrazolo[1,5-a]pyrimidin-7-amine (119 mg, 73%, 0.309 mmol). The product was then dissolved in 4.00 mL of DCM and 1.00 mL TFA and allowed to run at 25˚ C for 1 hour, at which point the reaction was concentrated and placed under vacuum overnight. The crude product was carried on to the next step. LC-MS (ESI) m/z: 386.28 $[M+H]^+$.

**Step 2:** To a solution of 5-chloro-N-(indolin-6-yl)pyrazolo[1,5-a]pyrimidin-7-amine (25.0 mg, 0.0870 mmol, 1.0 equiv) in DCM (4.4 mL, 0.02 M) was added DIPEA (0.110 mL, 0.610 mmol, 7.0 equiv). The mixture was cooled to 0˚ C and a 1% solution of acryloyl chloride in DCM (0.350 mL, 0.0443 mmol, 0.5 equiv) was added dropwise. 15 minutes later an additional 0.6 equiv of the 1% acryloyl chloride DCM solution was added dropwise and the reaction was allowed to stir for an additional 15 minutes. The reaction was concentrated *in vacuo* and purified by prep-HPLC (85% to 10% water (0.05% TFA) to MeOH (0.05% TFA)) to afford 5-chloro-N-(indolin-6-yl)pyrazolo[1,5-a]pyrimidin-7-amine (3.75 mg, 13%, 0.0110 mmol) as a white solid. LC-MS (ESI) m/z: 340.11 $[M+H]^+$. $^1H$ NMR (500 MHz, DMSO) δ 10.36 (s, 1H), 8.23 (t, *J* = 3.0 Hz, 2H), 7.34 (d, 1H), 7.12 (dd, *J* = 7.9, 2.1 Hz, 1H), 6.76 (dd, *J* = 16.6, 10.3 Hz, 1H), 6.52 (d, *J* = 2.3 Hz, 1H), 6.31 (dd, *J* = 16.7, 2.1 Hz, 1H), 6.01 (s, 1H), 5.84 (dd, *J* = 10.3, 2.1 Hz, 1H), 4.28 (t, *J* = 8.4 Hz, 2H), 3.20 (t, *J* = 8.5 Hz, 2H). $^{13}C$ NMR (126 MHz, DMSO) δ 163.54, 150.24, 147.69, 146.66, 144.52, 143.74, 135.18, 130.97, 129.77, 128.81, 125.48, 120.68, 113.44, 95.34, 85.28, 48.30, 27.13.

**Synthesis of SCA12 (N-(3-(pyrazolo[1,5-a]pyrimidin-7-ylamino)phenyl)acrylamide):**

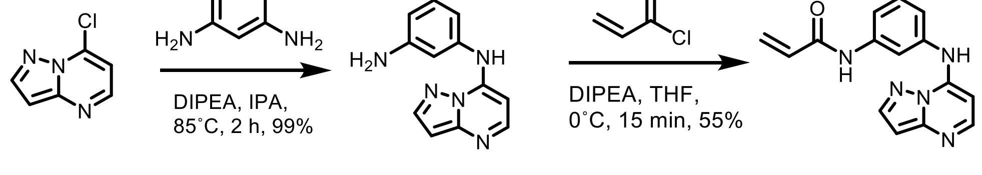

**Step 1:** To a solution of 7-chloropyrazolo[1,5-a]pyrimidine (410.0 mg, 2.67 mmol, 1.0 equiv) and 5-benzene-1,3-diamine (289 mg, 2.67 mmol, 1.0 equiv) in isopropanol (8.09 mL, 0.33 M) was added DIPEA (1.86 mL, 10.7 mmol, 4.0 equiv). The reaction was allowed to stir at 85˚ C for 2 hours. The reaction was concentrated *in vacuo* and purified through normal phase chromatography (0-100% ethyl acetate: hexane) to afford N1-(pyrazolo[1,5-a]pyrimidin-7-yl)benzene-1,3-diamine as a beige solid (595 mg, 99%, 2.64 mmol). LC-MS (ESI) m/z: 226.10 $[M+H]^+$.

**Step 2:** To a crude solution of N1-(pyrazolo[1,5-a]pyrimidin-7-yl)benzene-1,3-diamine (50.0 mg, 0.220 mmol, 1.0 equiv) in THF (0.02 mL, 0.05 M) was added DIPEA (0.12 mL, 0.67 mmol, 3.0 equiv). The mixture was cooled to 0˚ C and acryloyl chloride (0.0180 mL, 0.220 mmol, 1.0 equiv) was added dropwise and the reaction was allowed to stir for 15 minutes. The reaction was concentrated *in vacuo* and purified by prep-HPLC (85% to 10% water (0.05% TFA) to MeOH (0.05% TFA)) to afford N-(3-(pyrazolo[1,5-a]pyrimidin-7-ylamino)phenyl)acrylamide (34.4 mg, 55%, 0.123 mmol) as a white solid. LC-MS (ESI) m/z: 280.13 $[M+H]^+$. $^1$H NMR (500 MHz, DMSO) δ 10.44 (s, 1H), 8.38 (d, *J* = 2.3 Hz, 1H), 8.35 (d, *J* = 6.4 Hz, 1H), 7.94 (t, *J* = 2.1 Hz, 1H), 7.61 (ddd, *J* = 8.2, 2.0, 1.0 Hz, 1H), 7.48 (t, *J* = 8.0 Hz, 1H), 7.20 (ddd, *J* = 7.9, 2.1, 1.0 Hz, 1H), 6.67 (d, *J* = 2.3 Hz, 1H), 6.49 (d, *J* = 10.2 Hz, 1H), 6.45 (d, *J* = 10.2 Hz, 1H), 6.28 (dd, *J* = 17.0, 1.9 Hz, 1H), 5.79 (dd, *J* = 10.2, 1.9 Hz, 1H). $^{13}$C NMR (126 MHz, DMSO) δ 163.49, 148.07, 145.50, 145.31, 143.27, 140.18, 136.18, 131.67, 130.07, 127.45, 120.28, 118.10, 115.90, 93.14, 87.55.

**Synthesis of SCA13 (N-(3-(5-aminopyrazolo[1,5-a]pyrimidin-7-ylamino)phenyl)acrylamide):**

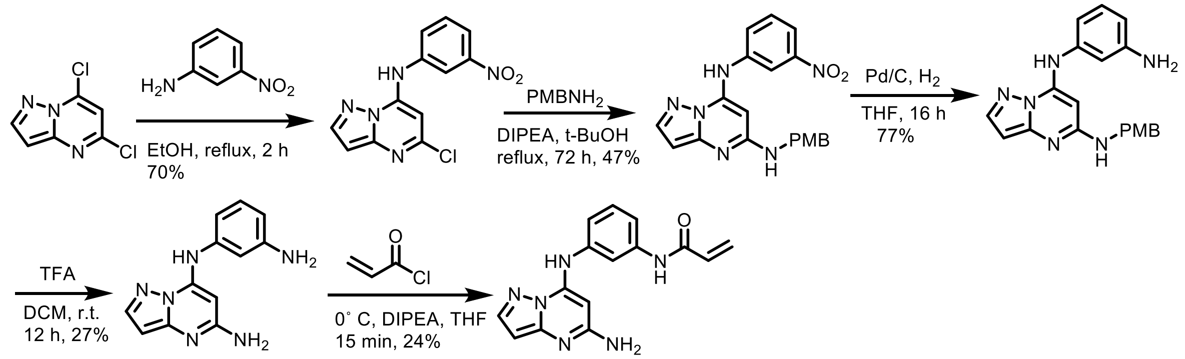

**Step 1:** A mixture of 5,7-dichloropyrazolo[1,5-a]pyrimidine (470.0 mg, 2.50 mmol, 1.0 equiv) and 3-nitroaniline (345 mg, 2.50 mmol, 1.0 equiv) in EtOH (10.0 mL, 0.25 M) was stirred at reflux for 2 h. After cooling to room temperature, the precipitation was collected by filtration, washed with EtOH (2.0 mL x 2) and dried under vacuum to afford 5-chloro-N-(3-nitrophenyl)pyrazolo[1,5-a]pyrimidin-7-amine as a yellow solid (500.0 mg, 70%, 1.72 mmol). LC-MS (ESI) m/z: 290 $[M+H]^+$.

**Step 2:** To a solution of 5-chloro-N-(3-nitrophenyl)pyrazolo[1,5-a]pyrimidin-7-amine (500.0 mg, 1.73 mmol, 1.0 equiv) and (4-methoxyphenyl)methanamine (356 mg, 2.60 mmol, 1.5 equiv) in t-BuOH (30.0 mL, 0.0577 M) was added DIPEA (670.0 mg, 5.19 mmol, 3.0 equiv). The mixture was stirred at reflux for 48 h. After removal of the solvent, the residue was purified with prep-HPLC (85% to 10% water (0.05% TFA) to MeOH (0.05% TFA)) to afford N5-(4-methoxybenzyl)-N7-(3-nitrophenyl)pyrazolo[1,5-a]pyrimidine-5,7-diamine, as a colorless oil (320.0 mg, 47%, 0.818 mmol). LC-MS (ESI) m/z: 391$[M+H]^+$.

**Step 3:** To a solution of N5-(4-methoxybenzyl)-N7-(3-nitrophenyl)pyrazolo[1,5-a]pyrimidine-5,7-diamine (320.0 mg, 0.820 mmol, 1.0 equiv) in MeOH (20.0 mL, 0.041 M) was added 10 % Pd/C (200.0 mg, 0.0532, 0.06 equiv). The mixture was stirred at room temperature under hydrogen (1 atm) overnight. The mixture was filtered and concentrated to afford crude N7-(3-aminophenyl)-N5-(4-methoxybenzyl)pyrazolo[1,5-a]pyrimidine-5,7-diamine as a yellow solid (226 mg, 77%, 0.626 mmol). LC-MS (ESI) m/z: 361$[M+H]^+$.

**Step 4:** To a solution of N7-(3-aminophenyl)-N5-(4-methoxybenzyl)pyrazolo[1,5-a]pyrimidine-5,7-diamine (226 mg, 0.630 mmol, 1.0 equiv) in THF (10.0 mL, 0.063 M) was added TFA (5.00 mL, 0.126 M). The mixture was stirred at room temperature overnight. After removal of the solvent, the residue was purified by prep-HPLC (85% to 10% water (0.05% TFA) to MeOH (0.05% TFA)) to afford N7-(3-aminophenyl)pyrazolo[1,5-a]pyrimidine-5,7-diamine as a brown solid (40.0 mg, 27%, 0.166 mmol). LC-MS (ESI) m/z: 241 $[M+H]^+$.

**Step 5:** To a solution of N7-(3-aminophenyl)pyrazolo[1,5-a]pyrimidine-5,7-diamine (40.0 mg, 0.170 mmol, 1.0 equiv) in THF (5.00 ml, 0.034 M) was added DIPEA (0.0550 mL, 0.340 mmol, 2.0 equiv). A solution of acryloyl chloride (0.0134 mL, 0.170 mmol, 1.0 equiv) in DCM (1.00 ml, 0.17 M) was added dropwise at 0˚ C. The mixture was stirred at 0˚ C for 15 min. After removal of the solvent, the residue was purified with prep-HPLC (85% to 10% water to MeOH) to afford N-(3-(5-aminopyrazolo[1,5-a]pyrimidin-7-ylamino)phenyl)acrylamide as a white solid (12 mg, 24%, 0.407). LC-MS (ESI) m/z: 295.14 $[M+H]^+$. $^1H$ NMR (500 MHz, DMSO) δ 10.26 (s, 1H), 9.23 (s, 1H), 7.81 (d, *J* = 2.1 Hz, 1H), 7.72 (t, *J* = 2.1 Hz, 1H), 7.56 (ddd, *J* = 8.2, 2.1, 1.0 Hz, 1H), 7.37 (t, *J* = 8.1 Hz, 1H), 7.10 (ddd, *J* = 7.8, 2.1, 0.9 Hz, 1H), 6.50 – 6.43 (m, 1H), 6.28 (dd, *J* = 17.0, 2.0 Hz, 1H), 6.22 (s, 2H), 5.86 (d, *J* = 2.1 Hz, 1H), 5.77 (dd, *J* = 10.1, 2.0 Hz, 1H), 5.62 (s, 1H). $^{13}C$ NMR (126 MHz, DMSO) δ 163.23, 158.72, 149.49, 144.89, 142.55, 139.87, 138.52, 131.77, 129.59, 127.17, 119.17, 115.88, 114.72, 90.49, 74.53.

**Synthesis of SCA14 (N-(3-(5-(dimethylamino)pyrazolo[1,5-a]pyrimidin-7-ylamino)phenyl)acrylamide):**

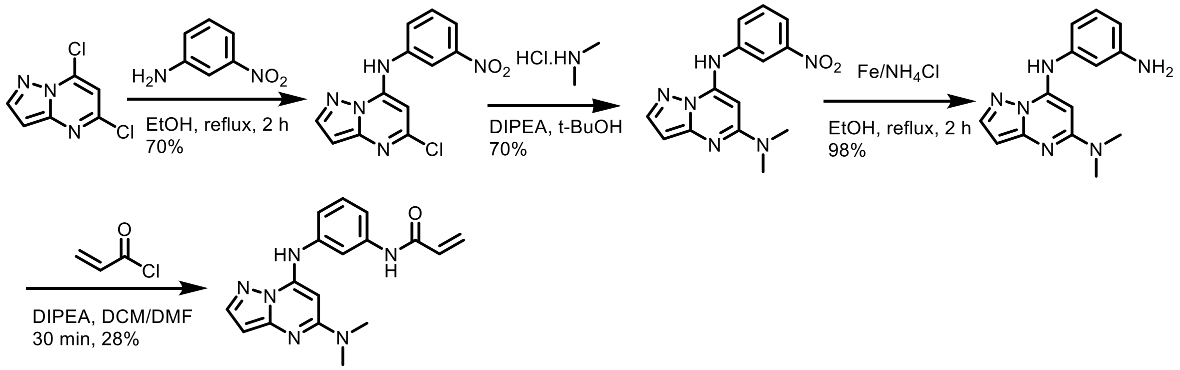

**Step 1:** A mixture of 5,7-dichloropyrazolo[1,5-a]pyrimidine (470.0 mg, 2.50 mmol, 1.0 equiv) and 3-nitroaniline (345 mg, 2.50 mmol, 1.0 equiv) in EtOH (10.0 mL, 0.25 M) was stirred at reflux for 2 h. After cooling to room temperature, the precipitate was collected by filtration, washed with EtOH (2.0 mL x 2) and dried under vacuum to afford 5-chloro-N-(3-nitrophenyl)pyrazolo[1,5-a]pyrimidin-7-amine as a yellow solid (500.0 mg, 70%, 1.72 mmol). LC-MS (ESI) m/z: 290 $[M+H]^+$.

**Step 2:** To a solution of 5,7-dichloropyrazolo[1,5-a]pyrimidine (300.0 mg, 1.00 mmol, 1.0 equiv) and dimethylamine hydrochloride (120.0 mg, 1.50 mmol, 1.5 equiv) in t-BuOH (30.0 mL, 0.0333 M) was added DIPEA (0.488 mL, 3.00 mmol, 3.0 equiv). The mixture was stirred at reflux for 48 h. After removal of the solvent, the residue was purified with prep-HPLC to afford N5,N5-dimethyl-N7-(3-nitrophenyl)pyrazolo[1,5-a]pyrimidine-5,7-diamine as a yellow solid (216 mg, 70%, 0.722 mmol). LC-MS (ESI) m/z: 299 $[M+H]^+$.

**Step 3:** To a solution of N5,N5-dimethyl-N7-(3-nitrophenyl)pyrazolo[1,5-a]pyrimidine-5,7-diamine (216 mg, 0.720 mmol, 1.0 equiv) in EtOH (20.0 mL, 0.036 M) and water (1.00 mL, 0.7 M) was added Fe powder (403 mg, 7.20 mmol, 10.0 equiv) and $NH_4Cl$ (193 mg, 3.60 mmol, 5.0 equiv). The mixture was stirred at reflux for 2 h. After cooling to room temperature, the mixture was filtered through Celite and the cake was washed with EtOH (10 mL x 2). The combined filtrate and wash was evaporated to afford N7-(3-aminophenyl)-N5,N5-dimethylpyrazolo[1,5-a]pyrimidine-5,7-diamine as a brown solid (190.0 mg, 98%, 3.00 mmol). LC-MS (ESI) m/z: 269 $[M+H]^+$.

**Step 4:** To a solution of N7-(3-aminophenyl)-N5,N5-dimethylpyrazolo[1,5-a]pyrimidine-5,7-diamine (190.0 mg, 0.710 mmol, 1.0 equiv) in DMF (5.00 ml, 0.742 M) was added DIPEA (0.344 mL, 2.13 mmol, 3 equiv). A solution of acryloyl chloride (0.0572 mL, 0.710 mmol, 1 equiv) in DCM (2.00 ml, 0.356 M) was added dropwise at 0° C. The mixture was stirred at 0° C for 30 min. After removal of the solvent, the residue was purified by prep-HPLC (85% to 10% water (0.05% TFA) to MeOH (0.05% TFA)) to afford N1-(5-chloropyrazolo[1,5-a]pyrimidin-7-yl)benzene-1,4-diamine as a white solid (63.0 mg, 28%, 0.195 mmol). LC-MS (ESI) m/z: 323.15 $[M+H]^+$. $^1$H-NMR (DMSO-$d_6$, 400 MHz): δ (ppm) 3.04 (s, 6H), 5.78 (d, J = 10.0 Hz, 1H), 5.92 (s, 1H), 5.99 (s, 1H), 6.25-6.30 (m, 1H), 6.42-6.49 (m, 1H), 7.17-7.19 (m, 1H), 7.35-7.39 (m, 2H), 7.88 (s, 1H), 8.00 (s, 1H), 9.39 (s, 1H), 10.28 (s, 1H). $^1$H NMR (500 MHz, DMSO) δ 10.26 (s, 1H), 9.44 (s, 1H), 8.00 (t, *J* = 2.0 Hz, 1H), 7.88 (d, *J* = 2.1 Hz, 1H), 7.40 – 7.31 (m, 2H), 7.17 (dt, *J* = 7.3, 2.0 Hz, 1H), 6.45 (dd, *J* = 17.0, 10.1 Hz, 1H), 6.27 (dd, *J* = 17.0, 2.0 Hz, 1H), 6.00 (d, *J* = 2.1 Hz, 1H), 5.90 (s, 1H), 5.77 (dd, *J* = 10.1, 2.0 Hz, 1H), 3.04 (s, 6H). $^{13}$C NMR (126 MHz, DMSO) δ 163.34, 157.35, 144.34, 143.05, 139.62, 138.40, 131.77, 129.67, 127.14, 118.31, 115.45, 113.67, 91.03, 72.48, 37.69.

**Synthesis of SCA15 (N-(3-(5-methoxypyrazolo[1,5-a]pyrimidin-7-ylamino)phenyl)acrylamide):**

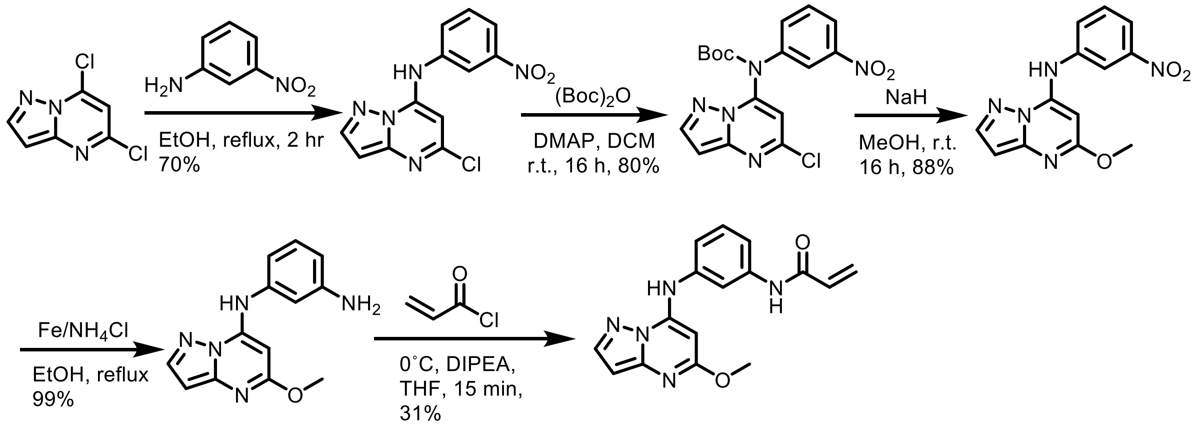

**Step 1:** A mixture of 5,7-dichloropyrazolo[1,5-a]pyrimidine (470.0 mg, 2.50 mmol, 1.0 equiv) and 3-nitroaniline (345 mg, 2.50 mmol, 1.0 equiv) in EtOH (10.0 mL, 0.25 M) was stirred at reflux for 2 hours. After cooling to room temperature, the precipitate was collected by filtration, washed with EtOH (2.0 mL x 2) and dried under vacuum to afford 5-chloro-N-(3-nitrophenyl)pyrazolo[1,5-a]pyrimidin-7-amine as a yellow solid (500.0 mg, 70%, 1.72 mmol). LC-MS (ESI) m/z: 290 $[M+H]^+$.

**Step 2:** To a solution of 5-chloro-N-(3-nitrophenyl)pyrazolo[1,5-a]pyrimidin-7-amine (289 mg, 1.0 mmol, 1.0 equiv) in DCM (10.0 mL, 0.1 M) was added DMAP (146 mg, 1.20 mmol, 1.2 equiv) and di-tert-butyl bicarbonate (218 mg, 1.00 mmol, 1.0 equiv). The mixture was stirred at room temperature overnight. The mixture was diluted with DCM (50 mL), washed with brine (50 mL), dried over anhydrous $Na_2SO_4$, concentrated, and purified with column chromatography on silica gel (ethyl acetate in petroleum ether, 10 %, v/v) to afford tert-butyl 5-chloropyrazolo[1,5-a]pyrimidin-7-yl(3-nitrophenyl)carbamate as a white solid (312 mg, 80%, 0.934 mmol). LC-MS (ESI) m/z: 334$[M-55]^+$.

**Step 3:** To a solution of 60% NaH (320.0 mg, 8.00 mmol, 8.0 equiv) in MeOH (15.0 mL, 0.0533 M) was added tert-butyl 5-chloropyrazolo[1,5-a]pyrimidin-7-yl(3-nitrophenyl)carbamate (310.0 mg, 0.800 mmol, 1.0 equiv). The mixture was stirred at room temperature overnight. The mixture was quenched with water (50 mL) and extracted with ethyl acetate (50 mL x 3), the combined organic layer was washed with brine (100 mL), dried over anhydrous $Na_2SO_4$, filtered and concentrated to afford 5-methoxy-N-(3-nitrophenyl)pyrazolo[1,5-a]pyrimidin-7-amine as a white solid (200.0 mg, 88%, 0.699 mmol). LC-MS (ESI) m/z: 286$[M+H]^+$.

**Step 4:** To a solution of 5-methoxy-N-(3-nitrophenyl)pyrazolo[1,5-a]pyrimidin-7-amine (200.0 mg, 0.700 mmol, 1.0 equiv) in EtOH (20.0 mL, 0.035 M) and water (1.00 mL, 0.7 M) was added Fe powder (392 mg, 7.00 mmol, 10.0 equiv) and $NH_4Cl$ (187 mg, 3.50 mmol, 5.0 equiv). The mixture was stirred at reflux for 2 h. After cooling to room temperature, the mixture was filtered through Celite and washed with EtOH (10 mL x 2). The filtrate was evaporated to afford N1-(5-methoxypyrazolo[1,5-a]pyrimidin-7-yl)benzene-1,3-diamine, as a brown solid (178 mg, 99%, 0.695 mmol). LC-MS (ESI) m/z: 256 $[M+H]^+$.

**Step 5:** To a solution of N1-(5-methoxypyrazolo[1,5-a]pyrimidin-7-yl)benzene-1,3-diamine (178 mg, 0.700 mmol, 1.0 equiv) in THF (5.00 ml, 0.14 M) was added DIPEA (271 mg, 2.10 mmol, 3 equiv). A solution of acryloyl chloride (0.0563 mL, 0.70 mmol, 1.0 equiv) in DCM (1.0 ml, 0.7 M) was added dropwise at 0˚ C. The mixture was stirred at 0˚ C for 15 min. After removal of the solvent, the residue was purified with prep-HPLC (85% to 10% water to MeOH) to afford N-(3-(5-methoxypyrazolo[1,5-a]pyrimidin-7-ylamino)phenyl)acrylamide as a white solid (67.0 mg, 31%, 2.78 mmol). LC-MS (ESI) m/z: 310.15 $[M+H]^+$. $^1$H NMR (500 MHz, DMSO) δ 10.27 (s, 1H), 9.77 (s, 1H), 8.05 (d, *J* = 2.1 Hz, 1H), 7.84 (t, *J* = 2.1 Hz, 1H), 7.50 (ddd, *J* = 8.1, 2.0, 0.9 Hz, 1H), 7.38 (t, *J* = 8.1 Hz, 1H), 7.15 (ddd, *J* = 8.0, 2.2, 1.0 Hz, 1H), 6.44 (dd, *J* = 17.0, 10.1 Hz, 1H), 6.32 (d, *J* = 2.1 Hz, 1H), 6.28 (dd, *J* = 17.0, 2.0 Hz, 1H), 5.78 (dd, *J* = 10.1, 2.0 Hz, 1H), 5.76 (s, 1H), 3.85 (s, 3H). $^{13}$C NMR (126 MHz, DMSO) δ 163.32, 163.19, 147.54, 146.10, 143.34, 139.78, 137.88, 131.75, 129.71, 127.20, 119.02, 116.19, 114.22, 93.87, 74.82, 53.28.

**Synthesis of SCA16 (N-(3-(5-chloropyrazolo[1,5-a]pyrimidin-7-yloxy)phenyl)acrylamide:**

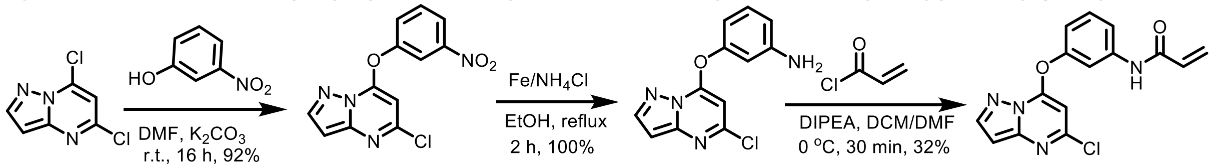

**Step 1:** To a solution of 5,7-dichloropyrazolo[1,5-a]pyrimidine (470.0 mg, 2.50 mmol, 1.0 equiv) and 3-nitrophenol (348 mg, 2.50 mmol, 1.0 equiv) in DMF (10.0 mL, 0.25 M) was added $K_2CO_3$ (690.0 mg, 5.00 mmol, 2.0 equiv). The mixture was stirred at room temperature overnight. The mixture was poured into water (100 mL) and stirred for 30 min. The precipitate was collected by filtration, washed with water (2.0 mL x 3) and dried under vacuum to afford 5-chloro-7-(3-nitrophenoxy)pyrazolo[1,5-a]pyrimidine as a yellow solid (672 mg, 92%, 2.31 mmol). LC-MS (ESI) m/z: 291 $[M+H]^+$.

**Step 2:** To a solution of 5-chloro-7-(3-nitrophenoxy)pyrazolo[1,5-a]pyrimidine (200.0 mg, 0.69 mmol, 1.0 equiv) in EtOH (8.00 mL, 0.0893 M) and water (1.00 mL, 0.690 M) was added Fe powder (386 mg, 6.90 mmol, 10.0 equiv) and $NH_4Cl$ (185 mg, 3.45 mmol, 5.0 equiv). The mixture was stirred at reflux for 2 hours. After cooling to room temperature, the mixture was filtered through Celite and the cake was washed with EtOH (10 mL x 2) to afford 3-(5-chloropyrazolo[1,5-a]pyrimidin-7-yloxy)aniline as a yellow solid (179 mg, 100%, 0.686 mmol). LC-MS (ESI) m/z: 261 $[M+H]^+$.

**Step 3:** To a solution of 3-(5-chloropyrazolo[1,5-a]pyrimidin-7-yloxy)aniline (179 mg, 0.690 mmol, 1.0 equiv) in DMF (4.00 ml, 0.173 M) was added DIPEA (356 mg, 2.76 mmol, 4.0 equiv). A solution of acryloyl chloride (186 mg, 2.07 mmol, 3.0 equiv) in DCM (13.8 ml, 0.05 M) was added dropwise at 0˚ C. The mixture was stirred at 0˚ C for 30 min. After removal of the solvent, the residue was purified by prep-HPLC (85% to 10% water to MeOH) to afford N-(3-(5-chloropyrazolo[1,5-

a]pyrimidin-7-yloxy)phenyl)acrylamide as a white solid (70.0 mg, 32%, 0.223 mmol). LC-MS (ESI) m/z: 315.05 $[M+H]^+$. $^1H$ NMR (500 MHz, DMSO) δ 10.46 (s, 1H), 8.33 (d, *J* = 2.3 Hz, 1H), 7.87 (t, *J* = 2.2 Hz, 1H), 7.61 (ddd, *J* = 8.3, 2.0, 1.0 Hz, 1H), 7.53 (t, *J* = 8.1 Hz, 1H), 7.19 (ddd, *J* = 8.1, 2.5, 1.0 Hz, 1H), 6.75 (d, *J* = 2.3 Hz, 1H), 6.44 (dd, *J* = 17.0, 10.1 Hz, 1H), 6.28 (dd, *J* = 17.0, 1.9 Hz, 1H), 6.13 (s, 1H), 5.80 (dd, *J* = 10.1, 1.9 Hz, 1H). $^{13}C$ NMR (126 MHz, DMSO) δ 163.49, 155.21, 151.53, 150.35, 148.56, 146.07, 141.02, 131.48, 130.97, 127.64, 117.93, 115.43, 111.32, 96.53, 90.35.

**Synthesis of SCA17: ((E)-N-(3-((5-chloropyrazolo[1,5-a]pyrimidin-7-yl)amino)phenyl)-4-(dimethylamino)but-2-enamide):**

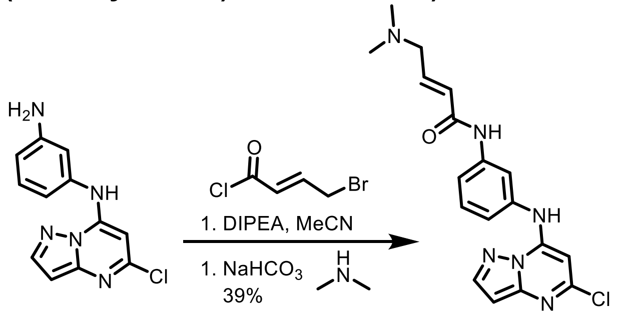

To a solution of N1-(5-chloropyrazolo[1,5-a]pyrimidin-7-yl)benzene-1,3-diamine (16.0 mg, 0.0623 mmol, 1.0 equiv) in acetonitrile at 0° C (0.620 mL, 0.1 M) was added DIPEA (0.032 mL, 0.180 mmol, 3.0 equiv) and (E)-4-bromobut-2-enoyl chloride (12.0 mg, 0.068 mmol, 1.1 equiv). After 1 hour, the reaction was extracted with ethyl acetate (3 x 10 mL) and the combined organic layer was concentrated *in vacuo.* To the crude product dissolved in acetonitrile (0.620 mL, 0.1 M) was added sodium bicarbonate (19.1 mg, 0.180 mmol, 3.0 equiv) and methylamine (0.0138 mL, 0.180 mmol, 3.0 equiv). The reaction was heated to 50° C and allowed to stir for 2 hours, at which point the reaction was purified by prep-HPLC (85% to 10% water (0.05% TFA) to MeOH (0.05% TFA)) to afford (E)-N-(3-((5-chloropyrazolo[1,5-a]pyrimidin-7-yl)amino)phenyl)-4-(dimethylamino)but-2-enamide (9.00 mg, 39%, 0.0243 mmol) as a white solid. LC-MS (ESI) m/z: 371.12 $[M+H]^+$. $^1H$ NMR (500 MHz, DMSO) δ 10.57 (s, 1H), 10.42 (s, 1H), 10.25 (s, 1H), 8.25 (d, *J* = 2.2 Hz, 1H), 7.85 (t, *J* = 2.1 Hz, 1H), 7.58 (dd, *J* = 8.1, 2.0 Hz, 1H), 7.45 (t, *J* = 8.0 Hz, 1H), 7.20 (dd, *J* = 8.0, 2.1 Hz, 1H), 6.77 (dt, *J* = 14.8, 7.1 Hz, 1H), 6.54 (d, *J* = 2.2 Hz, 1H), 6.50 (d, *J* = 15.3 Hz, 1H), 6.17 (s, 1H), 3.96 (d, *J* = 7.1 Hz, 2H), 2.81 (s, 6H). $^{13}C$ NMR (126 MHz, DMSO) δ 162.22, 158.28, 158.03, 150.33, 147.69, 146.18, 144.55, 139.78, 136.92, 132.04, 131.87, 129.98, 120.14, 117.35, 115.48, 95.47, 85.53, 56.75, 42.02.

**Synthesis of SCA18 (N-(2-amino-6-((5-chloropyrazolo[1,5-a]pyrimidin-7-yl)amino)pyridin-4-yl)acrylamide):**

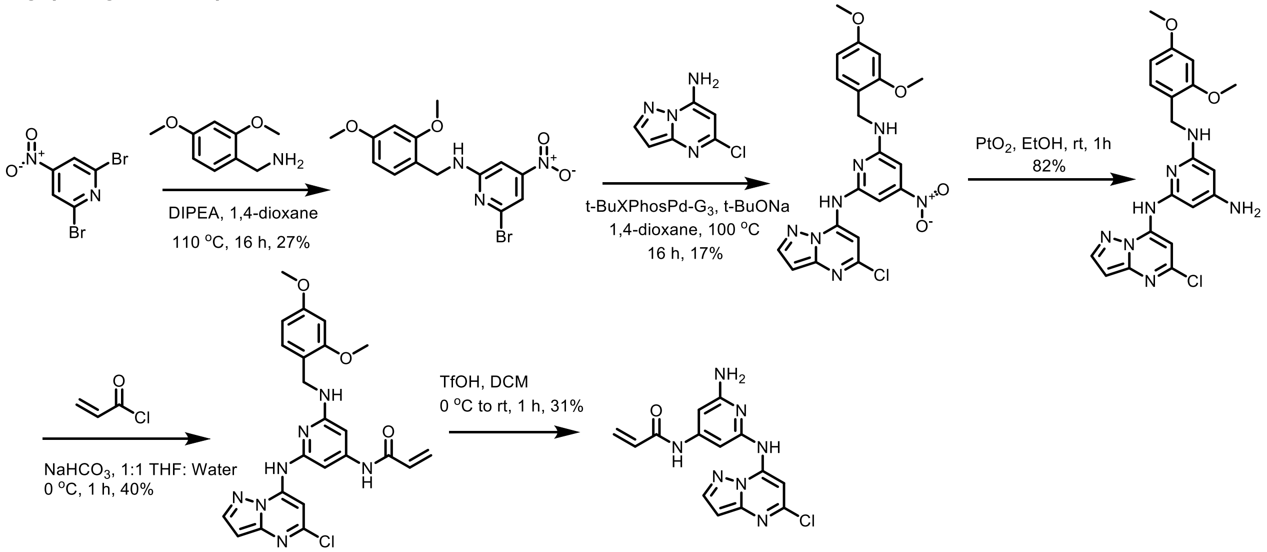

**Step 1:** To a solution of 2,6-dibromo-4-nitropyridine (5.00g, 17.7 mmol, 1.0 equiv) in 1,4-dioxane (177 mL, 0.1 M) was added (2,4-dimethoxyphenyl)methanamine (3.26g, 19.5 mmol, 2.0 equiv) and DIPEA (6.18 mL, 35.5 mmol, 4.59g), and the mixture was stirred at 110 °C for 16 hours under $N_2$. The reaction mixture was filtered, and the filtrate was diluted with 100 mL $H_2O$, extracted with EtOAc (200 mL * 3), dried over $Na_2SO_4$, filtered, and concentrated under reduced pressure. The resulting residue was purified by column chromatography ($SiO_2$,Petroleum ether/Ethyl acetate=1/0 to 5/1). 6-bromo-N-(2,4-dimethoxybenzyl)-4-nitropyridin-2-amine (1.75 g, 27%, 4.75 mmol) was obtained as a yellow solid.

**Step 2:** A solution of 6-bromo-N-(2,4-dimethoxybenzyl)-4-nitropyridin-2-amine (1.75g, 4.75 mmol, 1.0 equiv), 5-chloropyrazolo[1,5-a]pyrimidin-7-amine (801 mg, 4.75 mmol, 1.0 equiv), tBuXPhos Pd G3 (378 mg, 0.475 mmol, 0.1 equiv), and sodium tert-butoxide (914 mg, 9.51 mmol, 2.0 equiv) dissolved in 1,4-dioxane (4.75 mL, 0.1 M) was allowed to stir at 100 °C for 16 hours. The reaction mixture was then filtered, and the filtrate was diluted with 100 mL $H_2O$, extracted with EtOAc (200 mL * 3), dried over $Na_2SO_4$, filtered, and concentrated under reduced pressure. The resulting residue was purified by column chromatography ($SiO_2$,Petroleum ether/Ethyl acetate=1/0 to 5/1) to afford N2-(5-chloropyrazolo[1,5-a]pyrimidin-7-yl)-N6-(2,4-dimethoxybenzyl)-4-nitropyridine-2,6-diamine (360.0 mg, 0.790 mmol, 17%) as a white solid.

**Step 3:** A solution of N2-(5-chloropyrazolo[1,5-a]pyrimidin-7-yl)-N6-(2,4-dimethoxybenzyl)-4-nitropyridine-2,6-diamine (300.0 mg, 0.658 mmol, 1.0 equiv), EtOH (6.58 mL, 0.1 M), and platinum oxide (150.0 mg, 0.658 mmol, 1.0 equiv) was allowed to stir at 1 h, at which point the reaction was filtered. The filtrate was concentrated under reduced pressure to afford N2-(5-chloropyrazolo[1,5-a]pyrimidin-7-yl)-N6-(2,4-dimethoxybenzyl)pyridine-2,4,6-triamine as a crude solid (230.0 mg, 0.540 mmol, 82%).

**Step 4:** To a solution of N2-(5-chloropyrazolo[1,5-a]pyrimidin-7-yl)-N6-(2,4-dimethoxybenzyl)pyridine-2,4,6-triamine (160.0 mg, 0.375 mmol, 1.0 equiv) and sodium bicarbonate (94.7 mg, 1.14 mmol, 3.0 equiv) dissolved in a 1:1 mixture of THF: water (18.8 mL, 0.02 M) at 0 ˚C was added acryloyl chloride (0.191 mL, 0.375 mmol, 1.0 equiv) as a 10% THF solution, dropwise. The mixture was stirred at 15 °C for 1 hour. The reaction mixture was filtered and concentrated under reduced pressure and the resulting residue was purified by prep-HPLC

(85% to 10% water (0.05% TFA) to MeOH (0.05% TFA)) to afford N-(2-((5-chloropyrazolo[1,5-a]pyrimidin-7-yl)amino)-6-((2,4-dimethoxybenzyl)amino)pyridin-4-yl)acrylamide (70.0 mg, 0.15 mmol, 40%) as a white solid.

**Step 5:** To a solution of N-(2-((5-chloropyrazolo[1,5-a]pyrimidin-7-yl)amino)-6-((2,4-dimethoxybenzyl)amino)pyridin-4-yl)acrylamide (140.0 mg, 0.292 mmol, 1.0 equiv) in DCM (2.92 mL, 0.1 M) at 0 ˚C was added triflic acid (0.013 mL, 0.146 mmol, 0.5 equiv) dropwise. The reaction was allowed to warm to room temperature, at which point the reaction was quenched by the dropwise addition of saturated aqueous sodium bicarbonate solution on ice. The reaction was concentrated, then purified by prep-HPLC (85% to 10% water (0.05% TFA) to MeOH (0.05% TFA) to afford N-(2-amino-6-((5-chloropyrazolo[1,5-a]pyrimidin-7-yl)amino)pyridin-4-yl)acrylamide (30.1 mg, 31%, 0.0913 mmol) as a white solid. LC-MS (ESI) m/z = 330.10 $[M+H]^+$. $^1$H NMR (500 MHz, DMSO) δ 10.36 (s, 1H), 8.24 (d, *J* = 2.3 Hz, 1H), 7.71 (s, 1H), 6.90 (d, *J* = 1.6 Hz, 1H), 6.85 (d, *J* = 1.6 Hz, 1H), 6.57 (d, *J* = 2.2 Hz, 1H), 6.47 (dd, *J* = 17.0, 10.2 Hz, 1H), 6.30 (dd, *J* = 17.0, 1.9 Hz, 1H), 5.82 (dd, *J* = 10.1, 1.9 Hz, 1H). $^{13}$C NMR (126 MHz, DMSO) δ 163.97, 158.53, 158.18, 150.54, 148.79, 147.29, 144.16, 143.42, 131.44, 128.23, 95.71, 94.02, 92.76, 90.62.

**Synthesis of SCA19 (N-(2-((5-chloropyrazolo[1,5-a]pyrimidin-7-yl)amino)-6-(cyclopropylamino)pyridin-4-yl)acrylamide):**

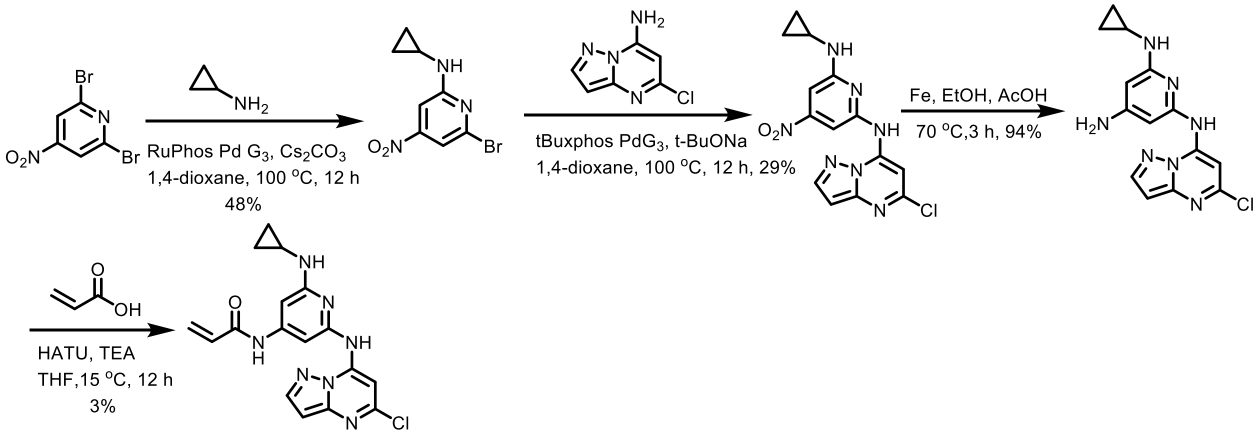

**Step 1:** A mixture of 2,6-dibromo-4-nitro-pyridine (5.00 g, 17.7 mmol, 1.1 equiv), cyclopropanamine (1.12 mL, 16.1 mmol, 1.0 equiv), RuPhos Pd $G_3$ (674 mg, 806 µmol, 0.05 equiv), $Cs_2CO_3$ (10.5 g, 32.3 mmol, 2.0 equiv) in 1,4-dioxane (20.0 mL, 0.885 M) was degassed and purged with $N_2$ for 3 times, and then the mixture was stirred at 100 °C for 12 hours under a $N_2$ atmosphere. The reaction mixture was diluted with $H_2O$ (20 mL) and extracted with EtOAc (30 mL * 2). The combined organic layers were dried over $Na_2SO_4$, filtered, and concentrated under reduced pressure. The resulting residue was purified by column chromatography ($SiO_2$, Petroleum ether/Ethyl acetate=100/1 to 20/1). 6-bromo-N-cyclopropyl-4-nitro-pyridin-2-amine (2.00 g, 48%, 7.75 mmol) was obtained as a yellow solid. LC-MS (ESI) m/z = 258.1 $[M+H]^+$.

**Step 2:** A mixture of 6-bromo-N-cyclopropyl-4-nitro-pyridin-2-amine (2.05 g, 7.94 mmol, 1.0 equiv), 5-chloropyrazolo[1,5-a]pyrimidin-7-amine (1.34 g, 7.94 mmol, 1.0 equiv), t-BuXPhos Pd $G_3$ (631 mg, 794 µmol, 0.1 equiv), NaOtBu (1.53 g, 15.9 mmol, 2.0 equiv) in 1,4-dioxane (20.0 mL) was degassed and purged with $N_2$ for 3 times, and then the mixture was stirred at 100 °C for 12 hours under $N_2$ atmosphere. The reaction mixture was diluted with $H_2O$ (20 mL) and extracted with EtOAc (30 mL * 2). The combined organic layers were dried over $Na_2SO_4$, filtered, and concentrated under reduced pressure and the resulting residue was purified by column chromatography ($SiO_2$, Petroleum ether/Ethyl acetate=100/1 to 0/1). N2-(5-chloropyrazolo[1,5-

a]pyrimidin-7-yl)-N6-cyclopropyl-4-nitro-pyridine-2,6-diamine (800.0 mg, 29%, 2.31 mmol) was obtained as a yellow solid. LC-MS (ESI) m/z: 346.3 $[M+H]^+$.

**Step 3:** A mixture of N2-(5-chloropyrazolo[1,5-a]pyrimidin-7-yl)-N6-cyclopropyl-4-nitro-pyridine-2,6-diamine (700.0 mg, 2.02 mmol, 1.0 equiv), Fe (565 mg, 10.1 mmol, 5.0 equiv) in AcOH (8.00 mL. 0.25 M) and EtOH (8.00 mL, 0.25 M) was degassed and purged with $N_2$ for 3 times, and then the mixture was stirred at 70 °C for 3 hours under a $N_2$ atmosphere. The reaction mixture was filtered and the filtrate was concentrated under reduced pressure. The resulting residue was diluted with $H_2O$ (20 mL) and extracted with DCM (30 mL * 2). The combined organic layers were dried over $Na_2SO_4$, filtered, and concentrated under reduced pressure and the resulting residue was purified by column chromatography ($SiO_2$, Petroleum ether/Ethyl acetate=50/1 to 1/1). N2-(5-chloropyrazolo[1,5-a]pyrimidin-7-yl)-N6-cyclopropyl-pyridine-2,4,6-triamine (600.0 mg, 94%, 1.90 mmol) was obtained as a deep green solid.

**Step 4:** A mixture of N2-(5-chloropyrazolo[1,5-a]pyrimidin-7-yl)-N6-cyclopropyl-pyridine-2,4,6-triamine (550.0 mg, 1.74 mmol, 1.0 equiv), acrylic acid (0.132 mL, 1.92 mmol, 1.1 equiv), HATU (993 mg, 2.61 mmol, 1.5 equiv), and TEA (0.485 mL, 3.48 mmol, 2.0 equiv) in THF (10.0 mL, 0.174 M) was degassed and purged with $N_2$ for 3 times, and then the mixture was stirred at 15 °C for 12 hours under a $N_2$ atmosphere. The reaction mixture was diluted with $H_2O$ (20 mL) and extracted with EtOAc (30 mL * 2). The combined organic layers were dried over $Na_2SO_4$, filtered, and concentrated under reduced pressure. The resulting residue was purified by prep-HPLC (85% to 10% water (0.05% TFA) to MeOH (0.05% TFA). N-[2-[(5-chloropyrazolo[1,5-a]pyrimidin-7-yl)amino]-6-(cyclopropylamino)-4-pyridyl]prop-2-enamide (47.7 mg, 7%, 129 μmol) was obtained as a light yellow white solid. LC-MS (ESI) m/z = 370.17 $[M+H]^+$. $^1H$ NMR (500 MHz, DMSO) δ 10.26 (d, *J* = 1.7 Hz, 2H), 8.28 (s, 1H), 8.23 (d, *J* = 2.2 Hz, 1H), 7.14 (s, 1H), 7.02 (d, *J* = 1.5 Hz, 1H), 6.80 (d, *J* = 1.5 Hz, 1H), 6.56 (d, *J* = 2.3 Hz, 1H), 6.48 (dd, *J* = 17.0, 10.1 Hz, 1H), 6.29 (dd, *J* = 17.0, 1.9 Hz, 1H), 5.79 (dd, *J* = 10.1, 1.9 Hz, 1H), 2.57 (dt, *J* = 7.0, 3.4 Hz, 1H), 0.83 – 0.76 (m, 2H), 0.54 – 0.48 (m, 2H). $^{13}C$ NMR (126 MHz, DMSO) δ 163.76, 159.30, 151.47, 150.53, 147.83, 147.25, 143.89, 143.13, 131.59, 127.80, 95.58, 93.13, 91.55, 91.30, 23.92, 6.71.

**Synthesis of SCA20: N-(2-((5-chloropyrazolo[1,5-a]pyrimidin-7-yl)amino)-6-(phenylamino)pyridin-4-yl)acrylamide:**

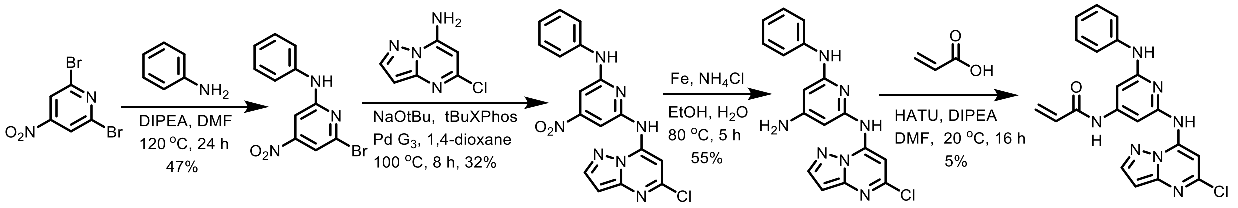

**T17**

**Step 1:** A mixture of 2,6-dibromo-4-nitro-pyridine (4.50 g, 16.0 mmol, 1.0 equiv), aniline (1.60 mL, 17.6 mmol, 1.1 equiv), DIPEA (5.56 mL, 31.9 mmol, 2.0 equiv) in DMF (5.00 mL, 6.38 M) was degassed and purged with $N_2$ for 3 times, and then the mixture was stirred at 120 °C for 24 hours under a $N_2$ atmosphere. The reaction mixture was diluted with $H_2O$ (20 mL) and extracted with EtOAc (30 mL * 2). The combined organic layers were dried over $Na_2SO_4$, filtered, and concentrated under reduced pressure and the resulting residue was purified by column chromatography (SiO2, Petroleum ether/Ethyl acetate=100/1 to 0/1). 6-bromo-4-nitro-N-phenyl-pyridin-2-amine (2.20 g, 47%, 7.48 mmol) was obtained as a yellow solid.

**Step 2:** A mixture of 6-bromo-4-nitro-N-phenyl-pyridin-2-amine (2.00 g, 6.80 mmol, 1.0 equiv) , 5-chloropyrazolo[1,5-a]pyrimidin-7-amine (1.38 g, 8.16 mmol, 1.2 equiv), NaOtBu (980.0 mg, 10.2 mmol, 1.5 equiv), and tBuXPhos Pd $G_3$ (540.0 mg, 680.0 μmol, 0.1 equiv) in 1,4-dioxane (5.00

mL, 1.4 M) was degassed and purged with $N_2$ for 3 times, and then the mixture was stirred at 100 °C for 8 hr under $N_2$ atmosphere. The reaction mixture was diluted with $H_2O$ (20 mL) and extracted with EtOAc (30 mL * 2). The combined organic layers were dried over $Na_2SO_4$, filtered, and concentrated under reduced pressure and the resulting residue was purified by column chromatography ($SiO_2$, Petroleum ether/Ethyl acetate=100/1 to 0/1). N2-(5-chloropyrazolo[1,5-a]pyrimidin-7-yl)-4-nitro-N6-phenyl-pyridine-2,6-diamine (0.82 g, 32%, 2.15 mmol) was obtained as a yellow solid.

**Step 3:** A mixture of N2-(5-chloropyrazolo[1,5-a]pyrimidin-7-yl)-4-nitro-N6-phenyl-pyridine-2,6-diamine (820.0 mg, 2.15 mmol, 1.0 equiv), Fe (599 mg, 10.7 mmol, 5.0 equiv) in EtOH (3.00 mL, 3.57 M) and saturated aqueous $NH_4Cl$ (3 mL) was degassed and purged with $N_2$ for 3 times, and then the mixture was stirred at 80 °C for 5 hours under $N_2$ atmosphere. The reaction mixture was filtered and the filtrate was concentrated under reduced pressure. The resulting residue was diluted with $H_2O$ (20 mL) and extracted with DCM (30 mL * 2). The combined organic layers were dried over $Na_2SO_4$, filtered, and concentrated under reduced pressure and the resulting residue was purified by column chromatography ($SiO_2$, DCM/MeOH=100/1 to 0/1). N2-(5-chloropyrazolo[1,5-a]pyrimidin-7-yl)-N6-phenyl-pyridine-2,4,6-triamine (415 mg, 55%, 1.18 mmol) was obtained as a black solid.

**Step 4:** A mixture of N2-(5-chloropyrazolo[1,5-a]pyrimidin-7-yl)-N6-phenyl-pyridine-2,4,6-triamine (400.0 mg, 1.14 mmol, 1.0 equiv), acrylic acid (78.0 uL, 1.14 mmol, 1.0 equiv), HATU (432 mg, 1.14 mmol, 1.0 equiv), and DIPEA (0.198 mL, 1.14 mmol, 1.0 equiv) in DMF (5.00 mL, 0.228 M) was degassed and purged with $N_2$ for 3 times, and then the mixture was stirred at 20 °C for 16 hours under $N_2$. The reaction mixture was diluted with $H_2O$ (10 mL) and extracted with EtOAc (20 mL * 2). The combined organic layers were dried over $Na_2SO_4$, filtered, and concentrated under reduced pressure and the resulting residue was purified by column chromatography ($SiO_2$, Petroleum ether/Ethyl acetate=100/1 to 0/1). N-[2-anilino-6-[(5-chloropyrazolo[1,5-a]pyrimidin-7-yl)amino]-4-pyridyl]prop-2-enamide (25.0 mg, 5%, 61.6 μmol) was obtained as a yellow solid. LC-MS (ESI) m/z = 406.18 $[M+H]^+$. $^1H$ NMR (500 MHz, DMSO) δ 10.45 (s, 1H), 10.42 (s, 1H), 9.20 (s, 1H), 8.25 (d, *J* = 2.3 Hz, 1H), 7.71 (s, 1H), 7.54 (d, *J* = 7.9 Hz, 2H), 7.31 (t, *J* = 7.7 Hz, 2H), 7.19 (s, 1H), 7.13 (s, 1H), 6.96 (t, *J* = 7.3 Hz, 1H), 6.58 (d, *J* = 2.3 Hz, 1H), 6.56 – 6.46 (m, 1H), 6.32 (dd, *J* = 17.0, 1.9 Hz, 1H), 5.83 (dd, *J* = 10.0, 1.9 Hz, 1H). $^{13}C$ NMR (126 MHz, DMSO) δ 163.92, 155.33, 150.84, 150.48, 148.13, 147.31, 144.05, 143.01, 141.07, 131.46, 128.90, 128.09, 121.10, 119.22, 95.66, 95.50, 94.70, 91.33.

**Synthesis of SCA21 (N-(2-((5-chloropyrazolo[1,5-a]pyrimidin-7-yl)amino)-6-methoxypyridin-4-yl)acrylamide):**

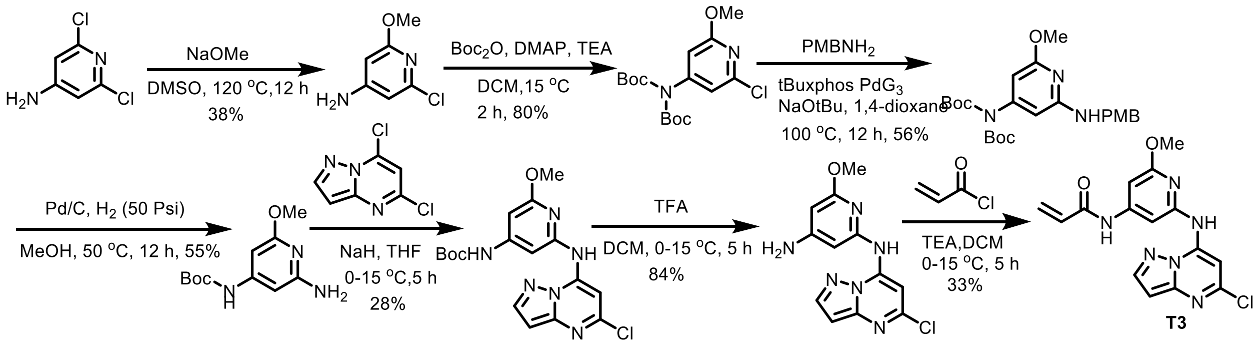

**Step 1:** A mixture of 2,6-dichloropyridin-4-amine (6.00 g, 36.8 mmol, 1.0 equiv), NaOMe (2.98 g, 55.2 mmol, 1.5 equiv) in DMSO (30.0 mL, 1.23 M) was degassed and purged with $N_2$ for 3 times, and then the mixture was stirred at 120 °C for 12 hours under a $N_2$ atmosphere. The reaction

mixture was diluted with $H_2O$ (30 mL) and extracted with EtOAc (30 mL * 3). The combined organic layers were dried over $Na_2SO_4$, filtered, and concentrated under reduced pressure. The resulting residue was purified by column chromatography ($SiO_2$, Petroleum ether/Ethyl acetate=100/1 to 20/1). 2-chloro-6-methoxy-pyridin-4-amine (2.20 g, 38%, 13.9 mmol) was obtained as a purple solid.

**Step 2**: A mixture of 2-chloro-6-methoxy-pyridin-4-amine (2.20 g, 13.9 mmol, 1.0 equiv), tert-butoxycarbonyl tert-butyl carbonate (6.06 g, 27.8 mmol, 6.37 mL, 2.0 equiv), DMAP (847 mg, 6.94 mmol, 0.5 equiv), and TEA (2.81 g, 27.8 mmol, 3.86 mL, 2.0 equiv) in DCM (25.0 mL, 0.556 M) was degassed and purged with $N_2$ for 3 times, and then the mixture was stirred at 15 °C for 2 hours under $N_2$ atmosphere. The reaction mixture was diluted with $H_2O$ (20 mL) and extracted with EtOAc (30 mL * 2). The combined organic layers were dried over $Na_2SO_4$, filtered, and concentrated under reduced pressure to yield a residue. The residue was purified by column chromatography ($SiO_2$, Petroleum ether/Ethyl acetate=100/1 to 10/1). Tert-butyl N-(2-chloro-6-methoxy-4-pyridyl)carbamate (2.90 g, 81%, 11.2 mmol) was obtained as a white solid.

**Step 3:** A mixture of tert-butyl N-tert-butoxycarbonyl-N-(2-chloro-6-methoxy-4-pyridyl)carbamate (4.50 g, 12.5 mmol, 1.0 equiv), $PMBNH_2$ (1.89 g, 13.8 mmol, 1.79 mL, 1.1 equiv), tBuXPhos Pd $G_3$ (996 mg, 1.25 mmol, 0.1 equiv) and NaOtBu (2.41 g, 25.1 mmol, 2.0 equiv) in 1,4-dioxane (25.0 mL, 0.552 M) was degassed and purged with $N_2$ for 3 times, and then the mixture was stirred at 100 °C for 12 hours under $N_2$ atmosphere. The reaction mixture was diluted with $H_2O$ (100 mL) and extracted with EtOAc (30 mL * 3). The combined organic layers were dried over $Na_2SO_4$, filtered, and concentrated under reduced pressure to yield a residue. The residue was purified by column chromatography (SiO2, Petroleum ether/Ethyl acetate=100/1 to 3/1). Tert-butyl N-tert-butoxycarbonyl-N-[2-methoxy-6-[(4-methoxyphenyl)methylamino]-4-pyridyl]carbamate (3.20 g, 55.5%, 6.96 mmol) was obtained as a yellow solid.

**Step 4:** To a solution of Pd/C (1.00 g, 1.67 mmol, 10%, 0.3 equiv) in MeOH (20 mL, 0.278 M) was added isopropyl N-isopropoxycarbonyl-N-[2-methoxy-6-[(4-methoxyphenyl)methylamino]-4-pyridyl]carbamate (2.40 g, 5.56 mmol, 1.0 equiv). The mixture was stirred at 50 °C for 12 hours under $H_2$ (50 Psi). The reaction mixture was filtered and the filtrate was concentrated under reduced pressure. The residue was purified by column chromatography ($SiO_2$, Petroleum ether/Ethyl acetate=100/1 to 0/1). Tert-butyl N-(2-amino-6-hydroxy-4-pyridyl)-N-tert-butoxycarbonyl-carbamate (1.00 g, 55%, 3.07 mmol) was obtained as a white solid.

**Step 5:** To a solution of tert-butyl N-(2-amino-6-methoxy-4-pyridyl)-N-tert-butoxycarbonyl-carbamate (500.0 mg, 1.47 mmol, 1.0 equiv) in THF (5.00 mL, 0.294 M) was added NaH (58.9 mg, 1.47 mmol, 60% purity, 1.0 equiv) slowly under $N_2$ at 0 °C. To the mixture was added 5,7-dichloropyrazolo[1,5-a]pyrimidine (277 mg, 1.47 mmol, 1.0 equiv) and the mixture was stirred at 15 °C for 5 hours. The reaction mixture was quenched by addition of aqueous $NH_4Cl$ (10 mL), and then diluted with $H_2O$ (20 mL) and extracted with EtOAc (30 mL * 2). The combined organic layers were dried over $Na_2SO_4$, filtered, and concentrated under reduced pressure. Tert-butyl N-tert-butoxycarbonyl-N-[2-[(5-chloropyrazolo[1,5-a]pyrimidin-7-yl)amino]-6-methoxy-4-pyridyl]carbamate (200.0 mg, 28%, 407 μmol) was obtained as a white solid. LC-MS (ESI) m/z = 391.2 $[M+H]^+$.

**Step 6:** To a solution of tert-butyl N-tert-butoxycarbonyl-N-[2-[(5-chloropyrazolo[1,5-a]pyrimidin-7-yl)amino]-6-methoxy-4-pyridyl]carbamate (500.0 mg, 1.02 mmol, 1.0 equiv) in DCM (5.00 mL, 0.204 M) was added TFA (0.0754 mL, 1.02 mmol, 1.0 equiv) slowly under $N_2$ at 0 °C. The mixture was stirred at 15 °C for 5 hours. The reaction mixture was diluted with $H_2O$ (20 mL) and extracted with EtOAc (30 mL * 2). The combined organic layers were dried over $Na_2SO_4$, filtered, and concentrated under reduced pressure. N2-(5-chloropyrazolo[1,5-a]pyrimidin-7-yl)-6-methoxy-pyridine-2,4-diamine (250.0 mg, 84%, 859 μmol) was obtained as a white solid.

**Step 7:** To a solution of N2-(5-chloropyrazolo[1,5-a]pyrimidin-7-yl)-6-methoxy-pyridine-2,4-diamine (100.0 mg, 343 μmol, 1.0 equiv) in DCM (5.00 mL, 0.0686 M) was added TEA (0.0957 mL, 688 μmol, 2.0 equiv) slowly under $N_2$ at 0 °C. Acryloyl chloride (0.0281 mL 344 μmol, 1.0 equiv) was then added dropwise and the mixture was stirred at 15 °C for 5 hours. The reaction mixture was concentrated under reduced pressure and the resulting residue was purified by prep-HPLC (85% to 10% water (0.05% TFA) to MeOH (0.05% TFA)) to afford N-[2-[(5-chloropyrazolo[1,5-a]pyrimidin-7-yl)amino]-6-methoxy-4-pyridyl]prop-2-enamide (30.0 mg, 34%, 116 μmol) as a white solid. LC-MS (ESI) m/z = 345.01 $[M+H]^+$. $^1$H NMR (500 MHz, DMSO) δ 10.71 (s, 1H), 10.59 (s, 1H), 8.26 (d, *J* = 2.3 Hz, 1H), 7.90 (s, 1H), 7.45 (d, *J* = 1.5 Hz, 1H), 6.96 (d, *J* = 1.4 Hz, 1H), 6.59 (d, *J* = 2.3 Hz, 1H), 6.49 (dd, *J* = 16.9, 10.1 Hz, 1H), 6.32 (dd, *J* = 17.0, 1.9 Hz, 1H), 5.84 (dd, *J* = 10.1, 1.9 Hz, 1H), 3.92 (s, 3H). $^{13}$C NMR (126 MHz, DMSO) δ 164.48, 163.70, 151.26, 150.77, 150.06, 147.83, 144.58, 143.42, 131.75, 128.90, 97.89, 96.29, 94.23, 91.86, 54.14.

**Synthesis of SCA22 (N-(2-((5-chloropyrazolo[1,5-a]pyrimidin-7-yl)amino)-6-(2-methoxyethoxy)pyridin-4-yl)acrylamide):**

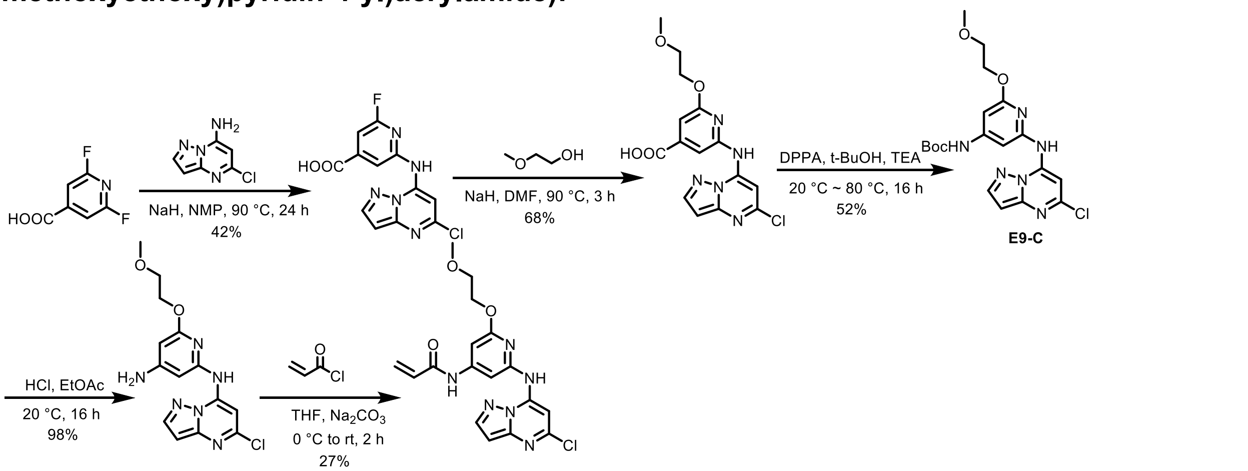

**Step 1:** To a solution of 2,6-difluoroisonicotinic acid (3.00 g, 17.8 mmol, 1.0 equiv) in NMP (30.0 mL) was added NaH (2.14 g, 53.3 mmol, 3.0 equiv) at 0 °C. The mixture was stirred at 0 °C for 30 minutes. Then 5-chloropyrazolo[1,5-a]pyrimidin-7-amine (2.83 g, 17.8 mmol, 1.0 equiv) was added and the mixture was stirred at 90 °C for 24 hours. The reaction mixture was quenched by the addition of $H_2O$ (30.0 mL) at 0 °C, and then adjusted to a pH of 2 with 2 M HCl. The reaction was filtered and product was obtained as a yellow solid. The crude product was triturated with EtOH (10.0 mL) at 20 °C for 1 hour and filtered. 2-((5-chloropyrazolo[1,5-a]pyrimidin-7-yl)amino)-6-fluoroisonicotinic acid (2.30 g, 42%, 7.48 mmol) was obtained as yellow solid. LC-MS (ESI) m/z = 308.1 $[M+H]^+$. $^1$H NMR: 400 MHz, DMSO-$d_6$ δ 13.96 (br s, 1 H), 11.29 (s, 1 H), 8.31 (d, J = 2.0 Hz, 1 H), 8.17 (s, 1 H), 7.83 (s, 1 H), 7.22 (s, 1 H), 6.66 (d, J = 2.0 Hz, 1 H).

**Step 2:** To a solution of 2-methoxyethan-1-ol (247 mg, 3.25 mmol, 2.0 equiv) in DMF (5.00 mL, 0.65 M) was added NaH (260.0 mg, 6.50 mmol, 4.0 equiv) at 0 °C. The mixture was stirred at 0 °C for 30 minutes. Then 2-((5-chloropyrazolo[1,5-a]pyrimidin-7-yl)amino)-6-fluoroisonicotinic acid (500.0 mg, 1.63 mmol, 1.0 equiv) was added, the mixture was stirred at 90 °C for 3 hours. The reaction mixture was quenched by the addition of $H_2O$ (5.00 mL) at 0 °C, and then adjusted to a pH of 2 with 2 M HCl. The reaction was filtered and the crude product was obtained as a yellow solid, which was then triturated with EtOAc (5.00 mL) at 20 °C for 1 hour, filtered, and dried under vacuum. 2-((5-chloropyrazolo[1,5-a]pyrimidin-7-yl)amino)-6-(2-methoxyethoxy)isonicotinic acid (400 mg, 68%, 1.10 mmol) was obtained as yellow solid. LC-MS (ESI) m/z = 364.2 $[M+H]^+$. $^1H$ NMR: 400 MHz, DMSO-$d_6$ δ 13.66 (br s, 1 H), 11.00 (s, 1 H), 8.29 (d, J = 2.4 Hz, 1 H), 7.84 - 7.91 (m, 1 H), 7.70 - 7.82 (m, 1 H), 6.92 (d, J = 0.8 Hz, 1 H), 6.63 (d, J = 2.4 Hz, 1 H), 4.42 - 4.53 (m, 2 H), 3.68 - 3.87 (m, 2 H), 3.33 (s, 3 H).

**Step 3:** To a solution of 2-((5-chloropyrazolo[1,5-a]pyrimidin-7-yl)amino)-6-(2-methoxyethoxy)isonicotinic acid (400.0 mg, 1.10 mmol, 1.0 equiv) in t-BuOH (8.00 mL, 0.138 M) was added DPPA (261 µL, 1.21 mmol, 1.1 equiv), and TEA (183 µL, 1.32 mmol, 1.2 equiv). The mixture was stirred at 20 °C for 1 hour under $N_2$, then warmed to 80 °C for 16 hours. The solvent was removed under vacuum. The residue was purified by column chromatography ($SiO_2$, Petroleum ether/Ethyl acetate = 100/0 to 0/1). Tert-butyl (2-((5-chloropyrazolo[1,5-a]pyrimidin-7-yl)amino)-6-(2-methoxyethoxy)pyridin-4-yl)carbamate (250.0 mg, 52%, 570 µmol) was obtained as white solid. LC-MS (ESI) m/z = 435.1 $[M+H]^+$.

**Step 4:** A mixture of tert-butyl (2-((5-chloropyrazolo[1,5-a]pyrimidin-7-yl)amino)-6-(2-methoxyethoxy)pyridin-4-yl)carbamate (120.0 mg, 275 µmol, 1.0 equiv) in HCl/EtOAc (5.0 mL, 0.055 M) was stirred at 20 °C for 16 hours. The solvent was removed under vacuum. N2-(5-chloropyrazolo[1,5-a]pyrimidin-7-yl)-6-(2-methoxyethoxy)pyridine-2,4-diamine (100 mg, 98%, 269 µmol) was obtained as a white solid. LC-MS (ESI) m/z = 335.1 $[M+H]^+$. $^1H$ NMR: 400 MHz, DMSO-$d_6$ δ 10.29 (br s, 1 H), 8.24 (d, J = 2.0 Hz, 1 H), 7.68 (br s, 1 H), 6.57 (d, J = 2.0 Hz, 1 H), 6.39 (s, 1 H), 5.72 (s, 1 H), 4.68 (br s, 3 H), 4.28 - 4.35 (m, 2 H), 3.59 - 3.76 (m, 2 H), 3.31 (s, 3 H).

**Step 5:** To a solution of N2-(5-chloropyrazolo[1,5-a]pyrimidin-7-yl)-6-(2-methoxyethoxy)pyridine-2,4-diamine (100.0 mg, 269 µmol, 1.0 equiv) in THF (5.00 mL, 0.0538 M) was added $Na_2CO_3$ (114 mg, 1.08 mmol, 4.0 equiv) and acryloyl chloride (43.7 µL, 538 µmol, 2.0 equiv) in THF (1.00 mL) at 0 °C. The mixture was stirred at 20 °C for 2 hours. The mixture was poured into ice-water (25.0 mL) and stirred for 30 minutes, which resulted in the formation of a white precipitate that was filtered and dried under vacuum. N-(2-((5-chloropyrazolo[1,5-a]pyrimidin-7-yl)amino)-6-(2-methoxyethoxy)pyridin-4-yl)acrylamide (30.0 mg, 27%, 73.7 µmol) was obtained as an off-white solid. LC-MS (ESI) m/z = 389.13 $[M+H]^+$. $^1H$ NMR: 400 MHz, DMSO-$d_6$ δ 10.76 (br s, 1 H), 10.54 (s, 1 H), 8.24 (s, 1 H), 7.73 (br s, 1 H), 7.39 (br s, 1 H), 6.96 (s, 1 H), 6.57 (br s, 1 H), 6.43 - 6.52 (m, 1 H), 6.27 - 6.36 (m, 1 H), 5.79 - 5.88 (m, 1 H), 4.29 - 4.55 (m, 2 H), 3.62 - 3.80 (m, 2 H), 3.32 - 3.33 (s, 3 H). $^{13}C$ NMR (126 MHz, DMSO) δ 164.03, 162.71, 150.87, 150.30, 149.64, 147.43, 144.08, 143.15, 131.27, 128.48, 97.68, 95.73, 94.09, 91.14, 70.32, 65.16, 58.21.

**Synthesis of SCA23 (N-(2-((5-chloropyrazolo[1,5-a]pyrimidin-7-yl)amino)-6-(2-hydroxyethoxy)pyridin-4-yl)acrylamide):**

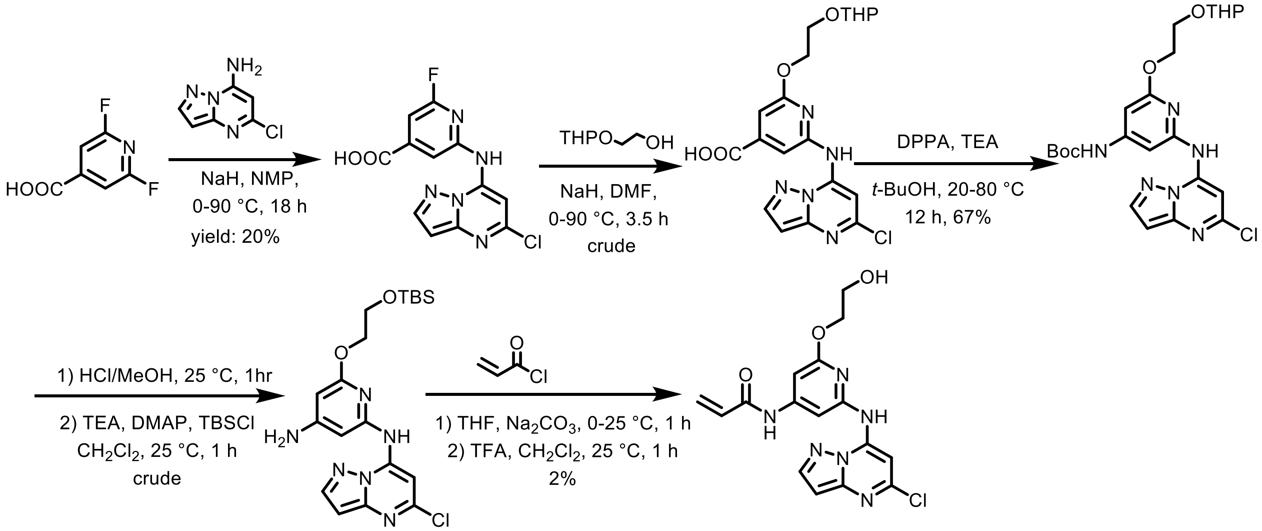

**Step 1:** To a solution of 2,6-difluoroisonicotinic acid (5.30 g, 31.4 mmol, 1.0 equiv) in NMP (40.0 mL) was added NaH (3.77 g, 94.3 mmol, 60.0% purity, 3.00 equiv) at 0 °C under $N_2$ for 1 hour. Then 5-chloropyrazolo[1,5-a]pyrimidin-7-amine (5.00 g, 31.4 mmol, 1.0 equiv) was added. The mixture was stirred at 90 °C for 17 hours. The reaction mixture was quenched by the addition of $H_2O$ (30 mL) at 0 °C, and then adjusted to a pH of 2 with 2 M HCl. Product was filtered and obtained as a yellow solid. The crude product 2-((5-chloropyrazolo[1,5-a]pyrimidin-7-yl)amino)-6-fluoroisonicotinic acid (1.94 g, 20.1%, 6.31 mmol) was used into the next step without further purification. MS (ESI) m/z = 307.8 $[M+H]^+$. $^1H$ NMR: 400 MHz, DMSO-$d_6$ δ 11.3 (s, 1H), 8.30 (d, J = 2.4 Hz, 1H), 8.16 (s, 1H), 7.83 (s, 1H), 7.21 (d, J = 1.2 Hz, 1H), 6.65 (d, J = 2.4 Hz, 1H).

**Step 2:** To a solution of 2-((5-chloropyrazolo[1,5-a]pyrimidin-7-yl)amino)-6-fluoroisonicotinic acid (2.77 g, 18.9 mmol, 2.57 mL, 3.0 equiv) in DMF (15.0 mL, 1.26 M) was added NaH (1.51 g, 37.8 mmol, 6.0 equiv) at 0 °C under $N_2$ for 0.5 hours. Then 2-((tetrahydro-2H-pyran-2-yl)oxy)ethan-1-ol (1.94 g, 6.31 mmol, 1.0 equiv) was added at 0 °C. The mixture was stirred at 90 °C for 3 hours. The reaction mixture was quenched by the addition of $H_2O$ (15.0 mL) at 0 °C under $N_2$ for 0.5 hours, and then adjusted to a pH of 2 with 2M HCl. The product was then filtered and 2-((5-chloropyrazolo[1,5-a]pyrimidin-7-yl)amino)-6-(2-((tetrahydro-2H-pyran-2-yl)oxy)ethoxy)isonicotinic acid (2.80 g, crude) was obtained as a yellow solid and used in the next step without further purification. MS (ESI) m/z = 433.8 $[M+H]^+$. $^1H$ NMR: 400 MHz, DMSO-$d_6$ δ 11.2 - 10.7 (m, 1H), 8.28 (d, J = 2.2 Hz, 1H), 7.88 (s, 1H), 7.74 (s, 1H), 6.92 (s, 1H), 6.62 (d, J = 2.2 Hz, 1H), 4.68 (br s, 1H), 4.57 - 4.48 (m, 2H), 4.03 - 3.95 (m, 1H), 3.84 - 3.72 (m, 3H), 1.68 - 1.58 (m, 2H), 1.49 - 1.41 (m, 4H).

**Step 3:** To a solution of 2-((5-chloropyrazolo[1,5-a]pyrimidin-7-yl)amino)-6-(2-((tetrahydro-2H-pyran-2-yl)oxy)ethoxy)isonicotinic acid (2.80 g, 6.45 mmol, 1.0 equiv) in t-BuOH (12.0 mL, 0.54 M) was added DPPA (1.53 mL, 7.10 mmol, 1.1 equiv) and TEA (1.08 mL, 7.74 mmol, 1.2 equiv) at 20 °C under $N_2$ for 1 hour. The mixture was stirred at 80 °C for 12 hours. The reaction mixture was concentrated under reduced pressure. The residue was purified by column chromatography ($SiO_2$, petroleum ether: ethyl acetate = 1/0 to 50/1). Tert-butyl (2-((5-chloropyrazolo[1,5-a]pyrimidin-7-yl)amino)-6-(2-((tetrahydro-2H-pyran-2-yl)oxy)ethoxy)pyridin-4-yl)carbamate (1.82 g, 67%, 4.32 mmol) was obtained as a yellow solid. MS (ESI) m/z = 505.0 $[M+H]^+$. $^1H$ NMR: 400 MHz, $CDCl_3$-d δ 8.88 - 8.71 (m, 1H), 8.04 (d, J = 2.4 Hz, 1H), 7.82 (s, 1H), 7.09 (d, J = 1.4 Hz,

1H), 6.68 (s, 1H), 6.53 (d, J = 2.4 Hz, 1H), 6.32 (d, J = 1.4 Hz, 1H), 4.76 - 4.72 (m, 1H), 4.60 - 4.56 (m, 2H), 4.15 - 4.11 (m, 1H), 1.65 (br s, 2H), 1.63 - 1.60 (m, 4H), 1.59 - 1.50 (m, 18H).

**Step 4:** A solution of tert-butyl (2-((5-chloropyrazolo[1,5-a]pyrimidin-7-yl)amino)-6-(2-((tetrahydro-2H-pyran-2-yl)oxy)ethoxy)pyridin-4-yl)carbamate (1.82 g, 3.98 mmol, 1.0 equiv) in HCl/MeOH (995 µL, 4.0 M, 1.0 equiv) was stirred at 25 °C for 1 hour. The mixture was concentrated under reduced pressure, then dissolved in DCM (15.0 mL), to which TEA (1.66 mL, 11.9 mmol, 3.0 equiv), DMAP (97.2 mg, 796 µmol, 0.20 equiv), and TBSCl (539 µL, 4.38 mmol, 1.1 equiv) were added. The mixture was stirred at 25 °C for 1 hour. The reaction mixture was quenched by the addition of $H_2O$ (20 mL) at 25 °C, and then extracted with DCM (10 mL x 3). The combined organic layers were washed with brine (20 mL), dried over $Na_2SO_4$, filtered, and concentrated under reduced pressure. The crude product 6-(2-((tert-butyldimethylsilyl)oxy)ethoxy)-N2-(5-chloropyrazolo[1,5-a]pyrimidin-7-yl)pyridine-2,4-diamine (2.56 g, crude) was used in the next step without further purification. 6-(2-((tert-butyldimethylsilyl)oxy)ethoxy)-N2-(5-chloropyrazolo[1,5-a]pyrimidin-7-yl)pyridine-2,4-diamine (2.56 g, crude) was obtained as a green solid. MS (ESI) m/z = 435.0 $[M+H]^+$.

**Step 5:** To a solution of 6-(2-((tert-butyldimethylsilyl)oxy)ethoxy)-N2-(5-chloropyrazolo[1,5-a]pyrimidin-7-yl)pyridine-2,4-diamine (1.50 g, 3.45 mmol, 1.0 equiv), $Na_2CO_3$ (1.10 g, 10.3 mmol, 3.0 equiv) in THF (15.0 mL, 0.23 M) was added acryloyl chloride (210 µL, 2.59 mmol, 0.750 equiv) at 0 °C. The mixture was stirred at 25 °C for 1 hour. Then the reaction mixture was concentrated under reduced pressure, dissolved in DCM (1.00 mL), to which TFA (0.330 mL) was added. The reaction was allowed to stir at 25 °C for 1 hour and was then concentrated under reduced pressure. The residue was purified by prep-HPLC (85% to 10% water (0.05% TFA) to MeOH (0.05% TFA). (N-(2-((5-chloropyrazolo[1,5-a]pyrimidin-7-yl)amino)-6-(2-hydroxyethoxy)pyridin-4-yl)acrylamide (39.0 mg, 2%, 79.3 µmol) was obtained as a white solid (TFA salt). MS (ESI) m/z = 374.8 $[M+H]^+$. $^1$H NMR (500 MHz, DMSO) δ 10.69 (s, 1H), 10.53 (s, 1H), 8.25 (d, *J* = 2.2 Hz, 1H), 7.81 (s, 1H), 7.42 (d, *J* = 1.5 Hz, 1H), 6.98 (d, *J* = 1.4 Hz, 1H), 6.58 (d, *J* = 2.2 Hz, 1H), 6.48 (dd, *J* = 17.0, 10.1 Hz, 1H), 6.32 (dd, *J* = 17.1, 1.9 Hz, 1H), 5.84 (dd, *J* = 10.1, 1.9 Hz, 1H), 4.92 (t, *J* = 5.5 Hz, 1H), 4.34 – 4.28 (m, 2H), 3.78 (q, *J* = 4.9 Hz, 2H). $^{13}$C NMR (126 MHz, DMSO) δ 164.48, 163.44, 151.10, 150.78, 150.04, 147.83, 144.58, 143.44, 131.74, 128.93, 97.88, 96.28, 94.63, 91.69, 68.57, 59.98.

**Synthesis of SCA24: N-(2-((5-chloropyrazolo[1,5-a]pyrimidin-7-yl)amino)-6-((2-(dimethylamino)ethyl)amino)pyridin-4-yl)acrylamide:**

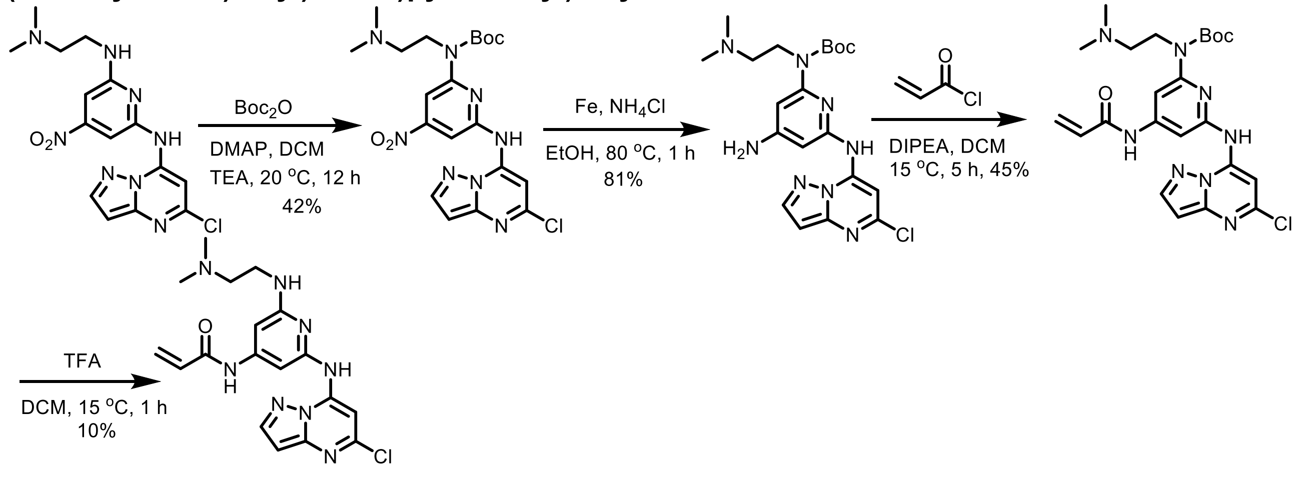

**Step 1:** To a solution of N2-(5-chloropyrazolo[1,5-a]pyrimidin-7-yl)-N6-[2-(dimethylamino)ethyl]-4-nitro-pyridine-2,6-diamine (1.50 g, 3.98 mmol, 1.0 equiv) in DCM (20.0 mL, 0.2 M) was added TEA (2.22 mL, 15.92 mmol, 4.0 equiv), DMAP (535 mg, 4.38 mmol, 1.1 equiv) and tert-butoxycarbonyl tert-butyl carbonate (4.57 mL, 19.9 mmol, 5.0 equiv) under $N_2$. The mixture was stirred at 20 °C for 12 hours under $N_2$. The reaction mixture was concentrated under reduced pressure and the resulting residue was purified by column chromatography ($SiO_2$, Petroleum ether/Ethyl acetate=1/0 to 0/1). Tert-butyl-N-[6-[tert-butoxycarbonyl-(5-chloropyrazolo[1,5-a]pyrimidin-7-yl)amino]-4-nitro-2-pyridyl]-N-[2-(dimethylamino)ethyl]carbamate (0.400 g, 16%, 645 µg) and tert-butyl-N-[6-[(5-chloropyrazolo[1,5-a]pyrimidin-7-yl)amino]-4-nitro-2-pyridyl]-N-[2-(dimethylamino)ethyl]carbamate (0.800 g, 42%, 1.68 mmol) were obtained as a white solid.

**Step 2:** To a solution of tert-butyl (4-amino-6-((5-chloropyrazolo[1,5-a]pyrimidin-7-yl)amino)pyridin-2-yl)(2-(dimethylamino)ethyl)carbamate (0.800 g, 1.39 mmol, 1.0 equiv) in saturated aqueous $NH_4Cl$ (5.00 mL, 0.278 M) and EtOH (5.00 mL, 0.278 M) was added Fe (232 mg, 4.16 mmol, 3.0 equiv) under $N_2$. The mixture was stirred at 80 °C for 1 hour under $N_2$. The reaction was filtered and concentrated under reduced pressure and the resulting residue was purified by column chromatography ($SiO_2$, Petroleum ether/Ethyl acetate=50/1 to 1/1). Tert-butyl N-[4-amino-6-[(5-chloropyrazolo[1,5-a]pyrimidin-7-yl)amino]-2-pyridyl]-N-[2-(dimethylamino)ethyl]carbamate (0.500 g, 81%, 1.12 mmol) was obtained as a white solid.

**Step 3:** To a solution of tert-butyl N-[4-amino-6-[(5-chloropyrazolo[1,5-a]pyrimidin-7-yl)amino]-2-pyridyl]-N-[2-(dimethylamino)ethyl]carbamate (500.0 mg, 1.12 mmol, 1.0 equiv) in DCM (20.0 mL, 0.056 M) was added DIPEA (195 uL, 1.12 mmol, 1.0 equiv) and acryloyl chloride (91.2 uL, 1.12 mmol, 1.0 equiv). The mixture was stirred at 15 °C for 5 hour. The reaction was filtered and concentrated under reduced pressure and the resulting residue was purified by column chromatography (SiO2, Petroleum ether/Ethyl acetate=50/1 to 1/1). Tert-butyl N-[6-[(5-chloropyrazolo[1,5-a]pyrimidin-7-yl)amino]-4-(prop-2-enoylamino)-2-pyridyl]-N-[2-(dimethylamino)ethyl]carbamate (250.0 mg, 45%, 499 µmol) was obtained as a white solid. LC-MS (ESI) m/z = 501.2 $[M+H]^+$.

**Step 4:** A mixture of tert-butyl N-[6-[(5-chloropyrazolo[1,5-a]pyrimidin-7-yl)amino]-4-(prop-2-enoylamino)-2-pyridyl]-N-[2-(dimethylamino)ethyl]carbamate (250.0 mg, 499 µmol, 1.0 equiv) in DCM (5.00 mL, 0.0998 M) and TFA (5.00 mL, 0.0998 M) was degassed and purged with $N_2$ for 3 times, and then the mixture was stirred at 15 °C for 1 hour under $N_2$ atmosphere. The reaction mixture was filtered and concentrated under reduced pressure and the resulting residue was purified by prep-HPLC (85% to 10% water (0.05% TFA) to MeOH (0.05% TFA). N-[2-[(5-chloropyrazolo[1,5-a]pyrimidin-7-yl)amino]-6-[2-(dimethylamino)ethylamino]-4-pyridyl]prop-2-enamide (0.0200 g, 10%, 49.9 µmol) was obtained as a white solid. LC-MS (ESI) m/z: 401.18 $[M+H]^+$. NMR. $^1H$ NMR (500 MHz, DMSO) δ 10.43 (s, 1H), 10.35 (s, 1H), 9.51 (s, 1H), 8.26 (d, *J* = 2.3 Hz, 1H), 7.70 (s, 1H), 7.05 (s, 1H), 6.95 – 6.90 (m, 2H), 6.58 (d, *J* = 2.3 Hz, 1H), 6.49 (dd, *J* = 17.0, 10.1 Hz, 1H), 6.29 (dd, *J* = 17.0, 1.9 Hz, 1H), 5.81 (dd, *J* = 10.1, 1.9 Hz, 1H), 3.62 (q, *J* = 5.3 Hz, 2H), 3.33 (q, *J* = 5.7 Hz, 2H), 2.84 (d, *J* = 3.9 Hz, 6H). $^{13}C$ NMR (126 MHz, DMSO) δ 163.92, 157.98, 151.05, 150.16, 148.02, 147.42, 144.14, 143.34, 131.48, 128.02, 95.70, 94.01, 93.07, 90.80, 55.73, 42.53, 36.53.

**Synthesis of SCA25 (N-(2-((5-chloropyrazolo[1,5-a]pyrimidin-7-yl)amino)-6-fluoropyridin-4-yl)acrylamide):**

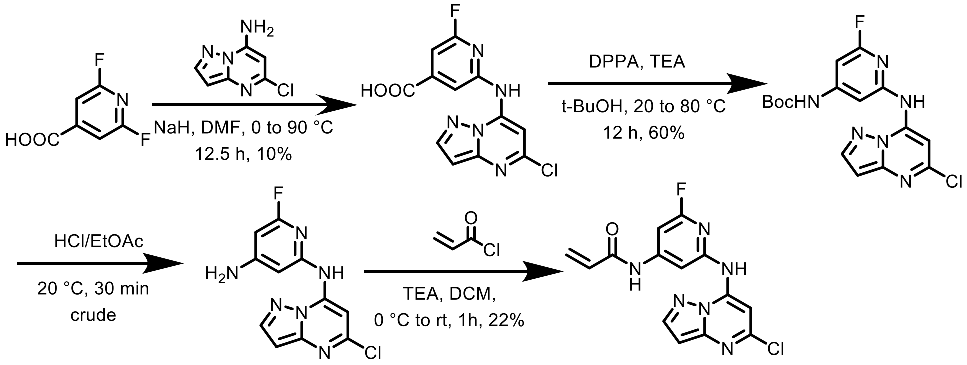

**Step 1:** To a solution of compound 2,6-difluoroisonicotinic acid (349 mg, 2.07 mmol, 1.1 equiv) in DMF (3.00 mL, 0.69 M) was added NaH (226 mg, 5.66 mmol, 3.0 equiv) at 0 °C. The mixture was stirred at 0 °C for 0.5 hour. Then 5-chloropyrazolo[1,5-a]pyrimidin-7-amine (300.0 mg, 1.89 mmol, 1.0 equiv) was added and the mixture was stirred at 90 °C for 12 hours. The reaction mixture was quenched by addition of $H_2O$ (4.00 mL) at 0 °C, and then adjusted to pH = 2 with 2M HCl. The solution was filtered and 2-((5-chloropyrazolo[1,5-a]pyrimidin-7-yl)amino)-6-fluoroisonicotinic acid (60.0 mg, 10%, 195 µmol) was obtained as a yellow solid. The residue was used directly in the next step. LC-MS (ESI) m/z = 308.0 $[M+H]^+$.

**Step 2:** To a solution of 2-((5-chloropyrazolo[1,5-a]pyrimidin-7-yl)amino)-6-fluoroisonicotinic acid (150.0 mg, 487 µmol, 1.0 equiv) in t-BuOH (3.00 mL, 0.16 M) was added DPPA (0.115 mL, 536 µmol, 1.1 equiv) and $Et_3N$ (0.0746 mL, 536 µmol, 1.1 equiv) at 20 °C. The mixture was stirred at 20 °C for 1 hour. Then the mixture was stirred at 80 °C for 11 hours. The reaction mixture was concentrated under reduced pressure to remove solvent. The residue was diluted with $NaHCO_3$ (aq, 5.00 mL) and extracted with EtOAc (5.00 mL * 3), dried over $Na_2SO_4$, filtered and concentrated under reduced pressure. The residue was purified by prep-TLC ($SiO_2$, DCM/MeOH = 1/0). Tert-butyl (2-((5-chloropyrazolo[1,5-a]pyrimidin-7-yl)amino)-6-fluoropyridin-4-yl)carbamate (110.0 mg, 60%, 290.0 µmol) was obtained as a yellow solid. LC-MS (ESI) m/z = 379.1 $[M+H]^+$.

**Step 3:** A mixture of tert-butyl (2-((5-chloropyrazolo[1,5-a]pyrimidin-7-yl)amino)-6-fluoropyridin-4-yl)carbamate (110 mg, 290 µmol, 1.0 equiv) in HCl/EtOAc (2.00 mL, 0.15 M) was stirred at 20 °C for 30 minutes under $N_2$ atmosphere. The reaction mixture was concentrated under reduced pressure to yield a residue. The residue was used directly in the next step. N2-(5-chloropyrazolo[1,5-a]pyrimidin-7-yl)-6-fluoropyridine-2,4-diamine (100 mg, crude, HCl) was obtained as a white solid. LC-MS (ESI) m/z = 278.9 $[M+H]^+$.

**Step 4:** To a solution of N2-(5-chloropyrazolo[1,5-a]pyrimidin-7-yl)-6-fluoropyridine-2,4-diamine (100.0 mg, 317 µmol, 1.0 equiv) in DCM (1.00 mL) was added TEA (88.3 µL, 634 µmol, 2.0 equiv) at 0 °C. Then acryloyl chloride (23.2 µL, 285 µmol, 0.9 equiv) was added. The mixture was stirred at 20 °C for 1 hour. The reaction mixture was quenched with $H_2O$ (5.00 mL) and extracted with DCM (5.00 mL * 3). The combined organic layers were concentrated under reduced pressure. The crude product was purified by reversed-phase HPLC (0.1% FA condition). (N-(2-((5-chloropyrazolo[1,5-a]pyrimidin-7-yl)amino)-6-fluoropyridin-4-yl)acrylamide) (24.4 mg, 22%, 69.6 µmol) was obtained as a white solid. LC-MS (ESI) m/z = 333.06 $[M+H]^+$. $^1$H NMR (500 MHz, DMSO) δ 11.03 (s, 1H), 10.87 (s, 1H), 8.28 (d, *J* = 2.3 Hz, 1H), 7.79 (t, *J* = 1.1 Hz, 1H), 7.75 (s, 1H), 7.18 (d, *J* = 1.3 Hz, 1H), 6.62 (d, *J* = 2.2 Hz, 1H), 6.49 (dd, *J* = 16.9, 10.1 Hz, 1H), 6.36 (dd, *J* = 17.1, 1.8 Hz, 1H), 5.89 (dd, *J* = 10.0, 1.8 Hz, 1H). $^{13}$C NMR (126 MHz, DMSO) δ 164.21,

161.93 (d, $J$ = 231.8 Hz), 151.28 (d, $J$ = 3.4 Hz), 151.26 (d, $J$ = 27.2 Hz), 150.29, 147.32, 144.22, 142.71, 130.95, 129.13, 101.19 (d, $J$ = 4.0 Hz), 95.99, 92.97, 92.64, 91.49.

**Synthesis of SCA26 (N-(2-((5-chloropyrazolo[1,5-a]pyrimidin-7-yl)amino)-5-methylpyridin-4-yl)acrylamide):**

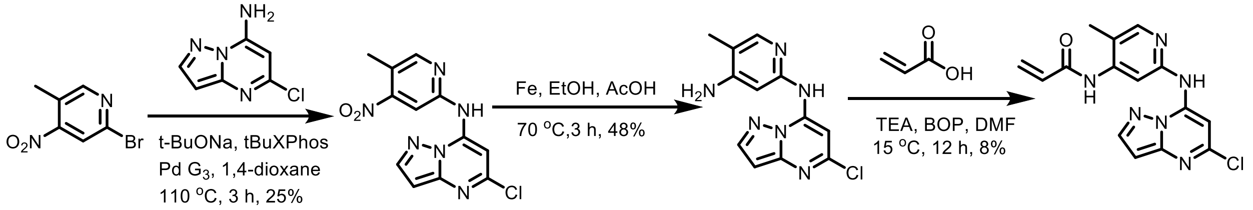

**Step 1:** A mixture of 2-bromo-5-methyl-4-nitro-pyridine (2.00 g, 9.22 mmol, 1.0 equiv), 5-chloropyrazolo[1,5-a]pyrimidin-7-amine (1.86 g, 11.1 mmol, 1.2 equiv), t-BuONa (1.77 g, 18.4 mmol, 2.0 equiv) and tBuXPhos Pd $G_3$ (732 mg, 922 µmol, 0.1 equiv) in 1,4-dioxane (35.0 mL, 0.263 M) was degassed and purged with $N_2$ for 3 times, and then the mixture was stirred at 110 °C for 3 hours under a $N_2$ atmosphere. The reaction mixture was diluted with $H_2O$ (40 mL) and extracted with EtOAc (50 mL * 2). The combined organic layers were dried over $Na_2SO_4$, filtered, and concentrated under reduced pressure. The residue was purified by column chromatography ($SiO_2$, Petroleum ether/Ethyl acetate=100/1 to 0/1). Compound 5-chloro-N-(5-methyl-4-nitro-2-pyridyl)pyrazolo[1,5-a]pyrimidin-7-amine (700.0 mg, 25%, 2.30 mmol) was obtained as a yellow solid. LC-MS (ESI) m/z = 302.7 $[M+H]^+$.

**Step 2:** A mixture of 5-chloro-N-(5-methyl-4-nitro-2-pyridyl)pyrazolo[1,5-a]pyrimidin-7-amine (700.0 mg, 2.30 mmol, 1.0 equiv), Fe (641.49 mg, 11.49 mmol, 5.0 equiv) in EtOH (8.00 mL, 0.288 M) and AcOH (8.00 mL, 0.288 M) was degassed and purged with $N_2$ for 3 times, and then the mixture was stirred at 70 °C for 3 hours under a $N_2$ atmosphere. The reaction mixture was filtered and the filtrate was concentrated under reduced pressure. The resulting residue was diluted with $H_2O$ (20 mL) and extracted with DCM (60 mL, 30 mL * 2). The combined organic layers were dried over $Na_2SO_4$, filtered, and concentrated under reduced pressure. The resulting residue was purified by column chromatography ($SiO_2$, Petroleum ether/Ethyl acetate=100/1 to 0/1). N2-(5-chloropyrazolo[1,5-a]pyrimidin-7-yl)-5-methyl-pyridine-2,4-diamine (300.0 mg, 48% 1.09 mmol) was obtained as a yellow solid. LC-MS (ESI) m/z = 275.0 $[M+H]^+$.

**Step 3:** A mixture of N2-(5-chloropyrazolo[1,5-a]pyrimidin-7-yl)-5-methyl-pyridine-2,4-diamine (200.0 mg, 728 µmol, 1.0 equiv), acrylic acid (54.9 uL, 801 µmol, 1.1 equiv), BOP (483 mg, 1.09 mmol, 1.5 equiv), and TEA (203 uL, 1.46 mmol, 2.0 equiv) in DMF (3.00 mL, 0.243 M) was degassed and purged with $N_2$ for 3 times, and then the mixture was stirred at 15 °C for 12 hours under $N_2$ atmosphere. The reaction mixture was concentrated under reduced pressure. The resulting residue was purified by prep-HPLC (85% to 10% water (0.05% TFA) to MeOH (0.05% TFA)) to afford N-[2-[(5-chloropyrazolo[1,5-a]pyrimidin-7-yl)amino]-5-methyl-4-pyridyl]prop-2-enamide (20.0 mg, 8%, 60.8 µmol) as a white solid. LC-MS (ESI) m/z = 329.05 $[M+H]^+$. $^1$H NMR (500 MHz, DMSO) δ 10.66 (s, 1H), 9.58 (s, 1H), 8.25 (d, $J$ = 2.1 Hz, 2H), 8.23 (s, 1H), 7.88 (s, 1H), 6.70 (dd, $J$ = 16.9, 10.2 Hz, 1H), 6.58 (d, $J$ = 2.2 Hz, 1H), 6.34 (dd, $J$ = 17.1, 1.9 Hz, 1H), 5.85 (dd, $J$ = 10.2, 1.9 Hz, 1H), 2.24 (s, 3H). $^{13}$C NMR (126 MHz, DMSO) δ 163.78, 151.15, 150.38, 148.31, 147.31, 145.31, 144.08, 143.22, 131.31, 128.23, 120.04, 107.43, 95.72, 90.60, 14.41.

**Synthesis of Desthiobiotinylated SCA1 (N-(2-(2-(3-((3-(3-acrylamido-5-((5-chloropyrazolo[1,5-a]pyrimidin-7-yl)amino)phenyl)propyl)amino)-3-oxopropoxy)ethoxy)ethyl)-6-((4R,5S)-5-methyl-2-oxoimidazolidin-4-yl)hexanamide):**

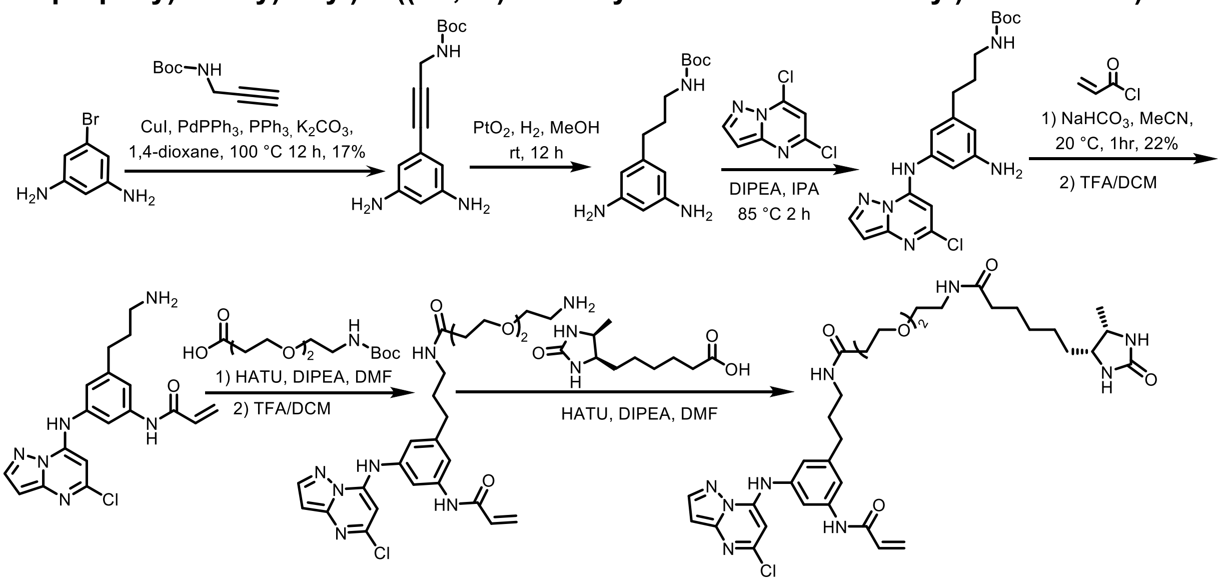

**Step 1**: A solution of 5-bromobenzene-1,3-diamine (100.0 mg, 0.400 mmol, 1.0 equiv), tert-butyl prop-2-yn-1-ylcarbamate (93.0 mg, 0.600 mmol, 1.5 equiv), copper iodide (15.0 mg, 0.0800 mmol, 0.2 equiv) $PdPPh_3$ (15.0 mg, 0.0800 mmol, 0.2 equiv), $PPh_3$ (21.0 mg, 0.0800 mmol, 0.2 equiv), and potassium carbonate (116 mg, 1.20 mmol, 3 equiv) in 1,4-dioxane (4.00 mL, 1.0 M) was flushed with nitrogen for 15 minutes and then heated at 100 ˚C for 12 hours. The mixture was allowed to cool, concentrated *in vacuo*, and purified by normal phase chromatography (0:40% DCM: MeOH) to afford tert-butyl (3-(3,5-diaminophenyl)prop-2-yn-1-yl)carbamate (18.0 mg, 17%, 0.0689 mmol). LC-MS (ESI) m/z = 261.88 $[M+H]^+$.

**Step 2:** To tert-butyl (3-(3,5-diaminophenyl)prop-2-yn-1-yl)carbamate (18.0 mg, 0.112 mmol, 1.0 equiv) was added $PtO_2$ (2.54 mg, 0.0112 mmol, 0.1 equiv). Upon addition of MeOH (1.12 mL, 0.1 M), the reaction was vacuum-flushed three times with a hydrogen balloon and allowed to stir overnight under hydrogen gas. The reaction was then filtered over cellite and concentrated as a crude residue to afford tert-butyl (3-(3-amino-5-((5-chloropyrazolo[1,5-a]pyrimidin-7-yl)amino)phenyl)propyl)carbamate.

**Step 3**: To a solution of tert-butyl (3-(3,5-diaminophenyl)propyl)carbamate (256 mg, 0.960 mmol, 1.0 equiv) and 5,7-dichloropyrazolo[1,5-a]pyrimidine (181 mg, 0.960 mmol, 1.0 equiv) in IPA (4.00 mL, 0.24 M) was added DIPEA (0.464 mL, 2.88 mmol, 3.0 equiv) and the reaction was allowed to stir at 85 ˚C for two hours. The reaction was then concentrated *in vacuo* and purified by normal phase chromatography (0:40% DCM: MeOH) to afford tert-butyl (3-(3-amino-5-((5-chloropyrazolo[1,5-a]pyrimidin-7-yl)amino)phenyl)propyl)carbamate. LC-MS (ESI) m/z = 417.27 $[M+H]^+$.

**Step 4:** To a solution of tert-butyl (3-(3-amino-5-((5-chloropyrazolo[1,5-a]pyrimidin-7-yl)amino)phenyl)propyl)carbamate (20.0 mg, 0.0480 mmol, 1.0 equiv) and sodium bicarbonate (12.0 mg, 0.140 mmol, 3.0 equiv) dissolved in THF (2.40 mL, 0.02 M) at 0˚C under nitrogen gas was added acryloyl chloride (0.00390 mL, 0.0480 mmol, 1.0 equiv) diluted in 1 mL acetonitrile, dropwise. The reaction was allowed to stir for 30 minutes, and was then purified by HPLC. Subsequently, it was dissolved in 4 mL DCM, 1 mL TFA, and allowed to stir for 1 hour. The mixture was then concentrated *in vacuo* and placed under vacuum for 12 hours to afford N-(3-(3-

aminopropyl)-5-((5-chloropyrazolo[1,5-a]pyrimidin-7-yl)amino)phenyl)acrylamide as a crude product.

**Step 5:** To a solution of 2,2-dimethyl-4-oxo-3,8,11,14-tetraoxa-5-azaheptadecan-17-oic acid (22.4 mg, 0.0809 mmol, 1.0 equiv) in DMF (1.62 mL, 0.05 M) and DIPEA (0.0705 mL, 0.404 mmol, 5.0 equiv) at 0 ˚C was added HATU (61.5 mg, 0.162 mmol, 2.0 equiv). The reaction was allowed to stir for 15 minutes before crude tert-butyl (2-(2-(3-((3-(3-acrylamido-5-((5-chloropyrazolo[1,5-a]pyrimidin-7-yl)amino)phenyl)propyl)amino)-3-oxopropoxy)ethoxy)ethyl)carbamate (30.0 mg, 0.809 mmol, 1.0 equiv) was added to the solution. The reaction was allowed to stir for 1 hour, at which point it was dissolved in 10 mL of water. The product was extracted using 3 x 20 mL ethyl acetate. The organic layer was washed with brine, dried over $Na_2SO4$, and concentrated *in vacuo*. The resulting residue was then dissolved in 4.00 mL DCM and 1.00 mL TFA and allowed to stir for 1 hour, at which point the reaction was concentrated *in vacuo* and placed under vacuum for 12 hours to afford N-(3-(3-(3-(2-(2-aminoethoxy)ethoxy)propanamido)propyl)-5-((5-chloropyrazolo[1,5-a]pyrimidin-7-yl)amino)phenyl)acrylamide as a crude product. LC-MS (ESI) m/z = 530.27 $[M+H]^+$.

**Step 6:** To a solution of 6-((4R,5S)-5-methyl-2-oxoimidazolidin-4-yl)hexanoic acid (11.2 mg, 0.0523 mmol, 1.0 equiv) in DMF (1.05 mL, 0.05 M) and DIPEA (0.0455 mL, 0.261 mmol, 5.0 equiv) at 0 ˚C was added HATU (39.7 mg, 0.105 mmol, 2.0 equiv). The reaction was allowed to stir for 15 minutes before crude N-(3-(3-(3-(2-(2-aminoethoxy)ethoxy)propanamido)propyl)-5-((5-chloropyrazolo[1,5-a]pyrimidin-7-yl)amino)phenyl)acrylamide (30.0 mg, 0.0524 mmol, 1.0 equiv) was added to the solution. The reaction was allowed to stir for 1 hour and was purified by prep-HPLC (85% to 10% water (0.05% TFA) to MeOH (0.05% TFA)) to afford N-(2-(2-(3-((3-(3-acrylamido-5-((5-chloropyrazolo[1,5-a]pyrimidin-7-yl)amino)phenyl)propyl)amino)-3-oxopropoxy)ethoxy)ethyl)-6-((4R,5S)-5-methyl-2-oxoimidazolidin-4-yl)hexanamide (8.41 mg, 21%, 0.0109 mmol) as a white solid. LC-MS (ESI) m/z = 726.35 $[M+H]^+$. $^1H$ NMR (500 MHz, DMSO) δ 10.32 (s, 1H), 10.27 (s, 1H), 8.24 (d, *J* = 2.3 Hz, 1H), 7.89 (t, *J* = 5.6 Hz, 1H), 7.79 (t, *J* = 5.7 Hz, 1H), 7.63 (t, *J* = 1.9 Hz, 1H), 7.46 (t, *J* = 1.7 Hz, 1H), 7.03 (t, *J* = 1.7 Hz, 1H), 6.53 (d, *J* = 2.3 Hz, 1H), 6.44 (dd, *J* = 17.0, 10.1 Hz, 1H), 6.27 (dd, *J* = 17.0, 1.9 Hz, 1H), 6.16 (s, 1H), 5.77 (dd, *J* = 10.1, 2.0 Hz, 1H), 3.59 (td, *J* = 7.0, 2.2 Hz, 3H), 3.46 (s, 5H), 3.36 (t, *J* = 5.9 Hz, 2H), 3.15 (q, *J* = 5.8 Hz, 2H), 3.12 – 3.05 (m, 2H), 2.63 – 2.57 (m, 2H), 2.31 (t, *J* = 6.4 Hz, 2H), 2.05 (q, *J* = 7.7 Hz, 2H), 1.77 – 1.67 (m, 2H), 1.46 (p, *J* = 7.3 Hz, 2H), 1.37 – 1.26 (m, 3H), 1.26 – 1.11 (m, 3H), 0.94 (d, *J* = 6.4 Hz, 3H). $^{13}C$ NMR (126 MHz, DMSO) δ 172.19, 169.97, 163.32, 162.81, 150.33, 147.67, 146.21, 144.51, 143.84, 139.89, 136.69, 131.74, 127.20, 119.97, 117.19, 113.09, 95.41, 85.54, 69.48, 69.44, 69.15, 66.88, 54.97, 50.22, 38.41, 38.07, 36.20, 35.21, 32.53, 30.61, 29.50, 28.68, 25.57, 25.15, 15.48.

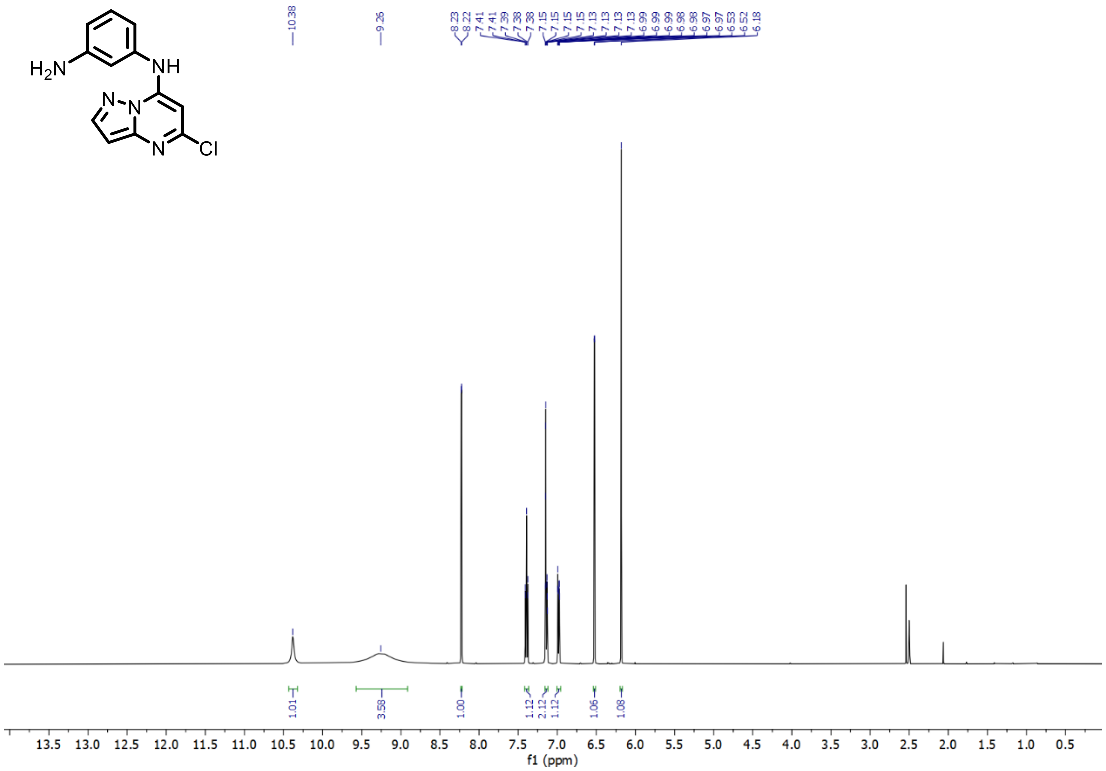

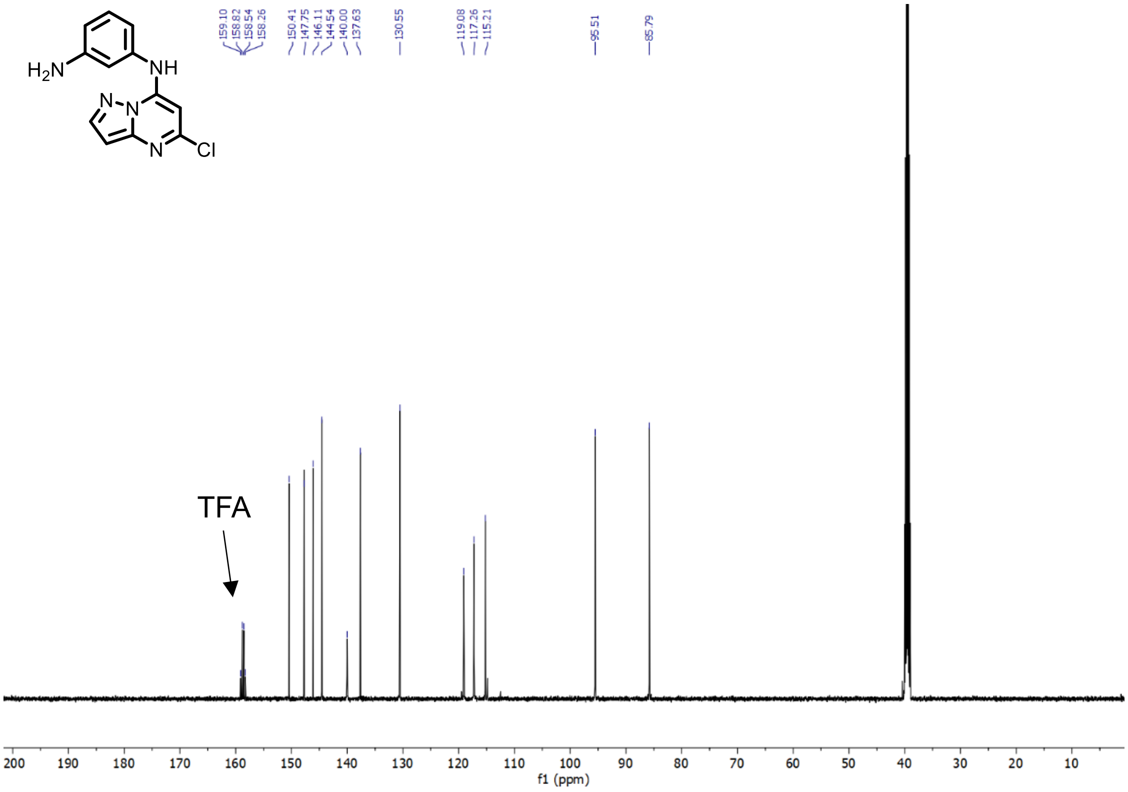

**Supplementary Figure 1.** $^{1}H$ and $^{13}C$ NMR of N1-(5-chloropyrazolo[1,5-a]pyrimidin-7-yl)benzene-1,3-diamine – SCA1 Intermediate 1 (TFA salt)

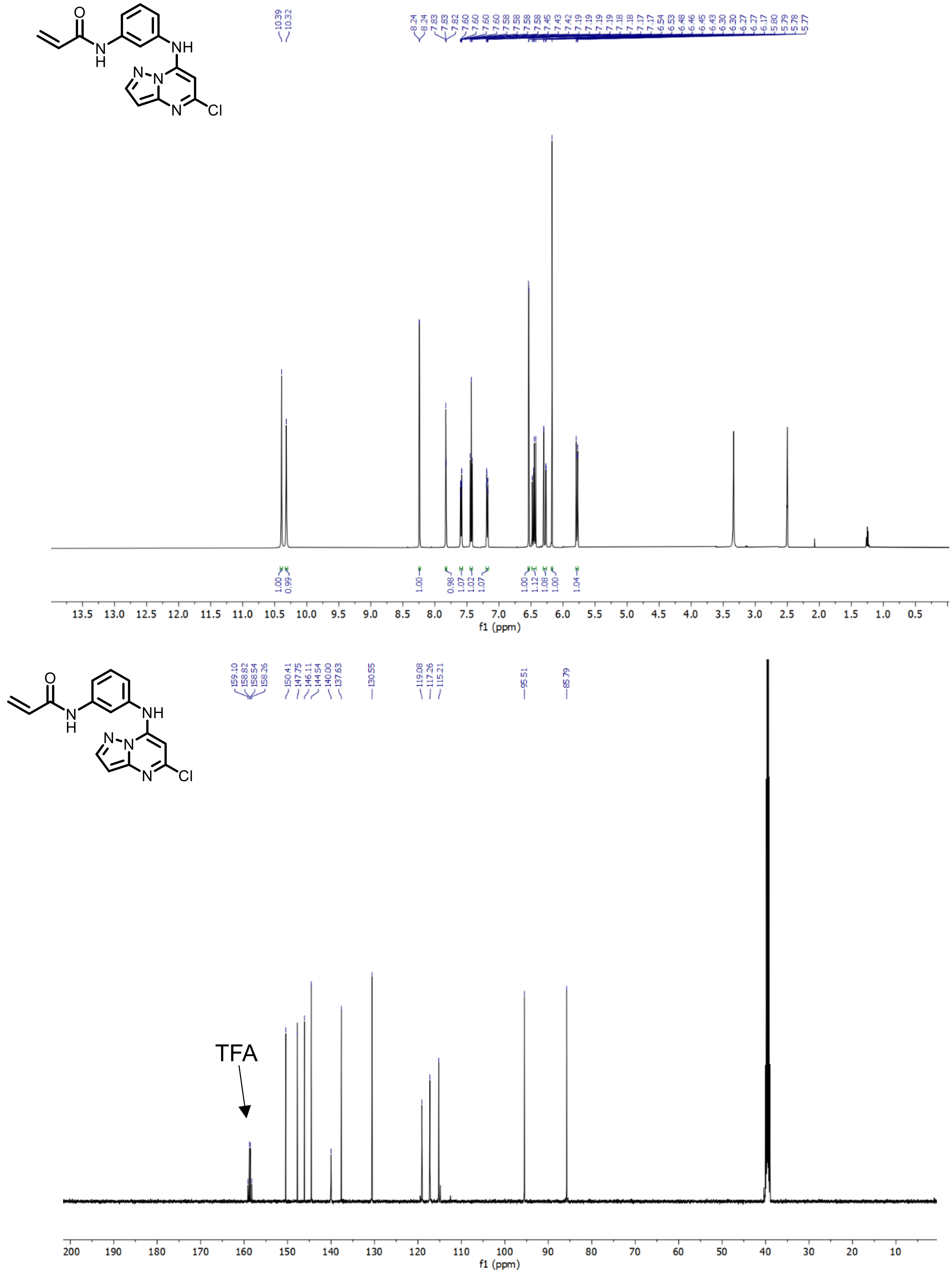

**Supplementary Figure 2.** $^{1}H$ and $^{13}C$ NMR of N-(3-((5-chloropyrazolo[1,5-a]pyrimidin-7-yl)amino)phenyl)propionamide – SCA1 (TFA salt)

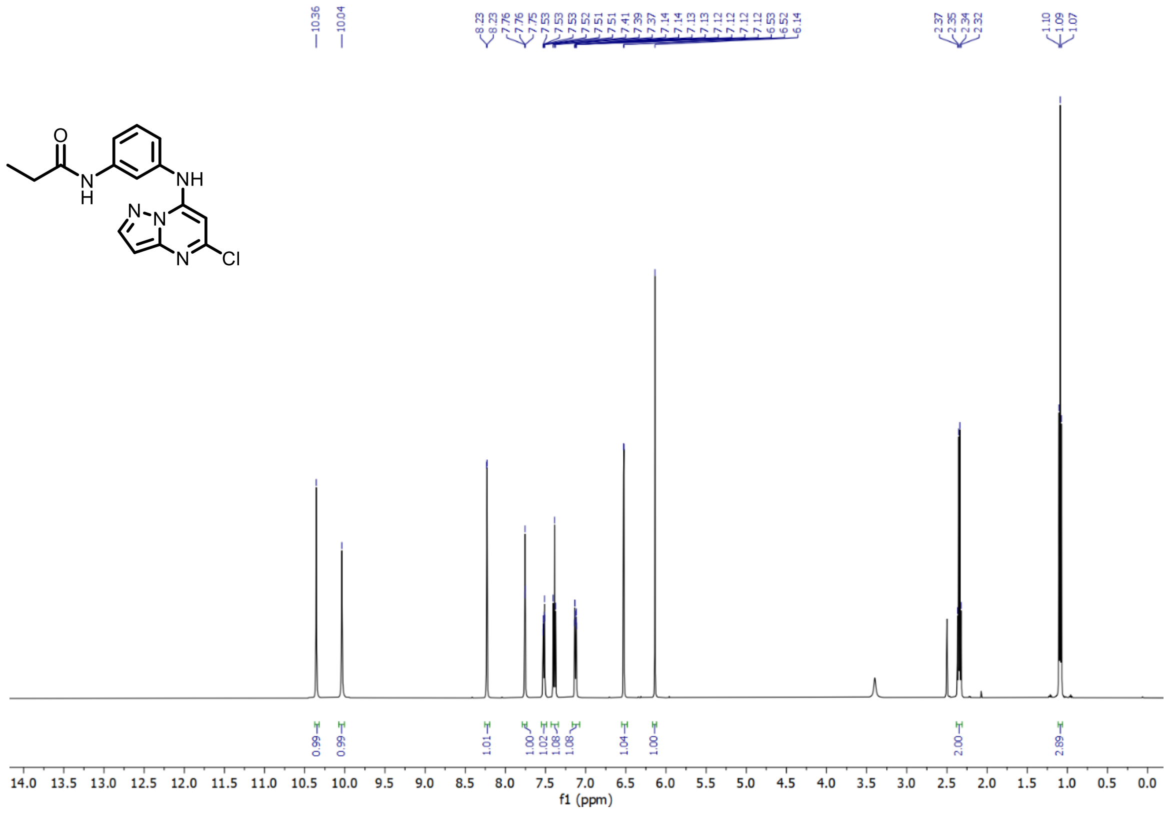

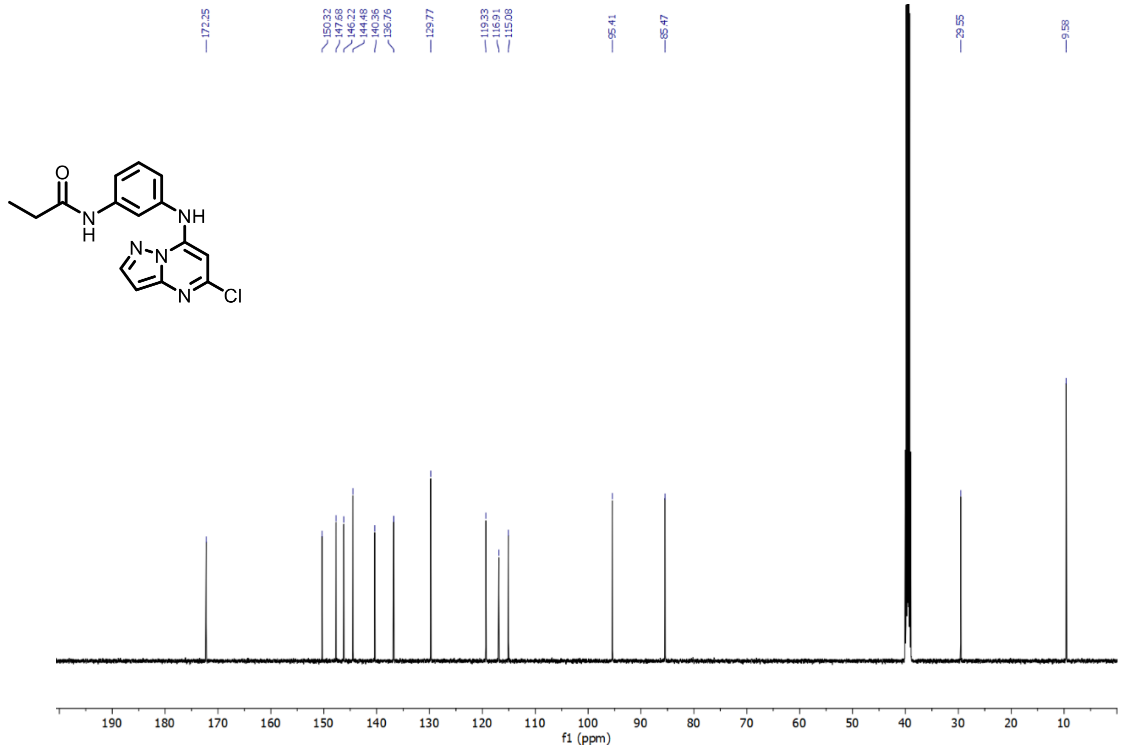

**Supplementary Figure 3.** $^{1}H$ and $^{13}C$ NMR of N-(3-((5-chloropyrazolo[1,5-a]pyrimidin-7-yl)amino)phenyl)propionamide – SCA1-NC

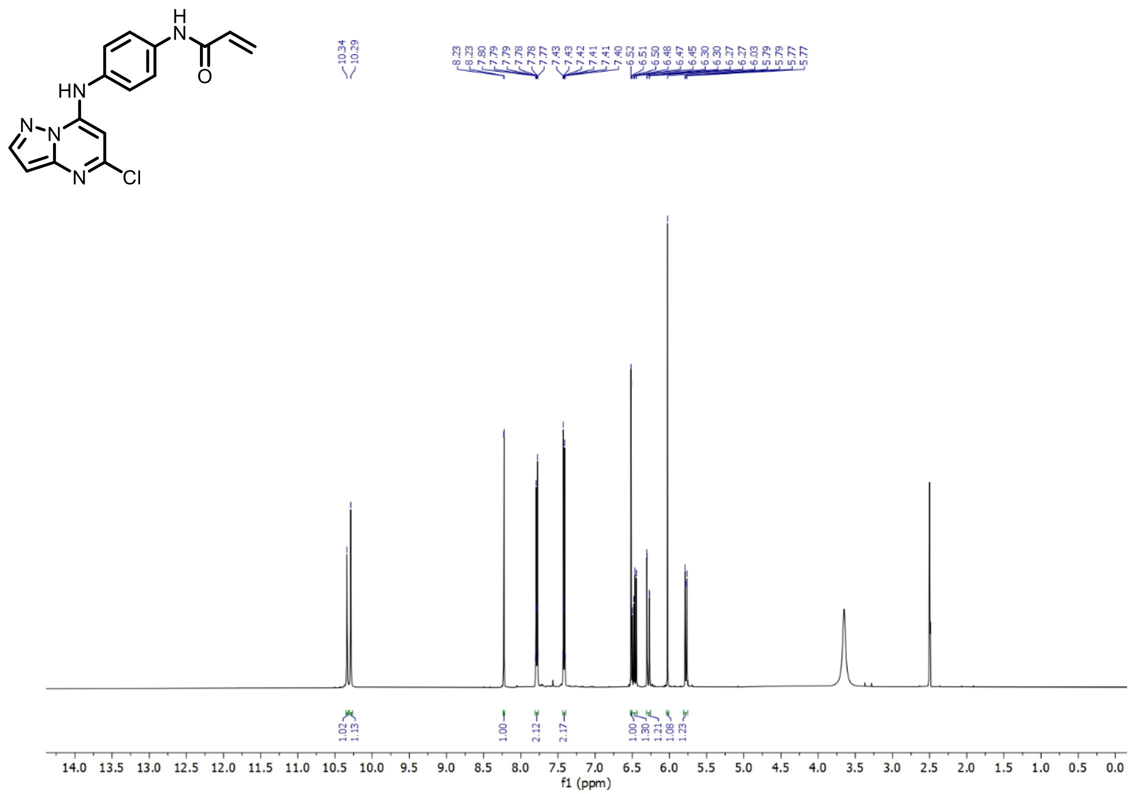

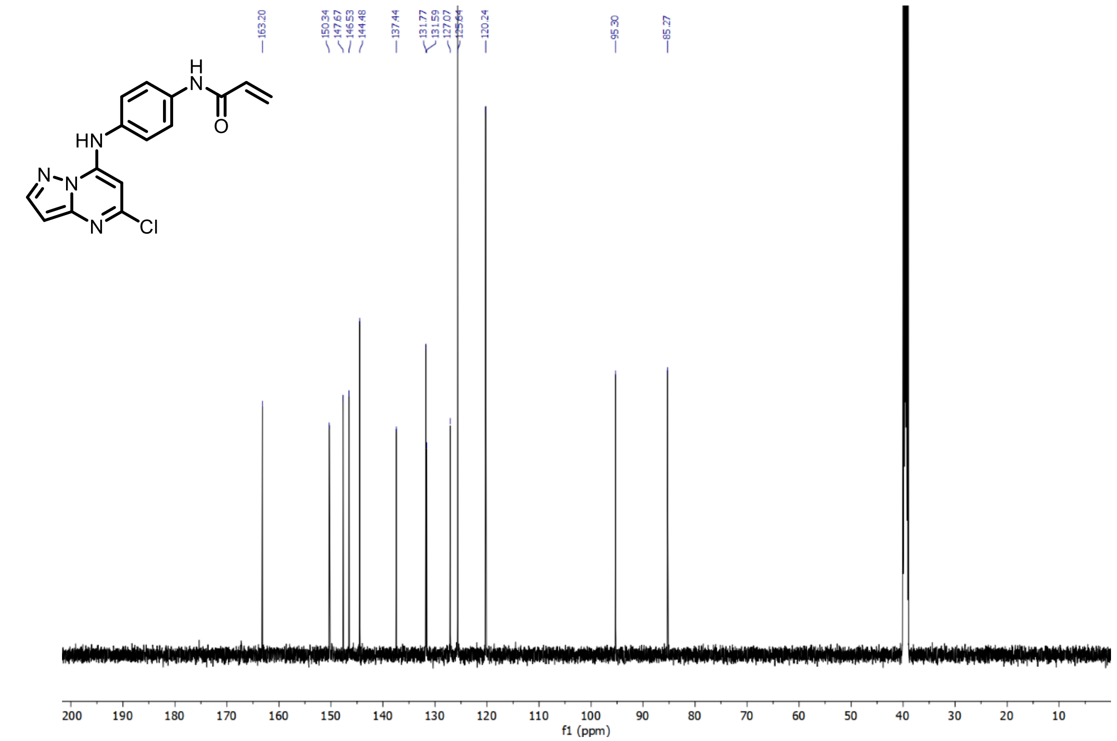

**Supplementary Figure 4.** $^{1}H$ and $^{13}C$ NMR of (N-(4-(5-chloropyrazolo[1,5-a]pyrimidin-7-ylamino)phenyl)acrylamide – SCA2

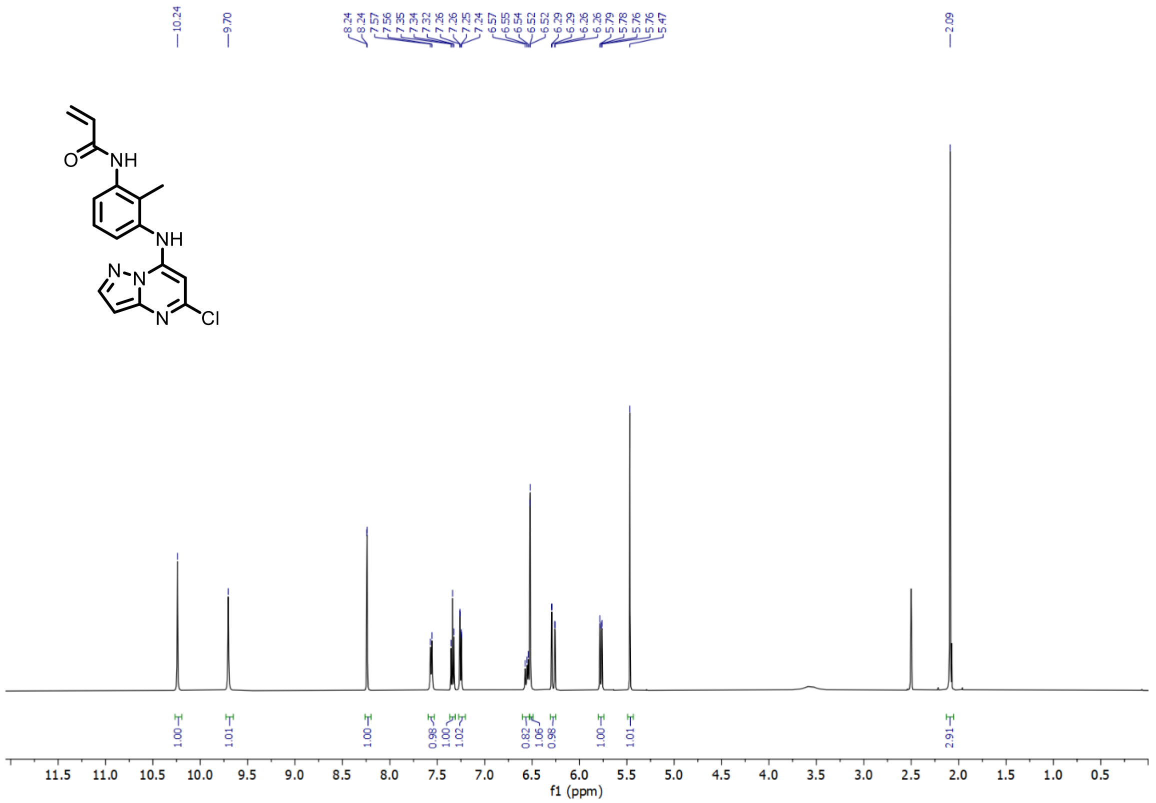

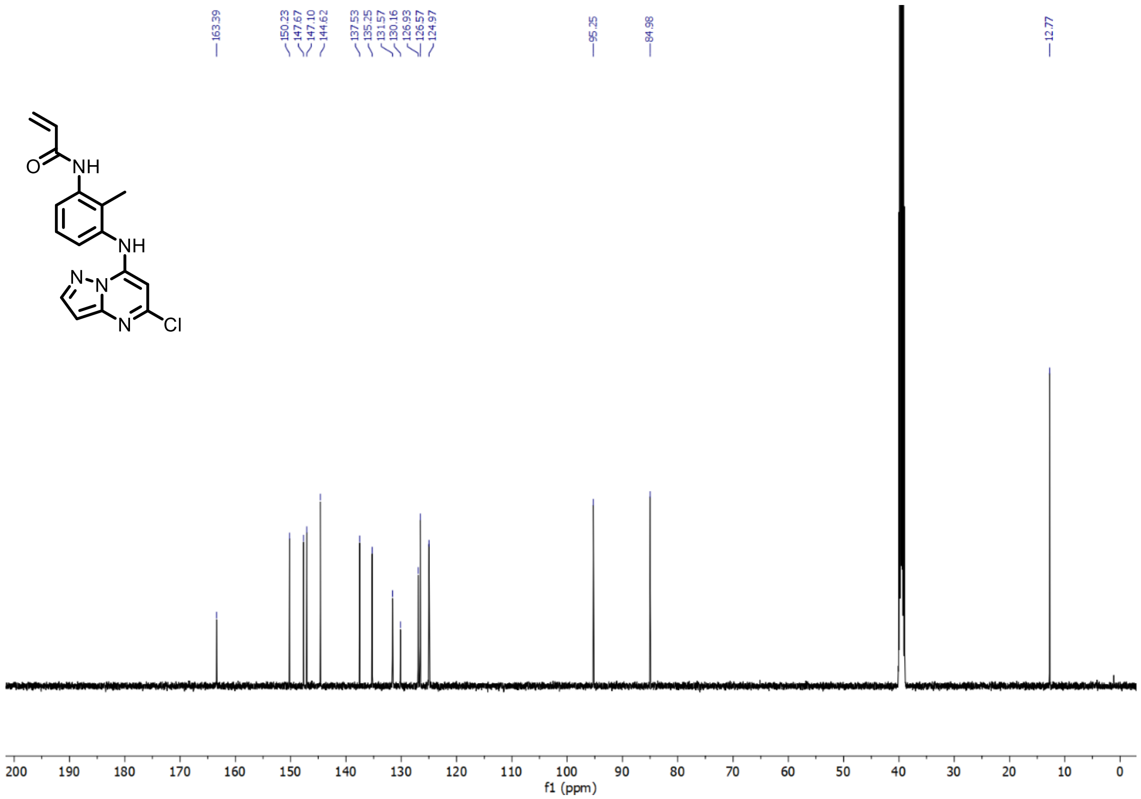

**Supplementary Figure 5.** $^{1}H$ and $^{13}C$ NMR of (N-(3-((5-chloropyrazolo[1,5-a]pyrimidin-7-yl)amino)-2-methylphenyl)acrylamide – SCA3

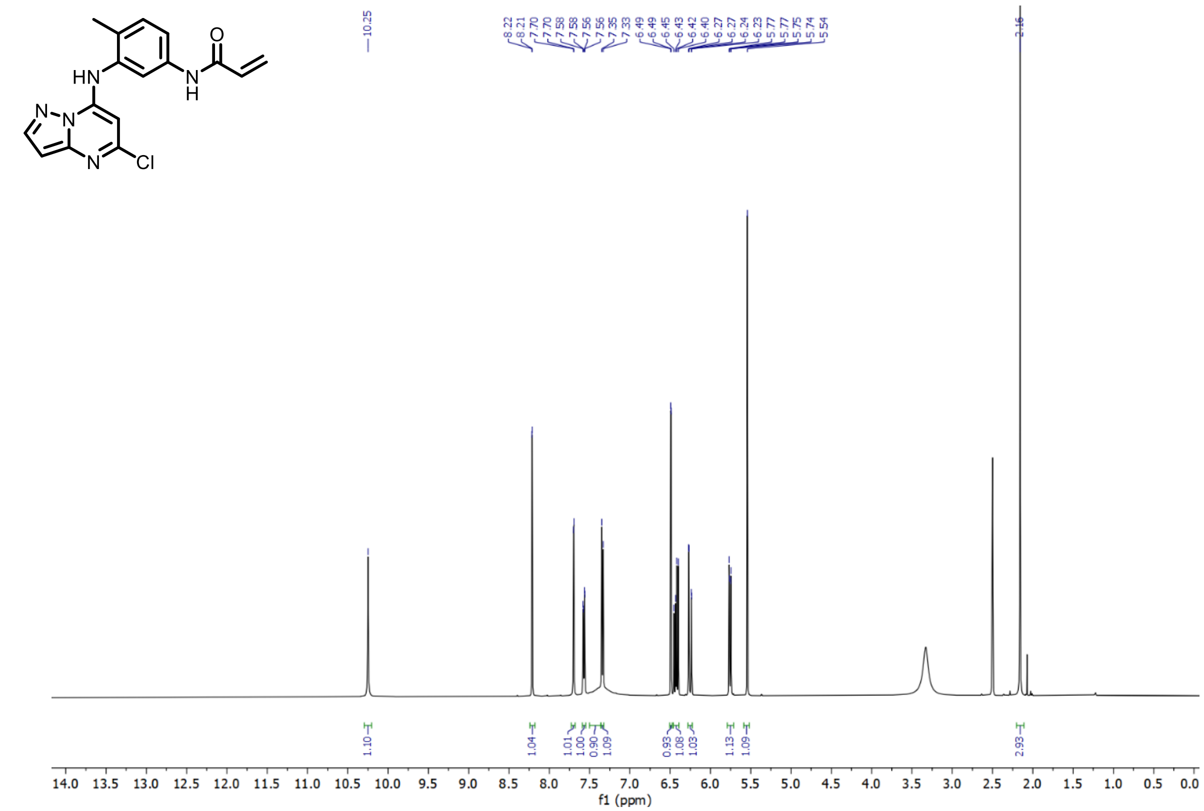

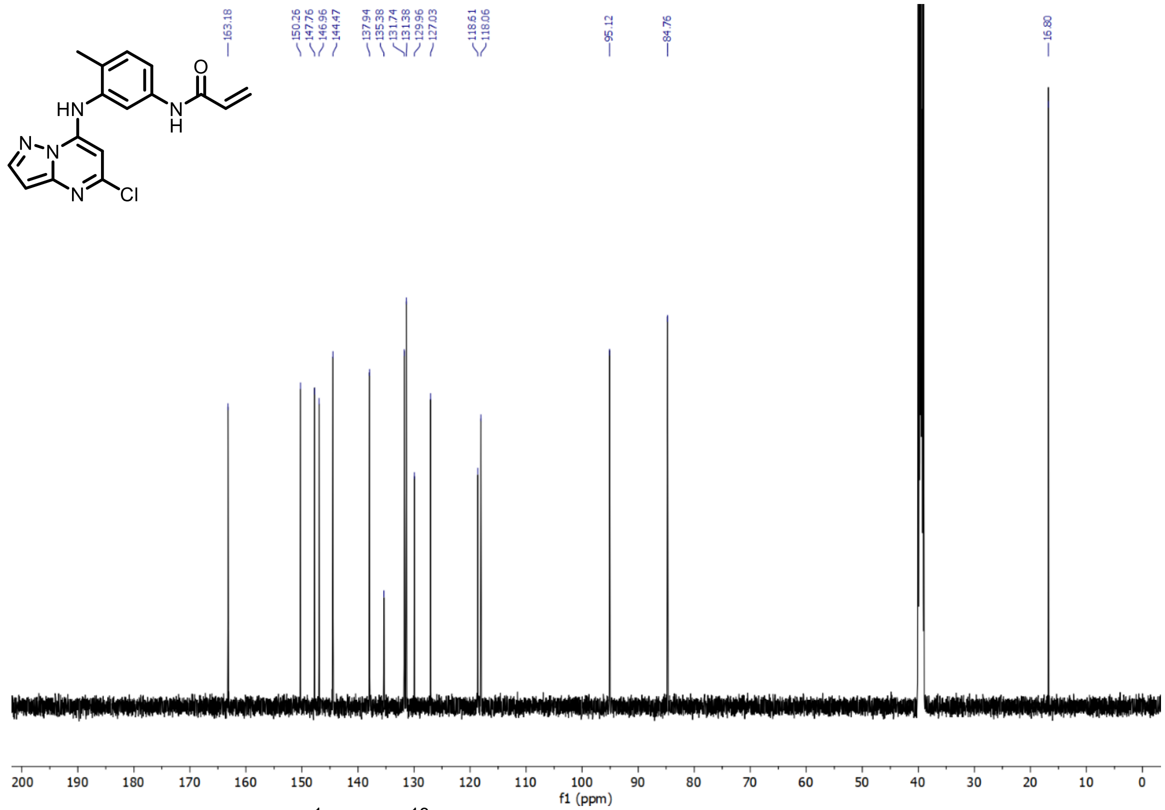

**Supplementary Figure 6**. $^{1}$H and $^{13}$C NMR N-(3-(5-chloropyrazolo[1,5-a]pyrimidin-7-ylamino)-4-methylphenyl)acrylamide – SCA4

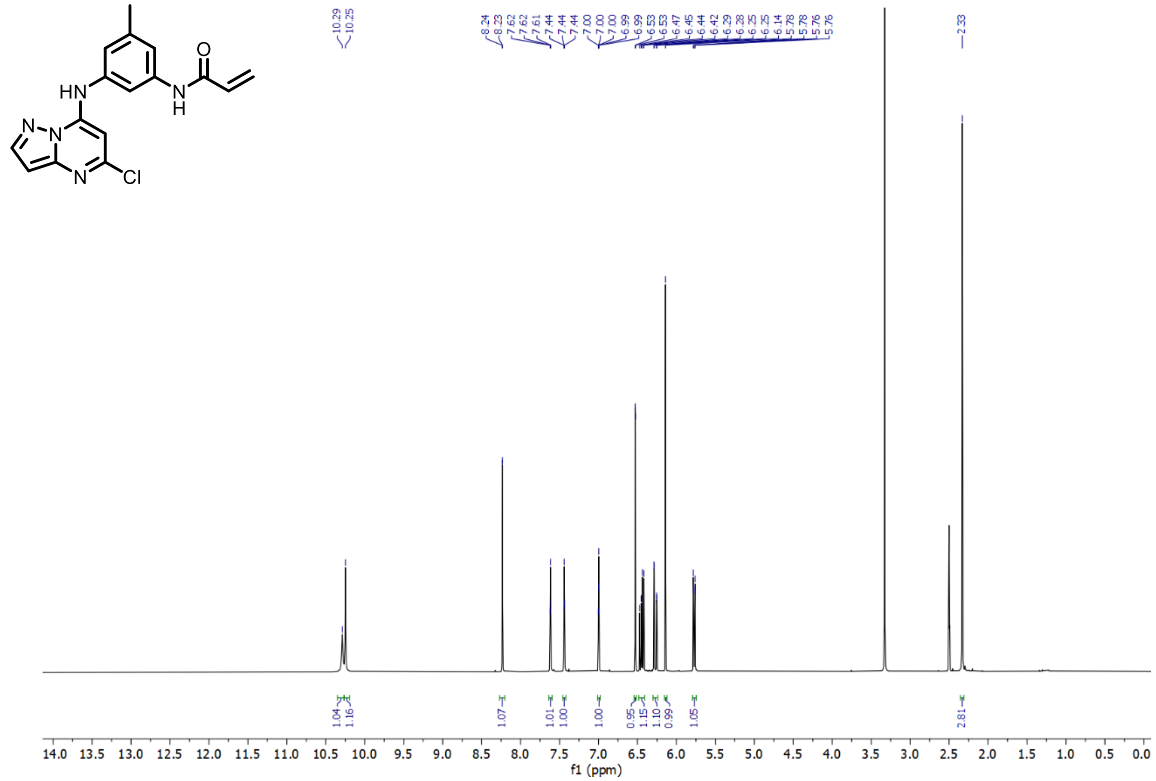

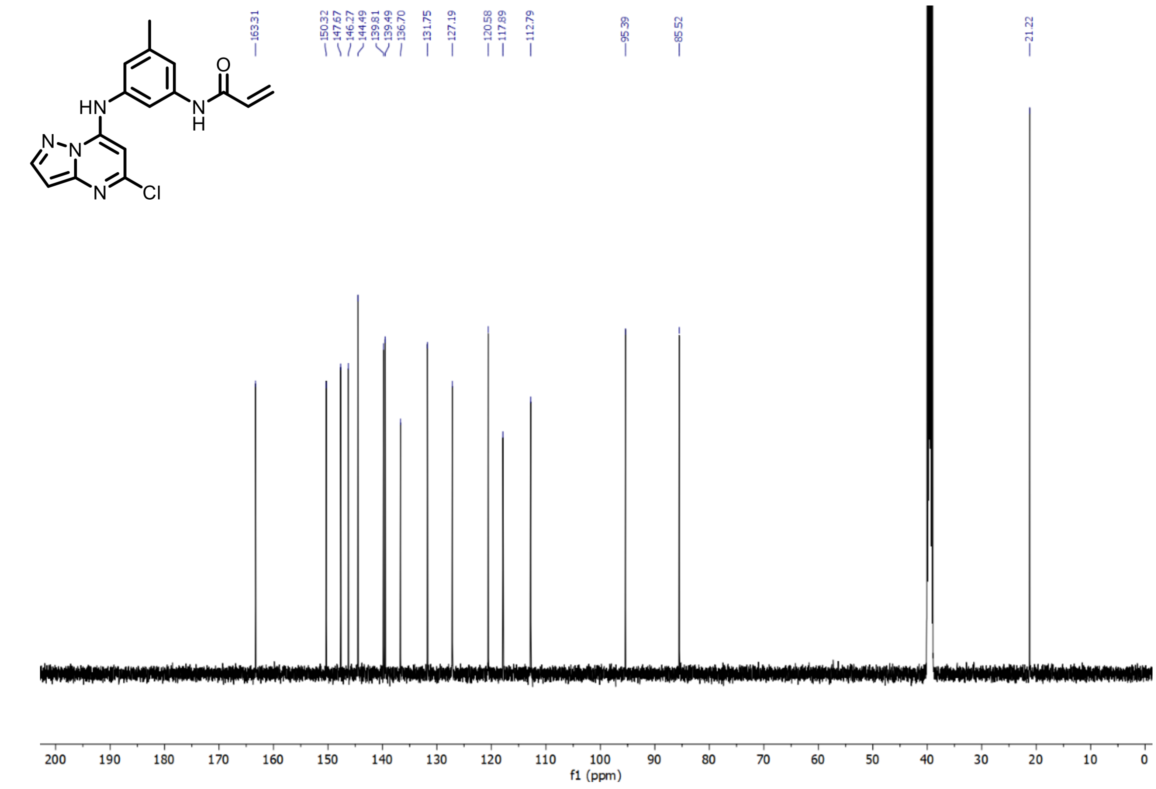

**Supplementary Figure 7.** $^{1}H$ and $^{13}C$ NMR N-(3-(5-chloropyrazolo[1,5-a]pyrimidin-7-ylamino)-5-methylphenyl)acrylamide – SCA5

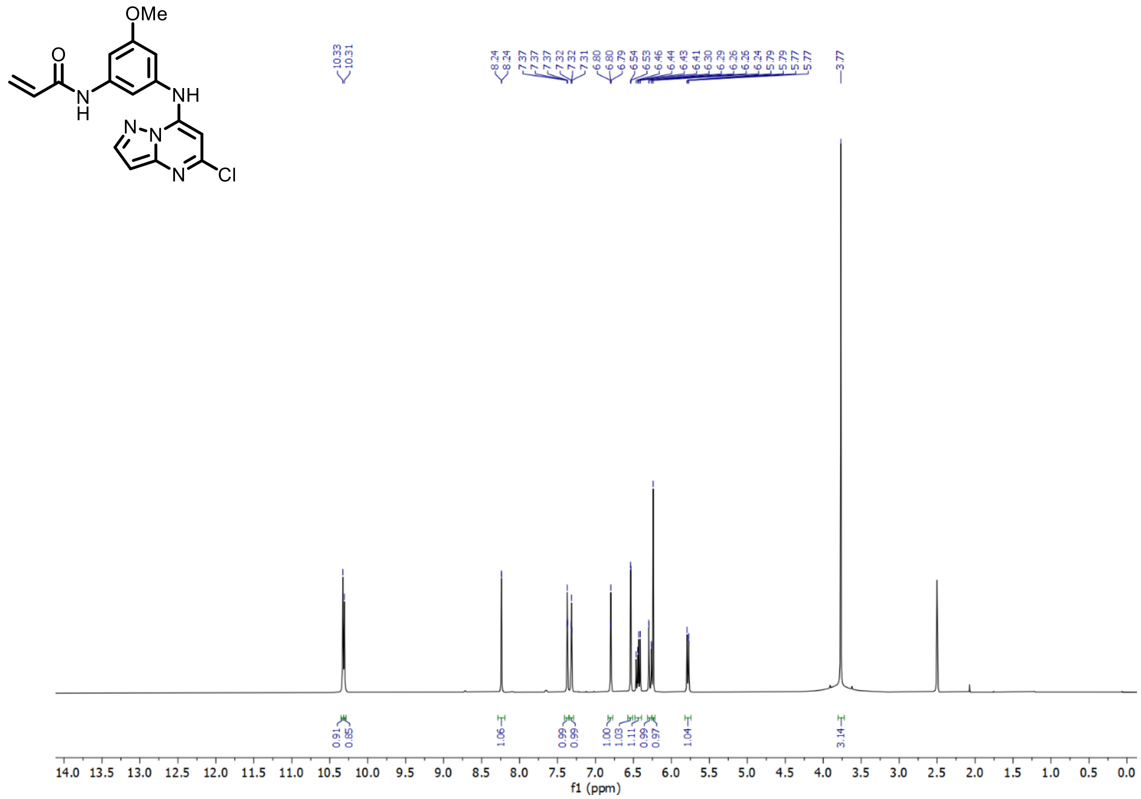

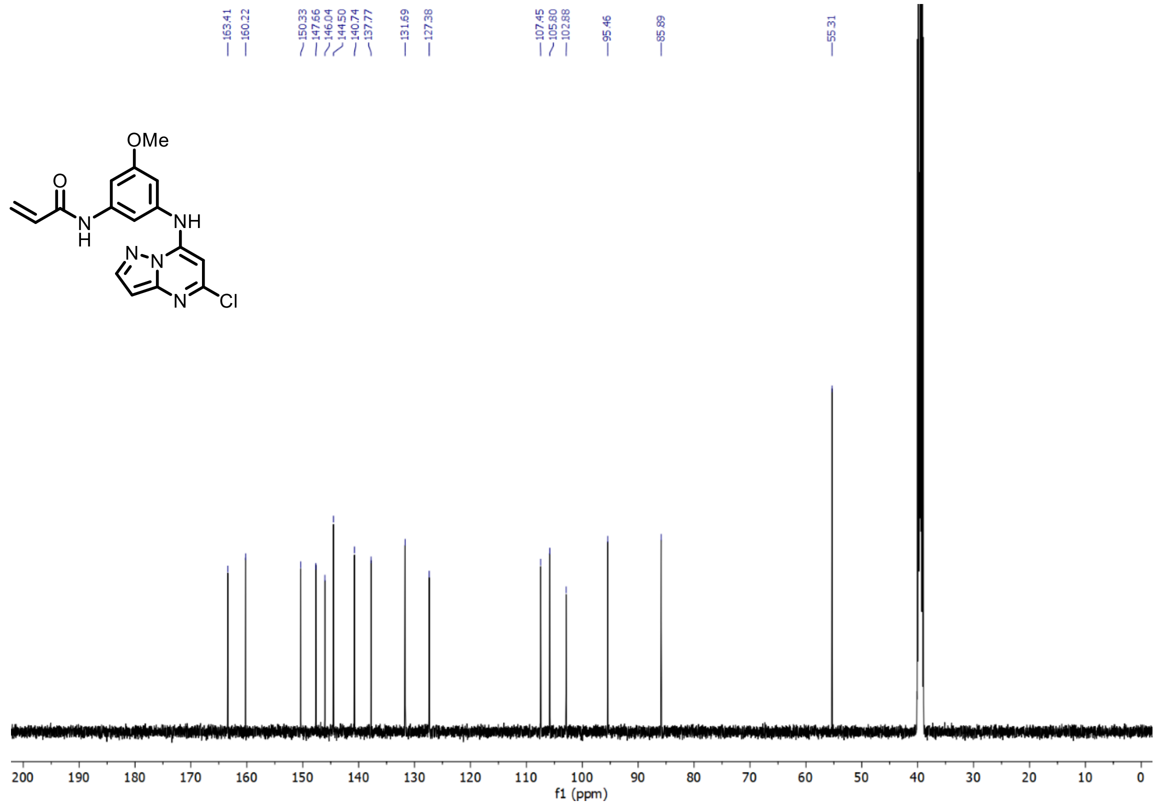

**Supplementary Figure 8.** $^{1}H$ and $^{13}C$ NMR of N-(3-((5-chloropyrazolo[1,5-a]pyrimidin-7-yl)amino)-5-methoxyphenyl)acrylamide – SCA6

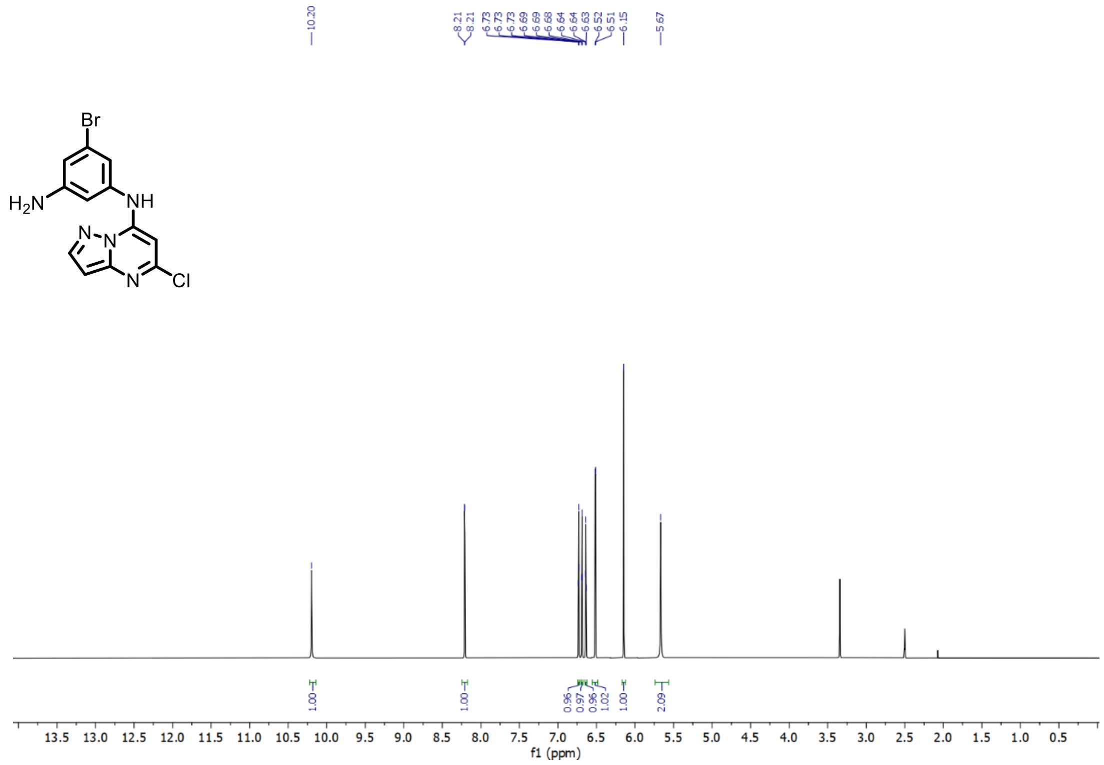

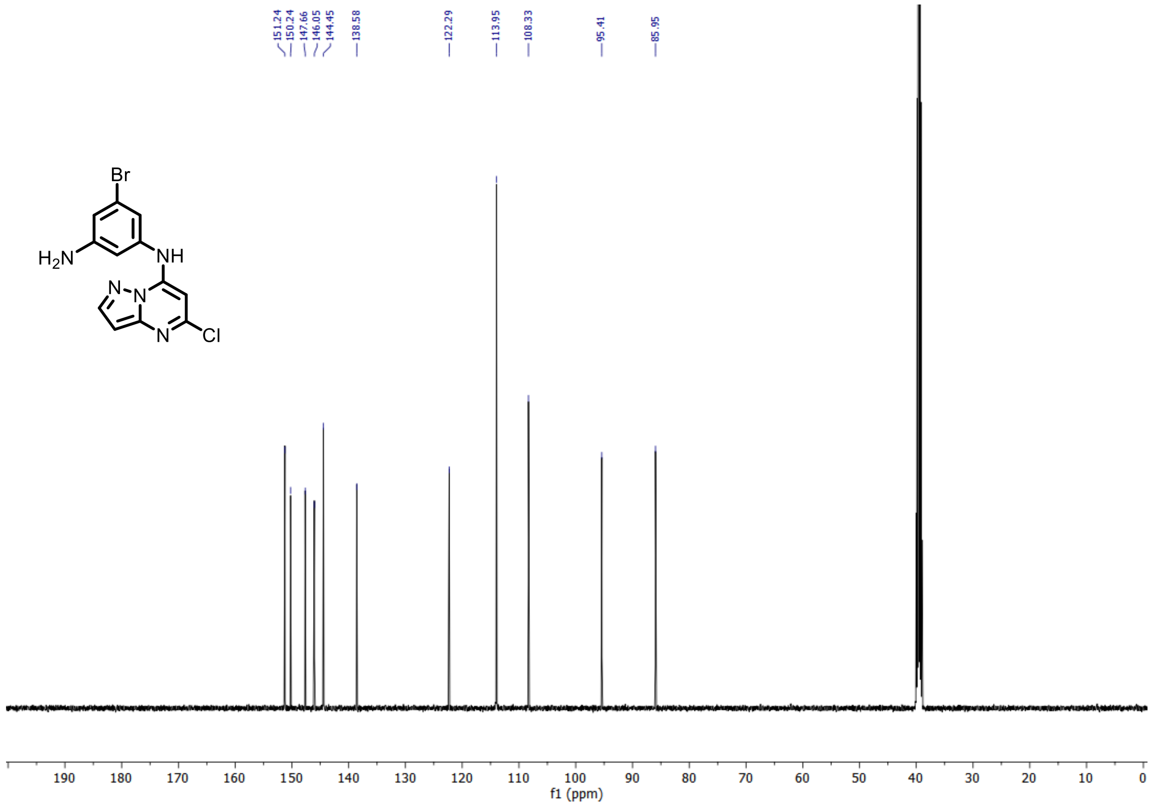

**Supplementary Figure 9.** $^{1}H$ and $^{13}C$ NMR of 5-bromo-N1-(5-chloropyrazolo[1,5-a]pyrimidin-7-yl)benzene-1,3-diamine – SCA7 Intermediate

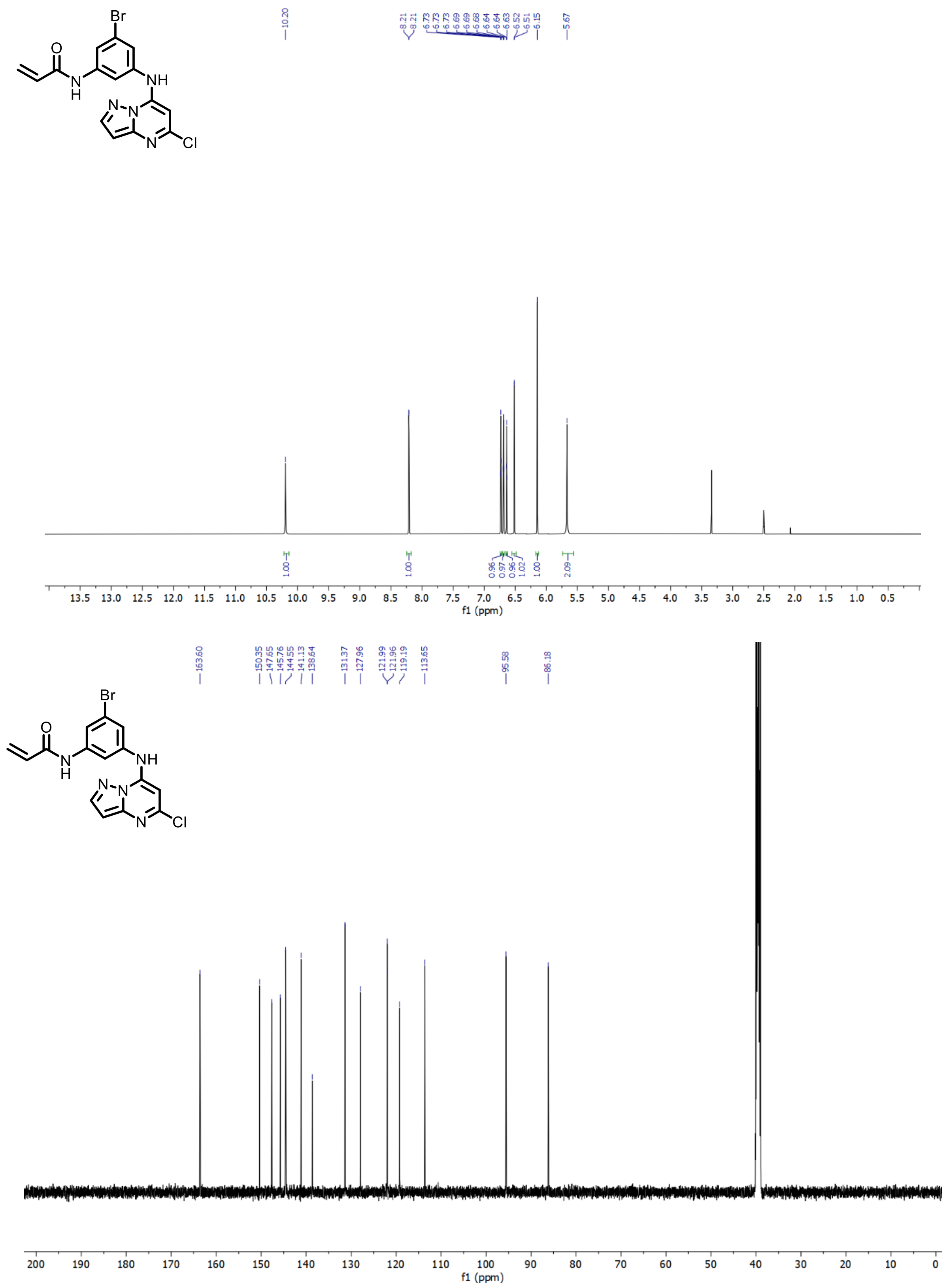

**Supplementary Figure 10.** $^{1}$H and $^{13}$C NMR of N-(3-bromo-5-((5-chloropyrazolo[1,5-a]pyrimidin-7-yl)amino)phenyl)acrylamide – SCA7 (HCl Salt)

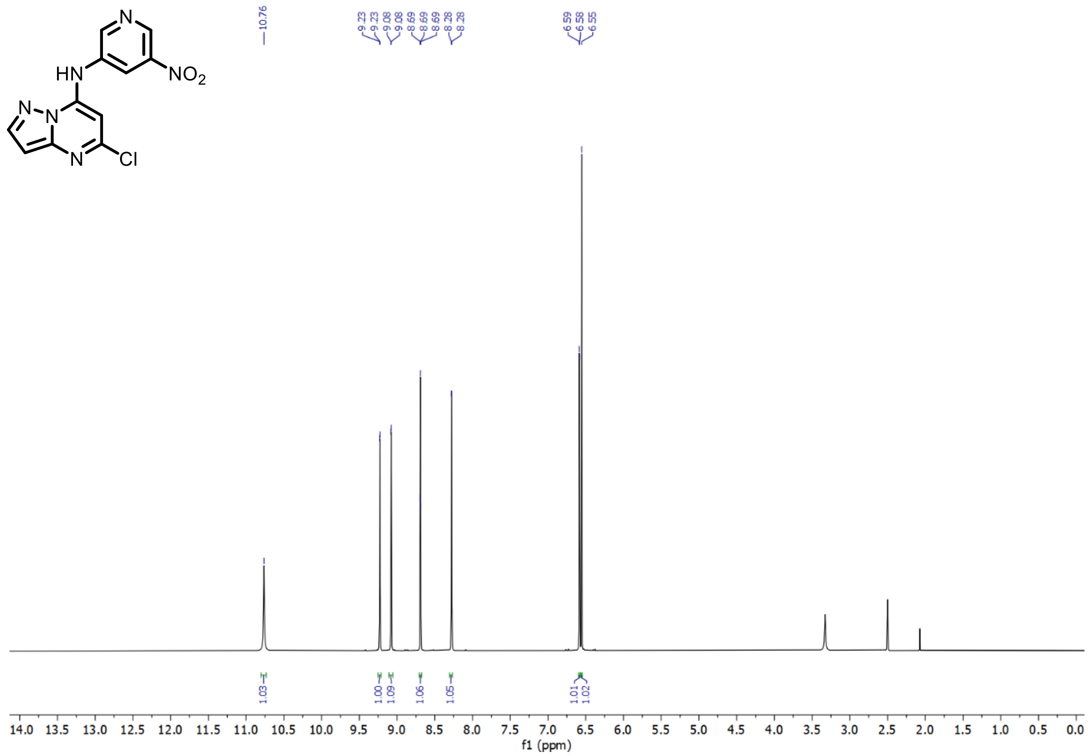

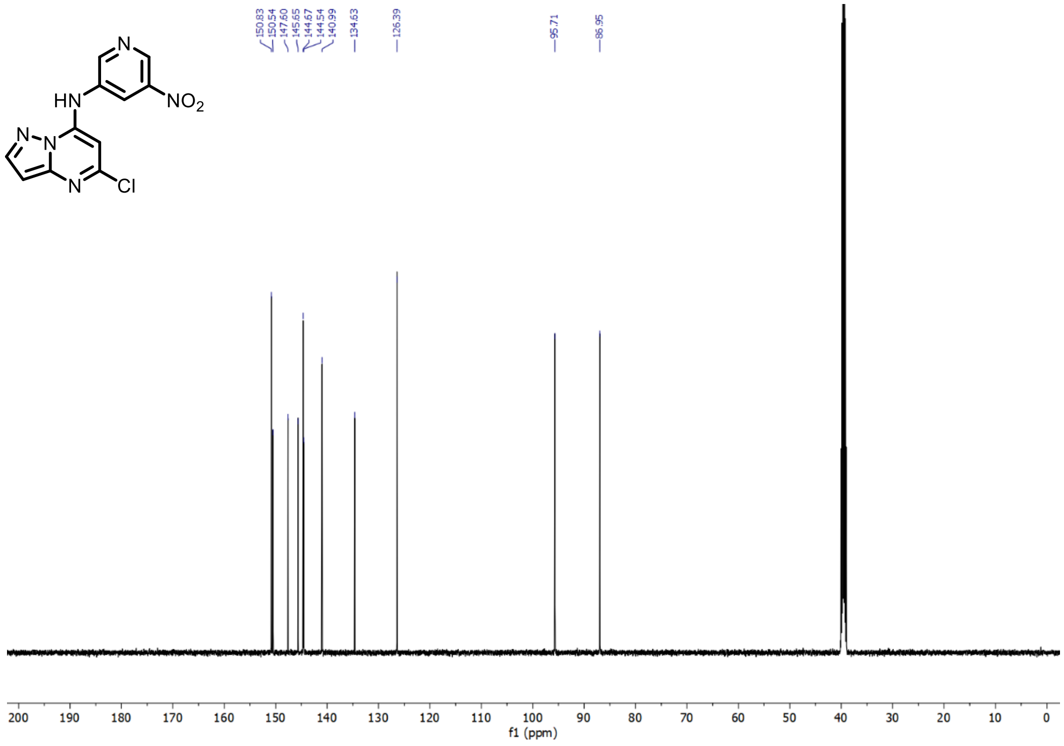

**Supplementary Figure 11.** $^{1}$H and $^{13}$C NMR 5-chloro-N-(5-nitro-3-pyridyl)pyrazolo[1,5-a]pyrimidin-7-amine – SCA8 Intermediate 1

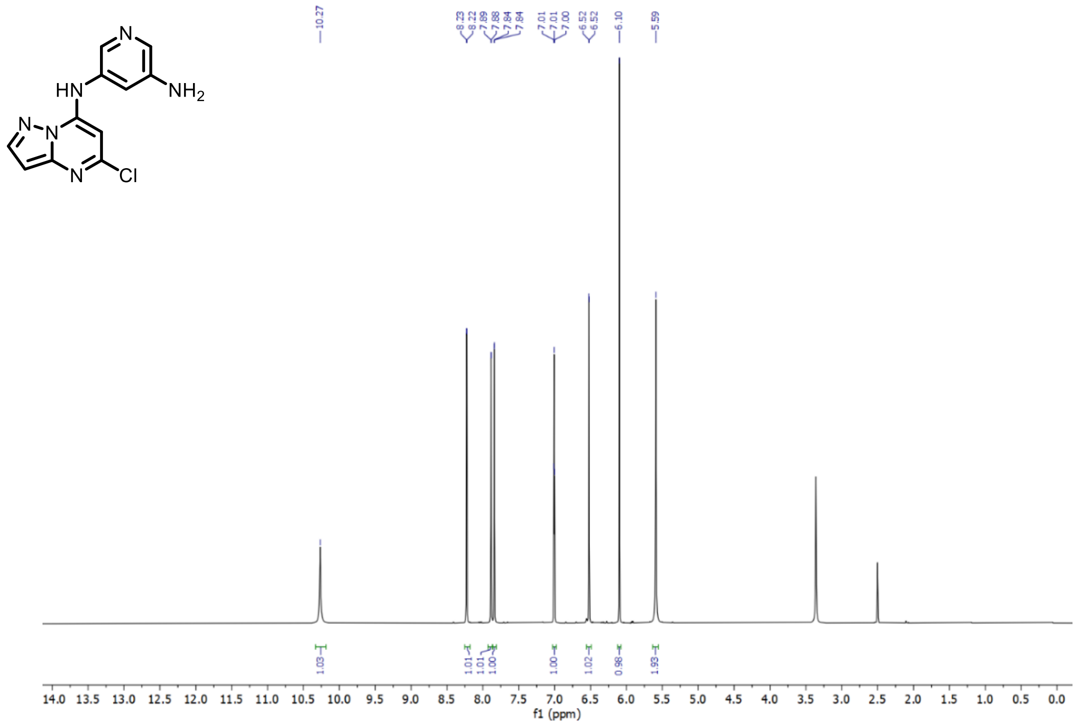

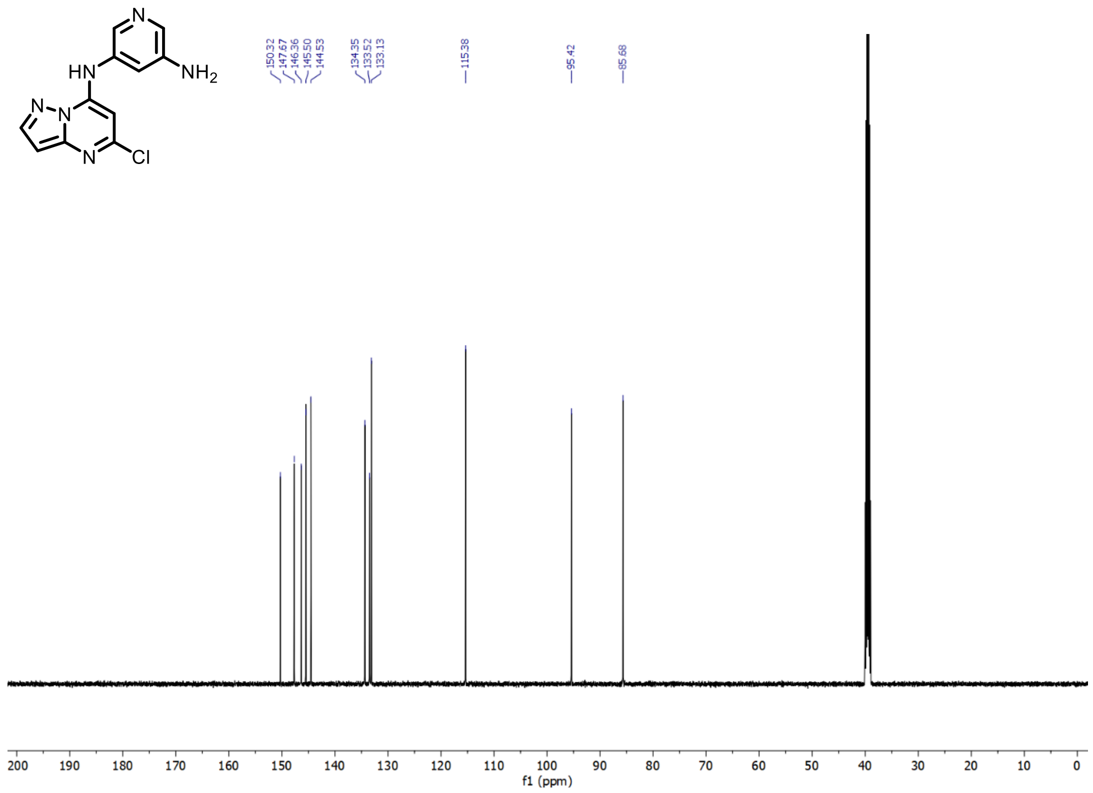

**Supplementary Figure 12.** $^{1}H$ and $^{13}C$ NMR of N5-(5-chloropyrazolo[1,5-a]pyrimidin-7-yl)pyridine-3,5-diamine – SCA8 Intermediate 2

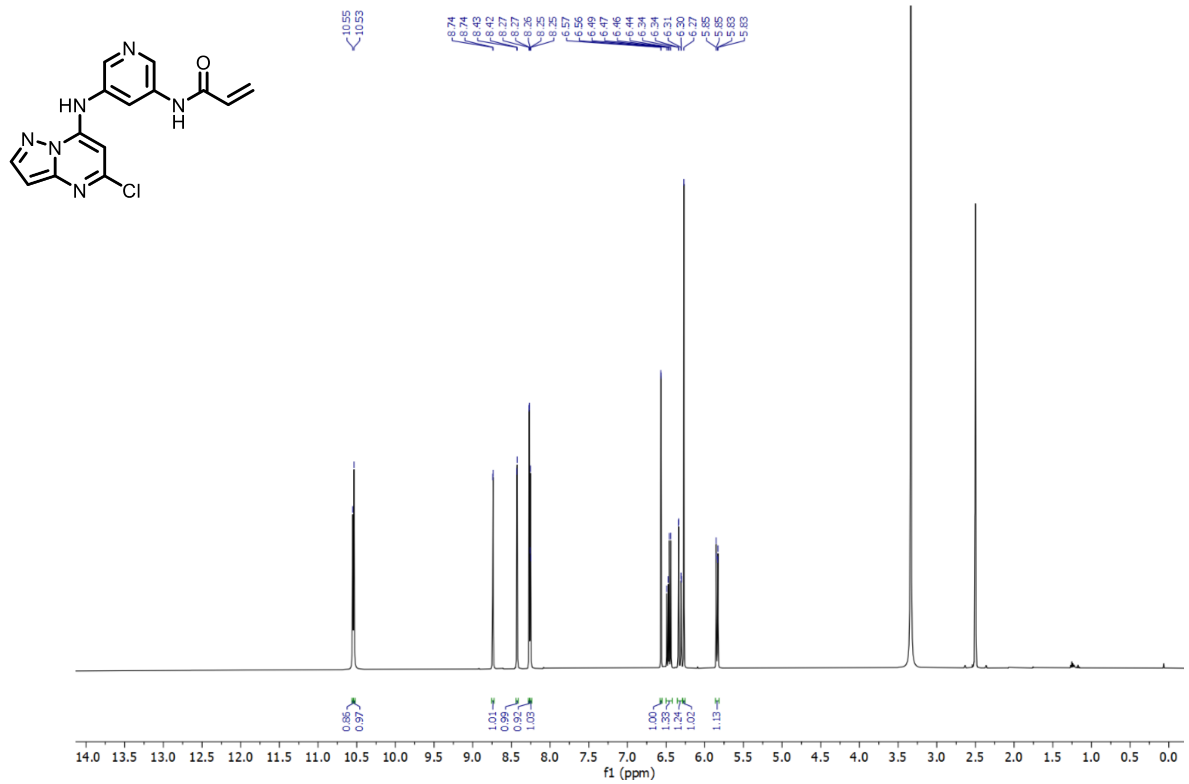

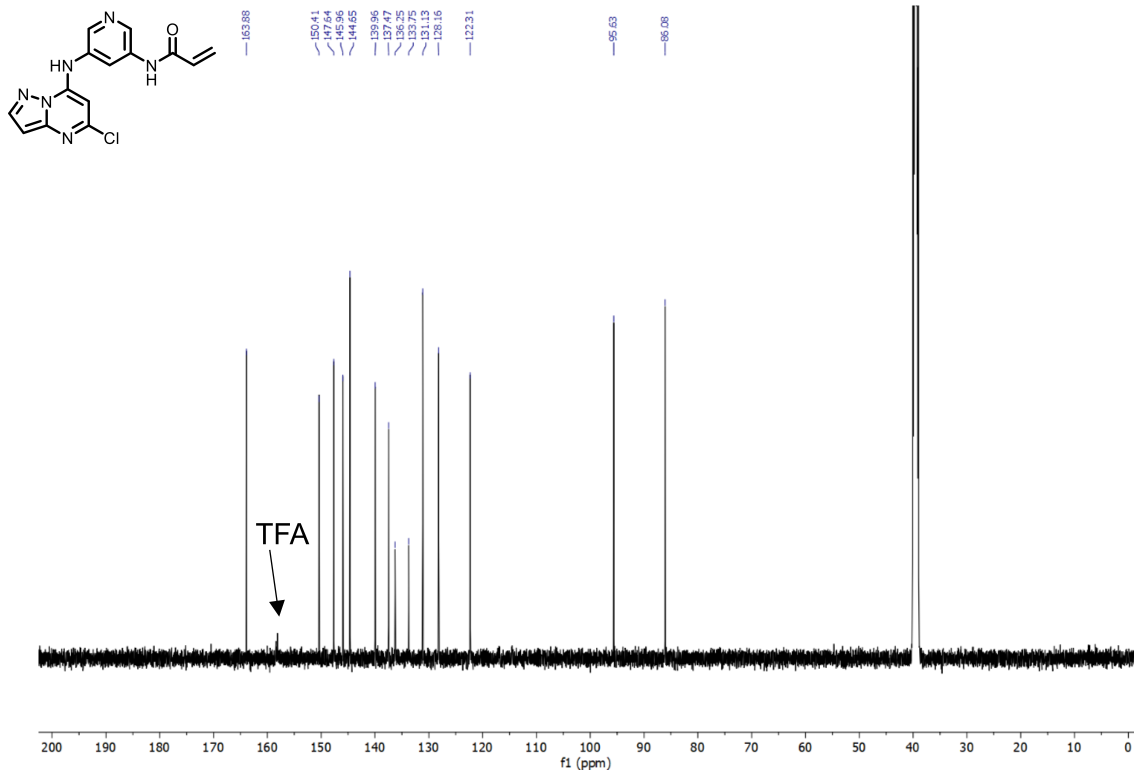

**Supplementary Figure 13.** $^{1}H$ NMR (HCl salt) and $^{13}C$ NMR (TFA salt) of N-(5-((5-chloropyrazolo[1,5-a]pyrimidin-7-yl)amino)pyridin-3-yl)acrylamide – SCA8

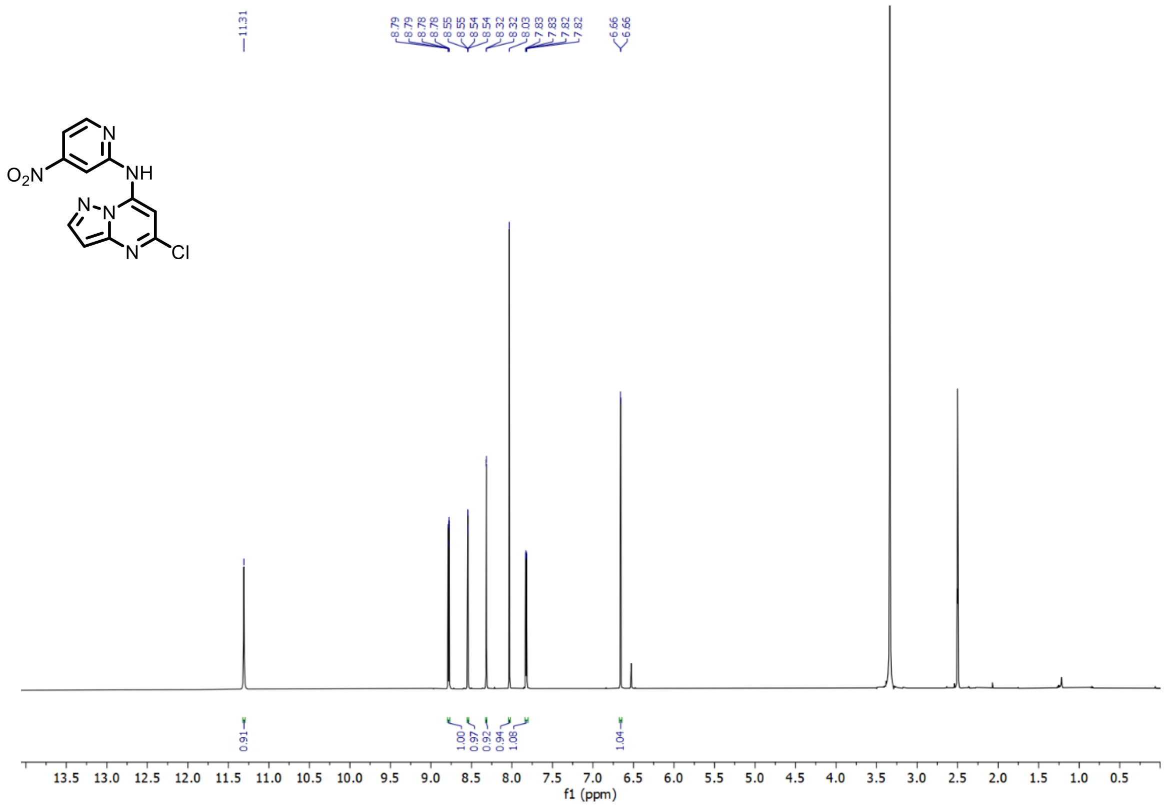

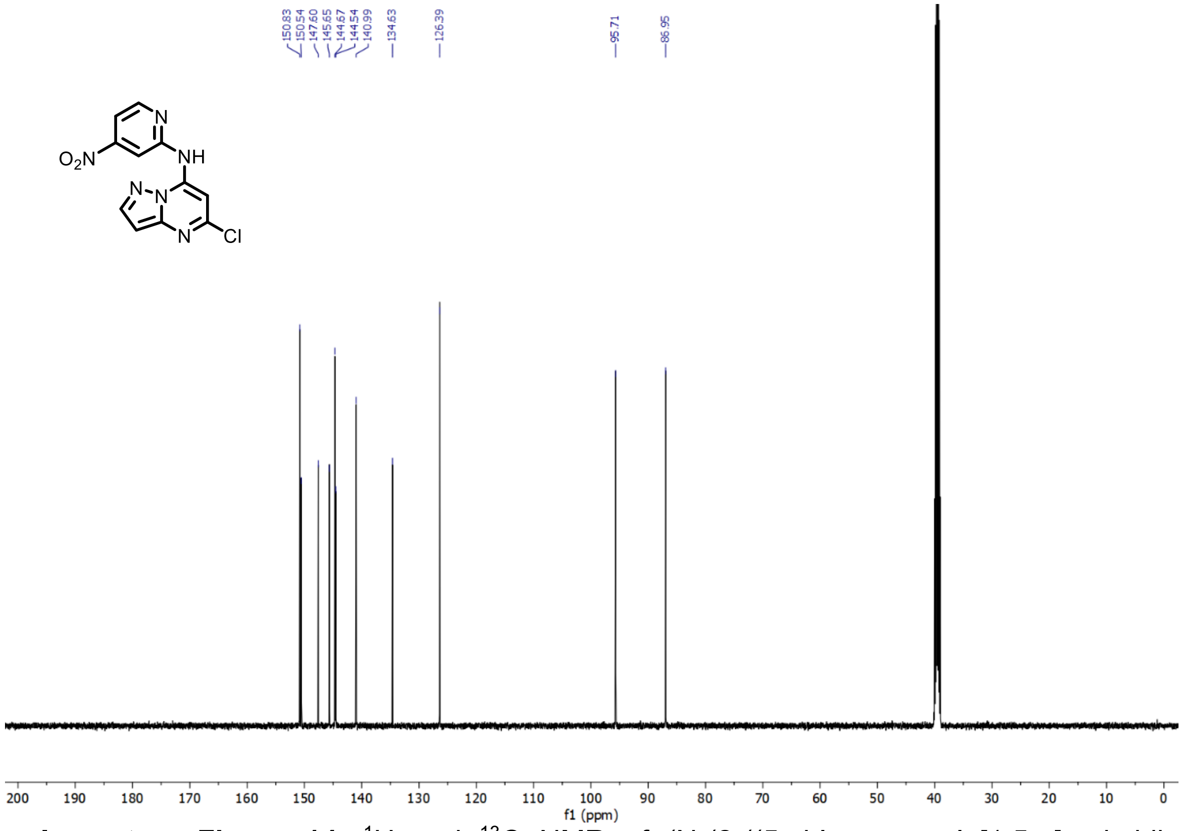

**Supplementary Figure 14.** $^{1}H$ and $^{13}C$ NMR of (N-(2-((5-chloropyrazolo[1,5-a]pyrimidin-7-yl)amino)pyridin-4-yl)acrylamide) – SCA9 Intermediate 1

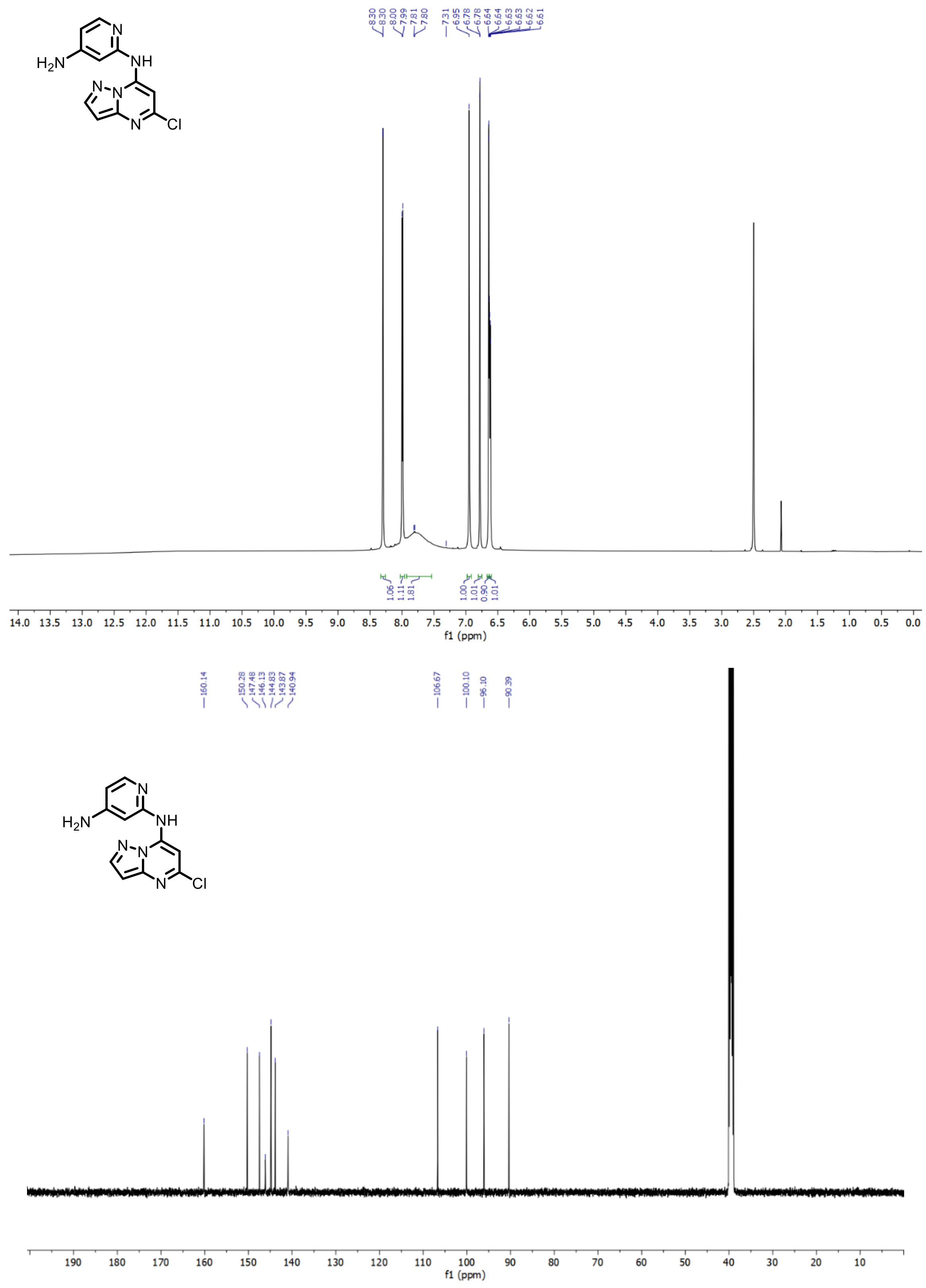

**Supplementary Figure 15.** $^{1}H$ and $^{13}C$ NMR of N2-(5-chloropyrazolo[1,5-a]pyrimidin-7-yl)pyridine-2,4-diamine – SCA9 Intermediate 2

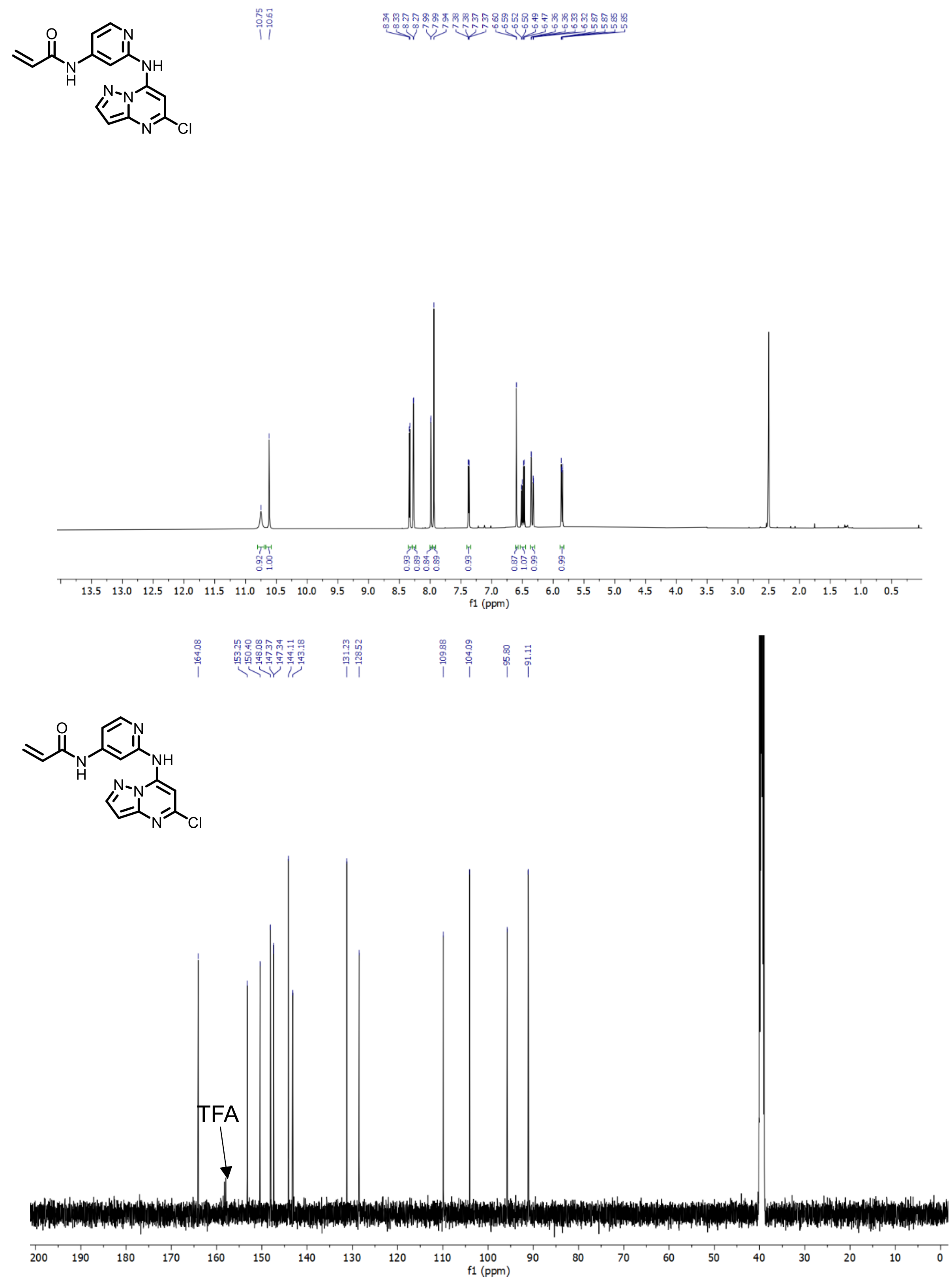

**Supplementary Figure 16.** $^{1}H$ and $^{13}C$ NMR of N-(2-((5-chloropyrazolo[1,5-a]pyrimidin-7-yl)amino)pyridin-4-yl)acrylamide – SCA9 (TFA salt)

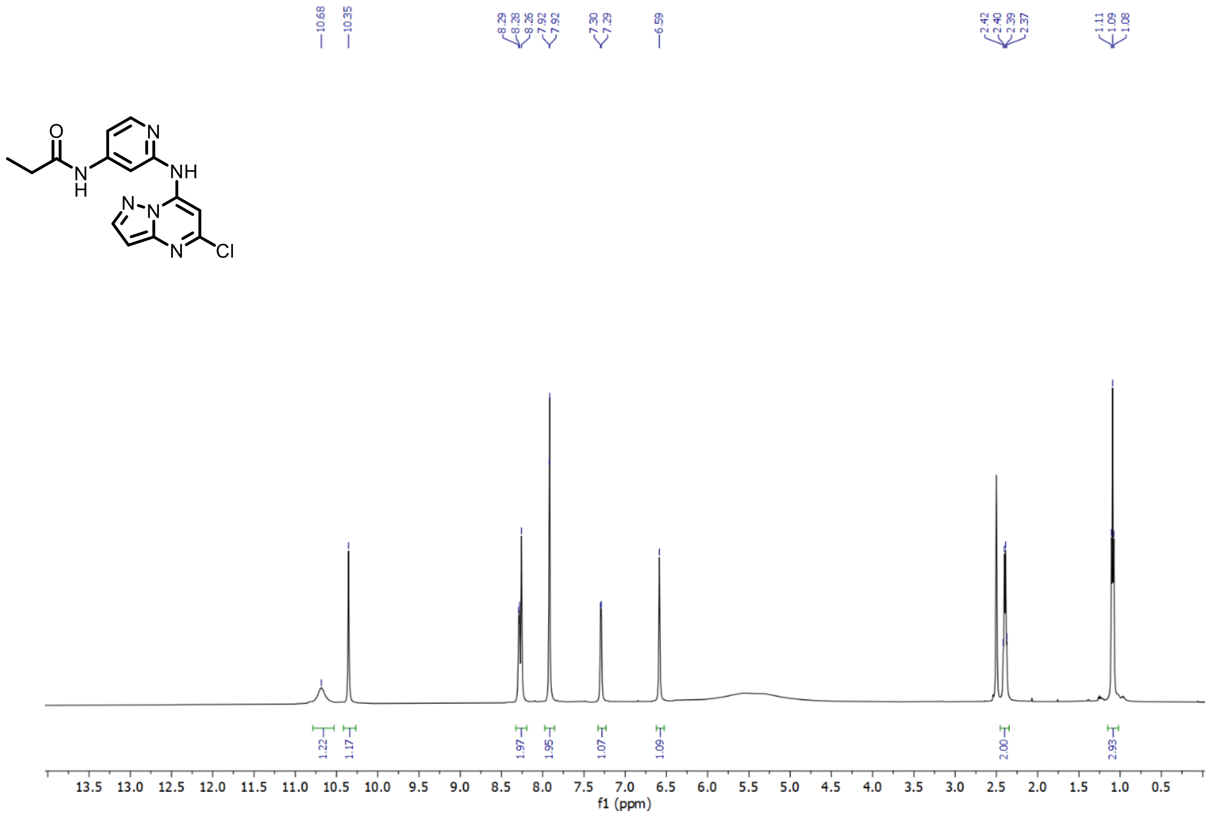

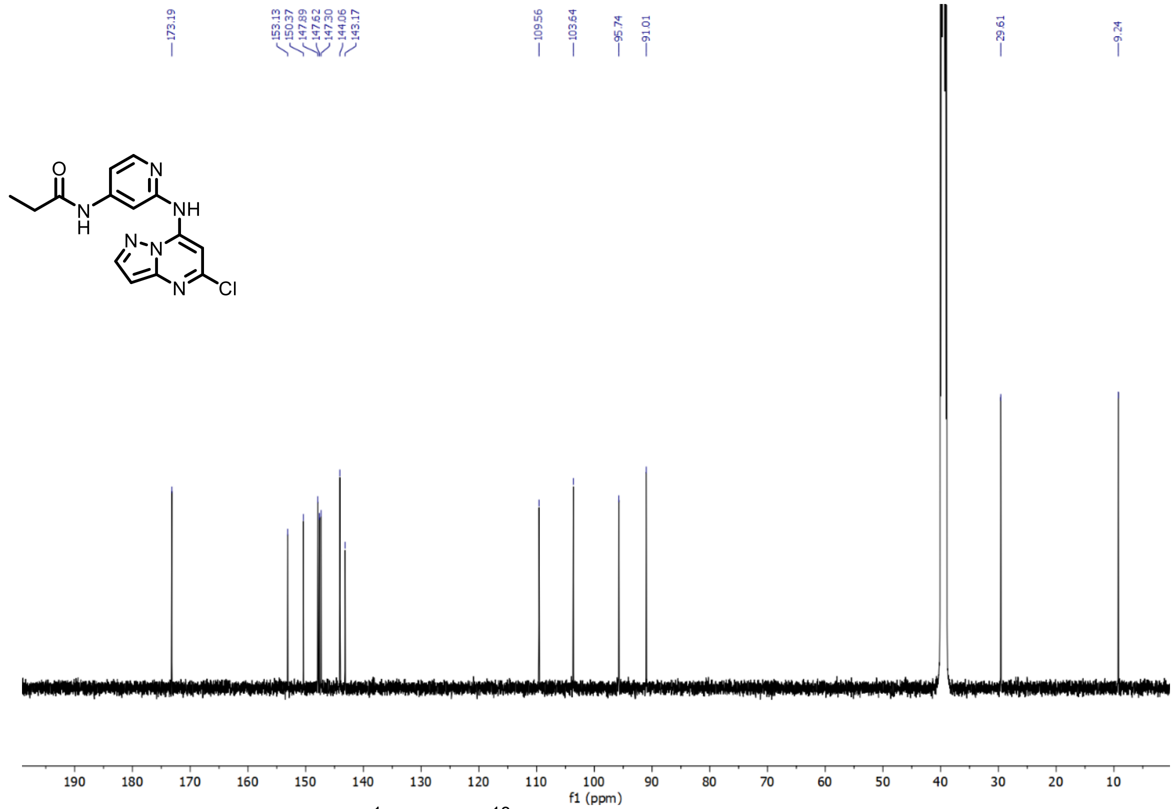

**Supplementary Figure 17.** $^{1}H$ and $^{13}C$ NMR of N-(2-((5-chloropyrazolo[1,5-a]pyrimidin-7-yl)amino)pyridin-4-yl) propionamide – SCA9-NC

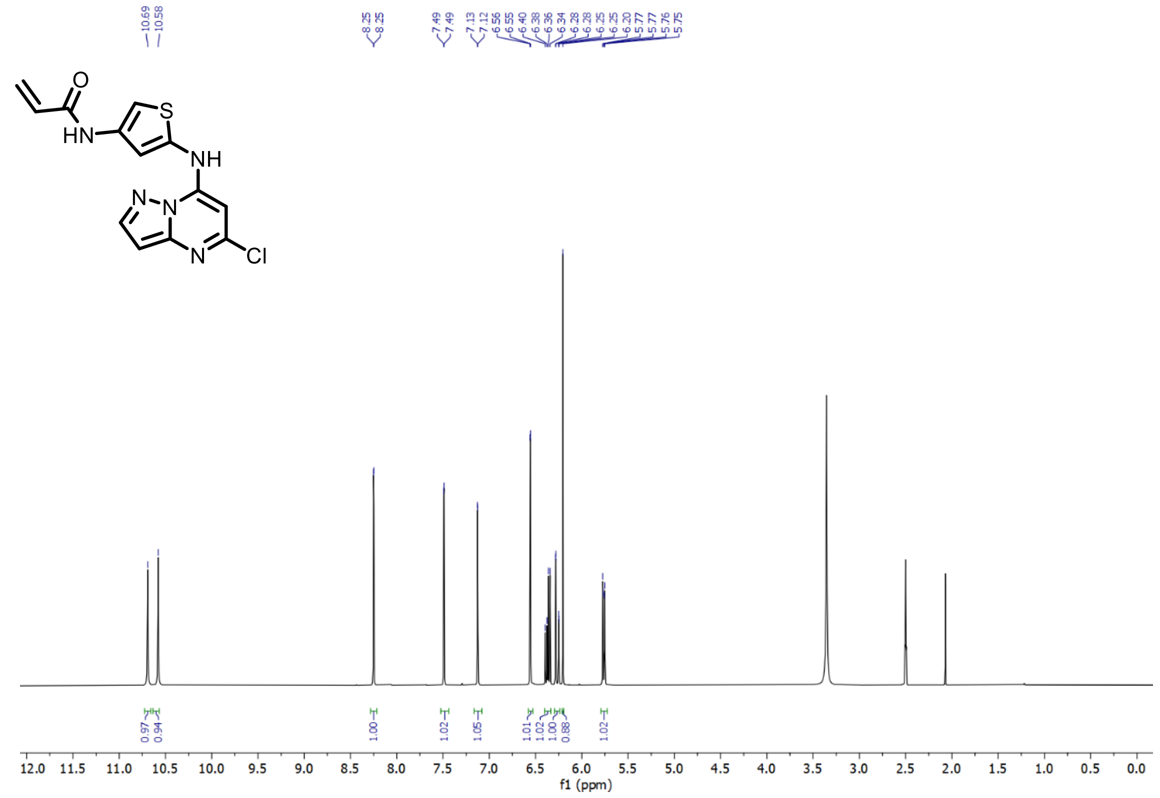

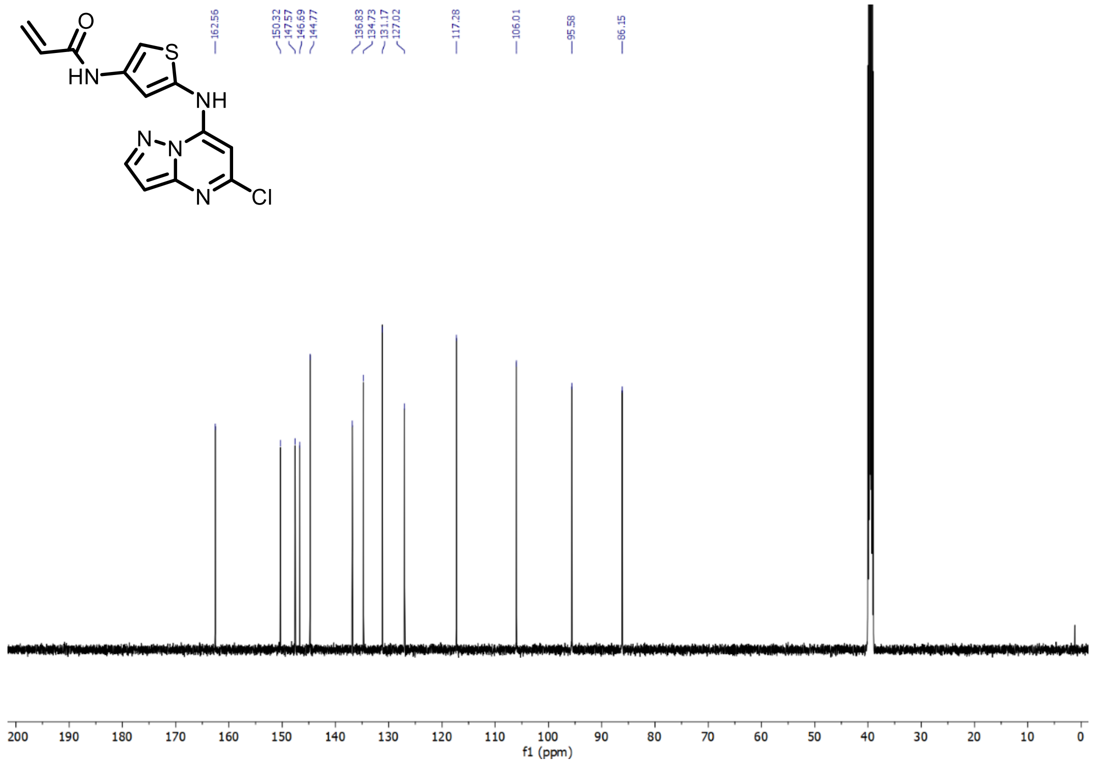

**Supplementary Figure 18.** $^{1}H$ and $^{13}C$ NMR of N-(5-((5-chloropyrazolo[1,5-a]pyrimidin-7-yl)amino)thiophen-3-yl)acrylamide – SCA10

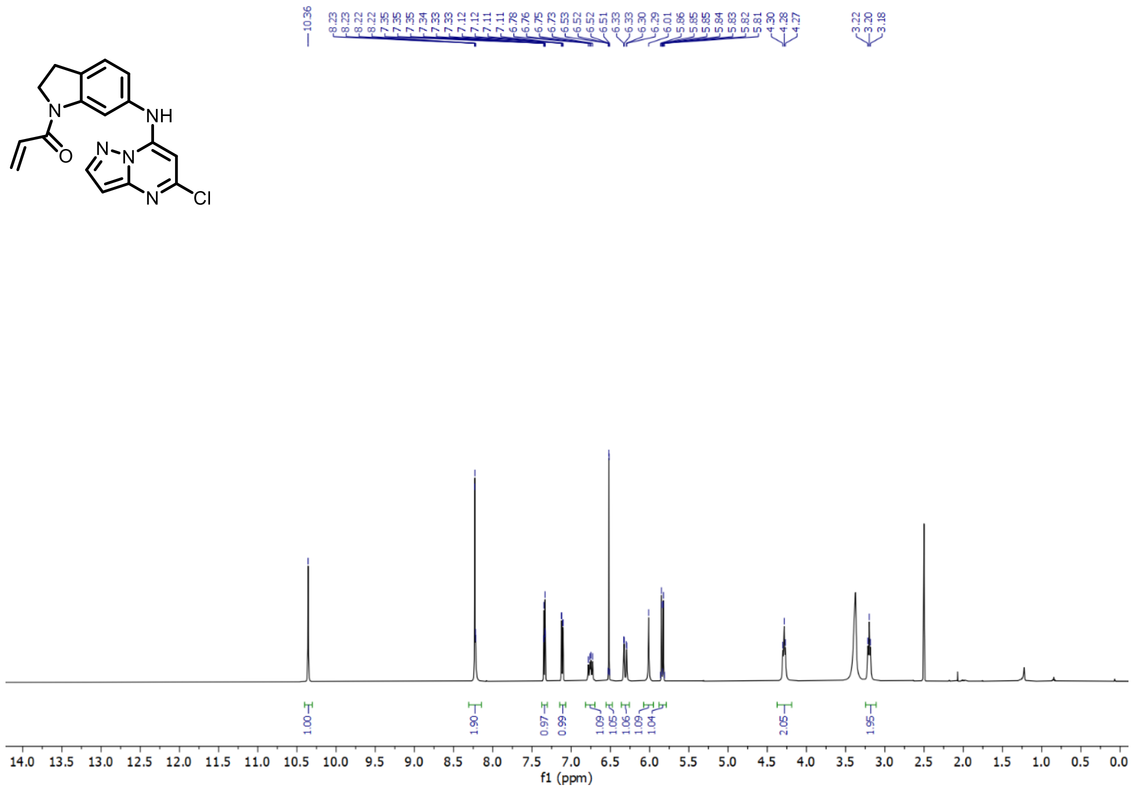

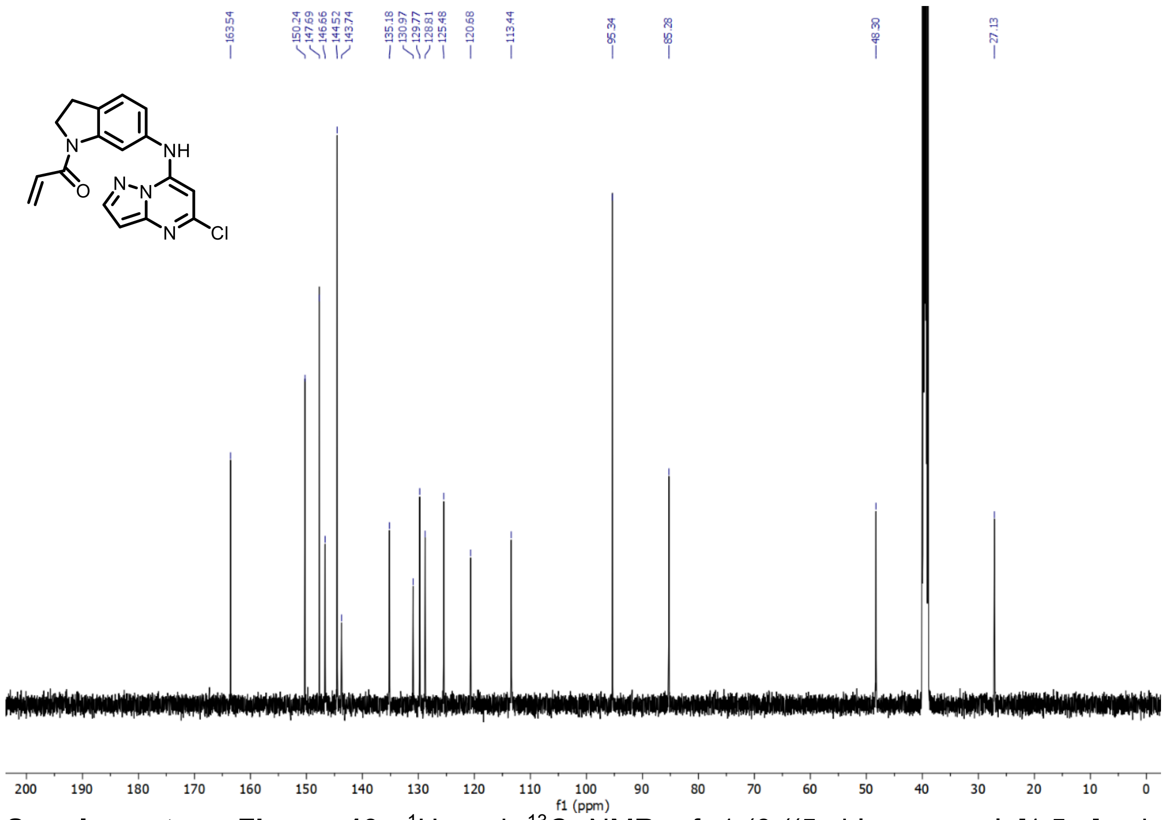

**Supplementary Figure 19.** $^{1}H$ and $^{13}C$ NMR of 1-(6-((5-chloropyrazolo[1,5-a]pyrimidin-7-yl)amino)indolin-1-yl)prop-2-en-1-one – SCA11

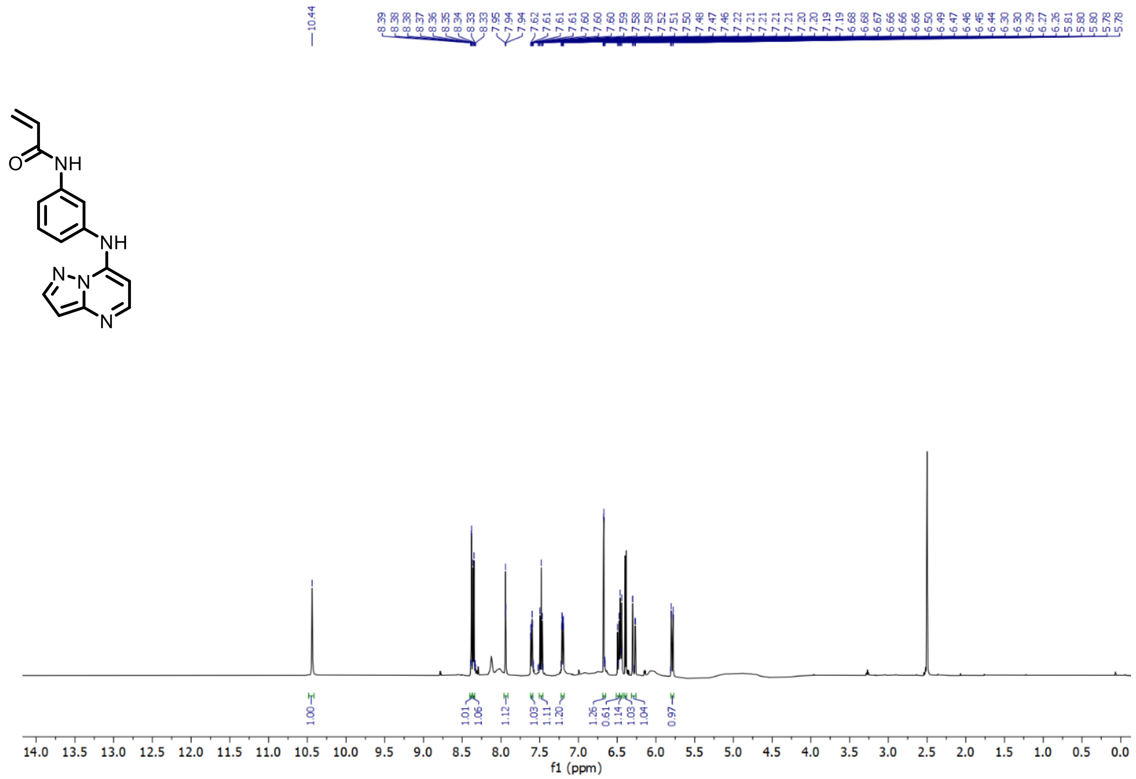

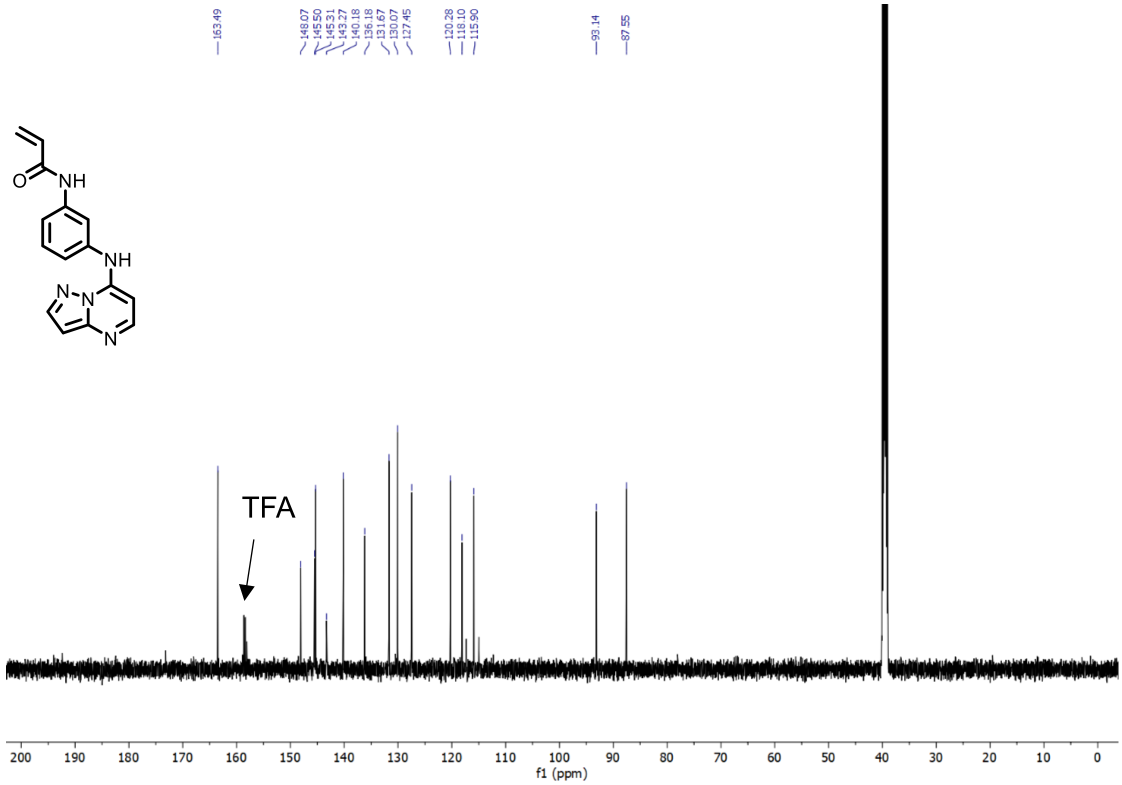

**Supplementary Figure 20.** $^{1}H$ and $^{13}C$ NMR of N-(3-(pyrazolo[1,5-a]pyrimidin-7-ylamino)phenyl)acrylamide – SCA12 (TFA salt)

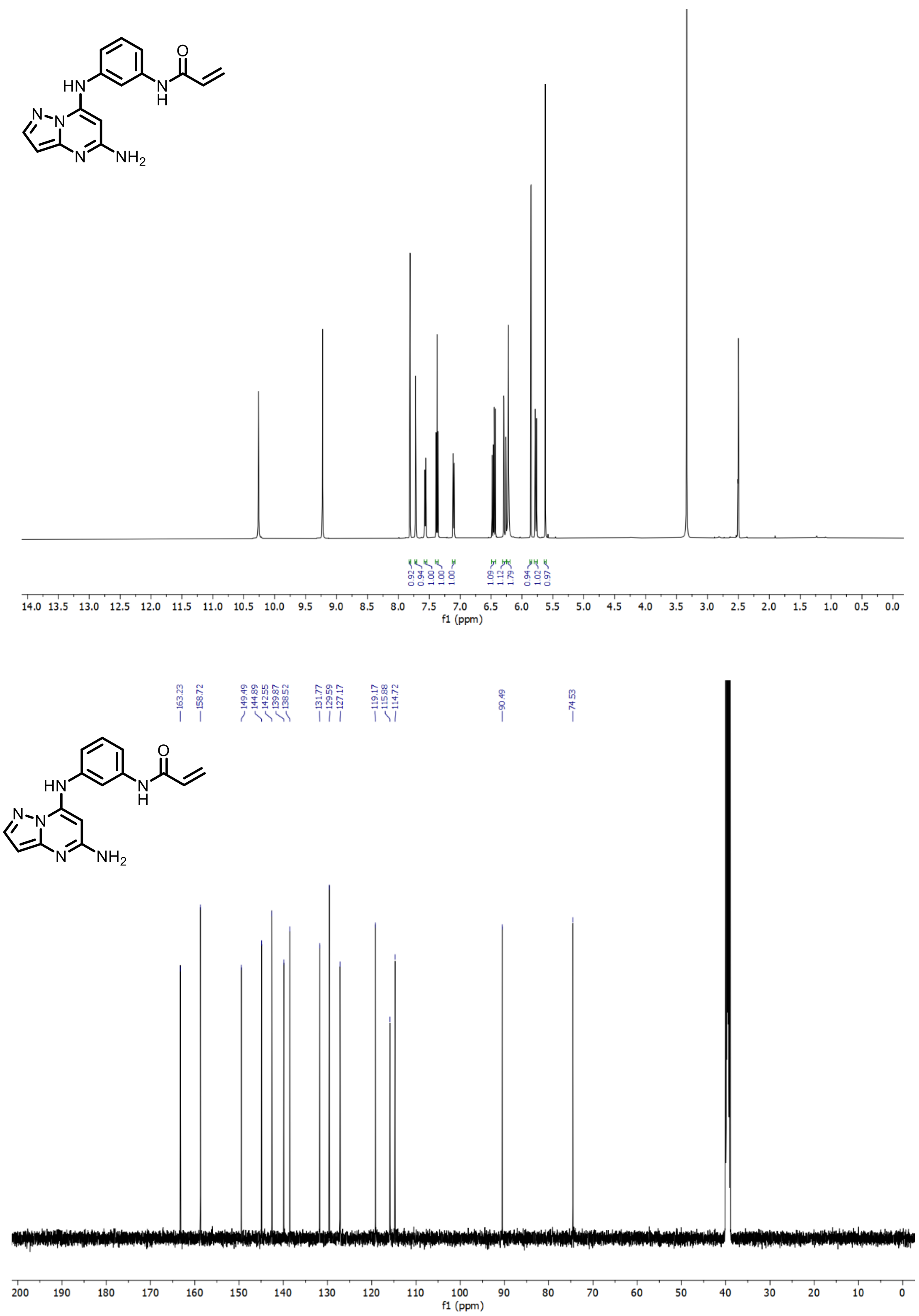

**Supplementary Figure 21.** $^{1}H$ and $^{13}C$ NMR of N-(3-(5-aminopyrazolo[1,5-a]pyrimidin-7-ylamino)phenyl)acrylamide – SCA13

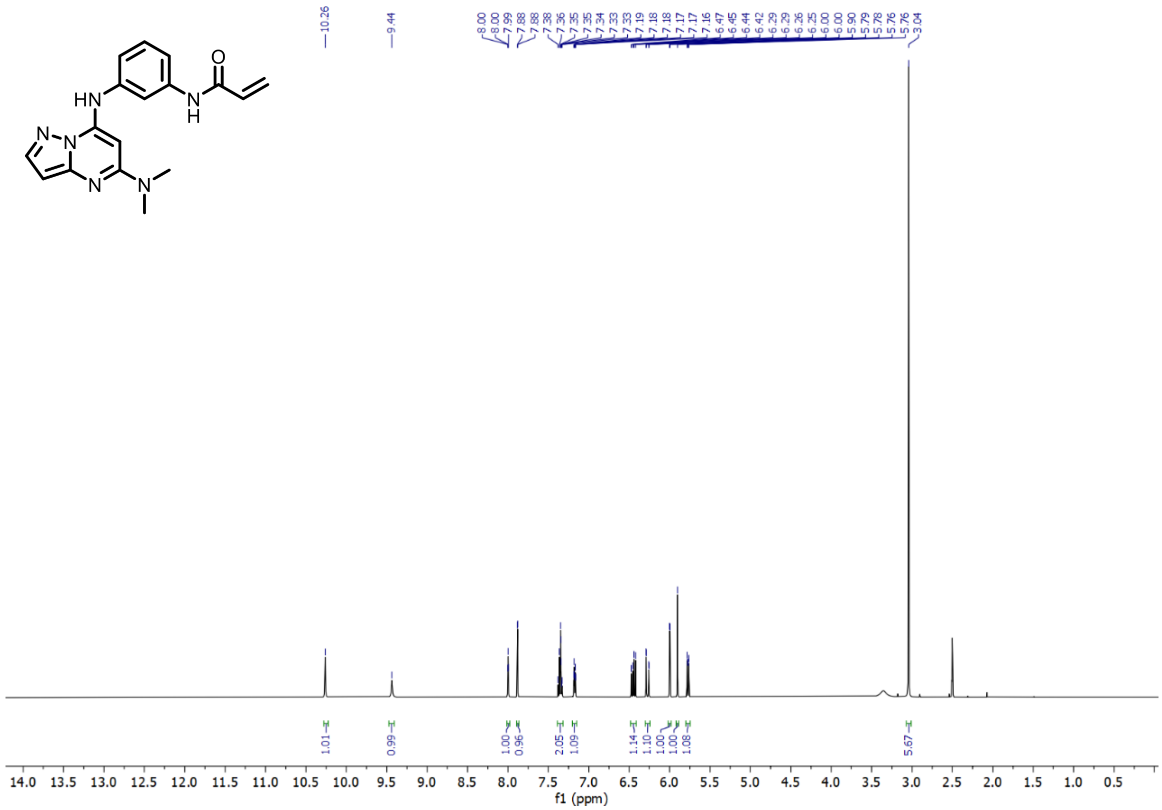

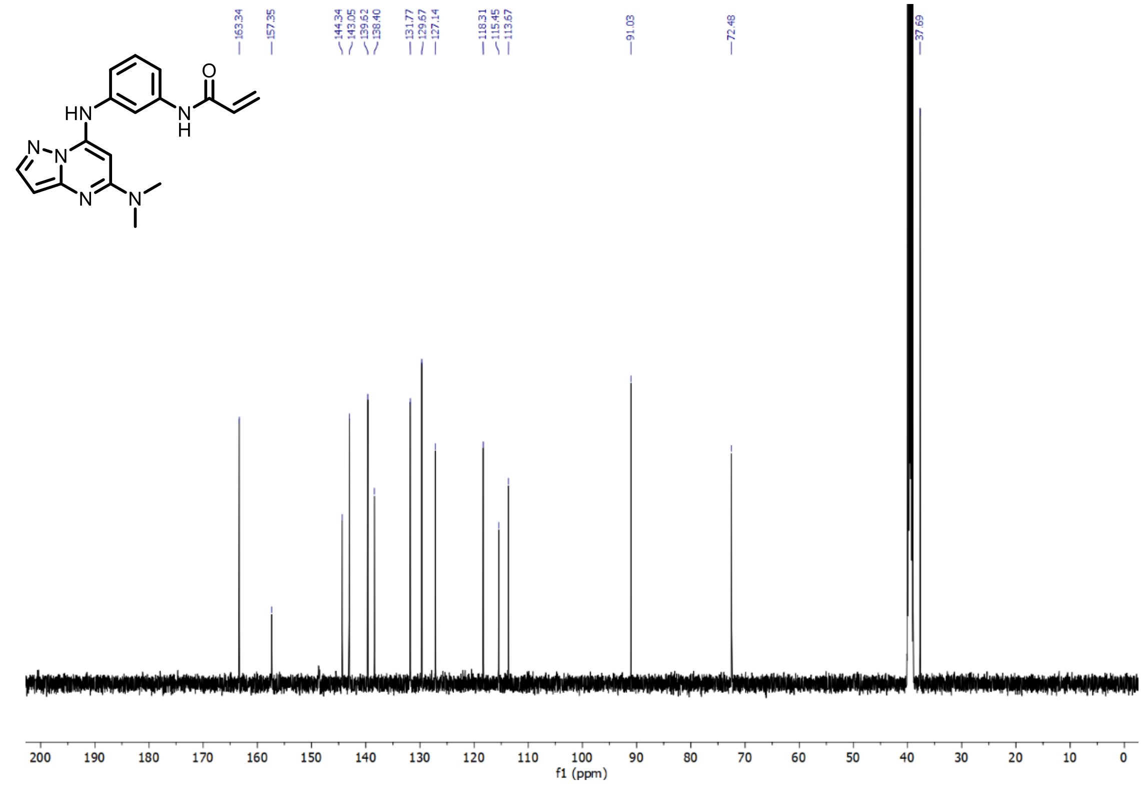

**Supplementary Figure 22.** $^{1}$H and $^{13}$C NMR N-(3-(5-(dimethylamino)pyrazolo[1,5-a]pyrimidin-7-ylamino)phenyl)acrylamide – SCA14

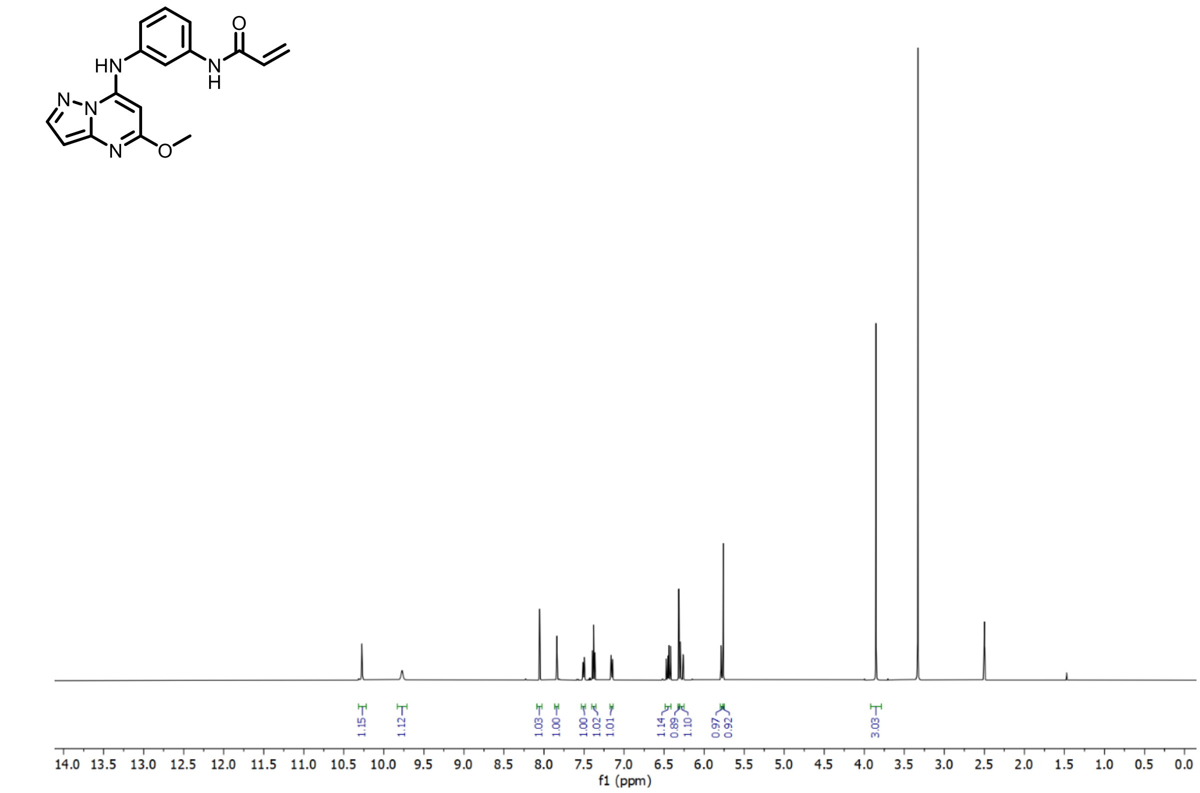

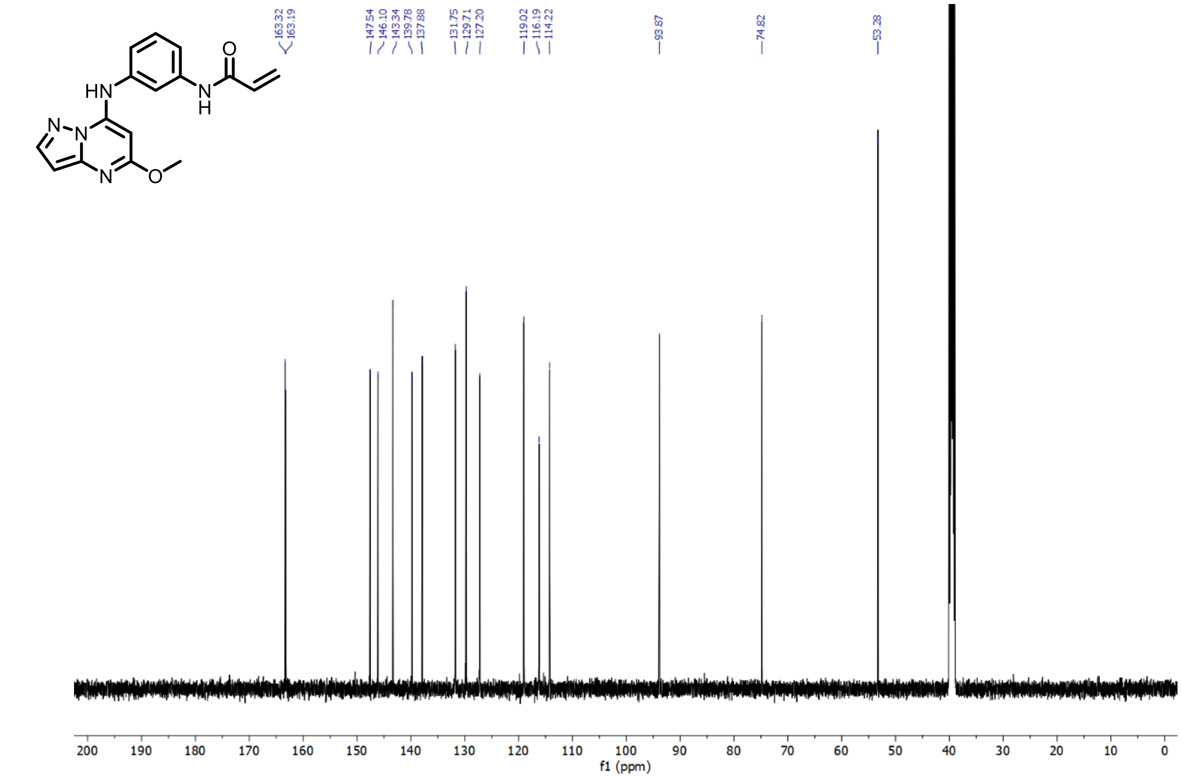

**Supplementary Figure 23.** $^{1}H$ and $^{13}C$ NMR 7 N-(3-(5-methoxypyrazolo[1,5-a]pyrimidin-7-ylamino)phenyl)acrylamide – SCA15

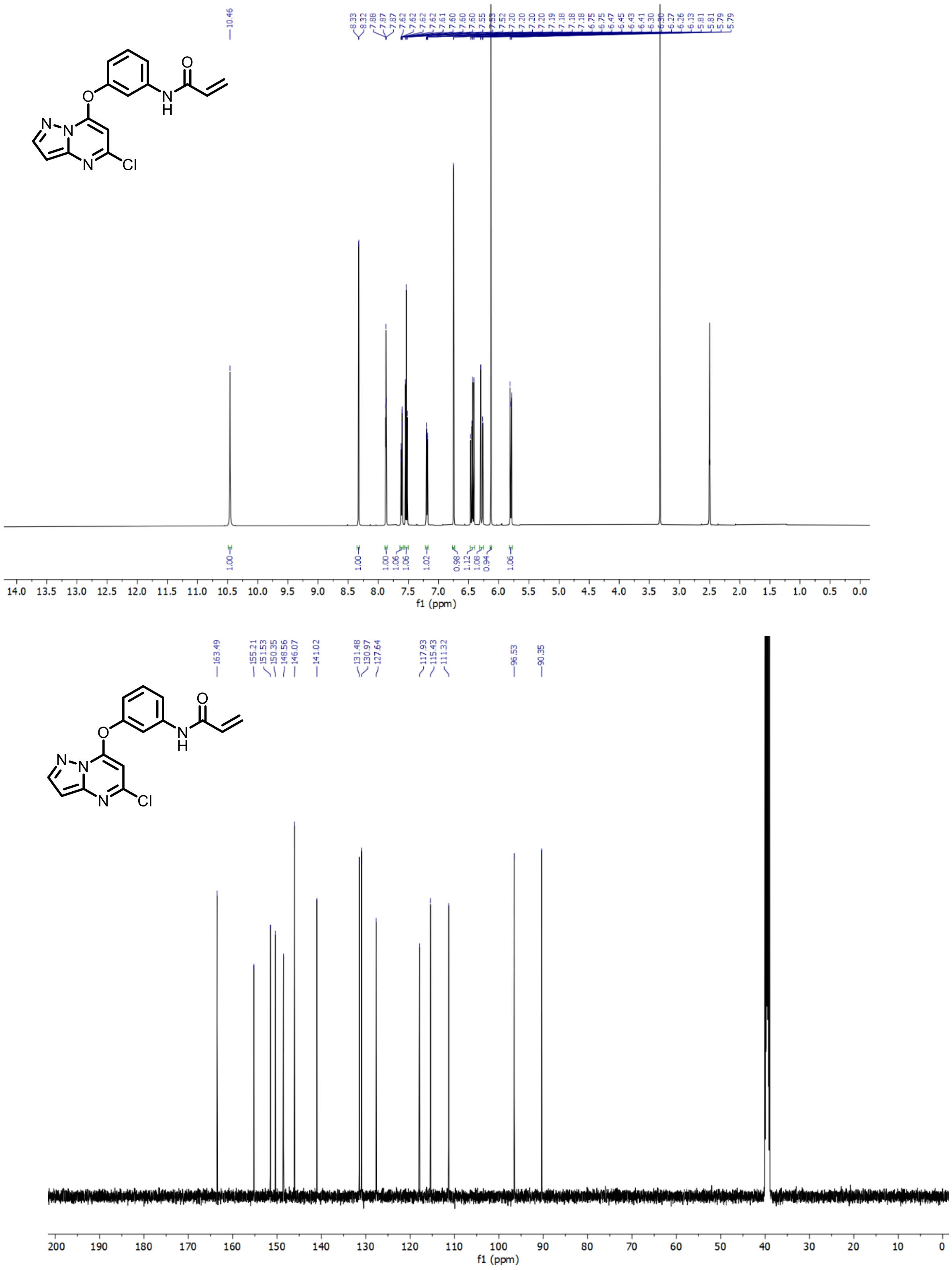

**Supplementary Figure 24.** $^{1}$H and $^{13}$C NMR (N-(3-(5-chloropyrazolo[1,5-a]pyrimidin-7-yloxy)phenyl)acrylamide – SCA16

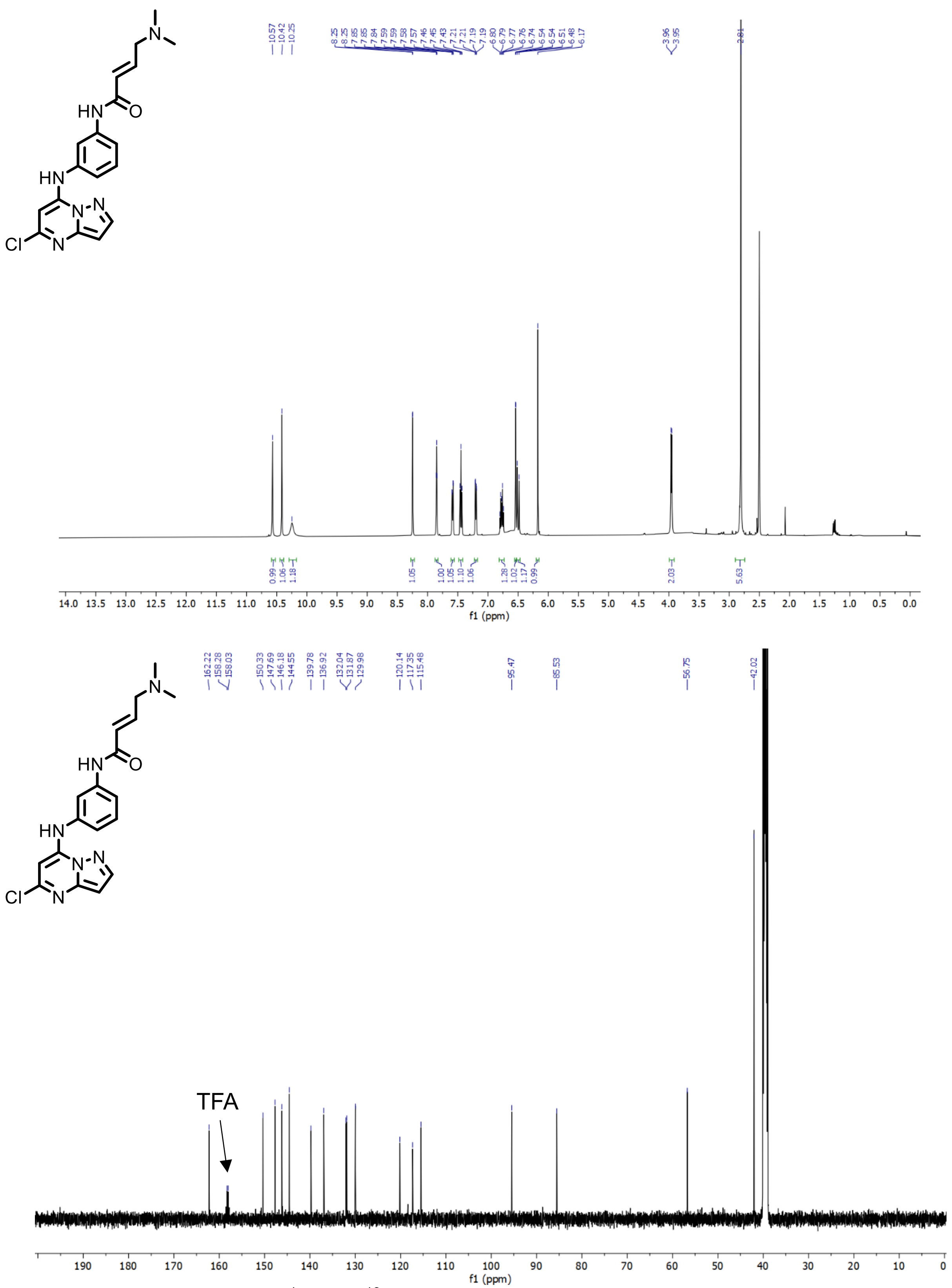

**Supplementary Figure 25.** $^{1}H$ and $^{13}C$ NMR of (E)-N-(3-((5-chloropyrazolo[1,5-a]pyrimidin-7-yl)amino)phenyl)-4-(dimethylamino)but-2-enamide – SCA17 (TFA Salt)

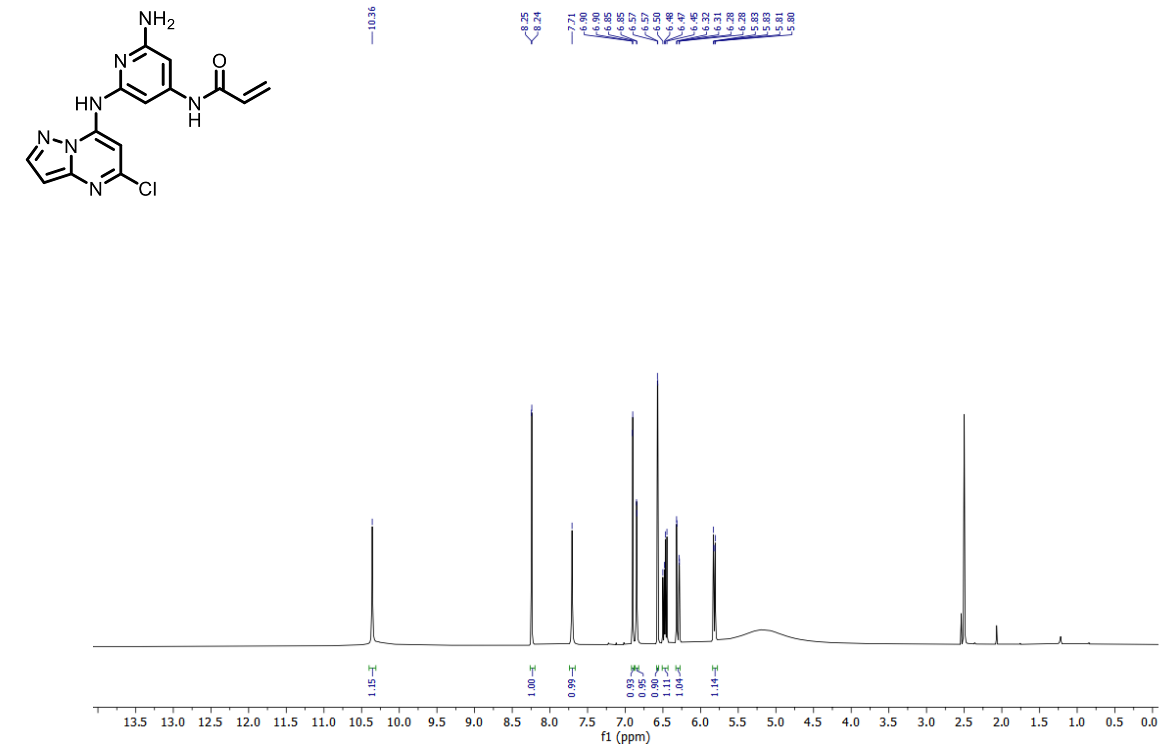

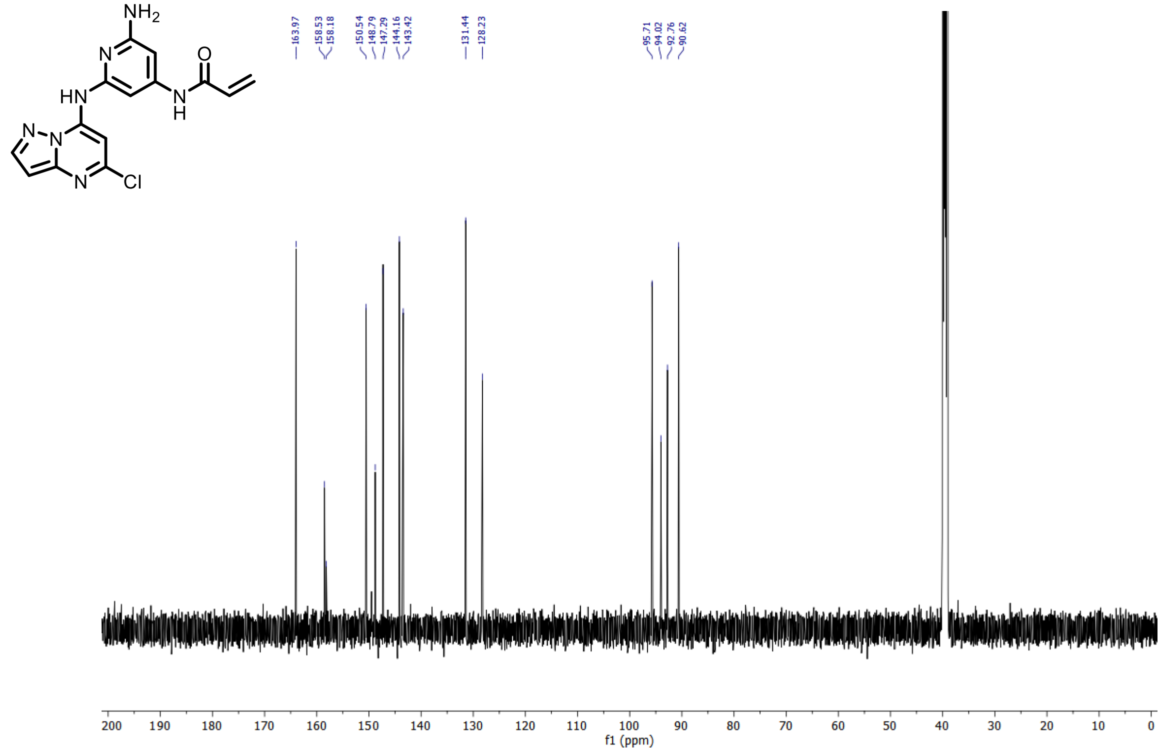

**Supplementary Figure 26.** $^1$H and $^{13}$C NMR of N-(2-amino-6-((5-chloropyrazolo[1,5-a]pyrimidin-7-yl)amino)pyridin-4-yl)acrylamide – SCA18

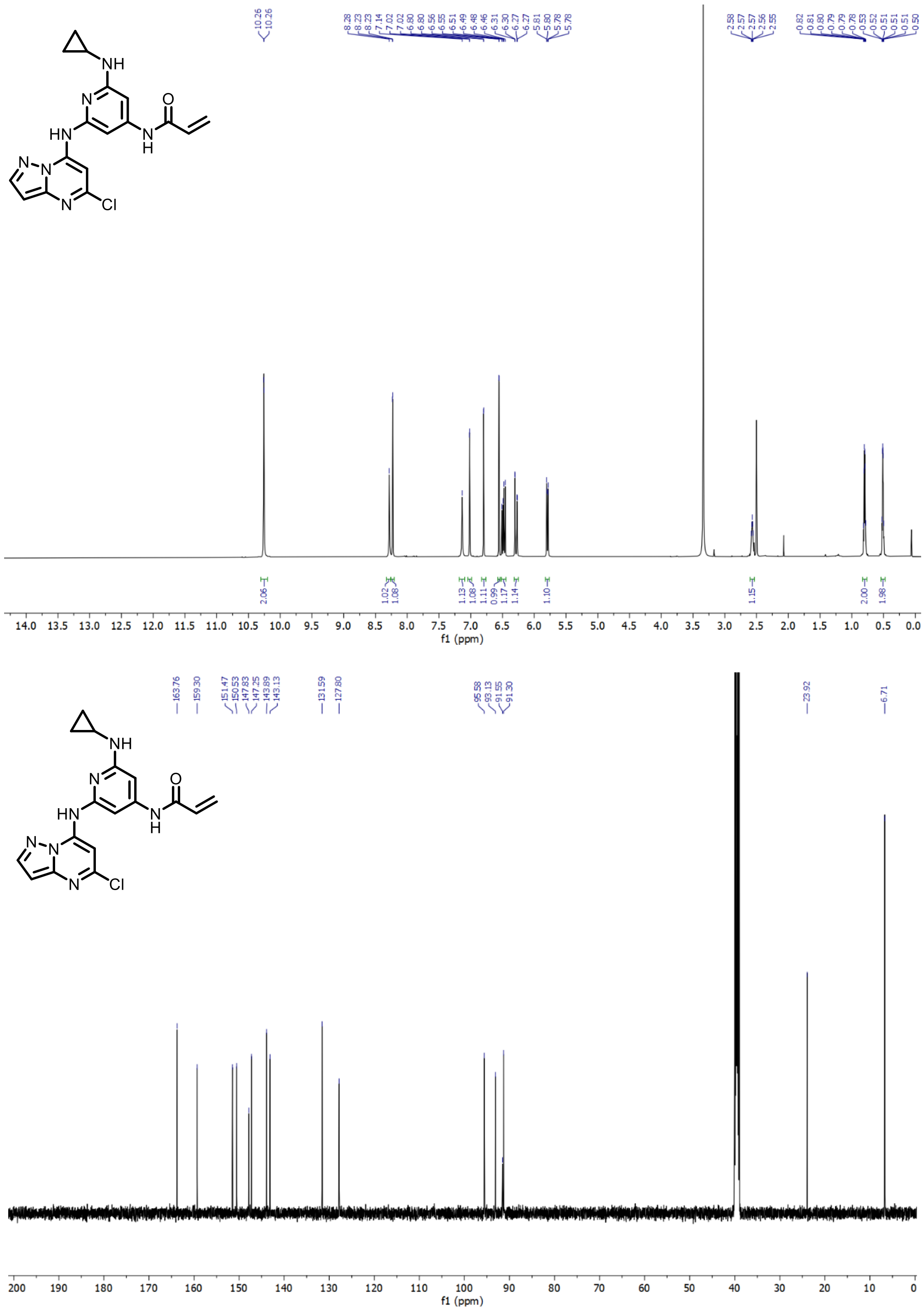

**Supplementary Figure 27.** $^{1}H$ and $^{13}C$ NMR of N-(2-((5-chloropyrazolo[1,5-a]pyrimidin-7-yl)amino)-6-(cyclopropylamino)pyridin-4-yl)acrylamide – SCA19

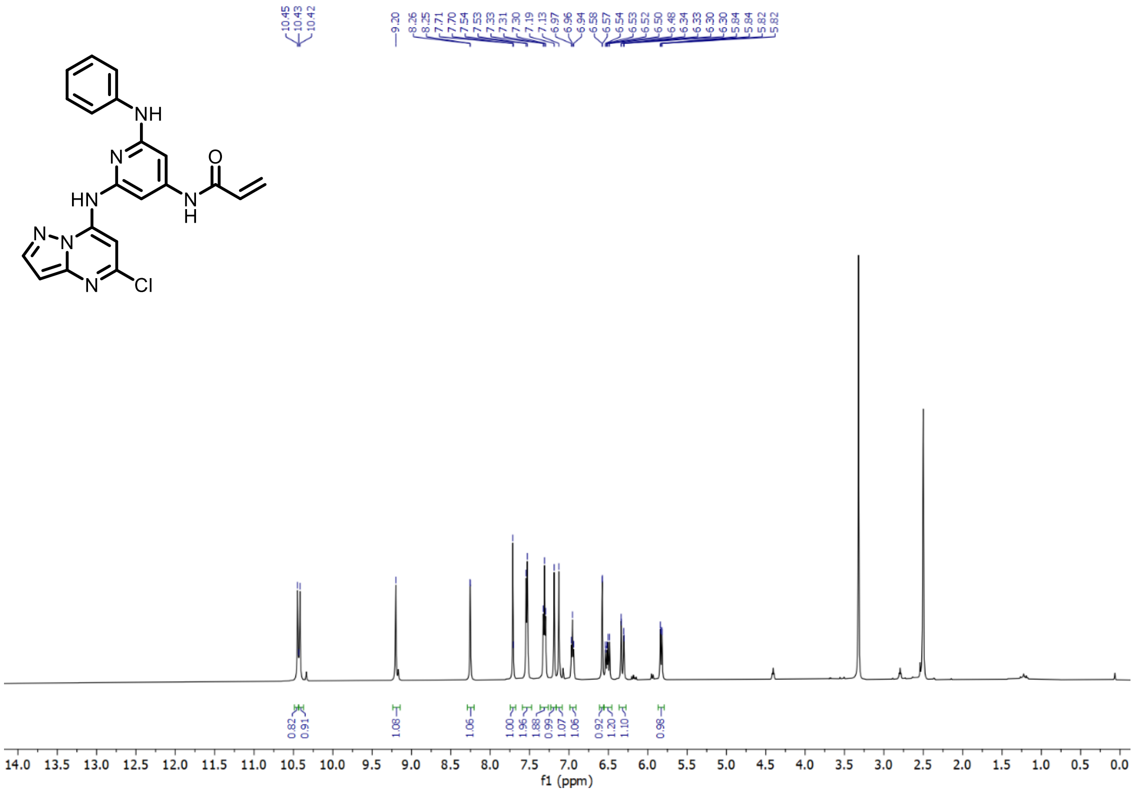

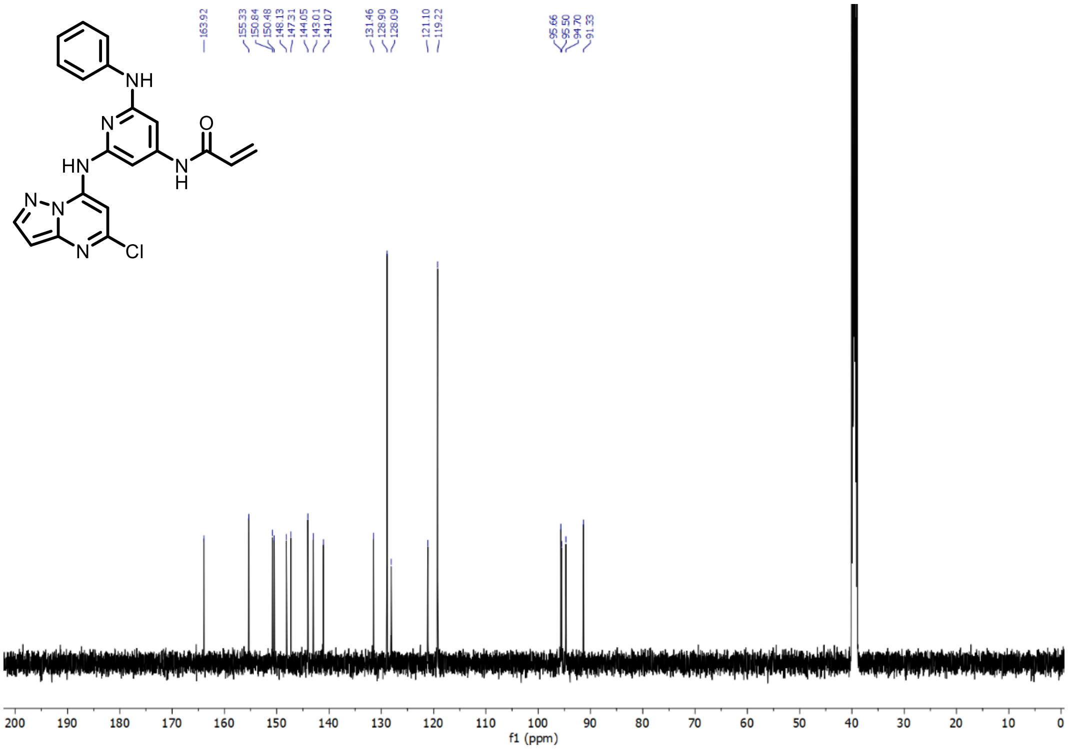

**Supplementary Figure 28.** $^{1}H$ and $^{13}C$ NMR of N-(2-((5-chloropyrazolo[1,5-a]pyrimidin-7-yl)amino)-6-(phenylamino)pyridin-4-yl)acrylamide – SCA20

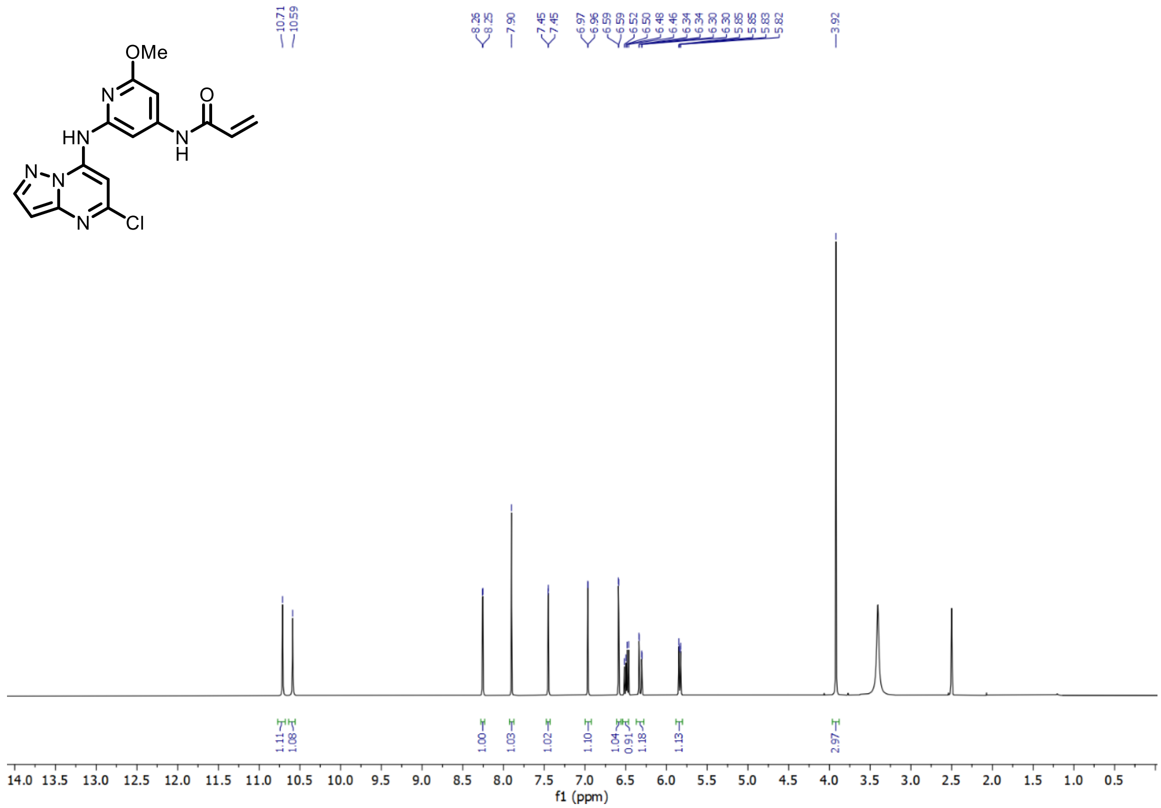

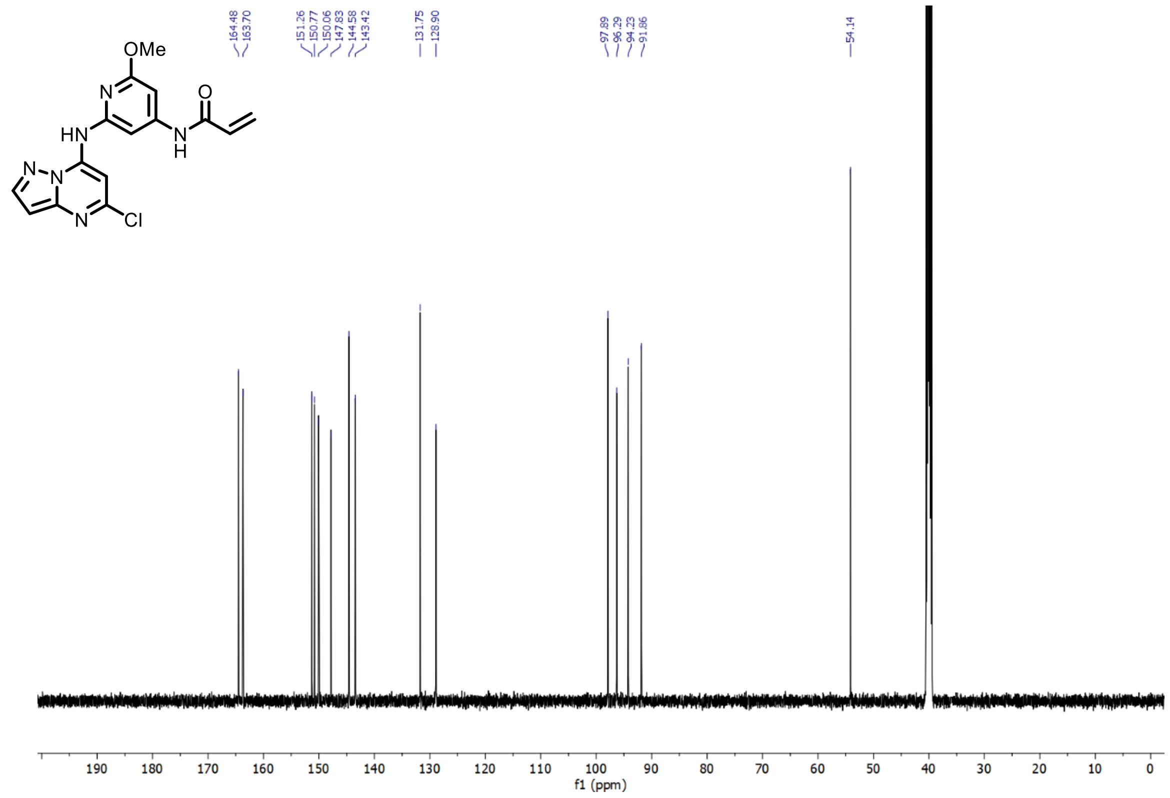

**Supplementary Figure 29.** $^{1}$H and $^{13}$C NMR of N-(2-((5-chloropyrazolo[1,5-a]pyrimidin-7-yl)amino)-6-methoxypyridin-4-yl)acrylamide – SCA21

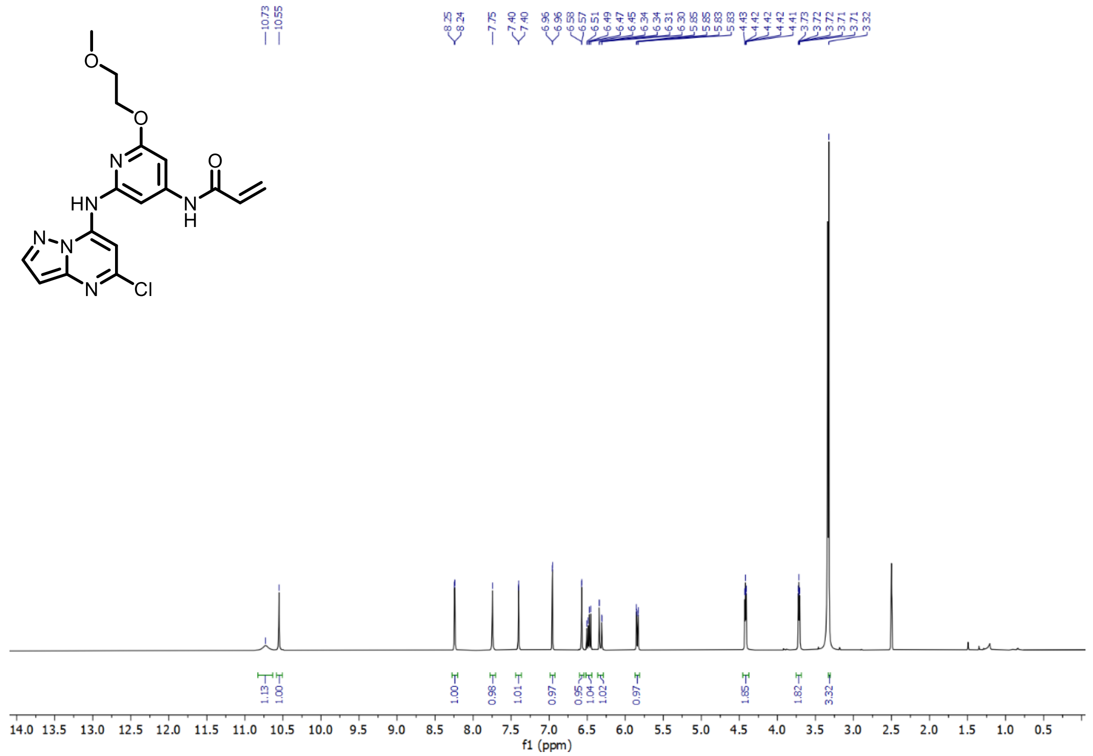

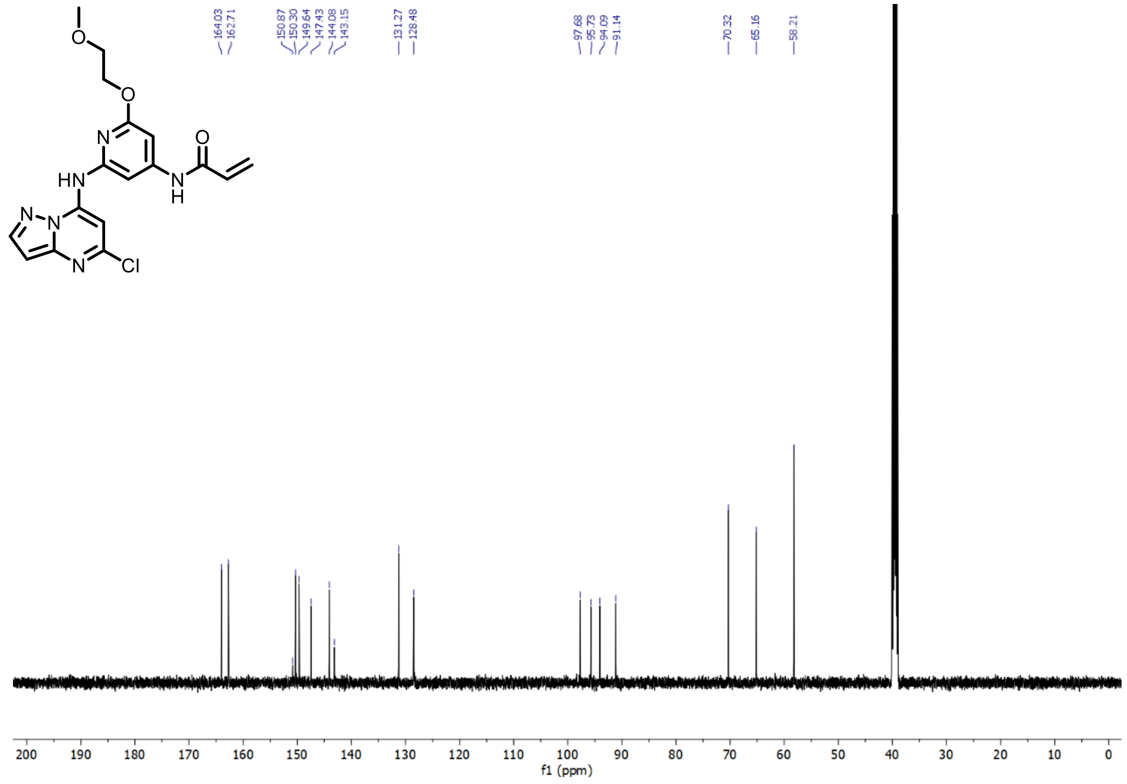

**Supplementary Figure 30.** $^{1}H$ and $^{13}C$ NMR of N-(2-((5-chloropyrazolo[1,5-a]pyrimidin-7-yl)amino)-6-(2-methoxyethoxy)pyridin-4-yl)acrylamide – SCA22

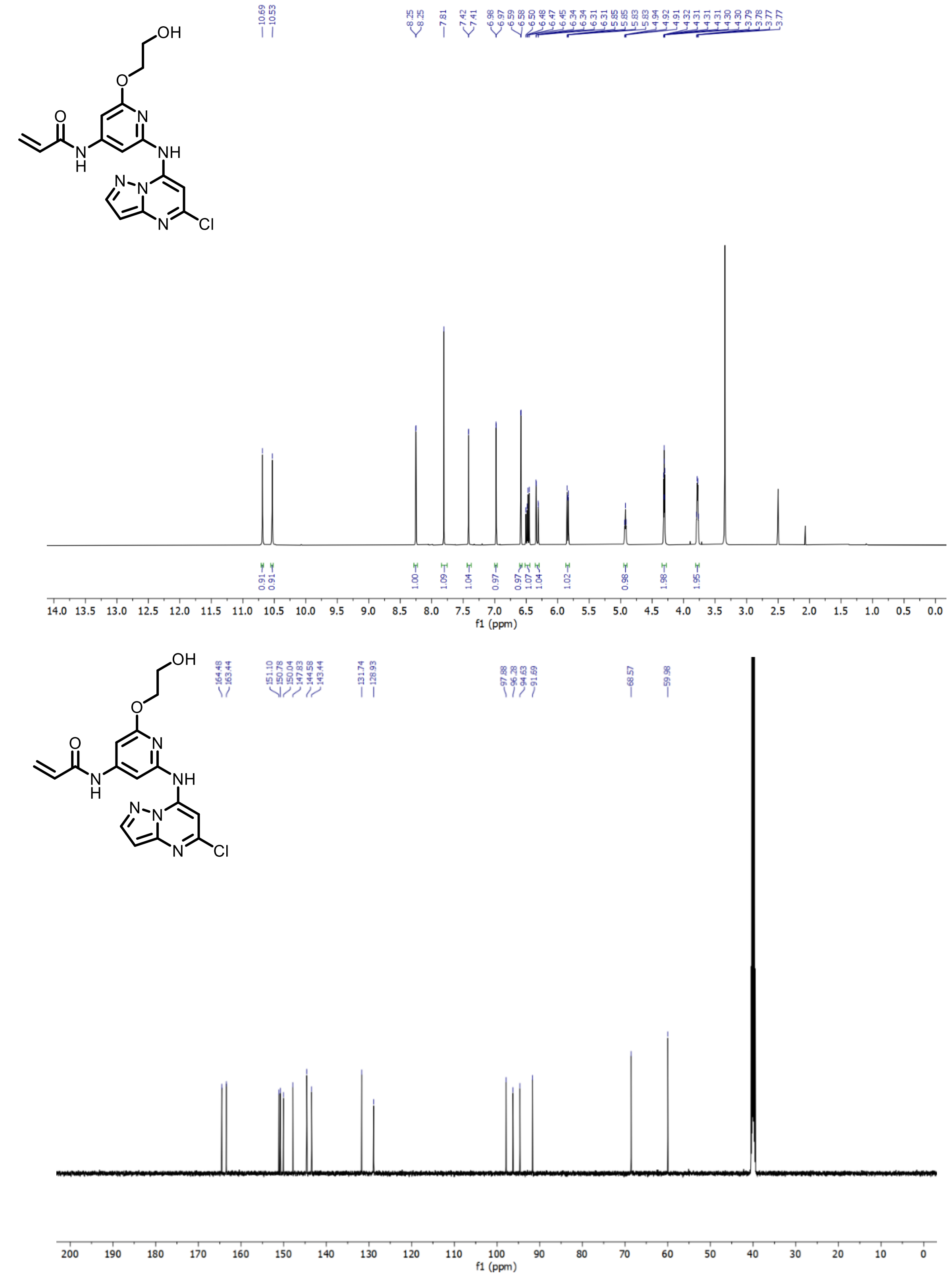

**Supplementary Figure 31.** $^{1}H$ and $^{13}C$ NMR of N-(2-((5-chloropyrazolo[1,5-a]pyrimidin-7-yl)amino)-6-(2-hydroxyethoxy)pyridin-4-yl)acrylamide – SCA23

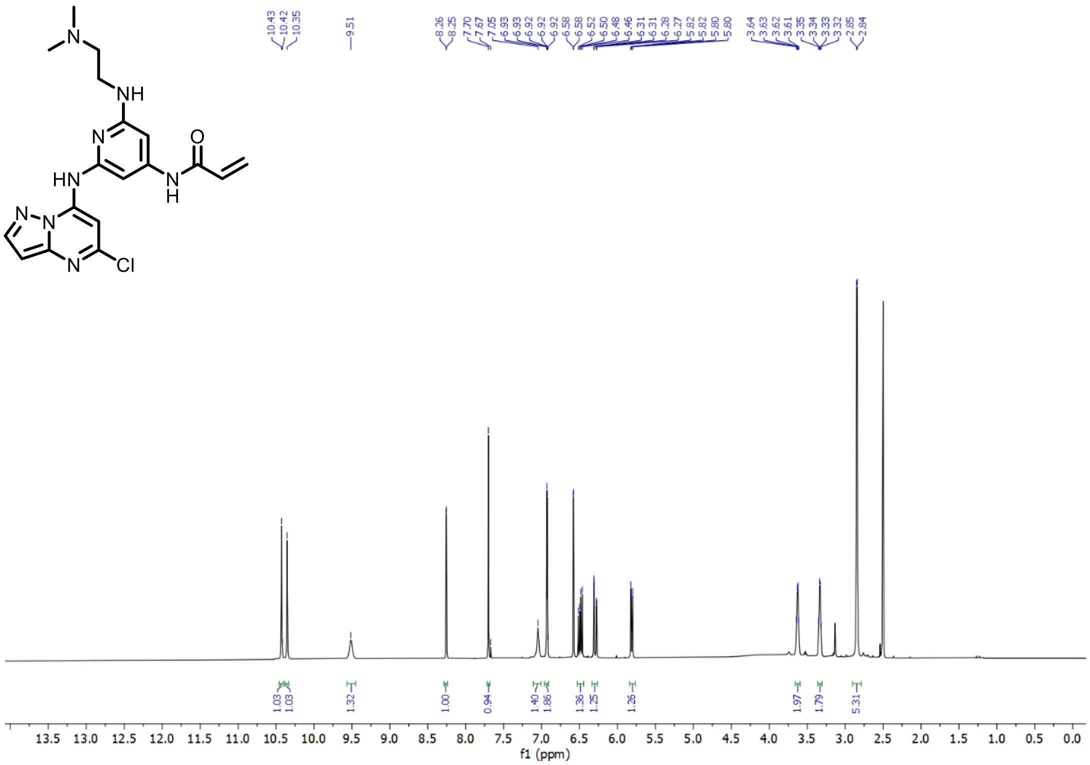

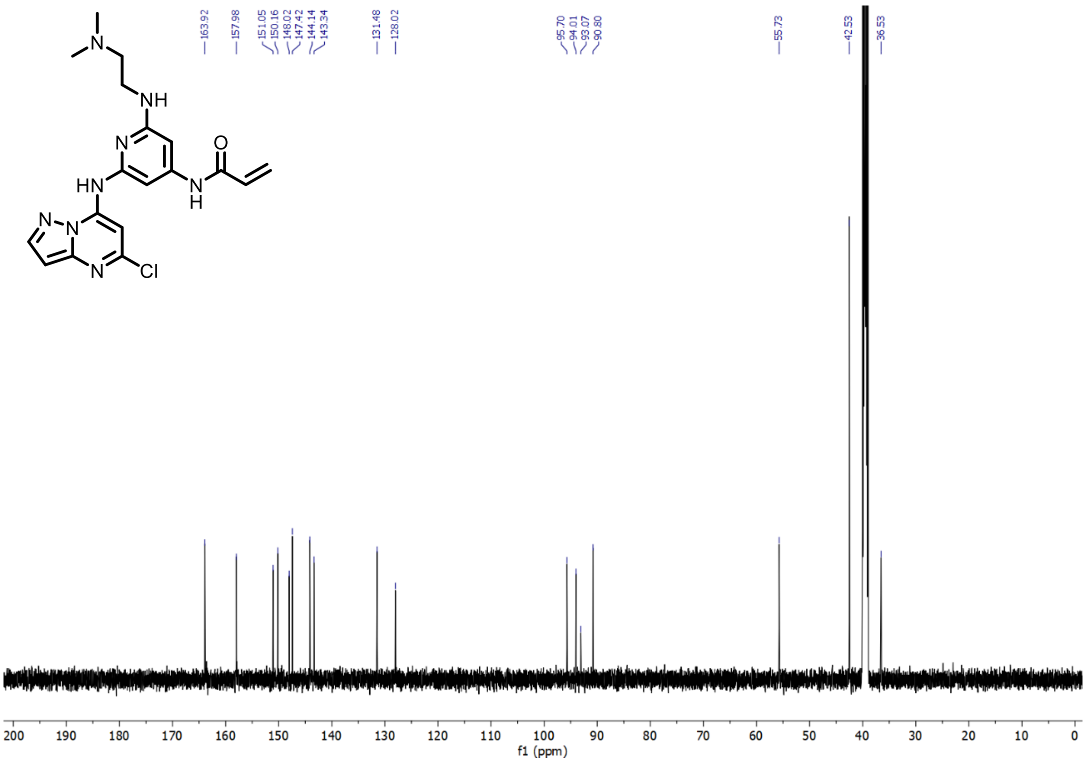

**Supplementary Figure 32.** $^{1}H$ and $^{13}C$ NMR of N-(2-((5-chloropyrazolo[1,5-a]pyrimidin-7-yl)amino)-6-((2-(dimethylamino)ethyl)amino)pyridin-4-yl)acrylamide – SCA24

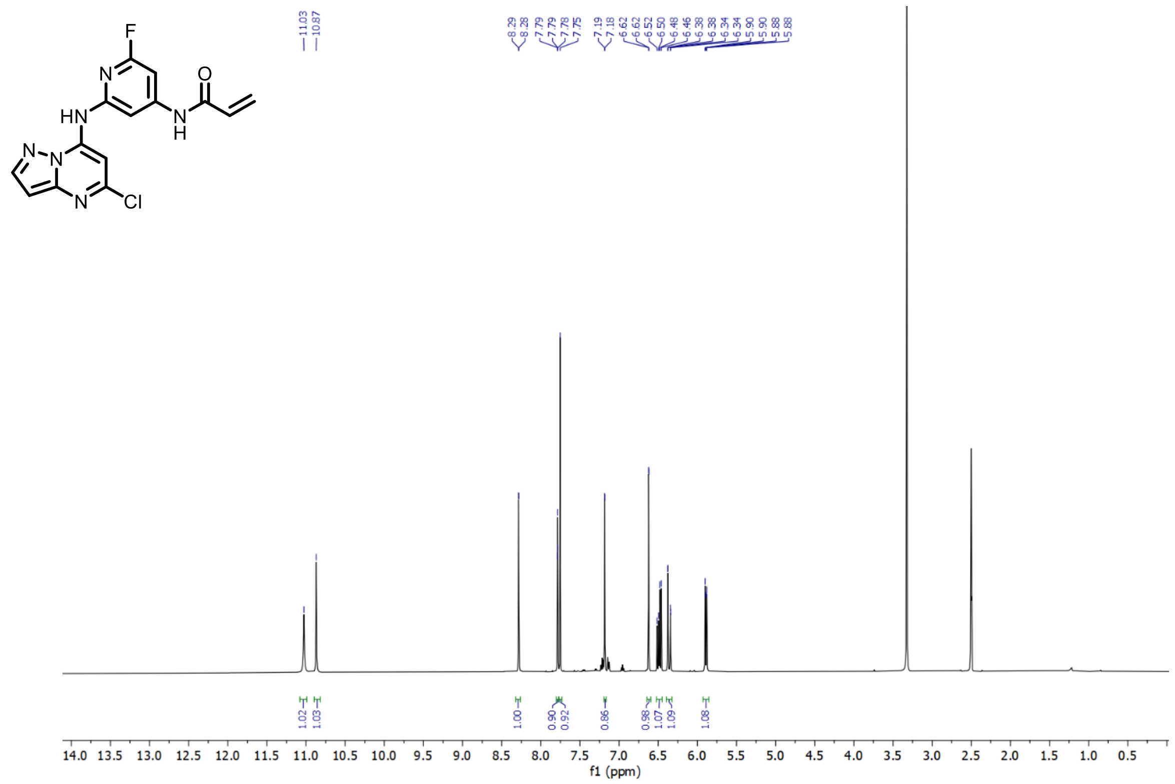
F
N
O
HN
N
H
N
N
N
Cl
11.03
10.87
8.29
8.28
7.79
7.79
7.78
7.75
7.19
7.18
6.62
6.62
6.52
6.50
6.48
6.46
6.38
6.38
6.34
6.34
5.90
5.90
5.88
5.88
1.02
1.03
1.00
0.90
0.92
0.86
0.98
1.07
1.09
1.08
14.0 13.5 13.0 12.5 12.0 11.5 11.0 10.5 10.0 9.5 9.0 8.5 8.0 7.5 7.0 6.5 6.0 5.5 5.0 4.5 4.0 3.5 3.0 2.5 2.0 1.5 1.0 0.5
f1 (ppm)

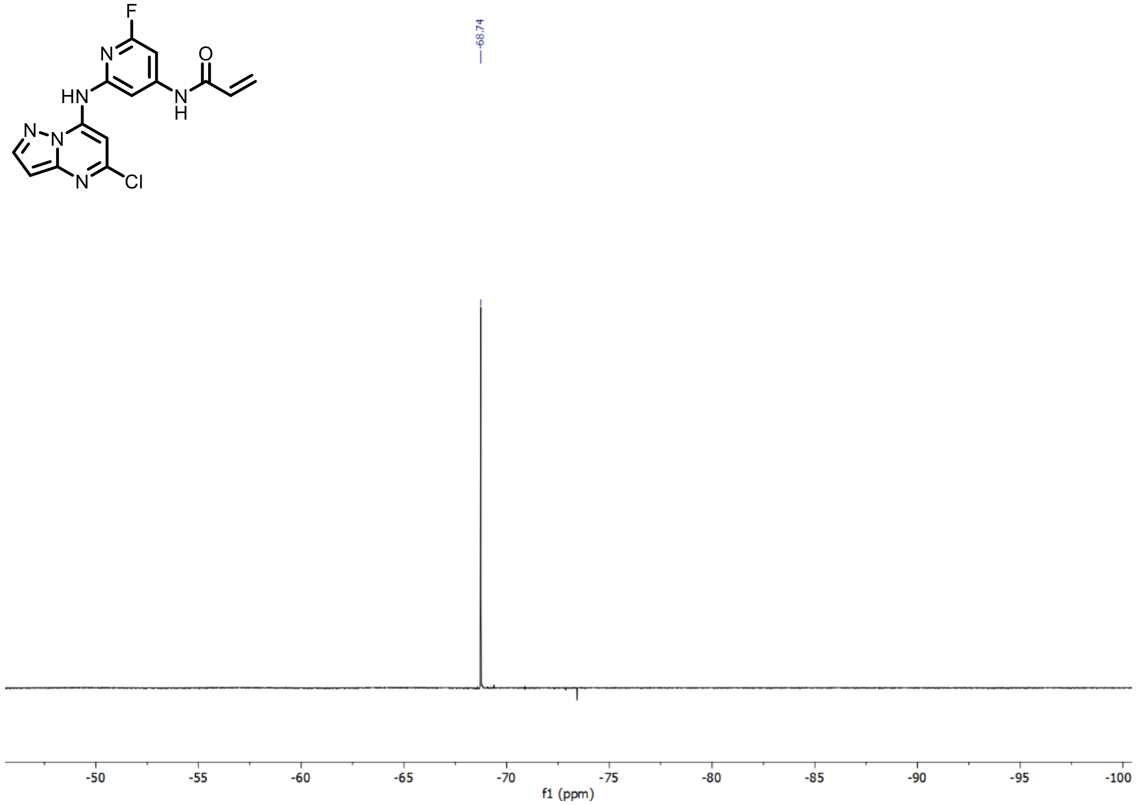
F
N
O
HN
N
H
N
N
N
Cl
-68.74
-50 -55 -60 -65 -70 -75 -80 -85 -90 -95 -100
f1 (ppm)

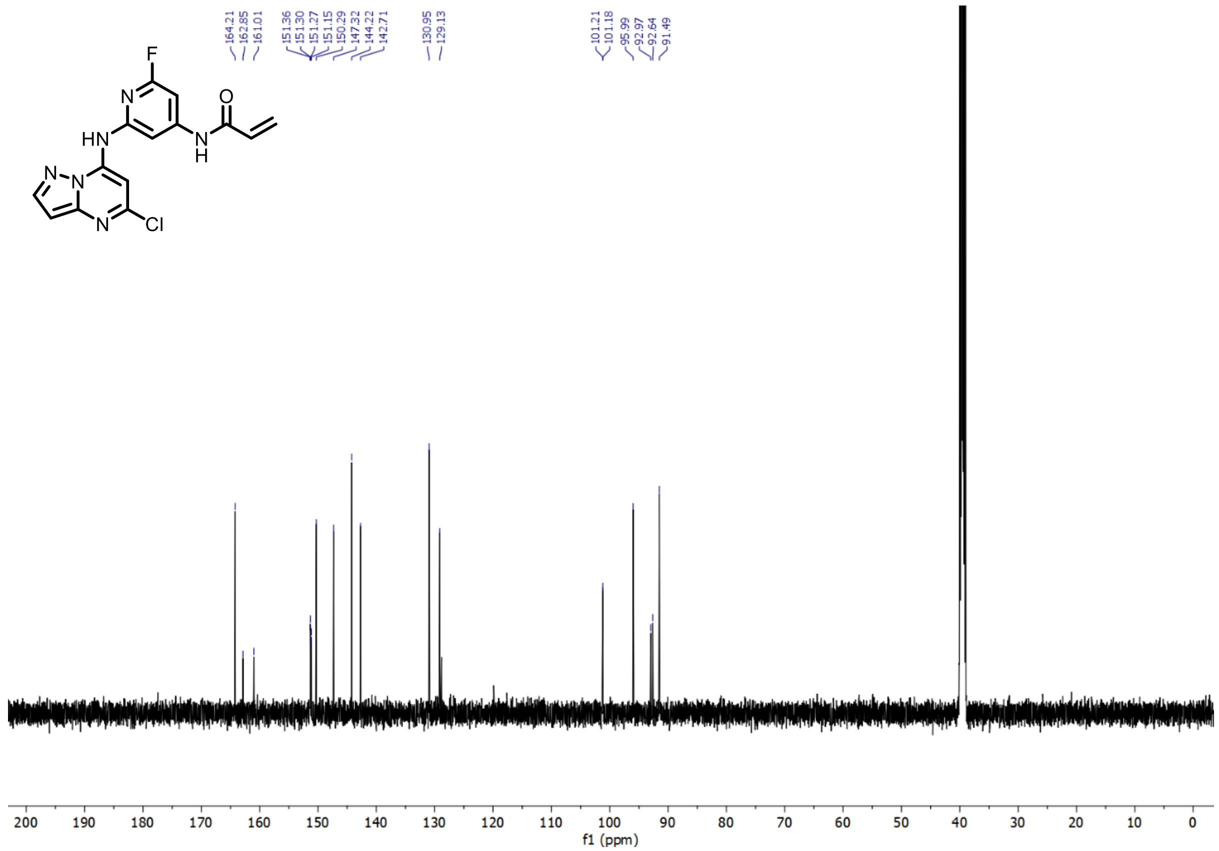

**Supplementary Figure 33.** $^{1}H$, $^{19}F$, and $^{13}C$ NMR of N-(2-((5-chloropyrazolo[1,5-a]pyrimidin-7-yl)amino)-6-fluoropyridin-4-yl)acrylamide – SCA25

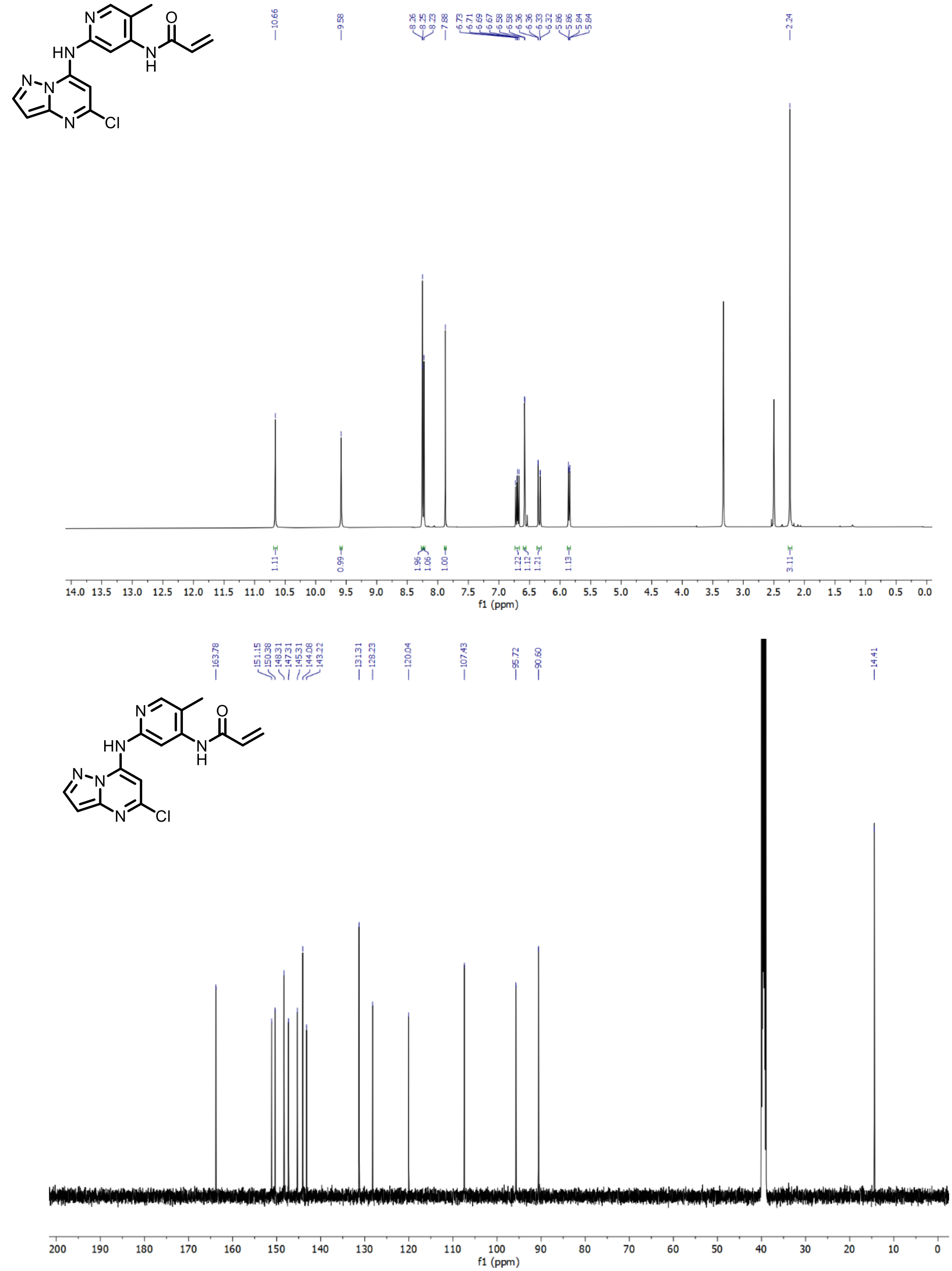

**Supplementary Figure 34.** $^{1}H$ and $^{13}C$ NMR of N-(2-((5-chloropyrazolo[1,5-a]pyrimidin-7-yl)amino)-5-methylpyridin-4-yl)acrylamide – SCA26

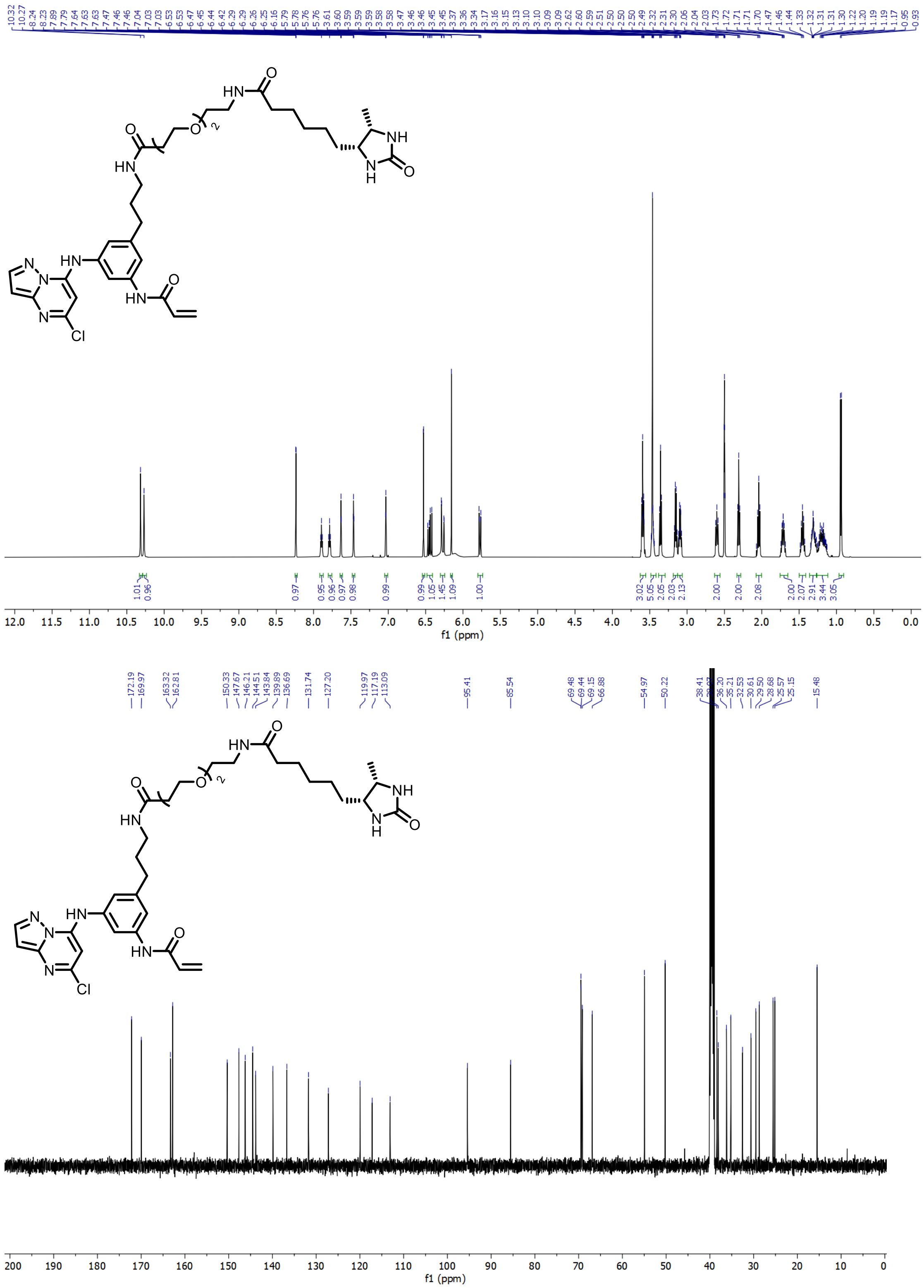

**Supplementary Figure 35.** $^1$H and $^{13}$C NMR of (N-(2-(2-(3-((3-(3-acrylamido-5-((5-chloropyrazolo[1,5-a]pyrimidin-7-yl)amino)phenyl)propyl)amino)-3-oxopropoxy)ethoxy)ethyl)-6-((4R,5S)-5-methyl-2-oxoimidazolidin-4-yl)hexanamide – Desthiobiotinylated SCA1
